# Identifying Combinations of Cancer Drivers in Individual Patients

**DOI:** 10.1101/674234

**Authors:** Michael I. Klein, Vincent L. Cannataro, Jeffrey P. Townsend, David F. Stern, Hongyu Zhao

**Author notes:** Equal contributors. Correspondences.

## Abstract

Identifying the subset of genetic alterations present in individual tumors that are essential and collectively sufficient for cancer initiation and progression would advance the development of effective personalized treatments. We present CRSO for inferring the combinations of alterations, i.e., rules, that cooperate to drive tumor formation in individual patients. CRSO prioritizes rules by integrating patient-specific passenger probabilities for individual alterations along with information about the recurrence of particular combinations throughout the population. We present examples in glioma, liver cancer and melanoma of significant differences in patient outcomes based on rule assignments that are not identifiable by consideration of individual alterations.

## 1 Background

Genome sequencing of patient tumors has revealed that most tumors harbor many genetic alterations—most abundantly, somatic mutations and somatic copy number variations (SCNVs). Large-scale datasets produced by cancer genomics projects such as The Cancer Genome Atlas (TCGA) [1, 2] and the International Cancer Genome Consortium (ICGC) [3] have further revealed tremendous inter-tumoral heterogeneity within all cancer types, motivating efforts to personalize treatment of individual patients based on patient-specific genetic data. Knowledge of the specific alterations responsible for cancer initiation and progression, i.e., *driver alterations*, in individual patients would be of great benefit to oncologists and precision medicine tumor boards. This information would help clinicians optimally decide the treatment course for a given patient, as well as help researchers to identify novel drug targets for single or combination treatments. Even if an identified driver is not currently druggable, such as the protein product of activating *KRAS* mutations in many cancers [4], knowledge of the full set of tumor-sustaining alterations could identify other vulnerabilities in the tumor that are druggable.

Identification of patient-specific driver alterations is obfuscated because the vast majority of alterations in each tumor are *passengers* that do not contribute to cancer formation. Many computational methods have greatly advanced our ability to distinguish drivers from passengers at the population level, including MutSigCV [5] and dNdScv [6] for identification of significantly mutated genes (SMGs), and GISTIC2 [7] for identification of significant SCNVs. These methods use empirical data to define expectations for the frequencies of specific mutations or SCNVs occurring as passengers, and then identify candidate drivers to be those alterations that are more frequent than are expected by chance. MutSigCV was the first algorithm to take into account sources of heterogeneity in mutation rates such as DNA replication timing, trinucleotide context and gene expression levels [5]. dNdScv assumes that synonymous substitutions occur as a result of neutral evolution and uses the ratio of non-synonymous to synonymous mutation rates (dN/dS), along with covariates that impact mutation rates, to define a null distribution for the expected number of non-synonymous mutations observed in each gene across the tumor population [6]. GISTIC2 identifies candidate driver SCNVs by estimating the background rate of SCNVs and then calculating a score for each region reflecting the likelihood of the observed alteration frequencies under the proposed background model. Regions with scores exceeding a significance threshold are predicted to be driver SCNVs, and peak gene targets are identified for each region [7]. These methods have helped to reveal novel driver alterations in many cancer types [8]—many of which have been studied experimentally and confirmed to drive tumor initiation or progression.

However, in most cases, direct application of frequency-based methods is inadequate for systematically identifying drivers in individual tumors. To demonstrate this, we analyzed the frequencies of driver alterations in patients from 19 TCGA cancer types [1, 2]. For each cancer type, dNdScv and GISTIC2 were used to respectively identify candidate driver mutations and SCNVs at the population level (see Methods for details). There is a wide distribution of the number of candidate drivers per patient within individual cancers (Figure 1). In 17 out of 19 cancer types, the median number of total candidate drivers (mutations plus SCNVs) is ≥ 6, and in 6 of them is ≥ 10. It seems unlikely that some tumors require more than 15 hits whereas others require only 2 or 3 hits, especially within a single cancer type. An alternative explanation is that just a few mutations are essential for cancer formation and progression, and that others contribute to make the cancer more aggressive but are not essential. Non-essential drivers would be expected to occur more frequently than neutral passengers because tumors harboring these alterations are more likely to affect patient health and therefore be detected clinically. This explanation is consistent with the emerging view that there are different degrees of driver-ness, with some alterations being “major” drivers that strongly promote tumor progression, whereas other alterations are “minor drivers” that slightly contribute to tumor progression [9, 10]. Further supporting the idea that only a few alterations are essential for tumor formation is an epidemiological study by Tomasetti *et al.* suggesting that only 3 hits are required for formation of lung and colon cancers [11]. Distinguishing between essential and non-essential drivers is very important for prioritizing vulnerabilities in the tumor that could be considered as targets for therapeutic intervention.

**Figure 1:**
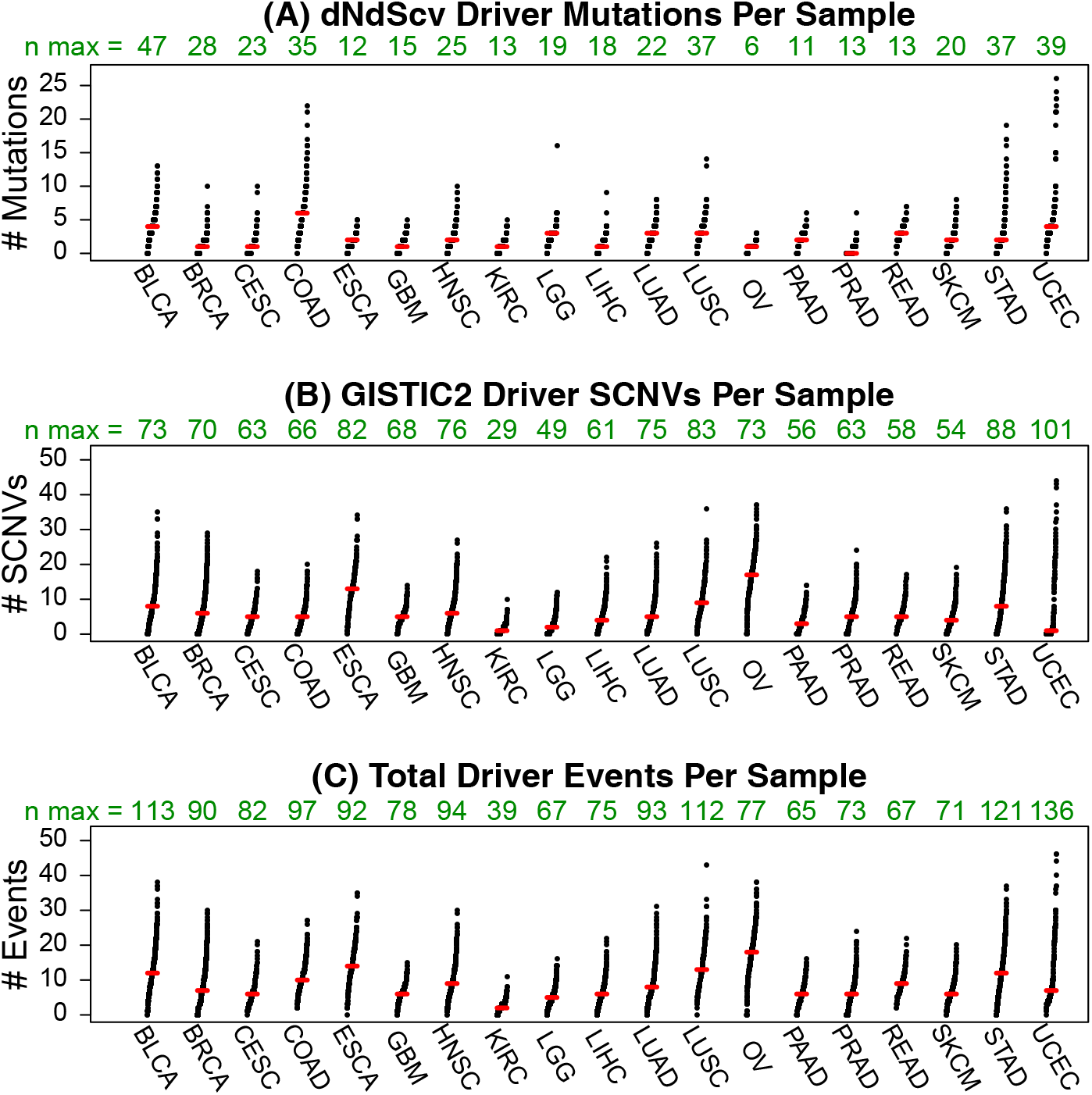
Number of driver events per patients across 19 TCGA cancer types. Driver events are SMGs identified by dNdScv and SCNVs identified by GISTIC2. Green numbers above each plot show the total number of candidate drivers identified for each cancer type by dNdScv (A), GISTIC2 (B) and both methods (C).

Statistical chance dictates that some of the predicted driver alterations tallied in Fig. 1 are expected to be falsely identified as drivers at the population level, even though they do not contribute to cancer formation in any patients. Alternatively, an alteration can be a true driver at the population level and yet a passenger in an individual sample. There is evidence that many tumors harbor passenger alterations within *bona fide* drivers. For example, Martincorena *et al.* [6] estimated that, based on the excess of non-synonymous mutation frequencies compared to the frequencies predicted by a covariate adjusted dN/dS model, greater than 95% of nonsense mutations in the tumor suppressors *ARID1A, RB1* and *APC* are estimated to be drivers, whereas the vast majority of missense mutations in these genes are estimated to be passengers. Meanwhile, for the tumor suppressor gene *TP53*, Martincorena *et al.* estimated that >95% of missense mutations are in fact drivers [6]. In theory, understanding the consequences of each alteration would help identify functionally neutral alterations, but most current applications consider any non-synonymous mutation in a known tumor suppressor to be a driver event. Nehrt *et al.* showed that consideration of functional domains within genes enables identification of candidate driver alterations that would be missed at the full gene level [12]. Adding to the complexity, there is evidence that the functional impact of a specific alteration can be different based on context. This situation is exemplified by the tumor suppressor *PTEN*, for which the vast majority of nonsense mutations are estimated to be drivers in breast cancer and kidney cancer. Surprisingly, based on the dN/dS ratios in each cancer type it is estimated that in kidney cancer most *PTEN* missense mutations are passengers, whereas 90% of *PTEN* missense mutations are estimated to be drivers in breast cancer [6].

To overcome the limitations of frequency-based methods, several methods have been developed to infer driver alterations in individual patients. One example is DawnRank, which outputs a prioritized list of drivers in each patient by incorporating gene-gene interactions networks with differential expression profiles between tumor and adjacent normal tissue [13]. The authors evaluated DawnRank based on how well it reproduced known drivers, but they did not explore whether the results could be used to explain clinical outcomes. The output of DawnRank is unique for each patient making it a good tool for identifying ultra rare drivers. Another method for inferring personalized drivers is iCAGES, which incorporates functional predictions of the impact of specific alterations in individual patients, along with prior knowledge of consensus cancer genes to prioritize cancer genes in individual patients [14]. The mutated genes identified by iCAGES are subsequently used to prioritize drugs for each patient by considering the relatedness of the mutated genes to the known gene targets of the drugs. A possible limitation of DawnRank and iCAGES is that they do not consider statistical dependencies between different alterations throughout the population.

Another approach for understanding the biological underpinnings of tumor formation has been to use information about the co-occurrence and mutual exclusivity of combinations of drivers (reviewed in [15]). Several methods have been developed to discover *driver modules*, which are sets of genes that exhibit statistically significant mutual exclusivity in individual patients and high coverage in the population [16, 17, 18, 19, 20, 21, 22]. Driver modules are very helpful for organizing the heterogeneous landscape of alterations into a small number of biological processes or pathways that are frequently perturbed in the population. Moreover, the alterations that are identified in driver modules are more likely to be actual driver alterations, because passenger alterations are not expected to display patterns of mutual exclusivity. These methods can help prioritize individual alterations that are part of discovered driver modules.

While there are many methods to identify sets of mutually exclusive candidate driver genes, there are comparatively very few methods that provide information about the co-occurrence of particular genes in individual patients. A likely reason for this is the surprising finding by Canisius *et al.* that there is little evidence of statistically significant co-occurrence between somatic mutations in cancer [23]. Zhang *et al.* found evidence of statistically significant co-occurrence between driver pathways [24]. The juxtaposition of this observation with the lack of significant co-occurrence at the alteration level suggests that when two pathways cooperate, most of the genes in each pathway are able to cooperate with several genes in the other pathway.

Several methods have been developed to detect drivers and pathways by integration of different types of data and prior knowledge. LNDriver showed improved ability compared to frequency methods to identify known cancer genes by incorporating a gene-gene interaction network into driver gene detection [25]. Chen *et al.* developed MAXDRIVER as a method for identifying candidate driver genes within copy number alterations by incorporating gene similarity networks and gene-cancer association networks [26].

The methods discussed so far can be grouped into four general categories: 1) frequency-based methods that predict drivers at the population level, 2) methods that output personalized ranked lists of driver genes in individual patients, 3) methods that detect sets of mutually exclusive driver modules, and 4) methods that predict drivers at the population level by integrating multiple types of data. In this paper, we introduce Cancer Rule Set Optimization (CRSO) to address a specific question that is not directly addressed by any of the aforementioned methods: what are the specific combinations of alterations in individual patients that are essential and collectively sufficient for tumor initiation and progression? In addition to prioritizing therapeutic vulnerabilities, this information would advance our understanding of driver gene cooperation. CRSO is developed as part of a theoretical framework that assumes the existence of specific combinations of two or more alterations called *rules* that cooperatively can drive cancer transformation when they co-occur in a host cell. Rules are assumed to be minimally sufficient, meaning that exclusion of any of the alterations renders the remaining collection of alterations as insufficient to drive cancer. CRSO seeks to find a collection of rules called a *rule set* that represent all of the different minimal ways for cancer to happen in the population, i.e., every sample is required to harbor all of the alterations in at least one of the rules. The output of CRSO is intended to provide insight into the ways different alterations can cooperate to cause cancer, as well as to infer the likely essential drivers in individual patients. When a patient is *assigned* to a rule it is assumed that the alterations comprising the rule are drivers and that all non-essential alterations in the patient are passengers. Despite this dichotomization, CRSO may be very robust to the presence of helper alterations.

Dash *et al.* [27] recently proposed an innovative method for identifying two-hit combinations that are likely responsible for carcinogenesis. The authors represented the problem as an instance of the weighted set cover problem [28] and showed that the identified set of combinations could discriminate normal and cancer samples with high accuracy. The combinations in Dash *et al.* are similar to the rules in CRSO and the set of combinations identified as the solution to the weighted set cover algorithm is similar to the optimal rule set identified by CRSO. However, there is a major difference in the optimization criteria in each of the methods. In [27] sets of combinations are prioritized based on how well they can discriminate normal and tumor samples in a training set. This choice of criteria assumes that combinations that occur only in tumors are likely to be cooperating drivers. There are many differences in the mutational landscapes of tumors and normal tissue and it is possible that many combinations involving passenger mutations are also likely to be observed almost exclusively in tumors. Alternatively, CRSO prioritizes rule sets that maximize the likelihood of observing the data by minimizing the statistical penalty caused by passenger alterations. Additionally, the present work expands upon the approach presented in [27] by considering combinations that involve more than two events, and by exploring clinical applications of stratifying patients based on predicted driver combinations.

One of the distinguishing features of CRSO is the incorporation of patient-specific, alteration-specific passenger probabilities that reflect how likely specific observations would have occurred by chance. The passenger probabilities for mutations are based on estimated passenger somatic nucleotide mutation rates for all nucleotide substitutions, calculated as in Cannataro *et al.* [29]. CRSO uses this information to prioritize rules that account for the most unlikely observations instead of just the most frequent observations. For example, recurrent hotspot mutations such as substitutions in codon 600 of *BRAF* or in codon 61 of *KRAS* are highly unlikely to happen by chance and are prioritized over non-recurrent substitutions in tumor suppressors. Nearly all of the methods that investigate driver interactions use binary alteration matrices to represent the data. CRSO is the only method we know of that searches for driver combinations based on high resolution passenger probabilities along with recurrence of combinations. CRSO can identify a particular alteration as a driver in one sample and as a passenger in another, based on the other alterations in each patient and patient specific passenger probabilities. We show that CRSO can identify subtypes with different clinical outcomes even though the optimal rule sets are found without any information about the outcomes.

## 2 Results

### 2.1 CRSO Overview

CRSO is an algorithm for finding combinations of genomic alterations that are predicted to cooperatively drive cancer in individual patients. The features considered by CRSO are mutations identified as candidate drivers by dNdScv [6] and copy number variations identified as candidate drivers by GISTIC2 [7]. We refer to these candidate driver alterations as **events**. The inputs into CRSO are specified by three event-by-sample matrices: **M**, **D** and **P**. **M** is a categorical matrix that describes the landscape of driver alterations in the population. Each event in **M** can take one of several values, called **observation types**, depending on the event type. For example, a mutation event can take values in the set {*WT, HS, L, S, I*}, corresponding to wild-type, hotspot mutation, loss mutation, splice site mutation or in-frame indel (see Methods section 5.1.1 for details). **D** is a binary matrix that is directly derived from **M**. The entries of **D** represent information about whether a particular event has happened in a particular patient, regardless of the specific observation type. Specifically, **D_ij_** equals 0 if **M_ij_** is wild-type, and equals 1 otherwise. **P** is the penalty matrix and represents information about how unlikely it would be for each observation in **M** to happen by chance, i.e., as a passenger event. The entry **P_ij_** is the negative log probability that **M_ij_** would be observed under the assumption that event *i* was not selected for in sample *j*. We make the simplifying approximation that wild-type events have passenger probability of 1, so that if **M_ij_** is wild-type then **P_ij_** = 0. Figure 2A shows an example of the input matrices using a miniature dataset that was extracted from TCGA melanoma (SKCM) data for the purposes of illustration.

**Figure 2:**
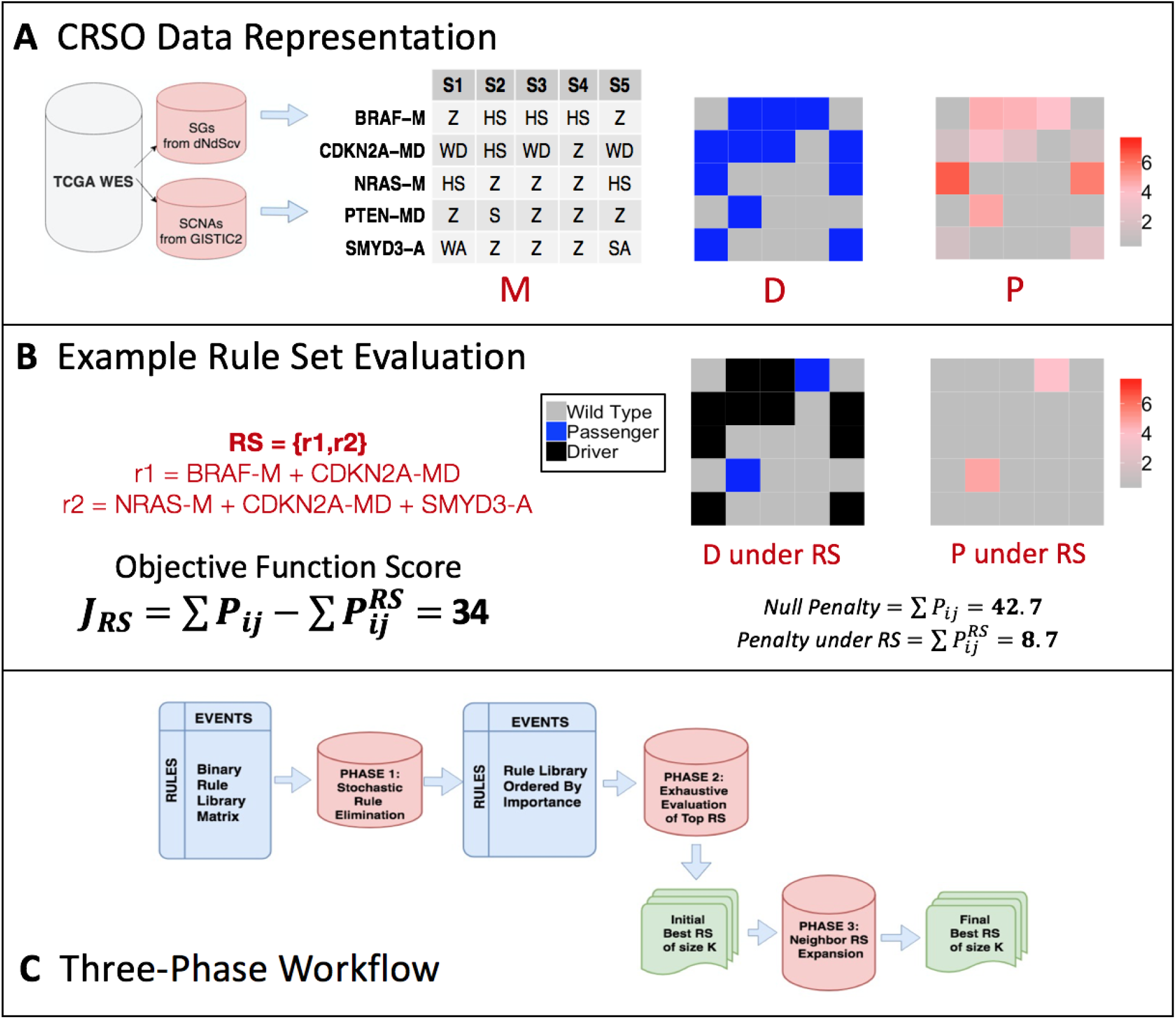
Visualization of CRSO using toy example. A) Candidate driver events identified by dNdScv and GISTIC2. Data is first represented as categorical matrix **M**, where each event can take one of several observation types, or *Z* if the event is wild-type. Events in **M** are represented as a binary matrix **D** for making rule assignments. The penalty matrix **P** contains penalties for each possible observation based patient-specific, event-specific and observation-type specific passenger probabilities. B) Objective function calculated for an example rule set. Under the proposed rule set, the assigned events are designated as drivers, and the corresponding penalties are reduced to 0. C) Workflow of three-phase procedure for identifying the best rule sets of size *K*, for a range of *K*s.

The CRSO model of cancer formation defines a **cancer rule** to be a set of two or more events that is hypothesized to drive cancer in tumors that satisfy the rule. A tumor satisfies a cancer rule if all of the events that comprise the rule co-occur in the tumor. Cancer rules are defined to be minimally sufficient, meaning that exclusion of any event renders the remaining collection of events as insufficient to cause cancer. A **cancer rule set** is defined to be a collection of cancer rules that collectively account for a population of tumors. In other words, a cancer rule set is assumed to represent all of the minimal ways cancer can happen in the population. The terms *rule* and *cancer rule* are used interchangeably throughout this paper, as are the terms *rule set* and *cancer rule set*. Two rules are defined to be *family members* if one rule is a strict subset of the other. Cancer rule sets cannot contain any two rules that are family members. To see why, consider that rule 1 is a strict subset of rule 2. Inclusion of rule 1 implies that rule 2 is not minimal, whereas inclusion of rule 2 implies that rule 1 is insufficient. The size of a rule set denotes the number of rules it contains.

CRSO is an optimization procedure over the space of possible rule sets. The ability of a rule set to account for the distribution of events in the population is quantified by an **objective function**. When a sample is assigned to a rule in a rule set, the events that comprise the rule are considered to be drivers within that sample, and all of the remaining events in that sample are considered passengers. The absence of a rule set is considered a null rule set, under which all of the events in the dataset are considered to be passengers. The **penalty** of a proposed rule set *RS* is the sum of the penalties of all of the events designated as passengers under *RS*. The objective function score of *RS*, *J*_*RS*_, is the total reduction in penalty under *RS* compared to the null rule set. Formal calculation of *J*_*RS*_ is present in Methods section 5.2.2. Figure 2B shows an example of a rule set consisting of two rules applied to the miniature melanoma dataset. The total penalty is greatly reduced once samples are assigned to rules and the corresponding events are designated as drivers. The objective function quantifies how well the rule set accounts for the observations in **M**.

The performance of a given rule set always increases or stays the same when a new rule is added to it. The goal of CRSO is to find the rule set that achieves the best balance of objective function score and rule set size, which is called the **core rule set**. CRSO uses a three phase algorithm to first find the highest scoring rule set of size *K* over a range of *K* values. The core rule set is then determined from among all of the solutions of size *K*. A subsampling process is used to identify an expanded list of **generalized core rules** (GCRs). A confidence score is determined for each GCR reflecting the probability that it is part of the theoretical “true” rule set. The full description of CRSO methodology is presented in the Methods section.

### 2.2 Application to TCGA Melanoma Data

CRSO was applied to 19 cancer types obtained from TCGA using default parameter values (Table 6). The number of samples analyzed for each tissue type ranged from a low of 120 in rectum adenocarcinoma (READ) to a high of 963 in breast invasive carcinoma (BRCA) (Table 1). For each tissue type, all samples that have both copy number and mutation profiles were included. The output of each CRSO run is a detailed report containing summaries and visual representations of the results. The reports for the 19 tissue types are included in the supplementary materials (supp. folder “TCGA.REPORTS”). In this section we present the results for one TCGA cancer type: skin cutaneous melanoma (SKCM). We choose to present one cancer type in detail in order to explain how CRSO works and how to interpret the results obtained from CRSO.

**Table 1:** Summary of 19 TCGA tissues types investigated with CRSO

| Cancer | Full Name | Samples | Events (M/C)<sup>a</sup> | RCT<sup>b</sup> | Rules | Core K | Core Cov. |
| --- | --- | --- | --- | --- | --- | --- | --- |
| BLCA | Bladder urothelial carcinoma | 392 | 113 (47/66) | 0.04 | 933 | 13 | 75 |
| BRCA | Breast invasive carcinoma | 963 | 90 (28/62) | 0.03 | 449 | 13 | 60 |
| CESC | Cervical and endocervical cancers | 191 | 82 (23/59) | 0.031 | 173 | 11 | 53 |
| COAD | Colon adenocarcinoma | 362 | 97 (35/62) | 0.048 | 986 | 7 | 79 |
| ESCA | Esophageal carcinoma | 184 | 92 (12/80) | 0.066 | 827 | 11 | 82 |
| GBM | Glioblastoma multiforme | 273 | 78 (15/63) | 0.03 | 186 | 12 | 78 |
| HNSC | Head and neck squamous cell carcinoma | 505 | 94 (25/69) | 0.03 | 926 | 9 | 69 |
| KIRC | Kidney renal clear cell carcinoma | 433 | 39 (13/26) | 0.03 | 18 | 6 | 47 |
| LGG | Brain lower grade glioma | 513 | 67 (19/48) | 0.03 | 210 | 5 | 65 |
| LIHC | Liver hepatocellular carcinoma | 366 | 75 (18/57) | 0.03 | 146 | 13 | 57 |
| LUAD | Lung adenocarcinoma | 478 | 93 (22/71) | 0.03 | 291 | 10 | 60 |
| LUSC | Lung squamous cell carcinoma | 178 | 112 (37/75) | 0.052 | 828 | 13 | 80 |
| OV | Ovarian serous cystadenocarcinoma | 455 | 77 (6/71) | 0.094 | 932 | 11 | 83 |
| PAAD | Pancreatic adenocarcinoma | 126 | 65 (11/54) | 0.048 | 210 | 3 | 75 |
| PRAD | Prostate adenocarcinoma | 492 | 73 (13/60) | 0.03 | 301 | 9 | 54 |
| READ | Rectum adenocarcinoma | 120 | 67 (13/54) | 0.05 | 735 | 7 | 82 |
| SKCM | Skin Cutaneous Melanoma | 290 | 71 (20/51) | 0.03 | 197 | 11 | 70 |
| STAD | Stomach adenocarcinoma | 391 | 121 (37/84) | 0.036 | 850 | 12 | 63 |
| UCEC | Uterine Corpus Endometrial Carcinoma | 242 | 136 (39/97) | 0.038 | 930 | 9 | 83 |

<sup>a</sup> Events shows the total number of events as determined by dNdScv and Gistic2, with (M/C) indicating the number of mutations and CNVs.
<sup>b</sup> RCT is the rule coverage threshold, and was chosen to be the larger value out of .03 or the smallest threshold satisfied by less than 1000 rules.

The SKCM dataset consists of 290 samples for which both mutation and copy number data are available. The features considered by CRSO were 20 SMGs identified by dNdScv, as well as 34 SCNV deletions and 20 SCNV amplifications identified by GISTIC2. SCNV amplifications and deletions were represented according to the narrow peak genes provided by GISTIC2. Three genes, *CDKN2A, PTEN* and *B2M*, were identified as SMGs and as part of significant deletion peaks. These events were represented as hybrid “mutDel” events by combining the gene-level mutations with the gene-containing deletions for each gene, resulting in a total of 71 events: 17 mutations, 20 amplifications, 31 deletions and 3 mutDels. Each non wild-type observation is represented as one of several observation-types. Mutations are subdivided into hotspot mutations (HS), loss mutations (L), in-frame indels (I) or splice mutations (S). Amplifications are subdivided into weak/strong amplifications (WA/SA), and deletions are subdivided into weak/strong deletions (WD/SD). Hybrid events take values from a larger set, since these events can involve copy number changes, mutations or both. Exact definitions of all observation types associated with each event type are provided in the Methods sections 5.1.1, 5.1.3 and 5.1.5.

Figure 3A shows the binary representation, **D**, of the SKCM dataset for the 25 most frequent events, as well as the types and frequencies of the events. The penalty matrix, **P**, represents information about how unlikely it would be for each observation to have happened by chance, as a passenger. The entries of **P** are negative-log probabilities of patient-specific, event-specific and observation-type specific passenger probabilities. Fig. 3B shows **P** for the 25 most frequent events in the SKCM dataset, as well as the total penalties contributed by each event. Two mutation events, *BRAF-M* and *NRAS-M*, have large penalties in many samples reflecting the fact that these two genes are known to contain highly recurrent hotspot alterations (*BRAFV600* and *NRASQ61*). The penalties provide information about how unlikely an event is to be a passenger that would be missed from consideration of frequency alone (Fig. 3A–B). For example, *TP53* mutations are observed in 17% of patients and have a total penalty of 150, whereas *ADAM18* mutations are observed in 19% of patients but have a lower total penalty of 92.

**D** was used to build a rule library consisting of all combinations of two or more events that co-occur in the same patient in at least 3% of patients. There were 197 eligible rules, of which 165 consist of two events and 32 consist of three events. Once the rule library was prepared, the three-phase algorithm was used to determine the best rule set for sizes *K* = 1 … 16. The three-phase algorithm (Figure 2C) is presented in detail in section 5.2.3. Briefly, in phase 1 all of the rules are ranked based on how much they contributed to random groups of rules. In phase 2, initial best rule sets of size *K* are determined by exhaustive evaluation over a subset of the rule library. Exhaustive evaluation of all rule sets from among the full rule library is computationally prohibitive for *K >* 3. To overcome this, exhaustive evaluation is performed instead over the top *n* rules based on phase 1 ranking, where *n* is chosen for each *K* to be the largest *n* such that there are at most 200,000 possible rule sets. In phase 3, additional rules are considered for each *K* beyond those considered in phase 2. This expansion of the search space is accomplished by systematically substituting small subsets of rules within the top rule sets with yet to be considered rules. Using 80 CPUs, the run-time for the three-phase algorithm was approximately 20 minutes for the melanoma dataset.

**Figure 3:**
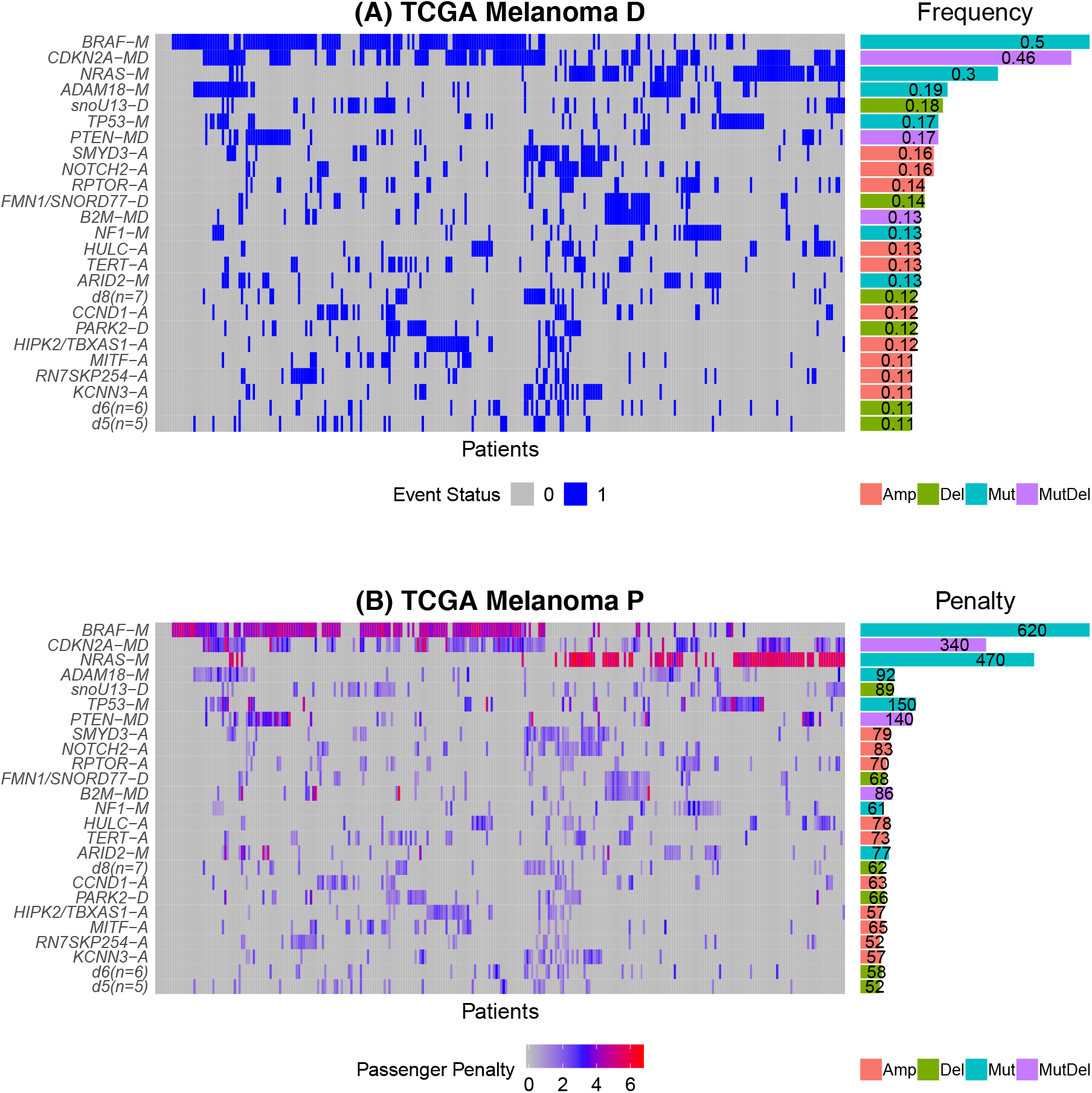
CRSO Representation of TCGA Melanoma Dataset. A) Binary representation of top-25 most frequent events. Horizontal bars on the right show event frequencies across the population. B) Penalty matrix for top-25 most frequent events. Horizontal bars on the right show total event penalties. Event types indicated by suffixes: -A for amplifications, -D for deletions, -M for mutations, and -MD for mutDel hybrid events.

Upon completion of the three-phase algorithm, a ranked list of the top 20,000 rule sets for *K* = 1 ... 16 was extracted. The results were filtered to only include rule sets for which each rule covers at least 3% of samples. No rule sets satisfied the filtering requirement for *K >* 12. We denote the best performing filtered rule set of size *K* as *RS*_*K*_, and we denote the performance and coverage of *RS*_*K*_ by *J*_*K*_ and *Cov*_*K*_, respectively. Figure 4A shows the objective function score (left) and coverage (right) of the best filtered rule sets for *K* = 1 … 12. In order to balance objective function score with over-fitting, CRSO defines a core rule set to be the rule set corresponding to the smallest *K* for which *Cov*_*K*_ ≥ 0.95 * *max*(*Cov*_*K*_) and *J_K_ ≥* 0.9 * *max*(*J*_*K*_). The core rule set was determined to be the best rule set of size *K* = 11, i.e., *RS*_*K*=11_. Out of the 11 core rules, 5 include *BRAF-M* but not *NRAS-M*, and 5 include *NRAS-M* but not *BRAF-M* (Fig. 4B). *BRAF* and *NRAS* mutations both activate the *RAS-RAF-MAPK* signaling pathway and are mutually exclusive in almost all melanoma patients [30]. The results suggest that *BRAF* and *NRAS* both cooperate with the tumor suppressors *CDKN2A* and *TP53*. The other partners are unique to each of the oncogenes, as *BRAF* is predicted to cooperate with *PTEN-MD*, *RN7SKP254-A* and *HIPK2/TBXAS-A*, while *NRAS* is predicted to cooperate with *ADAM-18*, *SMYD3-A* and *HULC-A*. In total only 70% of patients are assigned to at least one of the rules in the core rule set. Some of the core rules are not among the highest ranking in the rule library based on coverage or single rule performance (SJ) (Table 2). For example, *NRAS-M + SMYD3-A* is among the core rule set, although it has a low coverage of 5.5% (rank = 42.5), and is ranked 20th in SJ. This can happen because CRSO does not consider rules in isolation, but rather in the context of the other rules in the rule set. More extreme examples of this situation are observed in other tissue types. For example, in glioblastoma multiforme (GBM), the rule *TP53-M + RB1-M* is identified as part of the core rule set (*K*=12), event though it ranks outside of the top 50 in both coverage and SJ (see GBM tissue report in supplement).

**Table 2:**
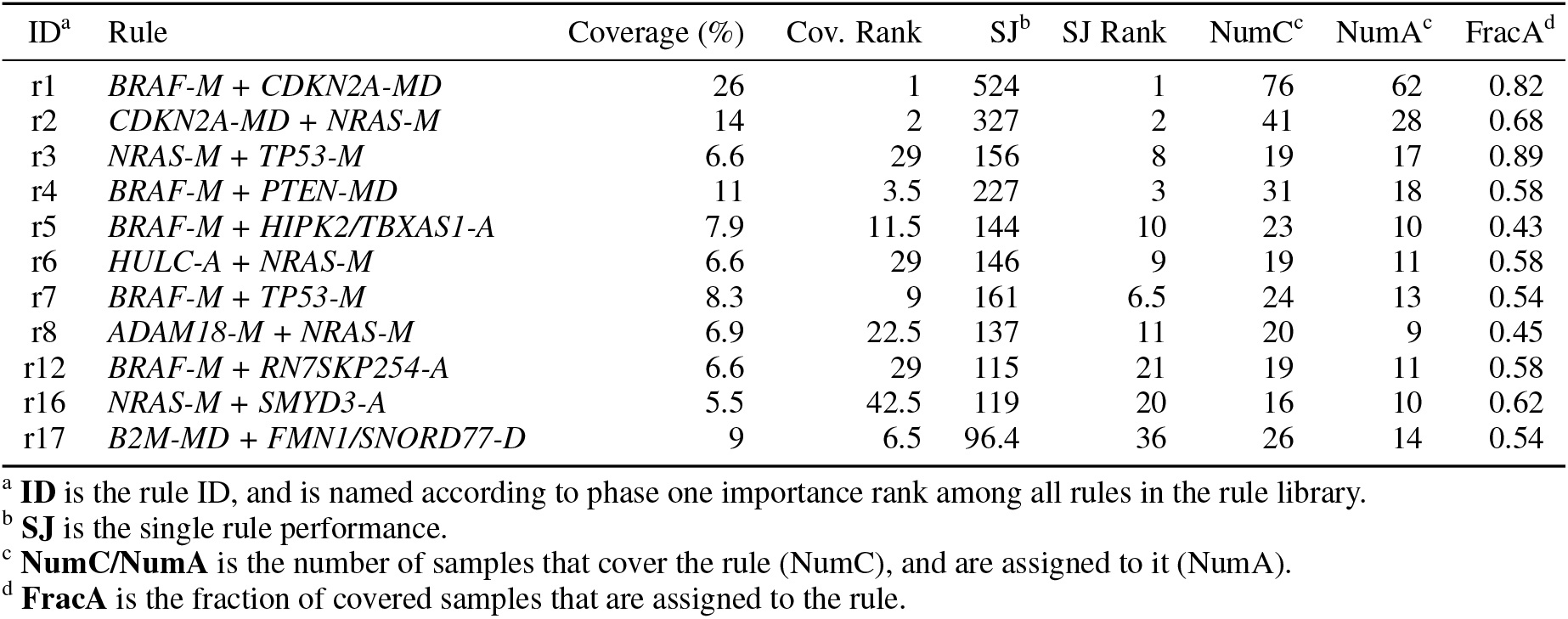
Characteristics of melanoma core rule set

| ID<sup>a</sup> | Rule | Coverage (%) | Cov. Rank | SJ<sup>b</sup> | SJ Rank | NumC<sup>c</sup> | NumA<sup>c</sup> | FracA<sup>d</sup> |
| --- | --- | --- | --- | --- | --- | --- | --- | --- |
| r1 | *BRAF-M + CDKN2A-MD* | 26 | 1 | 524 | 1 | 76 | 62 | 0.82 |
| r2 | *CDKN2A-MD + NRAS-M* | 14 | 2 | 327 | 2 | 41 | 28 | 0.68 |
| r3 | *NRAS-M + TP53-M* | 6.6 | 29 | 156 | 8 | 19 | 17 | 0.89 |
| r4 | *BRAF-M + PTEN-MD* | 11 | 3.5 | 227 | 3 | 31 | 18 | 0.58 |
| r5 | *BRAF-M + HIPK2/TBXAS1-A* | 7.9 | 11.5 | 144 | 10 | 23 | 10 | 0.43 |
| r6 | *HULC-A + NRAS-M* | 6.6 | 29 | 146 | 9 | 19 | 11 | 0.58 |
| r7 | *BRAF-M + TP53-M* | 8.3 | 9 | 161 | 6.5 | 24 | 13 | 0.54 |
| r8 | *ADAM18-M + NRAS-M* | 6.9 | 22.5 | 137 | 11 | 20 | 9 | 0.45 |
| r12 | *BRAF-M + RN7SKP254-A* | 6.6 | 29 | 115 | 21 | 19 | 11 | 0.58 |
| r16 | *NRAS-M + SMYD3-A* | 5.5 | 42.5 | 119 | 20 | 16 | 10 | 0.62 |
| r17 | *B2M-MD + FMN1/SNORD77-D* | 9 | 6.5 | 96.4 | 36 | 26 | 14 | 0.54 |

<sup>a</sup> **ID** is the rule ID, and is named according to phase one importance rank among all rules in the rule library.
<sup>b</sup> **SJ** is the single rule performance.
<sup>c</sup> **NumC/NumA** is the number of samples that cover the rule (NumC), and are assigned to it (NumA).
<sup>d</sup> **FracA** is the fraction of covered samples that are assigned to the rule.

**Figure 4:**
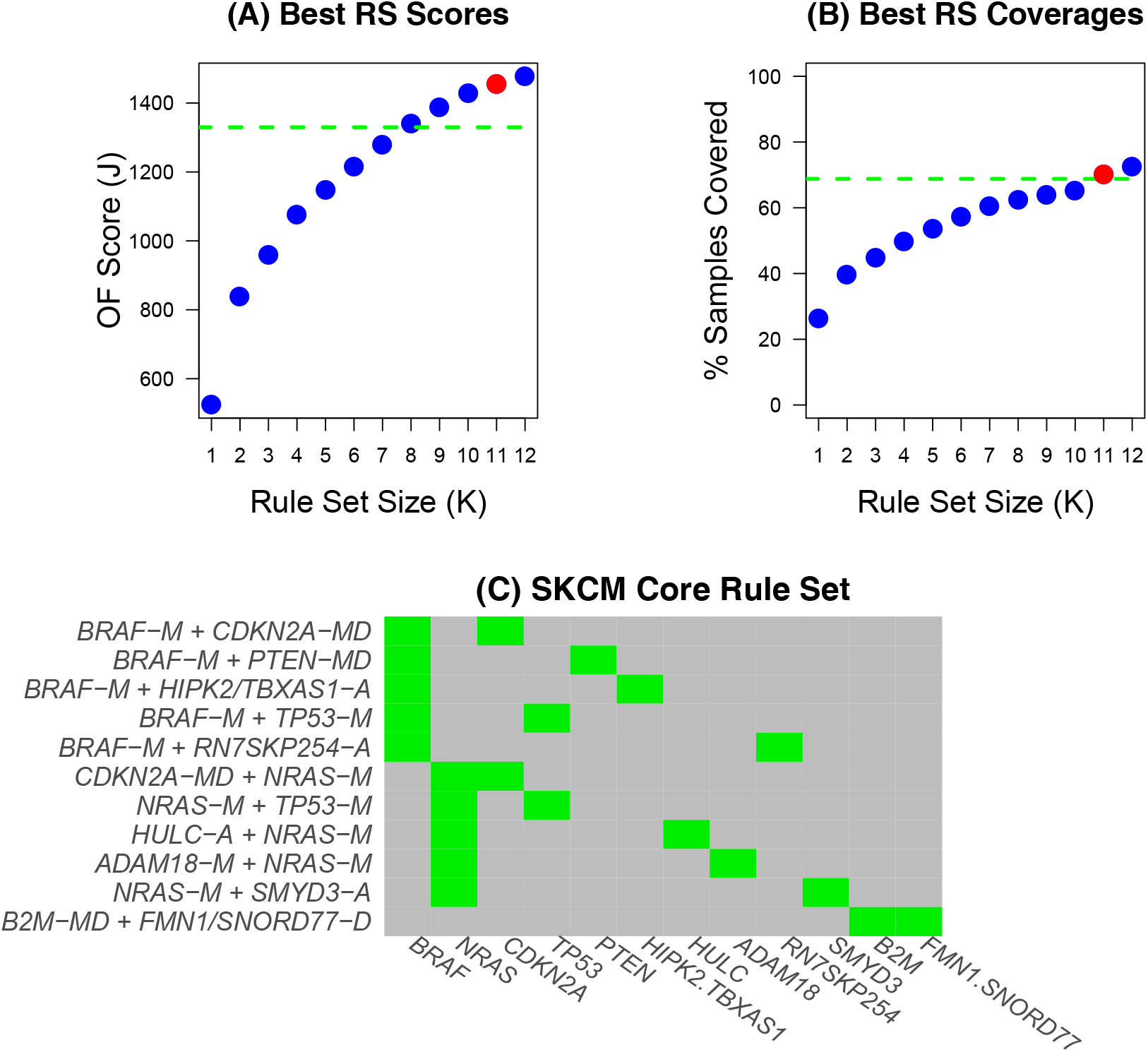
Melanoma CRSO results. A–B) The objective function score (A) and coverage (B) of the best filtered rule sets. The core rule set corresponds to *K* = 11 and is shown in red. The dashed green lines are the thresholds for determining the core rule set. C) Composition of the core rule set.

Each patient is assigned to a rule in the core rule set, or to the null rule. When a patient is assigned to a rule, the events that comprise the rule are designated as drivers (Figure 5A). Some samples satisfy multiple rules and these samples were assigned to the rule that minimizes the passenger penalty in each specific sample (Fig. 5B). Thirty percent of the SKCM patients do not satisfy any of the core rules, and are assigned to the null rule, indicated by the red color bar in Fig. 5A. Many of the unassigned samples contain high-penalty hotspot mutations in *BRAF* and *NRAS* (Fig. 5C), suggesting that the core rule set did not identify all of the possible partners for these oncogenes.

**Figure 5:**
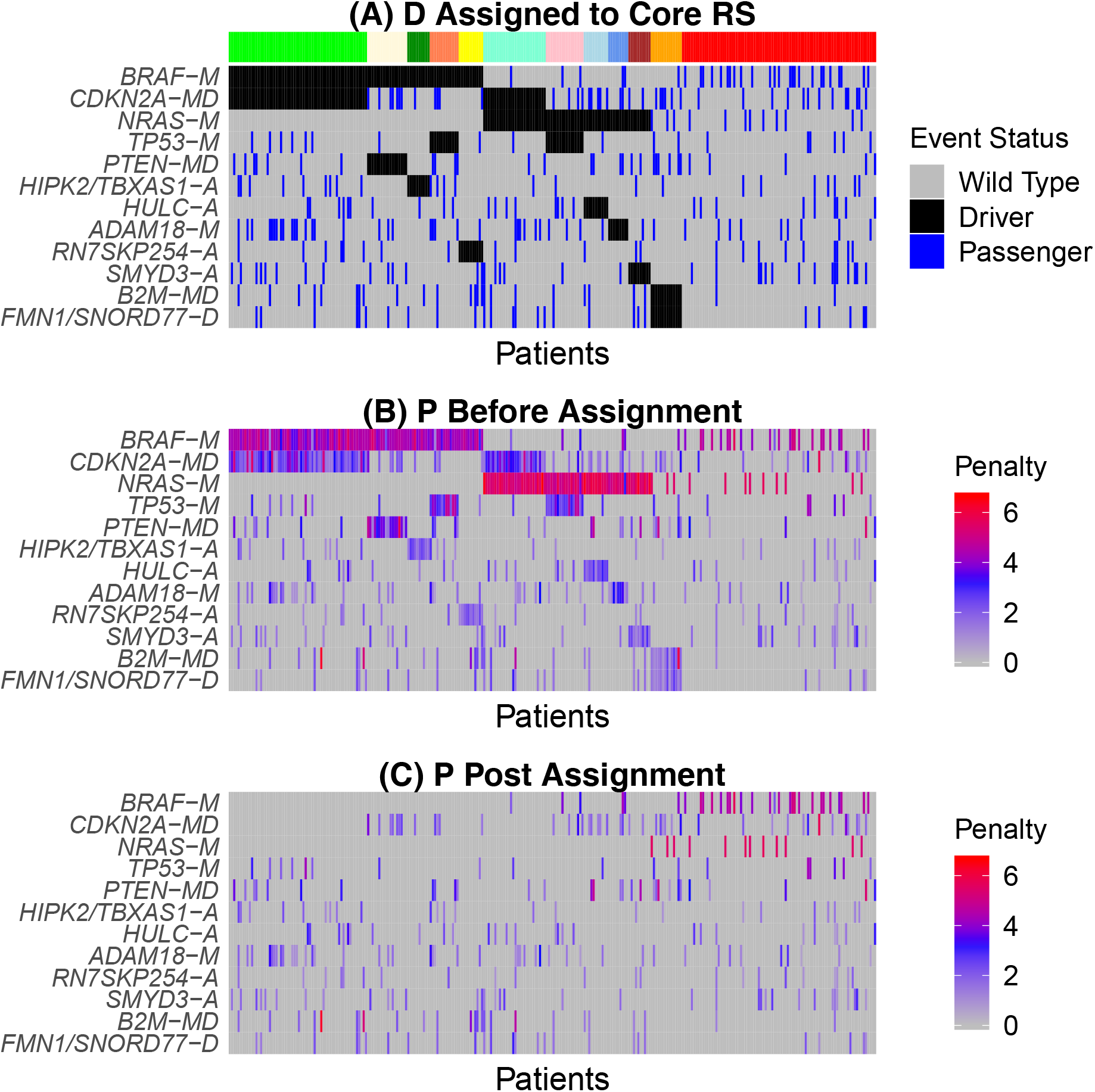
A) Heatmap of the binary alteration matrix **D** under core rule set assignment. Events are ordered by frequency. Samples are ordered according to rule set membership, as indicated by the color bar. The right-most group (red bar), are not assigned to any rule. For each sample, assigned events are designated as drivers (black), and unassigned events (blue) are assumed to be passengers. B–C) Heatmaps of the passenger penalty matrix **P** before and after assignment to the core rule set.

### 2.3 Generalized Core Rules

The core rule set represents the single best rule set that describes the given dataset. However, one limitation of the core rule set is that it can be sensitive to the inclusion/exclusion of a small number of samples. For melanoma, the coverage of the best rule set of size 9 is barely below the core coverage threshold (Fig. 4A), suggesting that exclusion of a few samples, or a slight change to the core coverage criteria might result in identification of a different core rule set.

To assess and improve the stability of the results, CRSO identifies a collection of **generalized core rules** (GCRs) by randomly sub-sampling 80% of the original dataset 100 times. For each subsample the best performing rule-sets of size *K* were identified for *K* = 1 … 12. To reduce computing time, only the top 2000 rule sets for each *K* determined from the original dataset were considered. A core rule set is extracted for each iteration by determining the smallest *K* for which *Cov*_*K*_ and *J*_*K*_ satisfy the core coverage and performance thresholds. Instead of fixing the coverage threshold at 0.95, it is chosen to be a uniform random value in [0.85, 0.98]. Similarly, the performance threshold is chosen to be a uniform random value in [0.8, 0.95]. The superset of the core rule sets identified in all 100 iterations are defined to be GCRs. Each GCR is associated with a confidence level that is defined to be the percentage of iterations for which the rule was identified. The GCRs and their associated confidence levels provide a more stable representation of the CRSO results. Because the collection of GCRs is the superset of many different core rule sets, we point out the collection of GCRs is not guaranteed to be a valid rule set, as it may contain rules that are family members.

In addition to GCRs, generalized core duos (GCDs) and generalized core events (GCEs) were also extracted from the collection of core rule sets determined by sub-sampling. A duo is a pair of events that appear together in at least one rule within a rule set. For each subsample the core duos are defined to be all of the unique duos contained within the associated core rule set. GCDs are then defined to be the superset of all core duos and are assigned confidence levels according to the percentage of iterations in which the duo is observed. GCEs are defined as the superset of all core events and are assigned confidence levels analogously. We are interested in GCEs because they reveal how consistently individual events are being identified as part of core driver rules.

GCRs are partitioned into three groups according to confidence level. High and intermediate confidence groups are demarcated by thresholds of 80 and 40, respectively. Rules with confidence level below 40 are low confidence. The same confidence level groupings are defined for GCDs and GCEs. The SKCM GCRs (Fig. 6A) overlap highly with the SKCM GCDs (Fig. 6B). This is because all but two GCRs consist of two events and are already duos. However, in other cancer types we observe duos that occur in multiple iterations as part of different rules. This can lead to a highly recurrent duo being overlooked because it is not part of any highly recurrent rules. An extreme example of this was observed in colon adenocarcinoma (COAD), for which 12 GCDs with confidence level ≥ 98 are identified, despite the fact that the highest confidence GCR has confidence level of 77 (see COAD TCGA report in supplemental materials).

**Figure 6:**
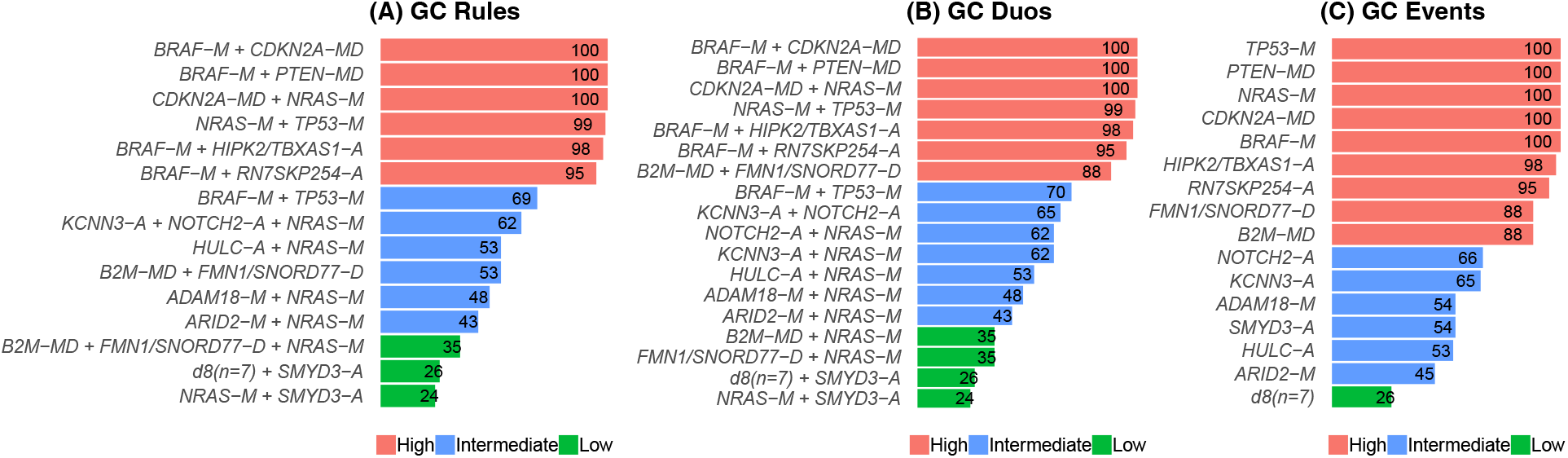
Summary of generalized core results. Bars show confidence levels, which are the percentage of sub-sample iterations containing the observation. GCRs (A), GCDs (B) and GCEs (C) that achieve a minimum confidence level of 10 are shown.

### 2.4 CRSO Reports from 19 TCGA Cancers

The CRSO reports in the supplemental materials provide a a detailed presentation of the CRSO findings for all 19 TCGA cancer types. Each report contains five sections, prefixed with SR to denote sections in the supplemental reports. SR Section 1 presents basic information about the dataset as well as heatmaps of the binary matrix **D** and the penalty matrix **P** for the 20 most frequent events. SR Section 2 shows the best rule sets of size *K* for *K* = 1 *... K*_*max*_. The performance and coverage curves are shown in SR Section 2.1 and the core rule set is highlighted in red. SR Section 2.2 presents a table of all of the rules comprising any of the *K* best rule sets. SR Section 3 presents a deeper dive into the core rule set. SR Section 3.1 is a table showing different characteristics of the core rules, including the coverage, phase 1 importance rank and single rule performance. SR Section 3.2 shows an event-by-rule breakdown of the core rule set. SR Section 3.3 shows heatmaps of **P** before and after assignment to the core rule sets. This is a visual representation of how well the rule set accounts for observations in **D**. SR Section 4 presents the generalized core results described above, and SR Section 5 is a dictionary of copy number events.

### 2.5 Performance and Coverage Convergence

Each cancer converges to the maximum performance and coverage at its own rate. For example, the core rule set in pancreatic adenocarcinoma (PAAD) consists of only three rules, whereas the core rule sets for most cancers contain more than 10 rules (see Table 1). Figure 7 shows the relative performance (A) and coverage (B) of the best rule sets of different sizes for each *K* for the 19 TCGA cancers.

**Figure 7:**
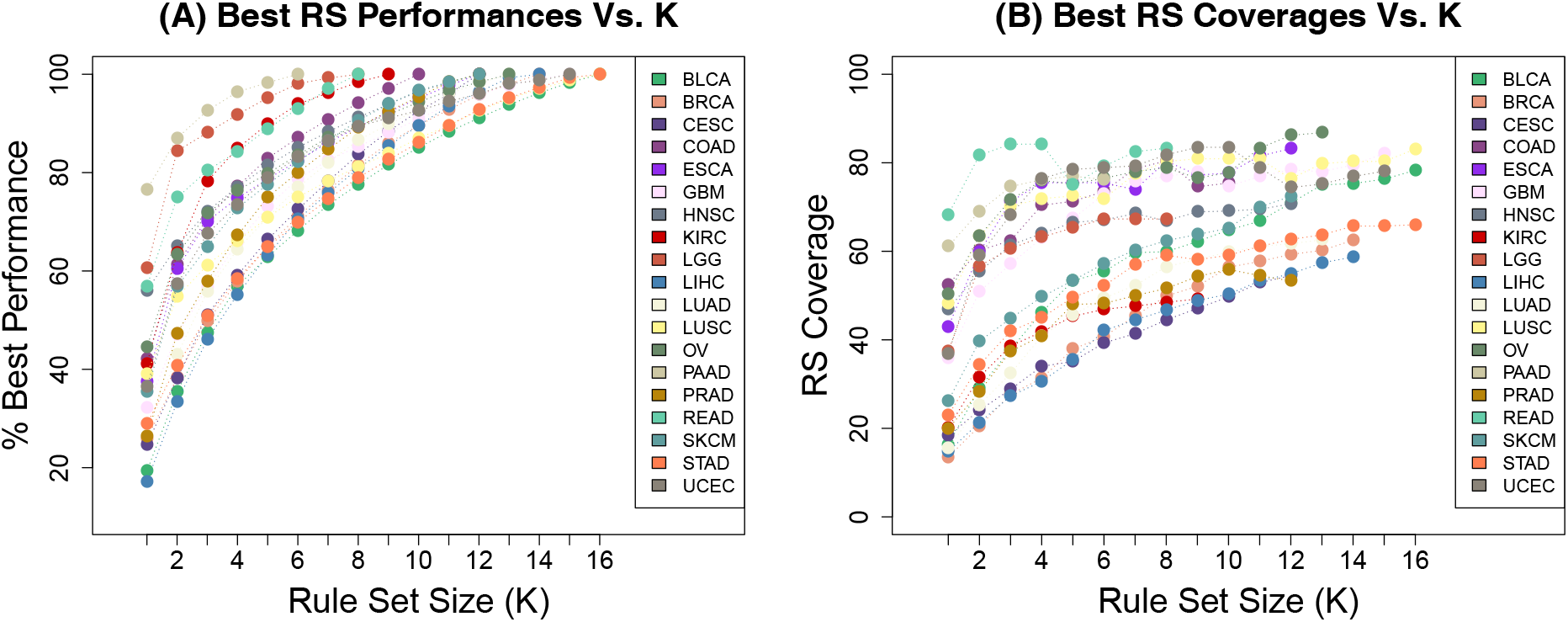
Performance and Coverage Convergence for Different Cancers.

### 2.6 Recurrent Driver Combinations Across Tissues

Many driver alterations are recurrently identified by GISTIC2 and dNdScv in multiple different cancer types. In order to determine whether any driver combinations are also shared across multiple cancers, we looked for overlap among the GCDs for each cancer type that achieve a minimum confidence value of 10. Overlap was investigated among duos instead of full rules in order to maximize the chances of finding recurrent combinations. Recurrence across multiple cancers is itself a measure of confidence, so that even low confidence GCDs that recur are interesting. Across the 19 cancer types there are 364 distinct duos and 58 of them (16%) were identified in at least two cancer types. In some cases the SCNV regions identified by GISTIC2 in different cancers can share overlapping genes. This analysis would consider these to be distinct events. For this reason we think that additional instances of recurrence involving driver genes that appear as part of non identical SCNV regions may have been missed.

Nineteen duos were identified in three or more cancers (shown in Table 3). The five most recurrent duos consist of *TP53* paired with the well known drivers: *PIK3CA, CDKN2A, RB1, PTEN* and *KRAS*. Some of the recurrent duos in Table 3 involve lesser known driver deletions, such as *LRP1B-D*, *FKSG52/MIR582/PDE4D-D* and *CCSER1/RN7SKP248-D*. The fact that these deletions are in duos that are identified in three or more cancer types suggests that they may be more important than has been appreciated.

**Table 3:**
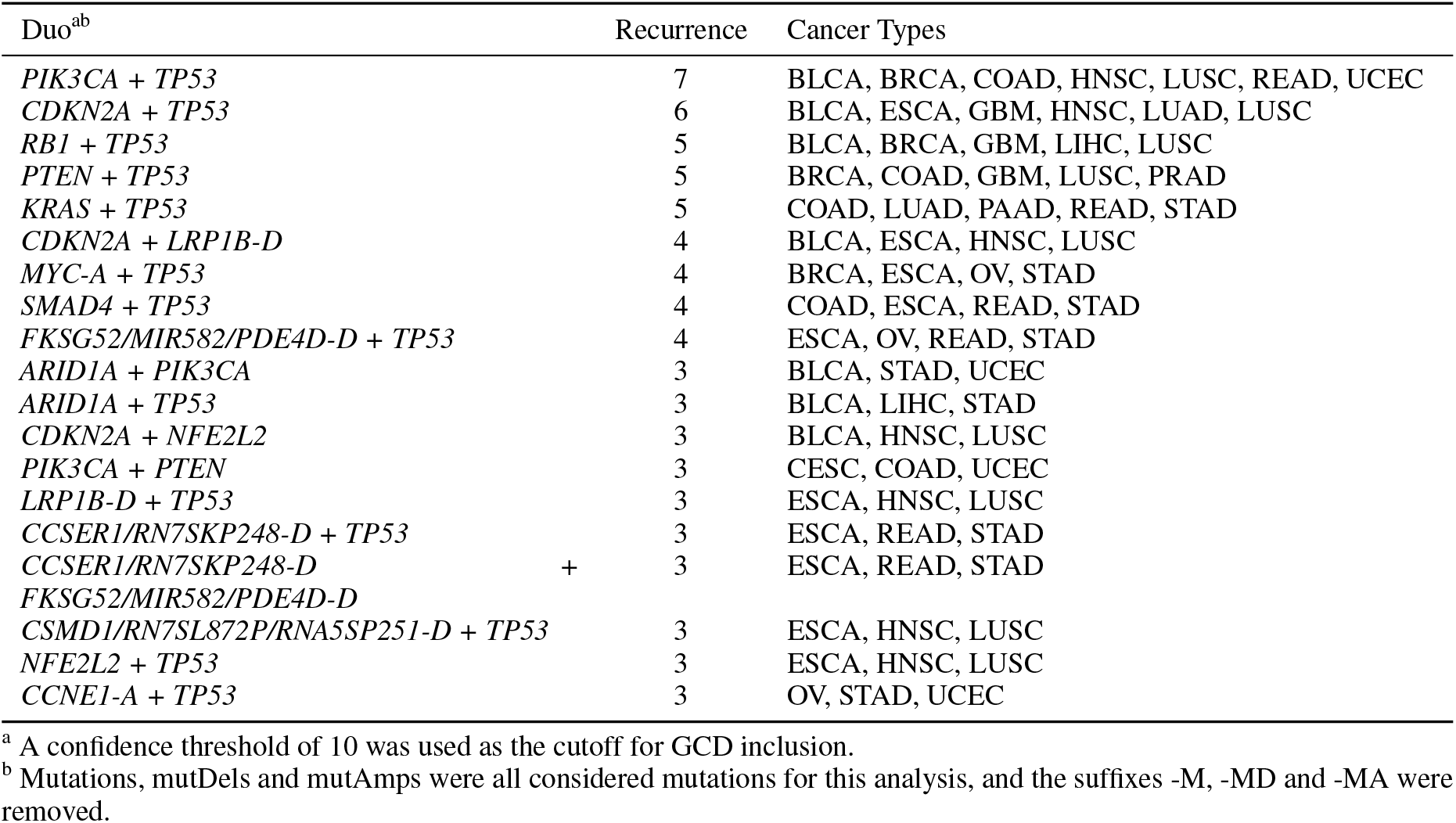
Generalized core duos (GCDs) that are identified in at least 3 cancer types

### 2.7 Association of Rules with Patient Prognosis

We next determined whether stratifying patients according to driver rules instead of individual driver events provides extra prognostic information. For each cancer type, GCRs were considered that have confidence level above 40. In cases where two GCRs were family members, the family member with lower confidence level was excluded. Note that all of the GCRs with confidence level greater than 50 are guaranteed to be non-family members. Driver events that appear in more than one GCR are of greatest interest, since these events are predicted by CRSO to occur in distinct genetic contexts. An event that is part of more than one GCR in a given cancer type is defined to be a **multi-rule event** for that cancer type.

Consider multi-rule event *E* that appears in rule *R*. The following test was performed to determine if there is statistical evidence of differential patient prognosis associated with *R*. The subset of patients that contain *E* are considered, and those that are wild-type for *E* are disregarded. Patients within this cohort were assigned to one of two classes: those that satisfy *R* are assigned to class “rule”, and those that harbor *E* but do not satisfy *R* are assigned to class “event”. Univariate cox-ph analysis was performed to test for differences in progression-free intervals (PFI) between the two classes. The result of each test is summarized with a *Z* score, as recommended in [31]. Compared to using *P* values for ascertaining statistical significance, *Z* scores provide additional information about the direction of the PFI differences. Positive *Z* scores indicate better PFI for patients in the rule class compared to patients in the event class, and negative *Z* scores indicate the opposite. PFI data were obtained from a recently published resource for TCGA outcome analysis [32], and the use of PFI as primary endpoint is consistent with the authors’ recommendations for best practices.

For each cancer type, each multi-rule event, *E*, was tested against all of the eligible GCRs that include *E*. A drawback to this approach is that there can be redundancy between different tests. For example, suppose most patients that contain *E* are assigned to one of *R*_1_ or *R*_2_, then testing *E* versus *R*_1_ is essentially the same as testing *E* versus *R*_2_ in the opposite direction. The tests should not be assumed to be independent, and conventional multiple hypothesis corrections may not be appropriate as a consequence. The reasons for testing *E* against each rule separately, instead of directly comparing all of the rules containing *E*, are two-fold. First, some multi-rule events occur in many GCRs making it difficult to interpret the results. Second, comparing multiple rules at once introduces the problem of dealing with patients that satisfy multiple rules.

A total of 220 tests were performed across the 19 cancer types, and 20 (9.1%) significant associations (|*Z*| ≥ 1.96) were detected (Table 4). Only those pairings for which both the event and rule classes contained at least ten patients were considered. It is tempting to argue that the expected percentage of significant associates should be 5% and that we are finding more associations than expected by chance. However, we are not comfortable making this assertion about the family-wise error rate because of the complicated dependences that may exist among the tests within each tissue type.

**Table 4:**
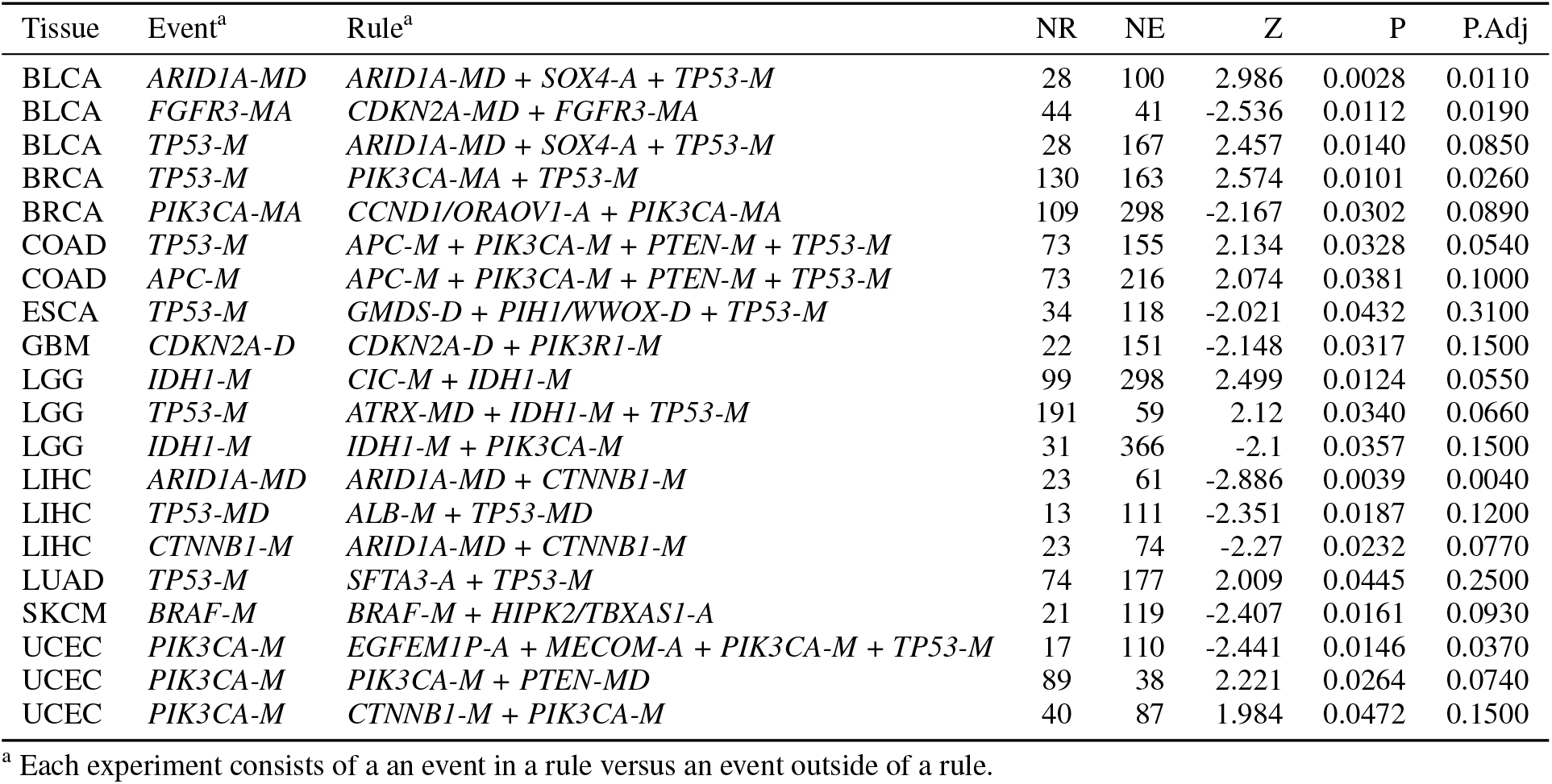
Significant PFI associations among consensus rule sets

A permutation test was performed in order to address the multiple hypotheses associated with each multi-event rule. For each multi-rule event, the PFIs of the samples that contain the event were scrambled 1000 times, and the smallest *P* value attained by any of the rule vs. event comparisons was stored for each iteration. An adjusted *P* value, *P*_*Adj*_, (Table 4), was defined to be the fraction of iterations that have permuted *P* value smaller than the real data *P* value.

The PFI associations identified in Table 4 have very different interpretations that are context dependent. For example, consider the pairing of the event *TP53-M* (*n* = 250 samples) and the rule *ATRX-MD* + *TP53-M* + *IDH1-M* (*n* = 191) in LGG (brain lower grade glioma) that has a *Z* score of 2.12 (*P*_*Adj*_ = 0.066). *IDH1* mutation is known to be a strong biomarker for improved outcome in LGG patients [33]. The observation that patents with *TP53-M*, *IDH1-M* and *ATRX-MD* have improved PFI compared to patients with *TP53* mutation that do not satisfy the full rule is just a reflection of this single event biomarker. In fact, comparison of patients with the duo *TP53-M* and *IDH1-M* (*n* = 233) against patients with *TP53-M* without *IDH1-M* (*n* = 17), reveals an even stronger association (*Z* = 3.61), confirming that *IDH1* mutation status is responsible for the differential outcomes. In this case the rule does not appear to provide prognostic information that cannot be ascertained from the single event *IDH1-M*. Alternatively, the pairing {*IDH1-M*, *IDH1-M* + *CIC-M*} (*Z* = 2.5, *P*_*Adj*_ = 0.055) appears to tell a very different story, as shown in Figure 8. Among the patients with *IDH1* mutation (*n* = 397), those that also have *CIC* mutation (*n* = 99) have significantly greater PFI than those that are *CIC* wild-type (*n* = 298, Fig. 8A). This result suggests that LGG patients should perhaps be stratified into three groups: *IDH1* wild-type, *IDH1* mutant + *CIC* wild-type, and *IDH1*, *CIC* double mutant (Fig. 8C). This stratification could not have been identified by consideration of individual events.

**Figure 8:**
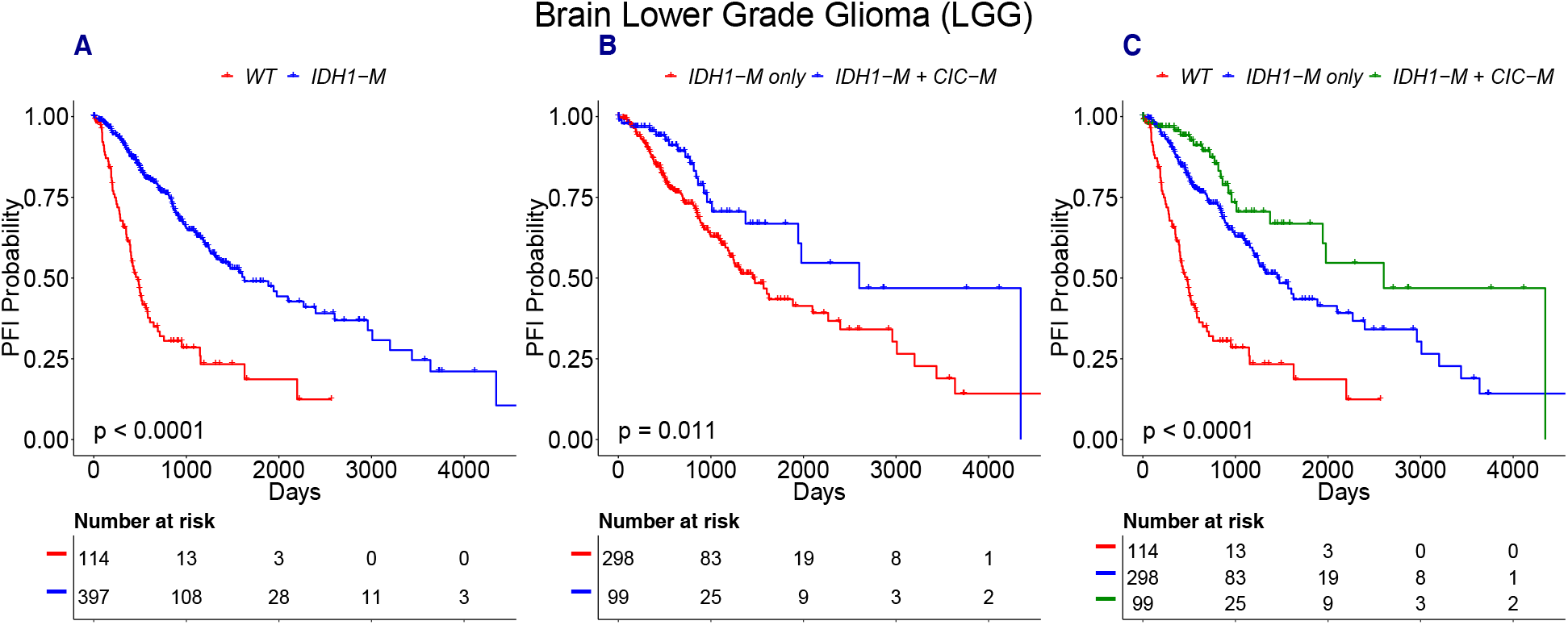
LGG patients with mutations in *CIC* and *IDH1* have longer PFI than patients with *IDH1* mutations who are *CIC* wild type. A) *IDH1* mutations define a subset of patients with longer PFI. B) Patients with *IDH1* and *CIC* have longer PFI than patients with *IDH1* only. C) LGG patients can be stratified into three classes: *IDH1* wild-type, *IDH1* without *CIC* and *IDH1* + *CIC*.

While a subtype defined by co-mutations in *IDH1* and *CIC* has not been described to our knowledge, the LGG TCGA paper [33] identified a subtype of *IDH1* mutant patients who also have a co-deletion in chromosome arms 1p and 19q. This subtype displays longer PFI compared to the patients with *IDH1* mutations that lack the co-deletion. Interestingly we observe that 100% of patients with *CIC* mutation also have the 1p/19q co-deletion, and about 70% of the patients with both *IDH1* mutation and p1/q19 co-deletion also have *CIC* mutations. The relationship between *CIC* mutations and 1p/19q co-deletion suggests that *CIC* mutations define a majority subtype of the patients harboring *IDH1* mutations and 1p/19q co-deletion.

The previous LGG examples show that the results in Table 4 require careful interpretation. We treat these results as a stepping stone for deeper case by case exploration. We highlight two additional unreported examples of differences in PFI that cannot be explained by single event biomarkers and may be useful for predicting patient outcomes

Liver hepatocellular carcinoma (LIHC) patients with both *ARID1A-MD* and *CTNNB1-M* (*n* = 23) have shorter PFI than patients with *ARID1A-MD* but not *CTNNB1-M* (Table 4, *n* = 61, *Z* = −2.89, *P*_*Adj*_ = 0.004). The patients satisfying this rule also have shorter PFI than those with *CTNNB1-M* but not *ARID1A-MD* (*n* = 74, *Z* = −2.27, *P*_*Adj*_ = 0.077). Neither *ARID1A-MD* nor *CTNNB1-M* are biomarkers individually (Fig. 9 A-B), and yet together they define a subtype with significantly worse prognosis (Fig. 9 C-D).

**Figure 9:**
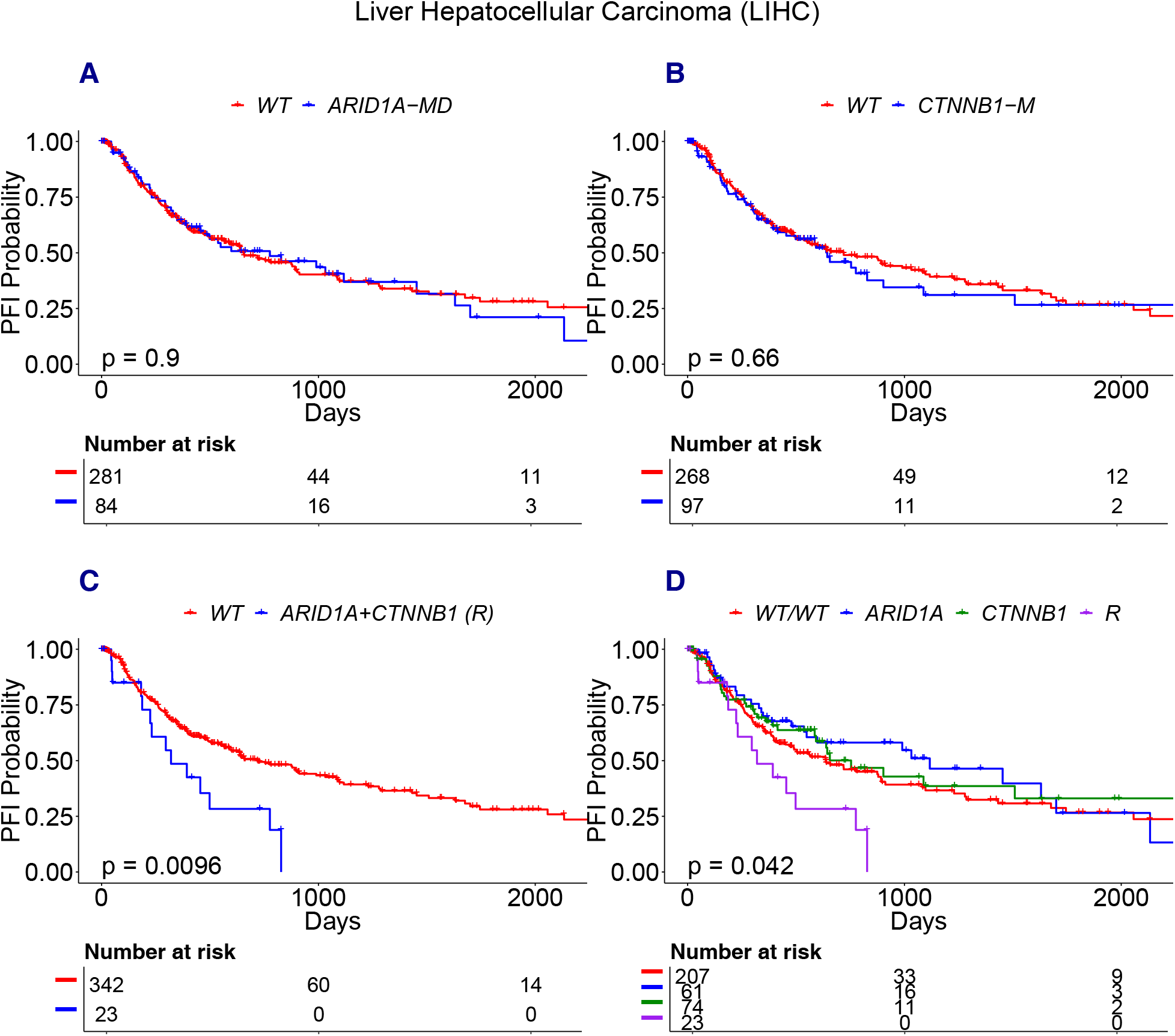
*ARID1A + CTNNB1* define a subtype with shorter PFI in LIHC. A) *ARID1A-MD* is not a biomarker. B) *CTNNB1-M* is not a biomarker. C-D) LIHC patients with *ARID1A-MD* and *CTNNB1-M* define a subtype with significantly worse prognosis compared to all other patients.

Skin-cutaneous melanoma (SKCM) patients with *BRAF-M* and *HIPK2/TBXAS-A* (*n* = 21) have worse PFI than patients with *BRAF-M* but not *HIPK2/TBXAS-A* (Table 4, *n* = 119, *Z* = −2.4, *P*_*Adj*_ = 0.093). Fig. 10A shows that *BRAF* mutant melanoma patients (*n* = 140) have better PFI than *BRAF* wild-type patients (*n* = 140, Kaplan-Meier *P* = .028). The combination of *BRAF-M* + *HIPK2/TBXAS-A* defines a subset of *BRAF-M* mutant patients with poor prognosis (Fig. 10B).

**Figure 10:**
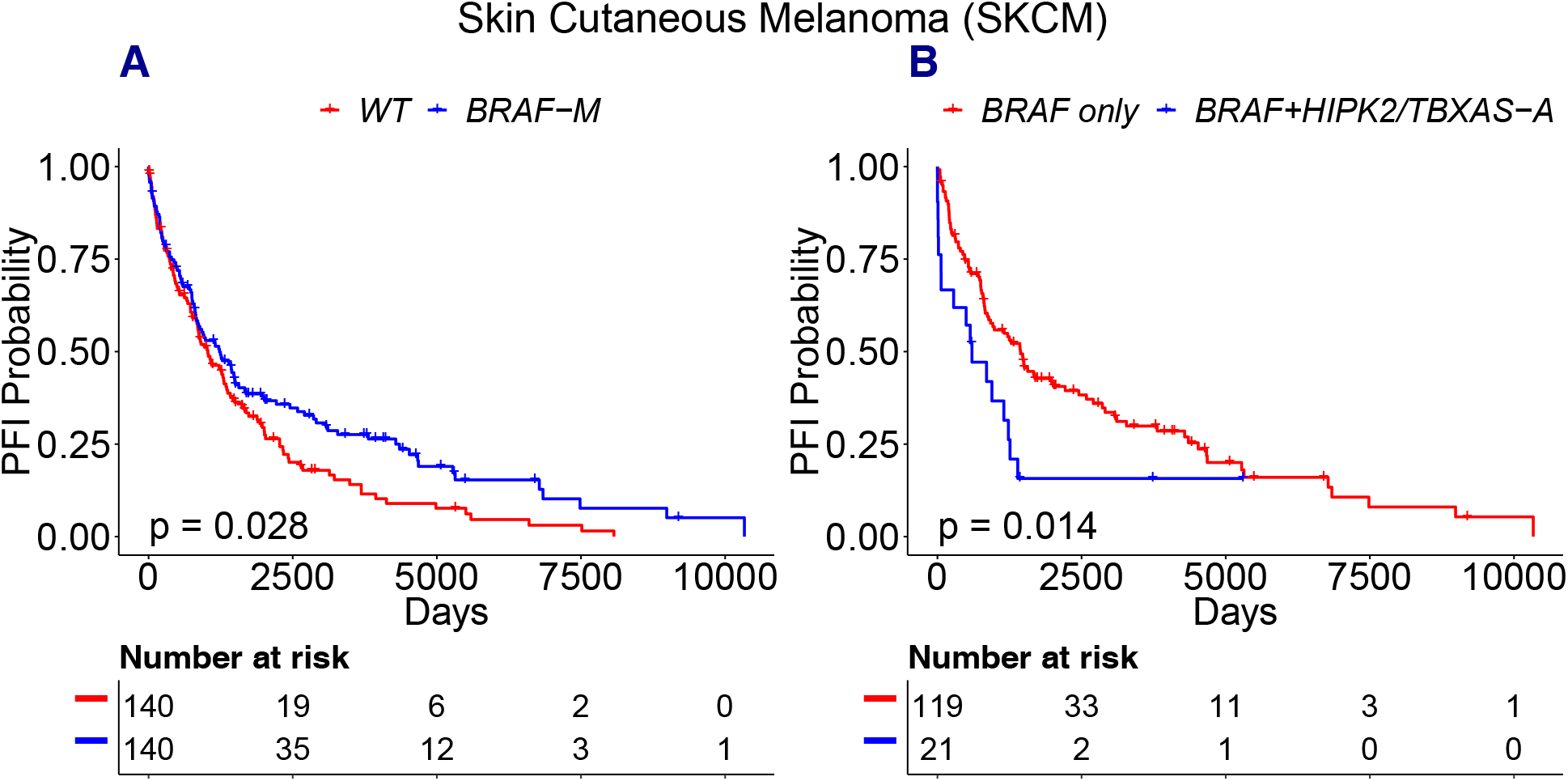
*HIPK2/TBXAS-A* define a subtype of *BRAF* mutant SKCM patients with poor PFI. A) *BRAF* patients have improved PFI compared to non-*BRAF* patients. B) SKCM patients with *BRAF + HIPK2/TBXAS-A* have worse PFI than patients with only *BRAF*.

Combined amplification of *HIPK2/TBXAS* has not been described previously. However, GISTIC2 reports amplified genes either as sets of “narrow peak genes” or as “wide peak genes”. The analyses here use narrow peak gene sets, as the wide peak sets often encompass too many genes for driver analysis. It is noteworthy that the wide peak *HIPK2* amplicon (7q34) includes *BRAF*, so *BRAF* dosage may contribute to outcome.

## 3 Discussion

In this study we developed CRSO as a stochastic optimization procedure for predicting combinations of alterations that drive cancer in individual patients. Ultimately the accuracy of these predictions are best tested with independent patient outcome data, and with mechanistic analysis and animal modeling. Experiments in patient-derived cell lines and animal models could reveal whether specific combinations of alterations are minimally sufficient to transform normal cells into cancer, and could provide a controlled genetic environment to measure the influence of additional alterations. We believe that CRSO can be used to prioritize a small number of combinations in each cancer type for deeper investigation. Specifically, we would recommend prioritizing the high confidence generalized core rules determined by CRSO.

The logic for believing that the generalized core rules identified by CRSO are good candidates to be minimally sufficient driver combinations merits further elaboration. CRSO prioritizes rules by incorporating multiple distinct criteria: 1) passenger probabilities of individual alterations in specific patients, 2) frequency of co-occurrence of combinations across the population, and 3) coverage of the full set of driver combinations across the population. The third criterion is a consequence of the fact that every sample is only assigned to at most one rule. This encourages CRSO to prioritize rules that cover patients who are not better explained by other rules in the rule library. The melanoma results demonstrate that many of the core rules identified by CRSO are mutually exclusive. Unlike methods that explicitly search for mutual exclusivity as the primary criteria for finding driver genes, in CRSO mutual exclusivity emerges naturally as the algorithm seeks to minimize the total penalty of the population using a constrained number of rules.

By simultaneously optimizing for all three of these criteria CRSO identifies rules that would not be prioritized based on frequency or passenger penalties alone. This is exemplified by the prioritization of the rules *NRAS-M + SMYD3-A* in SKCM and *TP53-M + RB1-M* in GBM (Section 2.2). The summary tables of the core rule sets within the supplementary TCGA reports show many additional examples of core rules that are not among the highest ranking rules based on frequency or single rule performance.

### 3.1 Associations of Rules with Patient Outcomes

Examples were presented in LGG, LIHC and SKCM of significant differences in patient PFIs that were found based on rules identified by CRSO. In all three examples the PFI differences could not have been identified by consideration of individual alterations. Rather, these differences appear to be a consequence of specific combinations of alterations who’s co-occurrences define subtypes with better or worse prognosis. These combinations, as well as other combinations in Table 4 that were not discussed, merit further investigation as prognostic biomarkers. Evaluating these findings on independent datasets is a necessary next step toward determining whether any of the combinations are reliable biomarkers that can assist clinical stratification. Incorporating information about patient treatments may help refine this analysis. The interpretability of potential biomarkers identified by CRSO may provide actionable information that goes beyond current clinical stratification, such as suggesting combination treatments or identifying specific alterations as drug targets.

### 3.2 Unassigned Samples

The coverages achieved by the core rule sets in different cancers ranges from a low of 47% in kidney cancer (KIRC), to a high of 83% in ovarian (OV) and uterine (UCEC) cancers, revealing that a substantial subset of patients in every cancer type were not assigned to any rule. The coverages for many of the cancer types appear to converge to a maximum value well before K = 16, as shown in Figure 7. This observation suggests that increasing the rule set size will not substantially increase the number of covered samples. For this reason we interpret the reduced coverage to be evidence that alterations that play an important role in carcinogenesis are likely missing from the input data matrices. One strategy for accounting for these samples would be to expand the set of mutations and SCNVs included as events. This can be accomplished by relaxing the significance threshold used for dNdScv or GISTIC2, or by including additional events identified as candidate drivers using other approaches.

### 3.3 CRSO in Context of Genetic and Transcriptomic Subtypes

In the present analysis we applied CRSO to the full TCGA cohorts for 19 cancer types. The molecular landscapes for nearly all of these cancer types have been extensively studied [1, 2]. These studies revealed distinct molecular subtypes with clinical implications. In some cases the subtypes are based on expression signatures that are difficult to interpret. Comparison of the rule assignments predicted by CRSO with previously established subtypes may help explain the genetic underpinnings of the different subtypes. Additionally, it may be of interest to apply CRSO to previously established subtypes to better understand the within-subtype heterogeneity. In such cases we recommend running dNdScv and GISTIC2 separately on each subtype to identify subtype-specific candidate drivers. Often subtypes have different mutational dynamics. CancerEffectSizeR [29] can precisely calculate passenger mutation probabilities within specific subtypes.

### 3.4 Inclusion of Additional Alteration Types

The cancer rules identified by CRSO are limited by the set of events that are used as inputs. We chose to use the SCNVs and SMGs because these event types are known to be important determinants of patient outcomes in most, if not all, cancer types. This choice, however, meant that some important candidate driver events were missing. Many other types of alterations may contribute to cancer formation, including germ-line alterations, arm level CNVs, gene fusions, chromosomal translocations and epigenetic alterations. For example, in LGG a recurrent p1/q19 chromosomal codeletion has been shown to be a biomarker of prognosis [33]. This event was omitted from our analysis because we chose to exclude arm-level copy number variations. It is possible, however, for users to include this alteration as an event in CRSO. Including unconventional drivers such as p1/q19 chromosomal codeletion poses the challenge of estimating passenger probabilities for such events. One strategy that we suggest would be to assign p1/q19 codeletion a penalty equal to 1.25 times the largest penalty in each sample that harbors this alteration. Doing so would ensure that the event has maximum priority in the samples that harbor it. Using a much larger penalty compared to those associated with SCNVs and mutations could adversely impact the coverage of CRSO by over-prioritizing rules that cover very few samples.

### 3.5 Inclusion of Functional Information

In this study mutations were prioritized based on passenger probabilities. However, it is sometimes possible to predict the functional consequences of specific alterations within candidate driver events. If this information was available, either from experimentation or computational prediction, it would be easy to modify the inputs into CRSO so that high confidence neutral alterations are ignored, i.e., represented as wild type.

### 3.6 Therapeutic Implications of the Cancer Rule Model

The theory that each tumor satisfies minimally sufficient collection of essential drivers would seem to imply that targeting any of these drivers would be sufficient to destroy or inhibit the tumor. How then, do so many tumors resist targeted treatments? In addition to known signaling and epigenetic mechanisms that mediate tumor resiliency, tumors may evolve robustness by accumulating a collection of alterations that satisfy multiple rules. According to this explanation, a logical strategy could be to design combination treatments that inhibit at least one essential driver in all of the rules satisfied by the tumor.

## 4 Conclusion

It has been almost two decades since the advent of next generation DNA sequencing. Projects such as TCGA have produced high quality datasets containing extensive molecular characterization of tens of thousands of cancer patients. Computational tools have been developed to extract information from these data that has translated into improved clinical decision making and improved patient outcomes. Despite all of this progress, it is not possible to explain much of the heterogeneity in patient responses to different treatments. We hope that CRSO will prove helpful in deepening our understanding of driver gene cooperation and personalized cancer treatments.

## 5 Methods

### 5.1 Representation of TCGA Inputs

The inputs into CRSO are genomic alterations that were identified as candidate drivers at the population level. Candidate driver mutations were defined to be the set of significantly mutated genes (SMGs) identified by dNdScv as significantly mutated above expectation using the threshold qsuball *<* 0.1—using tissue-specific mutational covariates developed within [29]. Candidate copy number variations were defined to be the set of genomic regions identified by GISTIC2 as amplified or deleted above expectation, using the threshold q-residual *<* 0.25. All of the TCGA data used were obtained from the January 28, 2016 GDAC Firehose version [34].

Three event types were considered: mutations, amplifications and deletions (not including the special case of hybrid events, see section 5.1.5). Mutations were represented at the gene level, whereas both copy number types were represented at the region level as defined by the GISTIC2 narrow peaks. Multiple types of alterations can be observed for single event. For example, suppose *TP53* is identified as an SMG within a cancer population. Some tumors may contain one of many nonsense point mutations within *TP53*, whereas other tumors may contain a highly recurrent missense mutation, or a splice site mutation that produces an alternative isoform of the TP53 protein. Although all of these alterations are *TP53* mutations, they occur with very different passenger probabilities. To address this, events were subdivided into multiple observation types having distinct passenger probabilities. Copy number events also have different observation types. For example, a particular deletion region can be observed as homozygous deletion or a hemizygous deletion. A tumor sample was said to contain an event if any of the alteration types are observed in the sample, however the penalty associated with the event depends on the specific observation type. This representation reflects the assumption that different types of alterations within the same event are functionally similar but probabilistically distinct.

#### 5.1.1 Mutational Observation Types

The MAF files for each TCGA dataset are annotated with many different mutation types (Table 5). To account for the fact that different kinds of mutations occur at different baseline probabilities, mutations were subdivivded into four observation types: hotspots (HS), loss mutations (L), splicing mutations (S) and in-frame insertions and deletions (I).

**Table 5:**
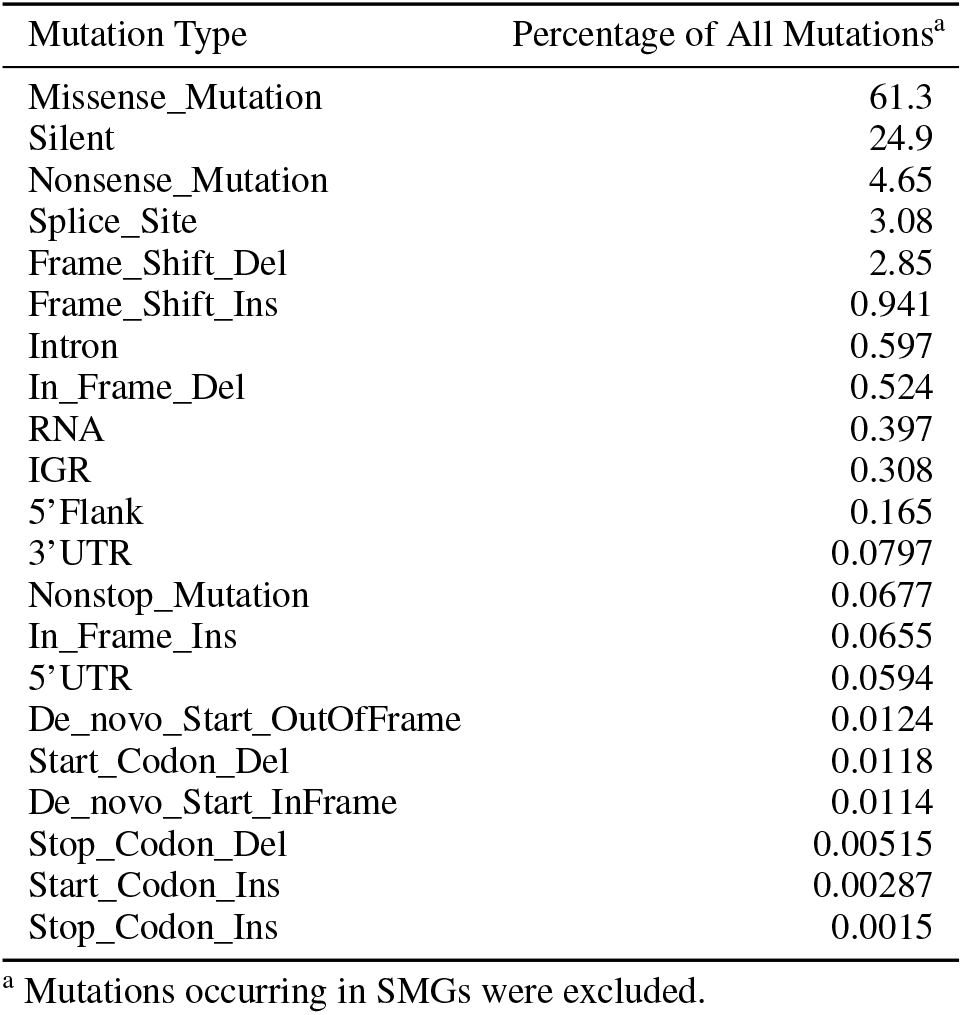
Distribution of mutation types observed across 19 TCGA cancers

##### Hotspot mutations

A hotspot mutation was defined to be any SNP that leads to an alteration at a specific amino acid position that is observed in at least three samples within the population. Silent mutations and intronic mutations do not lead to amino acid changes and by definition cannot be hotspots. Most hotspot mutations are missense mutations, but the definition allows for other recurrent SNPs, such as splice site mutation or nonsense mutations to be hotspots as well. Note that the definition of hotspot does not require three instances of the exact same substitution, but rather three instances of substitutions at the same amino acid position. This choice is motivated by the fact that multiple amino acid changes in known hotspots such BRAFV600 and NRASQ61 are observed.

##### Loss mutations

A loss mutation was defined as one occurring in a given gene if any mutation is detected except for those mutations that are silent, intronic, splice site, hotspot, in-frame insertions or in-frame deletions. The definition of loss mutations includes missense mutations, nonsense mutations, frame-shift indels and the other rarely observed mutations types shown in Table 5. All of these mutation types were combined under the general category of loss mutations because the majority of non-recurrent mutations will lead to loss of function.

##### In-frame Indels

In-frame indels are mutations that are in-frame deletions or in-frame insertions. In-frame insertions/deletions were categorized separately from frame-shift indels because in-frame indels have been shown to be much more likely than frame-shift indels to produce gain of function alterations [35].

##### Splice site mutations

The majority of splice site mutations present as point mutations, and they are the fourth most common class of point mutation behind missense, silent and nonsense mutations (Table 5). Splice site mutations can present as point mutations within exons that lead to exon-exclusion as well as point mutations within introns that lead to intron inclusion. Because of this, many splice site mutations are not annotated with a specific amino acid change. Splice site mutations encompass only those splice sites that are non-recurrent single amino acid substitutions. When a splice site mutation occurs as a SNP at a recurrent amino acid position it was designated as a hotspot mutation because this designation permits more accurate calculation of the associated passenger probabilities.

#### 5.1.2 Mutational Passenger Probabilities

Passenger probabilities were calculated for every observed mutation. These passenger probabilities are patient-specific, gene-specific and observation-type specific. Mutation rates for every possible amino acid substitution in each SMG were calculated as described in Cannataro *et al.* [29]. These rates are calculated by incorporating both gene-level estimates of mutation rate [6] and tumor-type-specific mutational processes that affect nucleotide substitution rates [36].

Hotspot mutation probabilities were calculated for each gene as the sum of the rates of all possible amino acid substitutions at hotspot positions. The loss mutation probability for a gene was calculated as the sum of the rates of all possible amino acid substitutions, except for those that occur at hotspot positions. Non-recurrent splice site substitutions were not excluded because the analysis did not include annotation of all possible splice site amino acid positions. The impact of excluding these sites will be minor, since the number of amino acids per protein is much larger than the number of splice junctions.

Control genes were used to calculate the in-frame indel and splice site probabilities. Control genes were defined to be all genes that are expressed above RSEM = 0 (RNAseq by Expectation Maximization) in at least 5% of samples and were not identified by dNdScv as SMGs. Population level in-frame indel mutation probabilities were calculated to be the frequency of in-frame indel mutations per control gene per sample. Population-level splice site mutation rates were calculated to be the average number of splice site mutations per sample per control gene. The frequencies of in-frame indels and splice site mutations were assumed to be proportional to the number of amino acids in the protein product of each gene. A gene length adjustment factor was defined for each gene to be the number of amino acids in the gene protein product divided by 480 (approximate mean number of amino acids per protein). Gene-specific probabilities for both in-frame indels and splice site mutations were calculated to be the respective population level probabilities multiplied by the gene length adjustment factor.

Mutation frequencies can vary greatly across patients within the same cancer type. To account for this, a patient adjustment factor was used that is based on the number of point mutations in each patient. Mutation counts for each patient were determined to be the total number of point mutations observed outside of the SMGs identified by dNdScv. Patient’s with 0 mutations were assigned a mutation count of 1. In general we do not want to remove outliers with large mutation counts because we want the penalties for observations in these samples to be down weighted accordingly. However, there were a few cases where one or two patients had such extreme outliers that they had mutation counts more than 100 times larger than the 90th percentile for the cohort. To mitigate the impact of these extreme outliers, a maximum mutation count was chosen to be the 10 times the 75th percentile of all mutation counts. Patients with mutation counts above the maximum were assigned the maximum mutation counts. Patient adjustment factors were defined to be the patient’s mutation count divided by the mean mutation count across the population. The patient-specific probabilities for every mutational observation are the product of the population-level probabilities and the patient adjustment factors.

#### 5.1.3 Copy Number Events

The outputs of GISTIC2 are a set of significantly amplified copy number regions and a set of significantly deleted copy number regions. Each of the significant amplifications/deletions were represented as a single event. To do so, the copy number results were first represented at the gene level by a discrete gene-by-sample matrix of focal copy number status, **M_G_**, and then the scores of individual genes within each region were combined to obtain event level features. The entries of **M_G_** take values in {SD,WD,Z,WA,SA}, corresponding respectively to strong deletions (SD), weak deletions (WD), wild type (Z), weak amplifications (WA) and strong amplifications (SA).

**M_G_** was constructed by thresholding the continuous value matrix from “focal_data_by_genes.txt”. Focal copy number values in [−0.3, 0.3] were designated as copy neutral, as per the noise threshold recommendation in the GDC CNV pipeline [37]. Values above 0.3 were designated as amplifications and values below −0.3 were designated as deletions. To designate copy number alterations as strong or weak the sample specific thresholds provided in the file “sample_cutoffs.txt” were used.

The peak genes for each amplification/deletion were extracted from the tables in the files “table_amp.conf_99”/“table_del.conf_99”. For each amplification peak, each sample was assigned to the maximum copy number value attained by any of the peak genes within that sample (i.e., the extreme method). Because the amplification peaks were selected for having evidence of significant amplification, amplification events are only allowed to take values in {Z,WA,SA}. If a deletion is observed within an amplification event it is assigned to be wild type. This procedure results in a discrete matrix of amplification peaks by samples, **M_AMP_**, that takes values in {Z,WA,SA}. Each row in **M_AMP_** corresponds to an amplification event identified by GISTIC2. A deletion event matrix, **M_DEL_**, was prepared analogously. Deletion peaks were assigned to the minimum copy number value attained by any of the genes in the peak. Amplifications observed within deletion peaks were assigned to be wild type, so that **M_DEL_** takes values in {SD,WD,Z}.

#### 5.1.4 Copy Number Passenger Probabilities

It is difficult to estimate copy number probabilities at the gene level because of the strong dependence between genes that are near each other. Copy number passenger probabilities were instead estimated at the cytoband level. Control cytobands were defined to be all cytobands that do not contain any genes that are within any of the significant wide regions reported in “table_amp.conf_99” and “table_del.conf_99”.

A control cytoband matrix, **M_C_**, was constructed by assigning each cytoband to the mode of the cytoband’s genes observed in each sample from **M_G_**. Consider **M_C_** to be an *n* x *m* matrix, and consider *C*_*SD*_, *C*_*W D*_, *C*_*W A*_ and *C*_*SA*_ to be the counts of each observation type observed in the population. A population rate for each observation type was defined as follows:

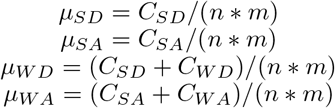

The probabilities for WA and WD were defined to be the frequency of observing any amplification or deletion in order to ensure that *µ*_*W D*_ and *µ*_*W A*_ are always larger than *µ*_*SD*_ and *µ*_*SA*_, respectively.

To account for variation in copy number rates between patients patient-specific adjustment factors were introduced for amplifications and deletions. Let 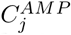 and 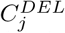 denote the number of control cytobands amplified and deleted in patient *j*, respectively. The amplification and deletion adjustment factors for each patient were defined to be:

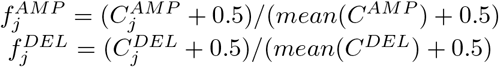

The addition of 0.5 to the counts of all patients ensures non-zero probabilities. The sample specific probabilities for SD and WD were respectively given by: 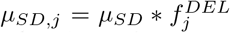 and 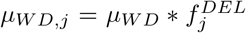. The sample specific probabilities for SA and WA were calculated analogously.

#### 5.1.5 Hybrid Events

When a particular gene within a tissue type is identified by dNdScv as an SMG and is also identified by GISTIC2 as part of a SCNV, the SMG and SCNV were combined into a special event type called hybrid events. This reflects the assumption that the SCNV and SMG are exerting similar functional changes in the tumor cells. Supporting this assumption is the observation that oncogenes are frequently amplified and tumor suppressors are frequently deleted in the same cancer types. If the SCNV was a deletion the hybrid event was denoted as “gene-MD”, for mutations/deletion, and if it was an amplification it was denoted as “gene-MA”, for mutation/amplification. The hybrid events could take values in any of the mutation types or any of the copy number types. A single hybrid event could also take two values if it was observed as an SCNV and an SMG in the same patient. For example, if *CDKN2A* loss mutation and *CDKN2A* weak deletion co-occur in a patient, this observation would have been denoted as “L,WD”. In such cases the penalty associated with the combined observation was the sum of the penalties of each observation independently (recall that penalties are −log probabilities). By increasing the penalty associated with co-occuring alterations of the same gene, the algorithm is encouraged to assign the event as a driver.

Two exceptions were encountered among the 19 TCGA cancer types that required special handling. In rectum adenocarcinoma (READ) *KRAS* was identified as part of an amplification peak and as part of a large deletion peak. Since *KRAS* is a known oncogene that is often part of amplification peaks in other cancer types we chose to ignore the deletion peak and represent *KRAS* as *KRAS-MA*. In kidney renal clear cell carcinoma (KIRC) both *ARID1* and *MTOR* were found to be part of the same deletion peak. Since *ARID1-MD* is observed in multiple cancer types whereas *MTOR-MD* is never observed in other cancer types, we chose to represent this deletion event as part of *ARID1-MD*.

### 5.2 CRSO Algorithm

This section presents the CRSO algorithm. CRSO involves several parameter choices that require specification. The methodology is presented using the default parameter values (Table 6) that were used for the presented applications to 19 TCGA cancer types.

**Table 6:** Default Parameters

| Parameter | Default Value | Description |
| --- | --- | --- |
| rule.cov.thresh | 3% (minimum 6 samples) | Min rule coverage |
| max.lib.size | 1000 | Max # rules in rule library |
| p1.spr | 100 | Random sets per rule in phase 1, for each sample size |
| p1.ss.vec | $v = [8, 12, 16, 20]$ when $> 200$ rules, $v = [6, 8, 10, 12]$ when $\leq 200$ rules | Sample sizes used in phase 1 for random sampling |
| p1.cut.size | 15% when $> 300$ rules, 10% when $\leq 300$ rules | Rules eliminated in each phase 1 iteration |
| p1.stop | 20 | Stop point for phase 1 |
| p2.k.max | 16 | Max K considered in phase 2 |
| p2.max.nrs | 200,000 | Max # rule sets evaluated for each K in phase 2 |
| p2.max.compute | 10 million | Max # non constrained rule sets considered for evaluation for each K in phase 2 |
| p3.max.nrs | 1 million | Max # rule sets considered for each K in phase 3 |
| min.rs.assign.thresh | 3% (minimum 6 samples) | Min rule assignment |
| core.cov.thresh | 95% | Coverage (relative to max) criteria for choosing core RS |
| core.perf.thresh | 90% | Performance (relative to max) criteria for choosing core RS |

#### 5.2.1 CRSO Input Matrices

The CRSO inputs are represented as two event-by-sample matrices, a binary alteration matrix **D** and passenger penalty matrix **P**. The entries of **D** are binary and indicate whether a particular event has happened in a particular patient. The entries of **P** are the negative log of the passenger probabilities associated with each observation in **D**. When an entry of **D** is zero, the corresponding entry in **P** is assigned to 0.

#### 5.2.2 CRSO Objective Function

The CRSO algorithm seeks to minimize the statistical penalty of the observed data under a proposed rule set. Consider a rule set *RS* = (*r*_1_, …, *r*_*k*_), and suppose each sample has been assigned to one rule in *RS*, or to the null rule. When a sample is assigned to a particular rule, the events of that rule are considered drivers within that sample. All non-driver events are assumed to be passenger events. In general there are multiple possible assignments for a given rule set, since some tumor samples may satisfy more than one rule in *RS*. We first formulate the objective function assuming each sample is assigned according to a single assignment, and then we show that it is easy to determine the best possible assignment for any rule set.

The statistical penalty of the data under a proposed rule assignment is defined to be the sum of the penalties of all of the unassigned events in the dataset. Every rule set is associated with a penalty matrix **P^RS^**. **P^RS^** is derived by modifying the full penalty matrix, **P**, such that the penalties of all assigned events are changed to 0. For example, suppose sample *j* is assigned to a rule containing events *x* and *y*. This is represented in **P^RS^** by assigning 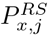 and 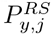 to be 0, instead of the original values they took in **P**. The statistical penalty of *RS* is defined to be:

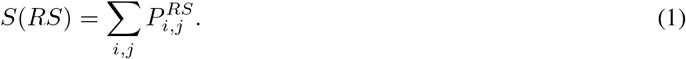

Under the null rule set every observed event is assumed to be a passenger, so that **P^0^** = **P**. The statistical penalty for the null rule set is given by *S*_0_ = ∑*P*_*i,j*_. The objective function, *J*(*RS*), is defined as:

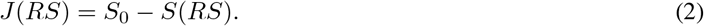

The CRSO algorithm seeks to identify the rule set of fixed size *K* that maximizes the objective function, *J* (*RS*), by minimizing the statistical penalty *S*(*RS*). Fixing the size of the rule set enforces competition between rules based on the number of events covered per sample, the rarity of events covered and the number of samples covered.

In general there are many possible assignments due to samples satisfying multiple rules in *RS*. The objective function score, *J* (*RS*), may be different for different assignments. Because *J* (*RS*) is a simple sum over the unassigned penalties, the optimal assignment is determined by assigning each sample to the rule that decreases the total penalty of that sample by the largest amount.

#### 5.2.3 Three Phase Algorithm

The three phase procedure seeks to find the best scoring rule set of a fixed size *K*. In general larger rule sets perform better than smaller rule sets because the samples in **D** have more assignment opportunities. On the other hand larger rule sets may be more likely to reflect noise in the dataset rather than true biological signal, i.e., over-fitting. To balance these considerations CRSO seeks the best rule set of size *K*, for *K* ∈ {1 … 16}. The overall best rule set is subsequently chosen from among the 16 best rule sets of size *K*. The choice of capping the best rule set size at *K* = 16 reflects the assumption that there are fewer than 16 driver rules for a given cancer type. Supporting this assumption, the best rule set coverage appears to converge before *K* = 16 for most of the 19 TCGA cancer types (Figure 7). In case there are more than 16 driver rules, CRSO can be applied to more homogeneous subsets of the data.

Given a binary-input matrix **D**, a starting rule library is built by identifying all rules that contain at least 2 events and occur in a minimum percentage of samples. A minimum rule coverage threshold is chosen to be either 3% of the population, or the minimum threshold that at most 1000 rules satisfy, depending on which threshold is larger. Since there will generally be a large number of rules in the starting rule library it is impossible to exhaustively evaluate every possible rule set, even for small *K*. The computational time of exhaustive evaluation grows exponentially according to the size of the rule pool, i.e., *O*(2^*n*^) for *n* rules. To address this computational limitation, phase 1 is used to prioritize rules according to how likely they are to be among the best rule set. In phase 2, a subset of the top-performing rules determined from phase 1 are exhaustively evaluated. The number of rules that are exhaustively evaluated is subject to a computational constraint on the number of total rule-sets that are considered for each *K*. In general a larger number of rules can be exhaustively evaluated for small *K*. The number of rules that can be exhaustively considered is generally small for *k >* 4, meaning that only the top 20-30 rules have the opportunity to be among the best rule sets. Since this subset may be too restrictive, the algorithm is exposed to additional rules in phase 3 by considering rulesets that involve a small number (e.g., 1-3) of unexplored rules within rule sets that overlap with the current best rule set.

##### Phase 1: Stochastic Rule Prioritization

In phase 1, an iterative stochastic procedure is used to rank all of the rules in the rule library according to how likely they are to be included in the best performing rule set. For each rule, a *rule importance score* is calculated based on the average contribution of the rule within many random subsets of rules. Consider a set of rules *RS*. The contribution of rule *r*_*j*_ within *RS* is defined to be the percentage decrease in performance when *r*_*j*_ is excluded from *RS*. To determine the rule importance score, multiple sampling sizes of rules are evaluated separately in order to make fair comparisons between rules across a broad range of rule sets. For each sample size a *Z* score is determined from the distribution of average contributions, and the rule importance score is determined to be the average of the *Z* scores from different sampling sizes. Once the importance scores are obtained for each rule in the rule pool, the 10% of rules (phase 1 cut size) that have the smallest contribution scores are eliminated. The procedure proceeds until there are at most 20 rules, at which point the remaining rules are ranked according to a final round of importance score calculation.

Two important design choices for phase 1 merit further elaboration. First, family members are allowed to be chosen within the same random sets of rules. This choice permits competition between all of the rules, including rules that are family members. Second, the reader may wonder why an iterative procedure is necessary instead of determining rule importance scores once for the full rule library and ranking the rules accordingly. The reason here too is to enforce direct, fair competition among the strongest scoring rules. The starting rule library will generally contain many rules that cover a small fraction of patients and are unlikely to be part of the best performing rulesets. The contribution scores of a rule will vary greatly according to the quality of the other rules in the rule set, i.e., a relatively unimportant rule can appear very important when sampled along with inferior rules. As weak rules are eliminated, the competition level of the remaining rules increase, leading to more fair comparisons among the strong rules. Experimentation shows that the iterative procedure leads to much more stable ordering compared to ranking rules in a single iteration.

##### Phase 2: Exhaustive rule set evaluation

In phase 2, a subset of the top rules from phase 1 are exhaustively evaluated to determine the best rule set of each *K*. In contrast to phase 1, only valid rule sets that do not contain family members are considered in phase 2. For each *K* the candidate rule pool is determined to be the maximum number of top rules that can be exhaustively evaluated with at most 200,000 rule sets. This computational parameter was chosen to balance run time with depth of coverage.

##### Phase 3: Neighbor Rule Set Expansion

Phase 2 results in identification of best rules of sizes *K* = 1 … 16. However, because of computational constraints only a subset of top rules can be considered by phase 2 for inclusion among the best size-*K* rule sets. In phase 3 the number of rules that can be included in the top rule sets is increased by making the assumption that the global best size-*K* rule sets will overlap highly with the size-*K* rule sets determined from phase 2. Consider as an example that in phase 2 the best rule set of size *K* = 8 is identified from among the top *n* = 30 rules. Denote this rule set as *RS_K_*. The *d*_*L*_ neighbors of *RS*_*K*_ are defined to be the set of all rule sets that contain *K − L* common rules with *RS*_*K*_. In phase 3 the search for the best performing rule sets is expanded to include rule sets that are *d*_*L*_ neighbors of *RS*_*K*_ and contain *L* rules from outside of the initial top 30 rule pool, for *L* = 1, 2, 3. For each *L* the number of new rules that can be considered is determined subject to a computational constraint. A similar expansion of candidate rule sets is performed by considering rule sets that overlap highly with *RS*_*K−*1_, allowing for consideration of new rule sets of size *K* that contain rules that are within *RS*_*K−*1_ but absent from *RS*_*K*_. This choice is motivated by the observation the best rule set of size *K* tends to overlap highly with the best rule set of size *K −* 1.

### 5.3 Software Details

CRSO was developed using R. RMarkdown was used for automatic generation of the TCGA reports. Many of the figures were generated with “ggplot2” package [38]. Survival analysis was performed using the “survival” package [39]. Kaplan-Meier plots were generated using the “survminer” package [40]. Parallelization was performed using the “foreach” [41] and “doMPI” [42] packages.

## 6 CRSO Availability

The CRSO R package is freely available for download at https://cran.r-project.org/web/packages/crso/.

## Supporting information

TCGA Reports

## 7 Acknowledgements

The results published here are based on data generated by the TCGA Research Network: http://cancergenome.nih.gov/ [43]. The authors thank Dr. Michael Kane for his help with developing the CRSO software package. The authors thank Dr. Christos Hatzis for his counsel.

HZ was partially supported by NIH grants P30CA016359 and P50CA1965305.

## 8 Author Contributions

MK, DS and HZ conceived of the project. MK programmed the CRSO software and drafted the manuscript. VC and JT provided SNV mutation rates and dNdScv values that were used to calculate TCGA passenger probabilities. JT, DS and HZ edited the manuscript. All authors read and approved the final manuscript.

## References

[1] John N. Cancer Genome Atlas Research Network, John N Weinstein, Eric A Collisson, Gordon B Mills, Kenna R Mills Shaw, Brad A Ozenberger, Kyle Ellrott, Ilya Shmulevich, Chris Sander, and Joshua M Stuart. The Cancer Genome Atlas Pan-Cancer analysis project. Nature genetics, 45(10):1113–20, oct 2013.

[2] Katarzyna Tomczak, Patrycja Czerwińska, and Maciej Wiznerowicz. Review The Cancer Genome Atlas (TCGA): an immeasurable source of knowledge. Współczesna Onkologia, 1A(1A):68–77, 2015.

[3] J. Zhang, J. Baran, A. Cros, J. M. Guberman, S. Haider, J. Hsu, Y. Liang, E. Rivkin, J. Wang, B. Whitty, M. Wong-Erasmus, L. Yao, and A. Kasprzyk. International Cancer Genome Consortium Data Portal–a one-stop shop for cancer genomics data. Database, 2011(0):bar026–bar026, sep 2011.

[4] Frank McCormick. Targeting KRAS Directly. Annual Review of Cancer Biology, 2(1):81–90, mar 2018.

[5] Michael S Lawrence, Petar Stojanov, Paz Polak, Gregory V Kryukov, Kristian Cibulskis, Andrey Sivachenko, Scott L Carter, Chip Stewart, Craig H Mermel, Steven A Roberts, Adam Kiezun, Peter S Hammerman, Aaron McKenna, Yotam Drier, Lihua Zou, Alex H Ramos, Trevor J Pugh, Nicolas Stransky, Elena Helman, Jaegil Kim, Carrie Sougnez, Lauren Ambrogio, Elizabeth Nickerson, Erica Shefler, Maria L Cortés, Daniel Auclair, Gordon Saksena, Douglas Voet, Michael Noble, Daniel DiCara, Pei Lin, Lee Lichtenstein, David I Heiman, Timothy Fennell, Marcin Imielinski, Bryan Hernandez, Eran Hodis, Sylvan Baca, Austin M Dulak, Jens Lohr, Dan-Avi Landau, Catherine J Wu, Jorge Melendez-Zajgla, Alfredo Hidalgo-Miranda, Amnon Koren, Steven A McCarroll, Jaume Mora, Brian Crompton, Robert Onofrio, Melissa Parkin, Wendy Winckler, Kristin Ardlie, Stacey B Gabriel, Charles W M Roberts, Jaclyn A Biegel, Kimberly Stegmaier, Adam J Bass, Levi A Garraway, Matthew Meyerson, Todd R Golub, Dmitry A Gordenin, Shamil Sunyaev, Eric S Lander, and Gad Getz. Mutational heterogeneity in cancer and the search for new cancer-associated genes. Nature, 499(7457):214–218, jul 2013.

[6] Iñigo Martincorena, Keiran M Raine, Moritz Gerstung, Kevin J Dawson, Kerstin Haase, Peter Van Loo, Helen Davies, Michael R Stratton, and Peter J Campbell. Universal Patterns of Selection in Cancer and Somatic Tissues. Cell, 171(5):1029–1041.e21, nov 2017.

[7] Craig H Mermel, Steven E Schumacher, Barbara Hill, Matthew L Meyerson, Rameen Beroukhim, and Gad Getz. GISTIC2.0 facilitates sensitive and confident localization of the targets of focal somatic copy-number alteration in human cancers. Genome Biology, 12(4):R41, 2011.

[8] Michael S. Lawrence, Petar Stojanov, Craig H. Mermel, James T. Robinson, Levi A. Garraway, Todd R. Golub, Matthew Meyerson, Stacey B. Gabriel, Eric S. Lander, and Gad Getz. Discovery and saturation analysis of cancer genes across 21 tumour types. Nature, 505(7484):495–501, jan 2014.

[9] S. De and S. Ganesan. Looking beyond drivers and passengers in cancer genome sequencing data. Annals of Oncology, 28(5):mdw677, dec 2016.

[10] Francesc Castro-Giner, Peter Ratcliffe, and Ian Tomlinson. The mini-driver model of polygenic cancer evolution. Nature Reviews Cancer, 15(11):680–685, nov 2015.

[11] Cristian Tomasetti, Luigi Marchionni, Martin A Nowak, Giovanni Parmigiani, and Bert Vogelstein. Only three driver gene mutations are required for the development of lung and colorectal cancers. Proceedings of the National Academy of Sciences of the United States of America, 112(1):118–23, jan 2015.

[12] Nathan L Nehrt, Thomas A Peterson, DoHwan Park, and Maricel G Kann. Domain landscapes of somatic mutations in cancer. BMC Genomics, 13(Suppl 4):S9, jun 2012.

[13] Jack P Hou and Jian Ma. DawnRank: discovering personalized driver genes in cancer. Technical report, 2014.

[14] Chengliang Dong, Yunfei Guo, Hui Yang, Zeyu He, Xiaoming Liu, and Kai Wang. iCAGES: integrated CAncer GEnome Score for comprehensively prioritizing driver genes in personal cancer genomes.

[15] Christos M Dimitrakopoulos and Niko Beerenwinkel. Computational approaches for the identification of cancer genes and pathways. WIREs Syst Biol Med, 9:1364, 2017.

[16] Christopher A Miller, Stephen H Settle, Erik P Sulman, Kenneth D Aldape, and Aleksandar Milosavljevic. Discovering functional modules by identifying recurrent and mutually exclusive mutational patterns in tumors. BMC Medical Genomics, 4(1):34, dec 2011.

[17] Mark Dm Leiserson, Hsin-Ta Wu, Fabio Vandin, and Benjamin J Raphael. CoMEt: a statistical approach to identify combinations of mutually exclusive alterations in cancer. Genome Biology, 16:160, 2015.

[18] Mark D. M. Leiserson, Dima Blokh, Roded Sharan, and Benjamin J. Raphael. Simultaneous Identification of Multiple Driver Pathways in Cancer. PLoS Computational Biology, 9(5):e1003054, may 2013.

[19] Jack P. Hou, Amin Emad, Gregory J. Puleo, Jian Ma, and Olgica Milenkovic. A new correlation clustering method for cancer mutation analysis. Bioinformatics, 32(24):3717, 2016.

[20] Hao Wu, Lin Gao, Feng Li, Fei Song, Xiaofei Yang, and Nikola Kasabov. Identifying overlapping mutated driver pathways by constructing gene networks in cancer. BMC Bioinformatics, 16(Suppl 5):S3, 2015.

[21] Yahya Bokhari and Tomasz Arodz. QuaDMutEx: quadratic driver mutation explorer. 18:458, 2017.

[22] Bo Gao, Guojun Li, Juntao Liu, Yang Li, and Xiuzhen Huang. Identification of driver modules in pan-cancer via coordinating coverage and exclusivity. Oncotarget, 8(22):36115–36126, may 2017.

[23] Sander Canisius, John W. M. Martens, and Lodewyk F. A. Wessels. A novel independence test for somatic alterations in cancer shows that biology drives mutual exclusivity but chance explains most co-occurrence. Genome Biology, 17(1):261, dec 2016.

[24] Junhua Zhang, Ling-Yun Wu, Xiang-Sun Zhang, and Shihua Zhang. Discovery of co-occurring driver pathways in cancer. BMC Bioinformatics, 15(1):271, dec 2014.

[25] Pi-Jing Wei, Di Zhang, Junfeng Xia, and Chun-Hou Zheng. LNDriver: identifying driver genes by integrating mutation and expression data based on gene-gene interaction network. BMC Bioinformatics, 17(S17):467, dec 2016.

[26] Yong Chen, Jingjing Hao, Wei Jiang, Tong He, Xuegong Zhang, Tao Jiang, and Rui Jiang. Identifying potential cancer driver genes by genomic data integration. Scientific reports, 3:3538, dec 2013.

[27] Sajal Dash, Nicholas A. Kinney, Robin T. Varghese, Harold R. Garner, Wu-chun Feng, and Ramu Anandakrishnan. Differentiating between cancer and normal tissue samples using multi-hit combinations of genetic mutations. Scientific Reports, 9(1):1005, dec 2019.

[28] V. Chvatal. A Greedy Heuristic for the Set-Covering Problem. Mathematics of Operations Research, 4(3):233–235, aug 1979.

[29] Vincent L Cannataro, Stephen G Gaffney, and Jeffrey P Townsend. Effect Sizes of Somatic Mutations in Cancer. JNCI: Journal of the National Cancer Institute, 110(11):1171–1177, nov 2018.

[30] The Cancer Genome Atlas Cancer Genome Atlas Network. Genomic Classification of Cutaneous Melanoma. Cell, 161(7):1681–96, jun 2015.

[31] Joan C Smith and Jason M Sheltzer. Systematic identification of mutations and copy number alterations associated with cancer patient prognosis. eLife, 7, dec 2018.

[32] Jianfang Liu, Tara Lichtenberg, and Katherine A Hoadley. An Integrated TCGA Pan-Cancer Clinical Data Resource to Drive High-Quality Survival Outcome Analytics In Brief Analysis of clinicopathologic annotations for over 11,000 cancer patients in the TCGA program leads to the generation of TCGA Clinical Data Reso. Cell, 173:400–416.e11, 2018.

[33] The Cancer Genome Atlas Research Network. Comprehensive, Integrative Genomic Analysis of Diffuse Lower-Grade Gliomas. New England Journal of Medicine, 372(26):2481–2498, jun 2015.

[34] doi:10.7908/C11G0KM9.

[35] Haiwang Yang, Yan Zhong, Cheng Peng, Jian-Qun Chen, and Dacheng Tian. Important role of indels in somatic mutations of human cancer genes. BMC medical genetics, 11:128, sep 2010.

[36] Rachel Rosenthal, Nicholas McGranahan, Javier Herrero, Barry S. Taylor, and Charles Swanton. deconstructSigs: delineating mutational processes in single tumors distinguishes DNA repair deficiencies and patterns of carcinoma evolution. Genome Biology, 17(1):31, dec 2016.

[37] https://docs.gdc.cancer.gov/Data/Bioinformatics_Pipelines/CNV_Pipeline/.

[38] Hadley Wickham. ggplot2: Elegant Graphics for Data Analysis. Springer-Verlag New York, 2016.

[39] Terry M Therneau. A Package for Survival Analysis in S, 2015. version 2.38.

[40] https://CRAN.R-project.org/package=survminer.

[41] https://CRAN.R-project.org/package=foreach.

[42] https://CRAN.R-project.org/package=doMPI.

[43] Robert L. Grossman, Allison P. Heath, Vincent Ferretti, Harold E. Varmus, Douglas R. Lowy, Warren A. Kibbe, and Louis M. Staudt. Toward a Shared Vision for Cancer Genomic Data. New England Journal of Medicine, 375(12):1109–1112, sep 2016.

