## Supplementary material for "Identifying Combinations of Cancer Drivers in Individual Patients": TCGA Reports: CRSO_Report_BLCA.pdf

### CRSO Output Report: BLCA

Michael Klein

Apr 19 2019

**4 Generalized Core Rules** **7**

**5 Dictionary of Copy Number Events** **8**

#### 1 Heatmaps of D and P

**Number of samples = 392.**
**Number of events = 113.**
**Rule coverage requirement = 12 samples.**
**Rule library size = 933 rules.**

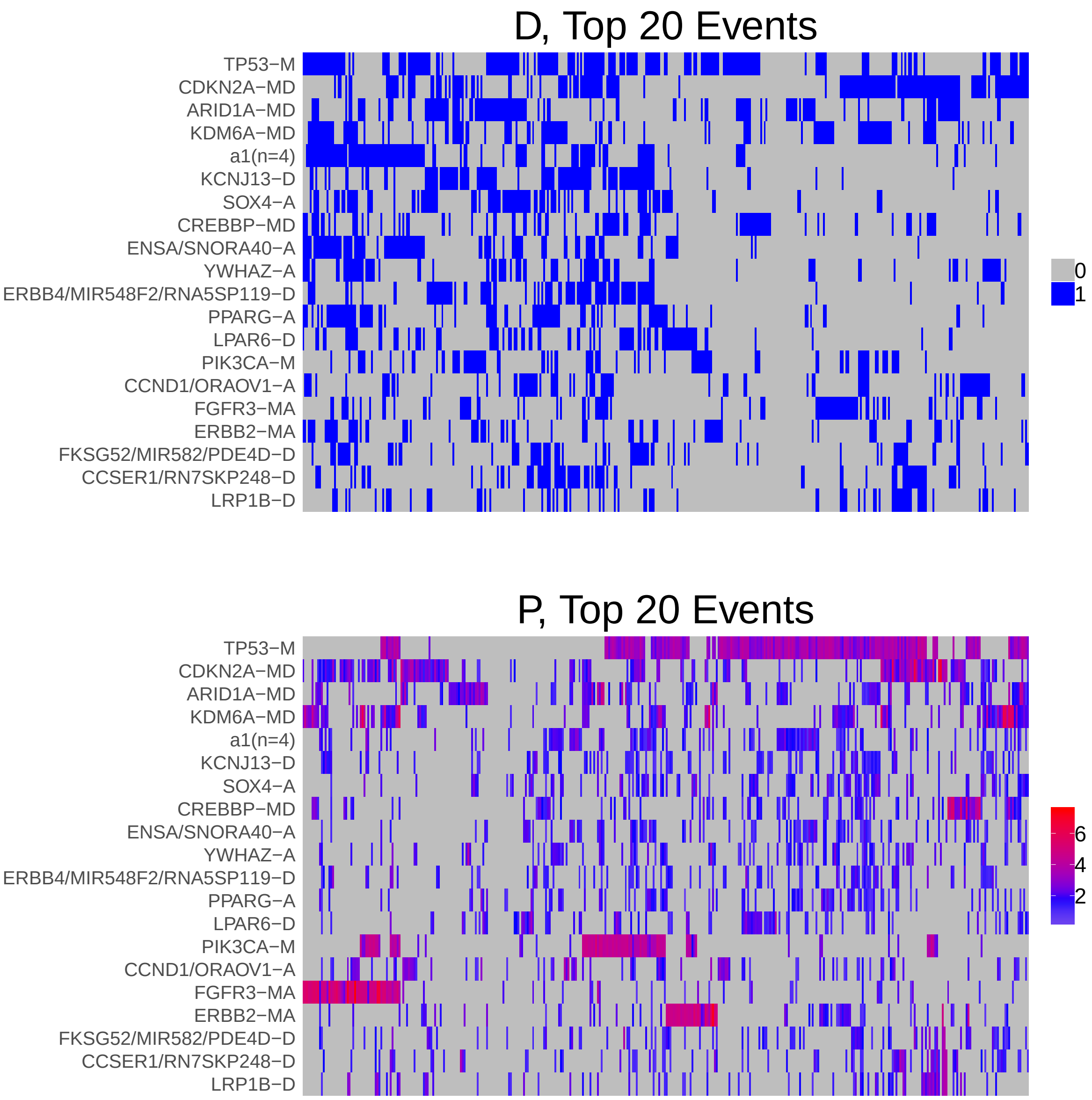

Figure 1: Heatmap of D and P. Events are ordered by frequency, from top to bottom. Event suffix -M: mutation, -A: amplification, -D: deletion, -MD: mutDel, -MA: mutAmp. Samples are ordered using hierarchical clustering. Wild-type events indicated in grey.

### 2 Summary of K Best Rule Sets

#### 2.1 Performance and coverage of best rule sets

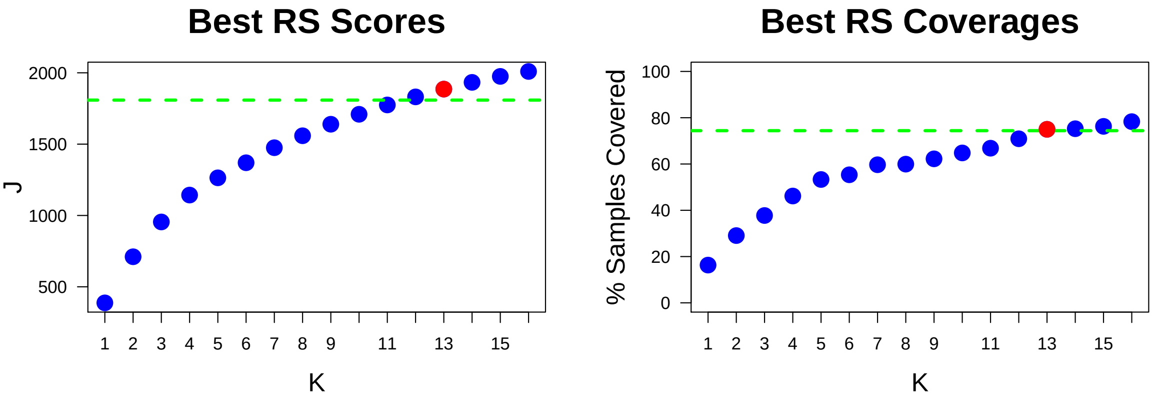

Figure 2: Performance and Coverage of Best Filtered RS for K. The core was determined to be the smallest RS that achieved at least 95% of maximum coverage and at least 90% of maximum performance.

#### 2.2 Table of rules that appear in any best rule set

| ID | Rule | PC | Ks |
|---|---|---|---|
| r4 | CDKN2A-MD + TP53-M | 16 | 1,2,3,4,5,6,7,8,9,10,11,12,13,14,15,16 |
| r1 | RB1-M + TP53-M | 14 | 2,3,4,5,6,7,8,9,10,11,12,13,14,15,16 |
| r2 | CDKN2A-MD + FGFR3-MA | 11 | 4,5,6,7,8,9,10,11,12,13,14,15,16 |
| r3 | FGFR3-MA + KDM6A-MD | 11 | 3,7,8,9,10,11,12,13,14,15,16 |
| r5 | PIK3CA-M + TP53-M | 12 | 6,7,8,9,10,11,12,13,14,15,16 |
| r6 | ERBB2-MA + TP53-M | 13 | 8,9,10,11,12,13,14,15,16 |
| r8 | ARID1A-MD + PIK3CA-M | 8.2 | 9,10,11,12,13,14,15,16 |
| r11 | CDKN2A-MD + KDM6A-MD | 15 | 9,10,11,12,13,14,15,16 |
| r57 | ARID1A-MD + SOX4-A + TP53-M | 7.1 | 8,9,10,11,12,14,15,16 |
| r7 | a1(n=4) + ENSA/SNORA40-A + TP53-M | 13 | 11,12,13,14,15,16 |
| r25 | CDKN2A-MD + NFE2L2-M | 5.6 | 10,11,12,13,14 |
| r12 | ARID1A-MD + CDKN2A-MD | 13 | 5,6,7,8 |
| r17 | ERBB4/MIR548F2/RNA5SP119-D + KCNJ13-D | 19 | 13,14,15,16 |
| r43 | CASC8-A + YWHAZ-A | 13 | 12,13,14,15 |
| r10 | KDM6A-MD + TP53-M | 17 | 4,5,6 |
| r62 | CREBBP-MD + KDM6A-MD + TP53-M | 5.4 | 14,15,16 |
| r41 | CCSER1/RN7SKP248-D + CDKN2A-MD + LRP1B-D | 5.6 | 15,16 |
| r73 | ATM-M + CDKN2A-MD | 6.4 | 15,16 |
| r9 | SOX4-A + TP53-M | 17 | 7 |
| r32 | ARID1A-MD + YWHAZ-A | 11 | 16 |
| r36 | ARID1A-MD + KDM6A-MD + TP53-M | 5.9 | 13 |
| r124 | ENSA/SNORA40-A + LPAR6-D | 10 | 16 |

**ID** = Rule IDs, rules are numbered according to importance rank determined from phase 1
**PC** = Percent of samples covered **Ks** = Membership in best RS

### 3 Core Rule Set

**Core K = 13.**
**Core rule set coverage = 75%.**

#### 3.1 Table of core rule set rules

| ID | Rule | CR | SJR | SJ | NSC | NSA | PC | PA | FracA |
|---|---|---|---|---|---|---|---|---|---|
| r1 | RB1-M + TP53-M | 15.5 | 3 | 343 | 54 | 32 | 14 | 8.2 | 0.59 |
| r2 | CDKN2A-MD + FGFR3-MA | 34 | 17 | 276 | 44 | 23 | 11 | 5.9 | 0.52 |
| r3 | FGFR3-MA + KDM6A-MD | 42.5 | 12 | 287 | 42 | 25 | 11 | 6.4 | 0.60 |
| r4 | CDKN2A-MD + TP53-M | 6 | 1 | 388 | 64 | 27 | 16 | 6.9 | 0.42 |
| r5 | PIK3CA-M + TP53-M | 27 | 6 | 330 | 46 | 34 | 12 | 8.7 | 0.74 |
| r6 | ERBB2-MA + TP53-M | 21.5 | 10 | 300 | 50 | 21 | 13 | 5.4 | 0.42 |
| r7 | a1(n=4) + ENSA/SNORA40-A + TP53-M | 21.5 | 8 | 302 | 50 | 24 | 13 | 6.1 | 0.48 |
| r8 | ARID1A-MD + PIK3CA-M | 96.5 | 47 | 189 | 32 | 18 | 8.2 | 4.6 | 0.56 |
| r11 | CDKN2A-MD + KDM6A-MD | 11 | 15 | 278 | 59 | 19 | 15 | 4.8 | 0.32 |
| r17 | ERBB4/MIR548F2/RNA5SP119-D + KCNJ13-D | 1 | 28 | 217 | 75 | 20 | 19 | 5.1 | 0.27 |
| r25 | CDKN2A-MD + NFE2L2-M | 337 | 127.5 | 144 | 22 | 15 | 5.6 | 3.8 | 0.68 |
| r36 | ARID1A-MD + KDM6A-MD + TP53-M | 293.5 | 56 | 182 | 23 | 18 | 5.9 | 4.6 | 0.78 |
| r43 | CASC8-A + YWHAZ-A | 17.5 | 97.5 | 154 | 52 | 18 | 13 | 4.6 | 0.35 |

**ID** = Rule IDs, rules are numbered according to importance rank determined from phase 1
**CR** = Coverage rank
**SJ/SJR** = Single rule performance / performance rank
**NSC/PC** = Number/percent of samples covered
**NSA/PA** = Number/percent of samples assigned
**FracA** = Fraction of covered samples assigned to rule

#### 3.2 Event breakdown of core rule set

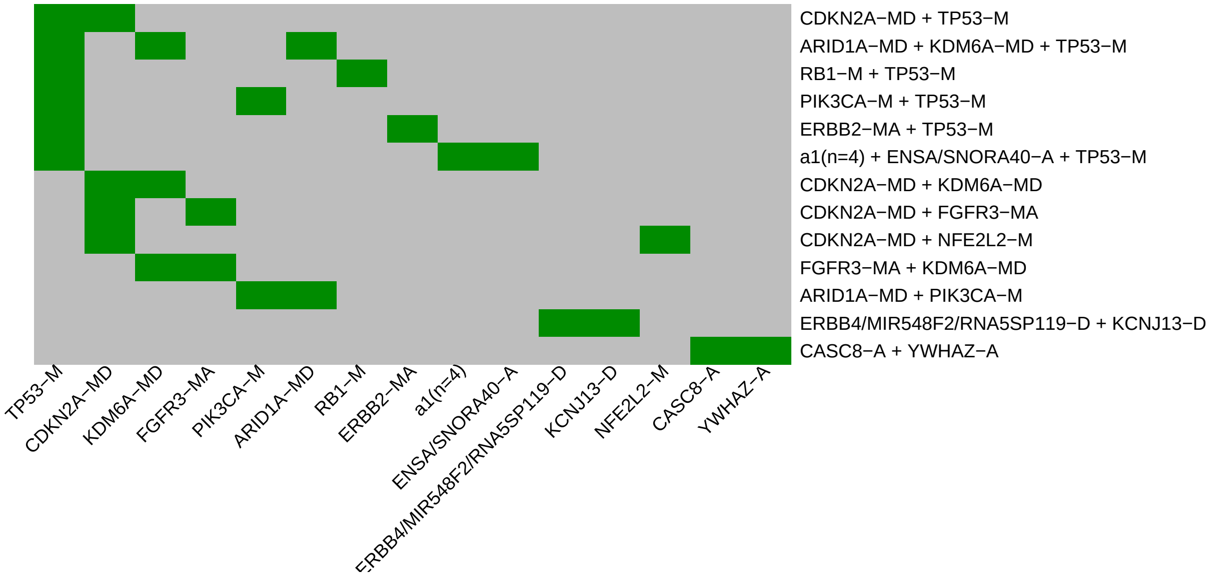

Figure 3: Visualization of core rule set. Rows are rules, columns are events. Events are ordered from left in decreasing rule membership frequency.

#### 3.3 Core Rule Set Penalties: Before and After Assignment

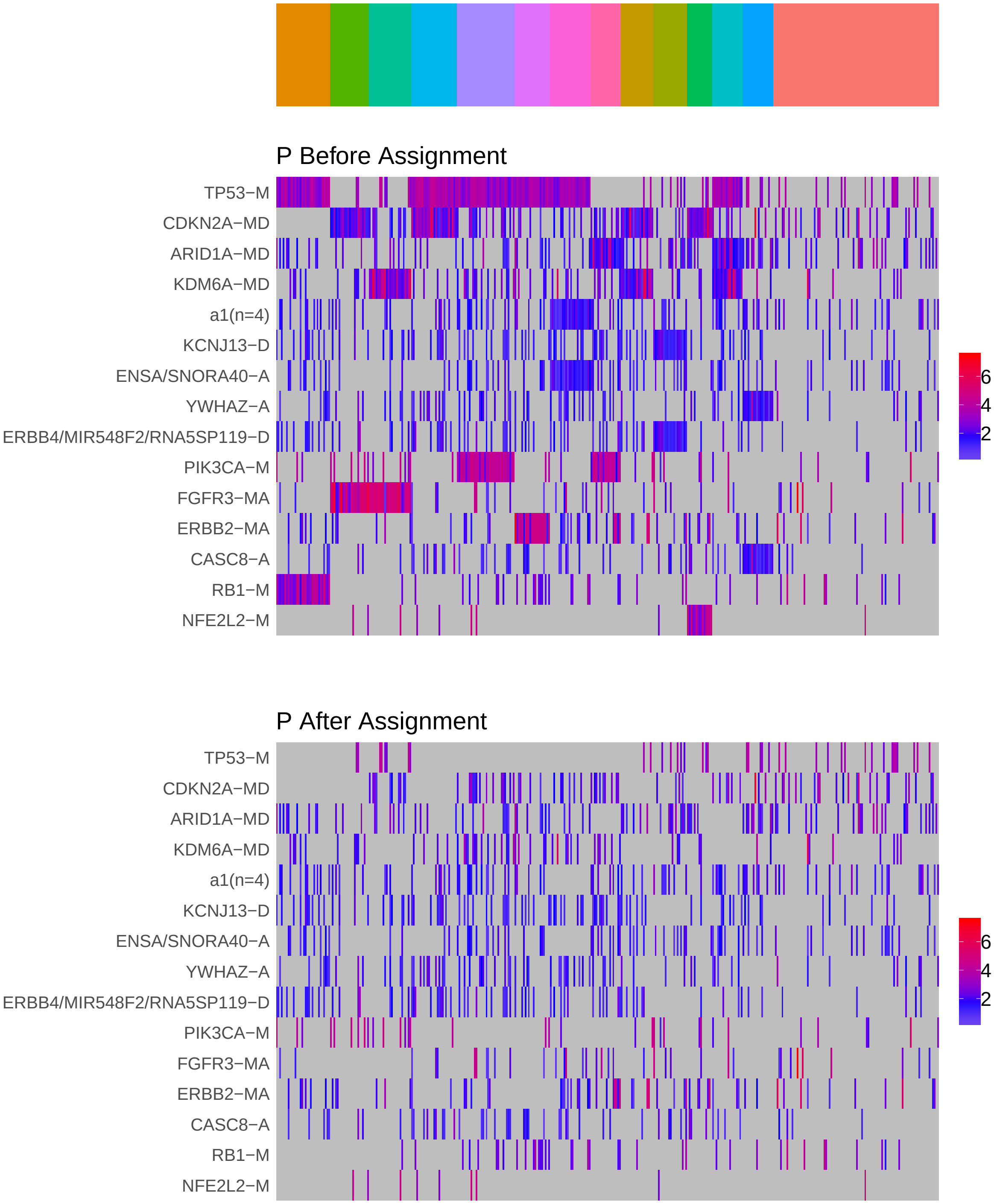

Figure 4: Heatmap of P before and after assignment to core rule set. Events are force ordered by frequency. Samples are force ordered according rule set membership, as indicated by the color bar. The right-most group of samples are not assigned to any rule.

### 4 Generalized Core Rules

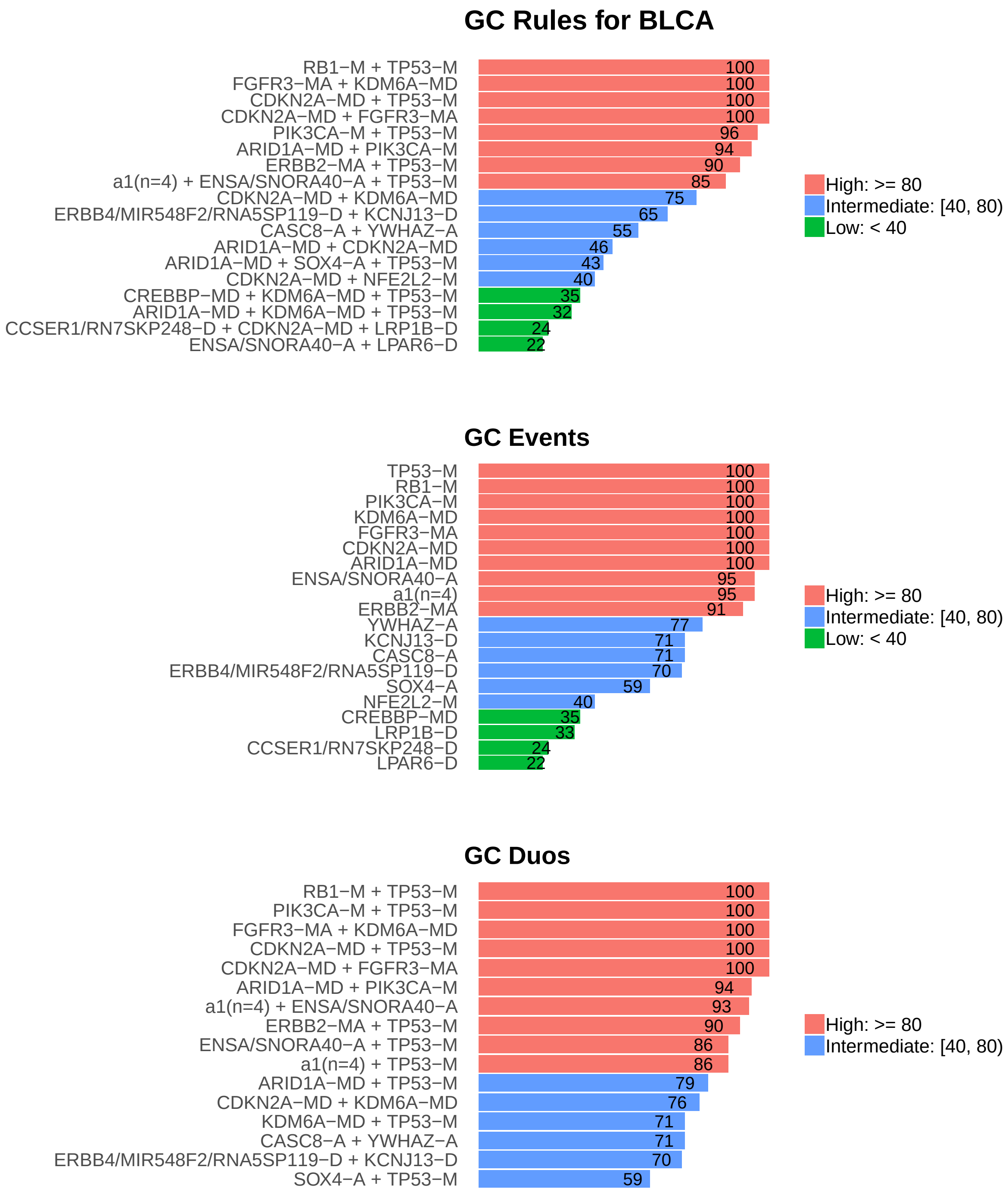

Figure 5: Summary of generalized core results. Bars show confidence levels, which are the percentage of sub-sample iterations containing the observation. Rules (top) and events (middle) that achieve a minimum confidence level of 20 are shown. Duos (bottom) with confidence of at least 50 are shown.

### 5 Dictionary of Copy Number Events

| CNV | Genes | Event__Name |
|---|---|---|
| a1 | PFDN2, PVRL4, KLHDC9, ARHGAP30 | a1(n=4) |
| a2 | SOX4 | SOX4-A |
| a3 | SNORA40\|ENSG00000253047.1, ENSA | ENSA/SNORA40-A |
| a4 | YWHAZ | YWHAZ-A |
| a5 | PPARG | PPARG-A |
| a6 | CCND1, ORAOV1 | CCND1/ORAOV1-A |
| a7 | CASC8 | CASC8-A |
| d2 | KCNJ13 | KCNJ13-D |
| d3 | MIR548F2, RNA5SP119, ERBB4 | ERBB4/MIR548F2/RNA5SP119-D |
| d4 | LPAR6 | LPAR6-D |
| d5 | FKSG52, MIR582, PDE4D | FKSG52/MIR582/PDE4D-D |
| d6 | RN7SKP248, CCSER1 | CCSER1/RN7SKP248-D |
| d7 | LRP1B | LRP1B-D |

Dictionary of important CNVs. Copy number events that are within any of the best K rule sets, or within the generalized core rules, or among the 20 most frequent events.
