## Supplementary material for "Identifying Combinations of Cancer Drivers in Individual Patients": TCGA Reports: CRSO_Report_BRCA.pdf

### CRSO Output Report: BRCA

Michael Klein

Apr 19 2019

**4 Generalized Core Rules** **6**

**5 Dictionary of Copy Number Events** **7**

#### 1 Heatmaps of D and P

**Number of samples = 963.**
**Number of events = 90.**
**Rule coverage requirement = 29 samples.**
**Rule library size = 449 rules.**

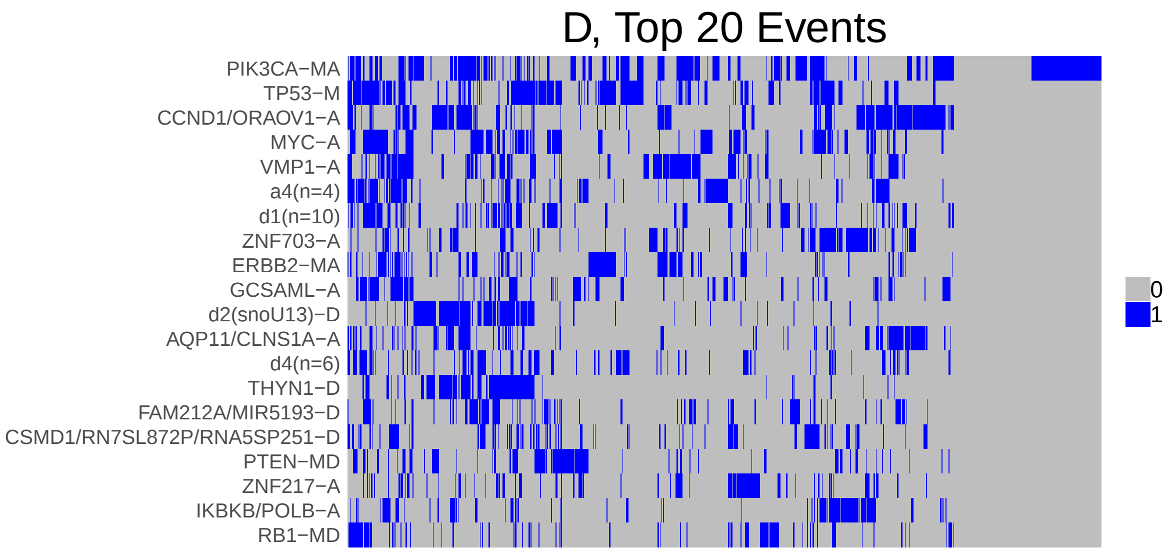

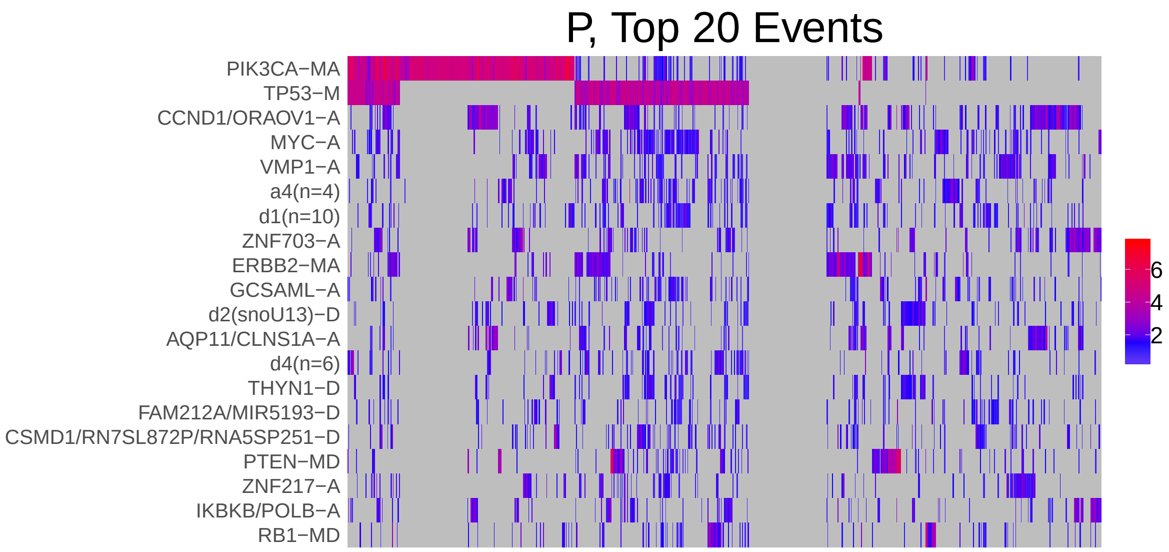

Figure 1: Heatmap of D and P. Events are ordered by frequency, from top to bottom. Event suffix -M: mutation, -A: amplification, -D: deletion, -MD: mutDel, -MA: mutAmp. Samples are ordered using hierarchical clustering. Wild-type events indicated in grey.

### 2 Summary of K Best Rule Sets

#### 2.1 Performance and coverage of best rule sets

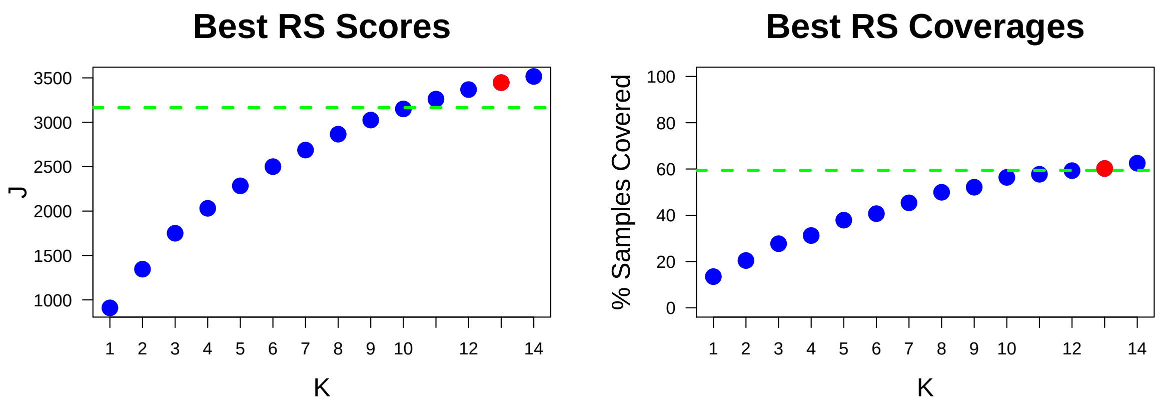

Figure 2: Performance and Coverage of Best Filtered RS for K. The core was determined to be the smallest RS that achieved at least 95% of maximum coverage and at least 90% of maximum performance.

#### 2.2 Table of rules that appear in any best rule set

| **ID** | **Rule** | **PC** | **Ks** |
|---|---|---|---|
| r1 | PIK3CA-MA + TP53-M | 13 | 1,2,3,4,5,6,7,8,9,10,11,12,13,14 |
| r4 | CCND1/ORAOV1-A + PIK3CA-MA | 11 | 2,3,4,6,7,8,9,10,11,12,13,14 |
| r5 | MYC-A + TP53-M | 13 | 3,4,5,6,7,8,9,10,11,12,13,14 |
| r2 | CDH1-M + PIK3CA-MA | 5.6 | 4,5,6,7,8,9,10,11,12,13,14 |
| r3 | MAP3K1-M + PIK3CA-MA | 3.7 | 5,6,7,8,9,10,11,12,13,14 |
| r6 | AQP11/CLNS1A-A + CCND1/ORAOV1-A | 13 | 5,6,7,8,9,10,11,12,13,14 |
| r10 | IKBKB/POLB-A + ZNF703-A | 8.9 | 7,8,9,10,11,12,13,14 |
| r12 | ERBB2-MA + VMP1-A | 9.3 | 8,9,10,11,12,13,14 |
| r9 | PTEN-MD + TP53-M | 7 | 9,10,11,12,13,14 |
| r11 | d2(snoU13)-D + THYN1-D | 12 | 10,11,12,13,14 |
| r8 | ERBB2-MA + TP53-M | 8.4 | 11,12,13,14 |
| r23 | GATA3-M + VMP1-A | 3.4 | 12,13,14 |
| r25 | RB1-MD + TP53-M | 6.4 | 13,14 |
| r37 | FBLN5-D + ZFP36L1-MD | 5.6 | 14 |

**ID** = Rule IDs, rules are numbered according to importance rank determined from phase 1
**PC** = Percent of samples covered **Ks** = Membership in best RS

### 3 Core Rule Set

**Core K = 13.**
**Core rule set coverage = 60.2%.**

#### 3.1 Table of core rule set rules

| ID | Rule | CR | SJR | SJ | NSC | NSA | PC | PA | FracA |
|---|---|---|---|---|---|---|---|---|---|
| r1 | PIK3CA-MA + TP53-M | 1 | 1 | 911 | 130 | 93 | 13 | 9.7 | 0.72 |
| r2 | CDH1-M + PIK3CA-MA | 70 | 10 | 424 | 54 | 45 | 5.6 | 4.7 | 0.83 |
| r3 | MAP3K1-M + PIK3CA-MA | 241.5 | 49.5 | 282 | 36 | 31 | 3.7 | 3.2 | 0.86 |
| r4 | CCND1/ORAOV1-A + PIK3CA-MA | 5 | 3 | 627 | 109 | 50 | 11 | 5.2 | 0.46 |
| r5 | MYC-A + TP53-M | 2 | 2 | 676 | 123 | 64 | 13 | 6.6 | 0.52 |
| r6 | AQP11/CLNS1A-A + CCND1/ORAOV1-A | 3 | 7 | 448 | 121 | 48 | 13 | 5.0 | 0.40 |
| r8 | ERBB2-MA + TP53-M | 13.5 | 4 | 478 | 81 | 32 | 8.4 | 3.3 | 0.40 |
| r9 | PTEN-MD + TP53-M | 31.5 | 17 | 386 | 67 | 38 | 7 | 3.9 | 0.57 |
| r10 | IKBKB/POLB-A + ZNF703-A | 10 | 36 | 320 | 86 | 39 | 8.9 | 4.0 | 0.45 |
| r11 | d2(snoU13)-D + THYN1-D | 4 | 32 | 327 | 118 | 41 | 12 | 4.3 | 0.35 |
| r12 | ERBB2-MA + VMP1-A | 7 | 28 | 330 | 90 | 37 | 9.3 | 3.8 | 0.41 |
| r23 | GATA3-M + VMP1-A | 308.5 | 175 | 168 | 33 | 31 | 3.4 | 3.2 | 0.94 |
| r25 | RB1-MD + TP53-M | 41 | 22 | 345 | 62 | 31 | 6.4 | 3.2 | 0.50 |

**ID** = Rule IDs, rules are numbered according to importance rank determined from phase 1
**CR** = Coverage rank
**SJ/SJR** = Single rule performance / performance rank
**NSC/PC** = Number/percent of samples covered
**NSA/PA** = Number/percent of samples assigned
**FracA** = Fraction of covered samples assigned to rule

#### 3.2 Event breakdown of core rule set

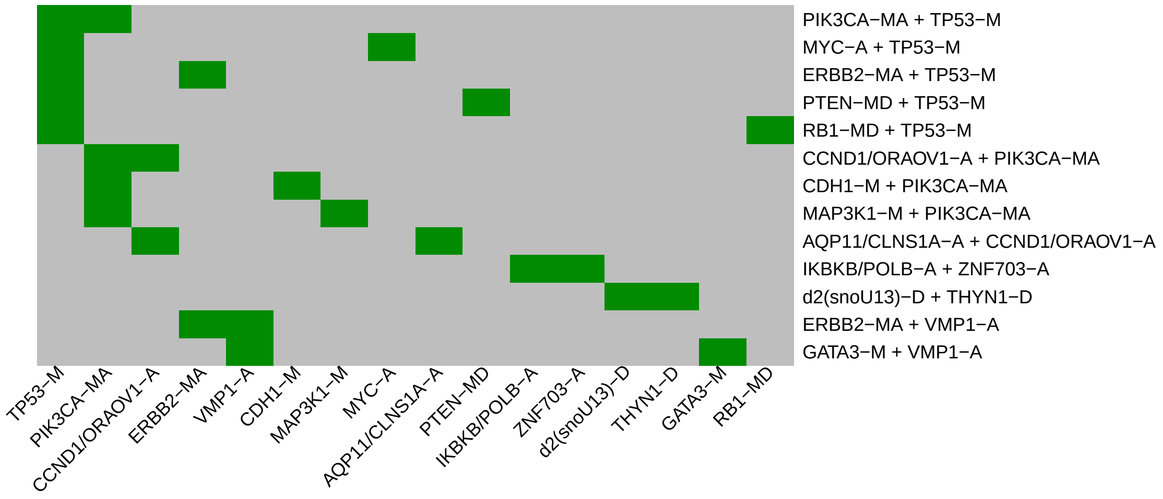

Figure 3: Visualization of core rule set. Rows are rules, columns are events. Events are ordered from left in decreasing rule membership frequency.

#### 3.3 Core Rule Set Penalties: Before and After Assignment

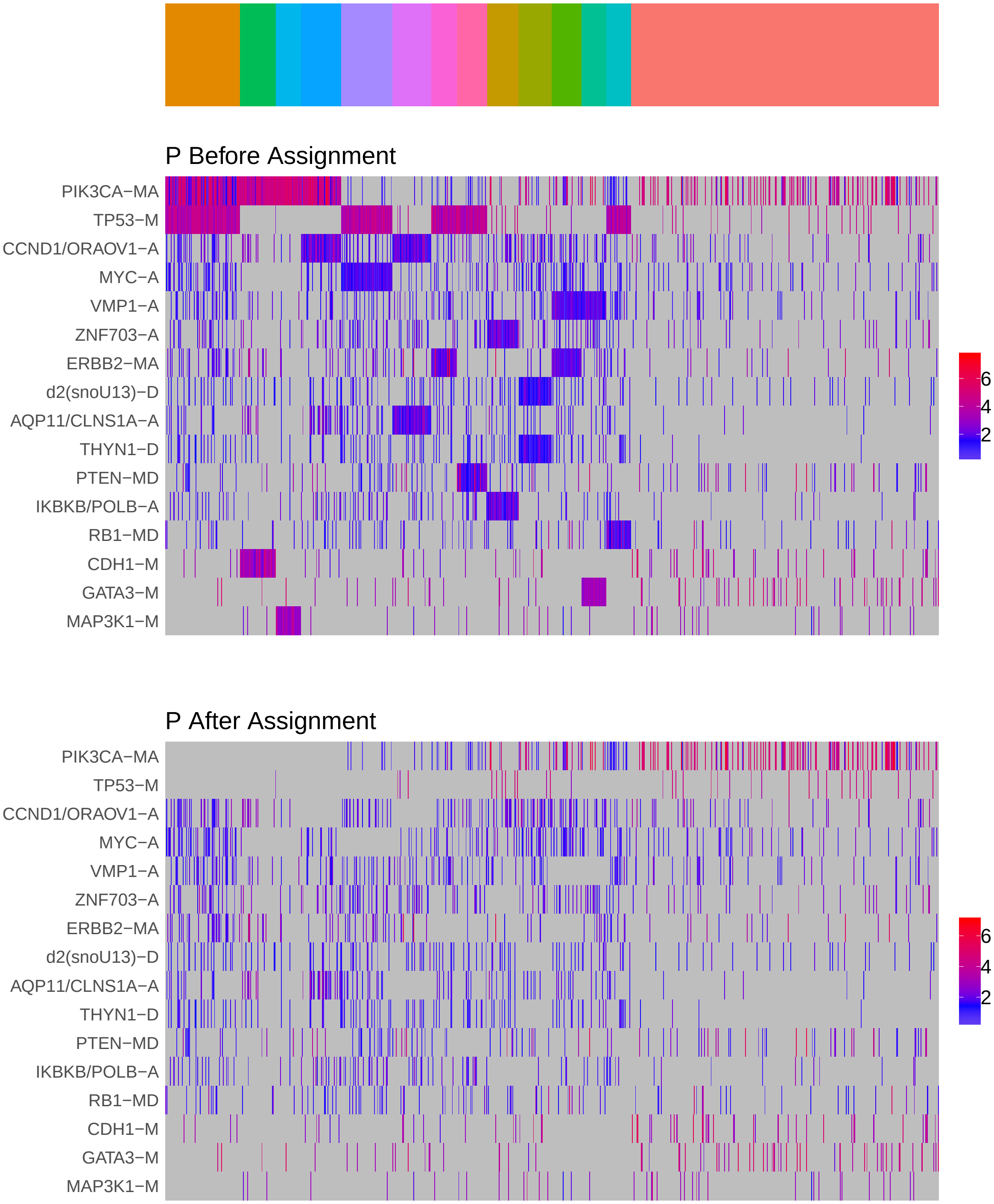

Figure 4: Heatmap of P before and after assignment to core rule set. Events are force ordered by frequency. Samples are force ordered according rule set membership, as indicated by the color bar. The right-most group of samples are not assigned to any rule.

### 4 Generalized Core Rules

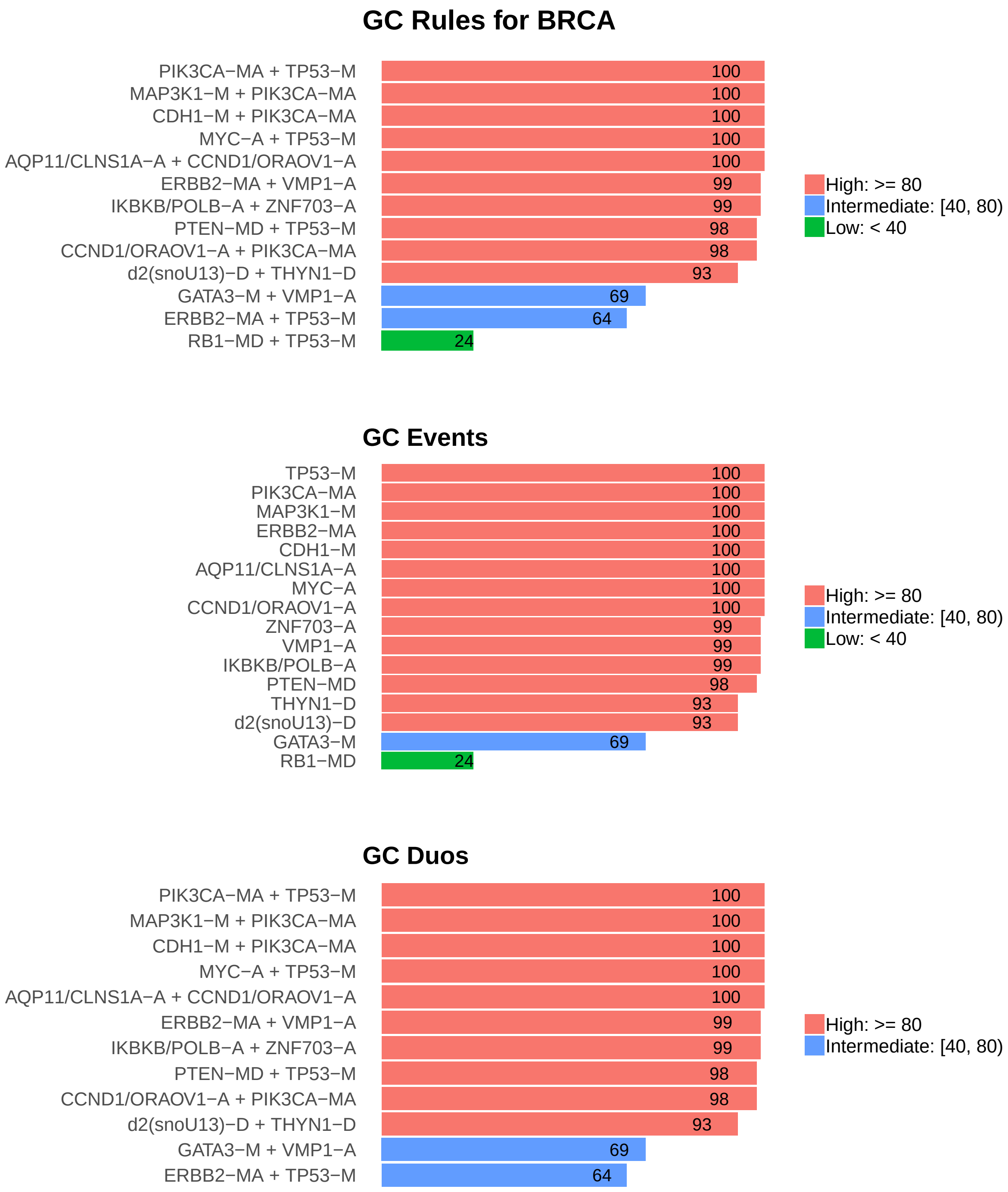

Figure 5: Summary of generalized core results. Bars show confidence levels, which are the percentage of sub-sample iterations containing the observation. Rules (top) and events (middle) that achieve a minimum confidence level of 20 are shown. Duos (bottom) with confidence of at least 50 are shown.

### 5 Dictionary of Copy Number Events

| CNV | Genes | Event_Name |
|---|---|---|
| a1 | CCND1, ORAOV1 | CCND1/ORAOV1-A |
| a2 | MYC | MYC-A |
| a3 | VMP1 | VMP1-A |
| a4 | SNORA40\|ENSG00000253047.1, ENSA, MCL1, GOLPH3L | a4(n=4) |
| a5 | ZNF703 | ZNF703-A |
| a6 | GCSAML | GCSAML-A |
| a7 | CLNS1A, AQP11 | AQP11/CLNS1A-A |
| a10 | ZNF217 | ZNF217-A |
| a11 | IKBKB, POLB | IKBKB/POLB-A |
| d1 | U1\|ENSG00000228549.2, MST1L, ESPNP, CROCCP2, U1\|ENSG00000233421.3, MFAP2, CROCC, ATP13A2, NBPF1, MIR3675 | d1(n=10) |
| d2 | snoU13\|ENSG00000239153.1 | d2(snoU13)-D |
| d3 | THYN1 | THYN1-D |
| d4 | UQCR11, RN7SL477P, TCF3, UQCR11, MBD3, MEX3D | d4(n=6) |
| d5 | MIR5193, FAM212A | FAM212A/MIR5193-D |
| d6 | RN7SL872P, RNA5SP251, CSMD1 | CSMD1/RN7SL872P/RNA5SP251-D |
| d27 | FBLN5 | FBLN5-D |

Dictionary of important CNVs. Copy number events that are within any of the best K rule sets, or within the generalized core rules, or among the 20 most frequent events.
