## Supplementary material for "Identifying Combinations of Cancer Drivers in Individual Patients": TCGA Reports: CRSO_Report_CESC.pdf

### CRSO Output Report: CESC

Michael Klein

Apr 19 2019

**4 Generalized Core Rules** **6**

**5 Dictionary of Copy Number Events** **7**

### 1 Heatmaps of D and P

**Number of samples = 191.**
**Number of events = 82.**
**Rule coverage requirement = 6 samples.**
**Rule library size = 173 rules.**

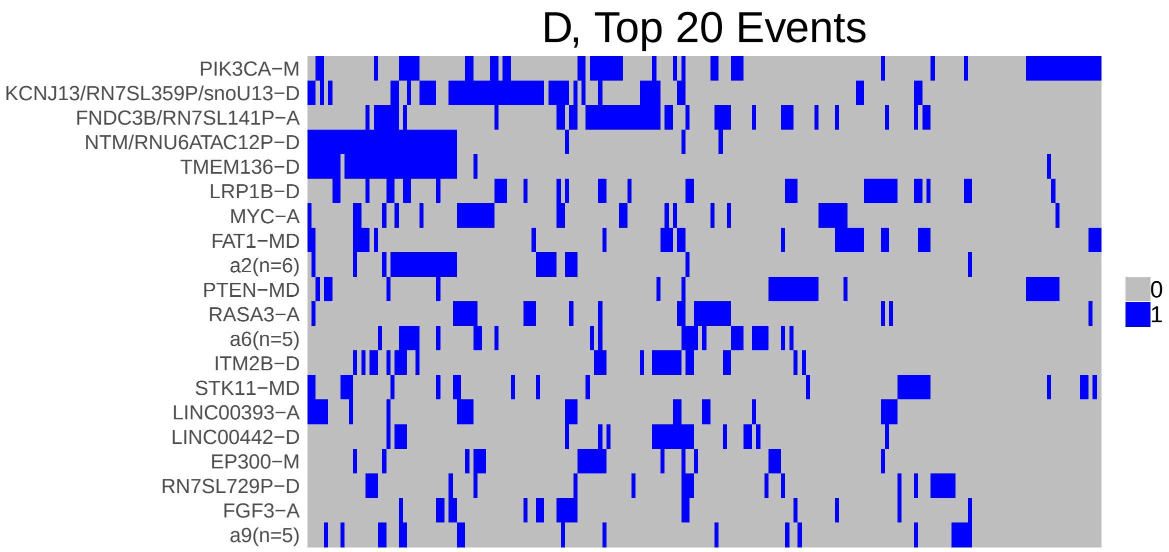

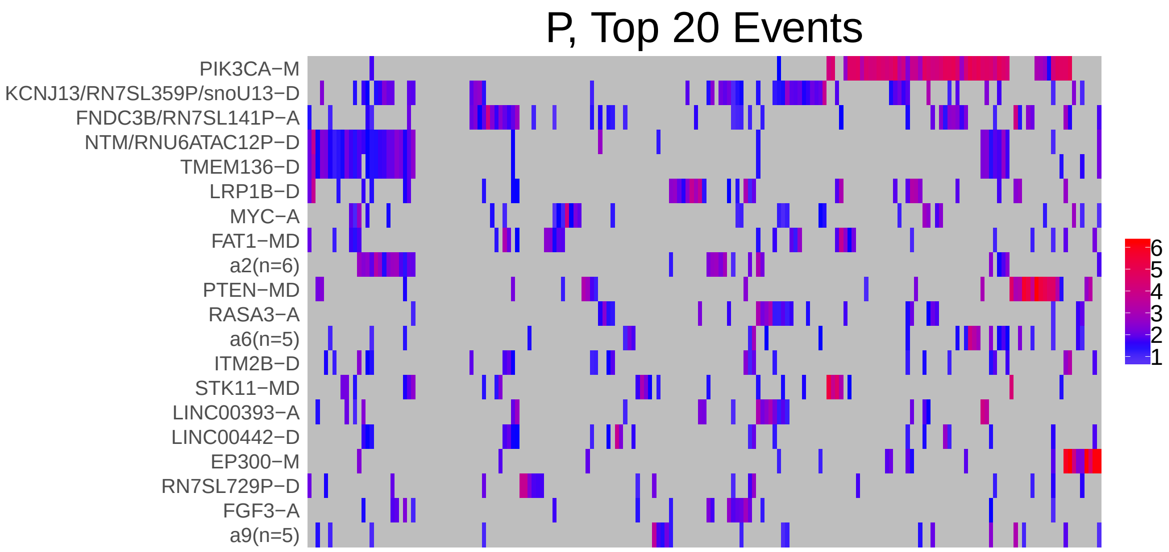

Figure 1: Heatmap of D and P. Events are ordered by frequency, from top to bottom. Event suffix -M: mutation, -A: amplification, -D: deletion, -MD: mutDel, -MA: mutAmp. Samples are ordered using hierarchical clustering. Wild-type events indicated in grey.

### 2 Summary of K Best Rule Sets

#### 2.1 Performance and coverage of best rule sets

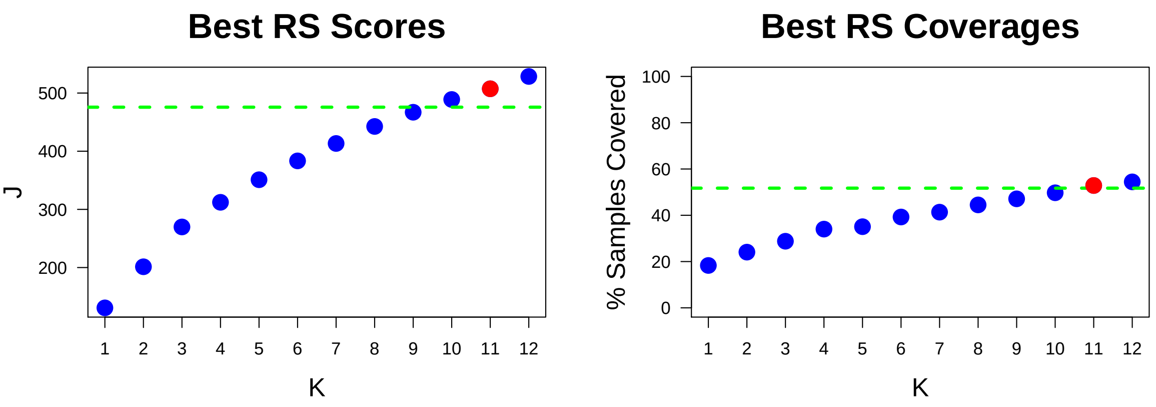

Figure 2: Performance and Coverage of Best Filtered RS for K. The core was determined to be the smallest RS that achieved at least 95% of maximum coverage and at least 90% of maximum performance.

#### 2.2 Table of rules that appear in any best rule set

| ID | Rule | PC | Ks |
|---|---|---|---|
| r1 | NTM/RNU6ATAC12P-D + TMEM136-D | 18 | 1,2,3,4,5,6,7,8,9,10,11,12 |
| r2 | PIK3CA-M + PTEN-MD | 5.2 | 3,4,5,6,7,8,9,10,11,12 |
| r3 | FNDC3B/RN7SL141P-A + PIK3CA-M | 6.8 | 2,3,4,5,6,7,8,9,10 |
| r10 | LINC00393-A + RASA3-A | 5.8 | 4,5,6,7,8,9,10,11,12 |
| r9 | FNDC3B/RN7SL141P-A + LRP1B-D | 7.9 | 6,7,8,9,10,11,12 |
| r11 | a2(n=6) + KCNJ13/RN7SL359P/snoU13-D | 8.4 | 8,9,10,11,12 |
| r14 | FBXW7-M + PIK3CA-M | 3.1 | 5,6,7,8,9 |
| r38 | a11(n=6) + ZNF750-MD | 3.7 | 9,10,11,12 |
| r6 | a6(n=5) + PIK3CA-M | 6.3 | 10,11,12 |
| r17 | KCNJ13/RN7SL359P/snoU13-D + PIK3CA-M | 6.3 | 10,11,12 |
| r20 | PIK3CA-M + ZNF750-MD | 3.7 | 7,8,9 |
| r5 | BCL2L1/COX4I2-A + PIK3CA-M | 3.7 | 11,12 |
| r19 | a6(n=5) + FBXW7-M | 3.1 | 11,12 |
| r30 | FNDC3B/RN7SL141P-A + ITM2B-D | 7.9 | 11,12 |
| r25 | FAT1-MD + PIK3CA-M | 3.7 | 12 |
| r41 | FNDC3B/RN7SL141P-A + ITM2B-D + LINC00442-D | 5.2 | 10 |

**ID** = Rule IDs, rules are numbered according to importance rank determined from phase 1
**PC** = Percent of samples covered **Ks** = Membership in best RS

### 3 Core Rule Set

**Core K = 11.**
**Core rule set coverage = 52.9%.**

#### 3.1 Table of core rule set rules

| ID | Rule | CR | SJR | SJ | NSC | NSA | PC | PA | FracA |
|---|---|---|---|---|---|---|---|---|---|
| r1 | NTM/RNU6ATAC12P-D + TMEM136-D | 1 | 1 | 131 | 35 | 19 | 18 | 9.9 | 0.54 |
| r2 | PIK3CA-M + PTEN-MD | 28 | 8 | 73 | 10 | 9 | 5.2 | 4.7 | 0.90 |
| r5 | BCL2L1/COX4I2-A + PIK3CA-M | 84.5 | 27 | 47.1 | 7 | 7 | 3.7 | 3.7 | 1.00 |
| r6 | a6(n=5) + PIK3CA-M | 17.5 | 7 | 73.5 | 12 | 6 | 6.3 | 3.1 | 0.50 |
| r9 | FNDC3B/RN7SL141P-A + LRP1B-D | 7.5 | 16 | 58.1 | 15 | 11 | 7.9 | 5.8 | 0.73 |
| r10 | LINC00393-A + RASA3-A | 21.5 | 30 | 45.2 | 11 | 9 | 5.8 | 4.7 | 0.82 |
| r11 | a2(n=6) + KCNJ13/RN7SL359P/snoU13-D | 5 | 11 | 65.3 | 16 | 12 | 8.4 | 6.3 | 0.75 |
| r17 | KCNJ13/RN7SL359P/snoU13-D + PIK3CA-M | 17.5 | 10 | 69.4 | 12 | 9 | 6.3 | 4.7 | 0.75 |
| r19 | a6(n=5) + FBXW7-M | 138.5 | 48 | 38.7 | 6 | 6 | 3.1 | 3.1 | 1.00 |
| r30 | FNDC3B/RN7SL141P-A + ITM2B-D | 7.5 | 23 | 50.9 | 15 | 7 | 7.9 | 3.7 | 0.47 |
| r38 | a11(n=6) + ZNF750-MD | 84.5 | 77.5 | 30.4 | 7 | 6 | 3.7 | 3.1 | 0.86 |

**ID** = Rule IDs, rules are numbered according to importance rank determined from phase 1
**CR** = Coverage rank
**SJ/SJR** = Single rule performance / performance rank
**NSC/PC** = Number/percent of samples covered
**NSA/PA** = Number/percent of samples assigned
**FracA** = Fraction of covered samples assigned to rule

#### 3.2 Event breakdown of core rule set

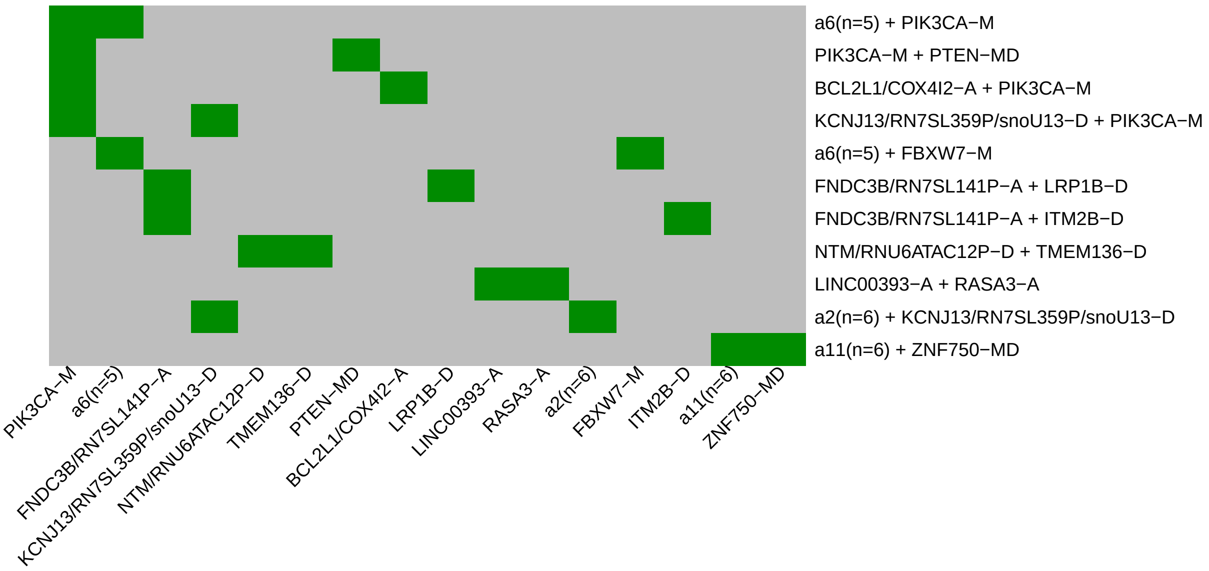

Figure 3: Visualization of core rule set. Rows are rules, columns are events. Events are ordered from left in decreasing rule membership frequency.

#### 3.3 Core Rule Set Penalties: Before and After Assignment

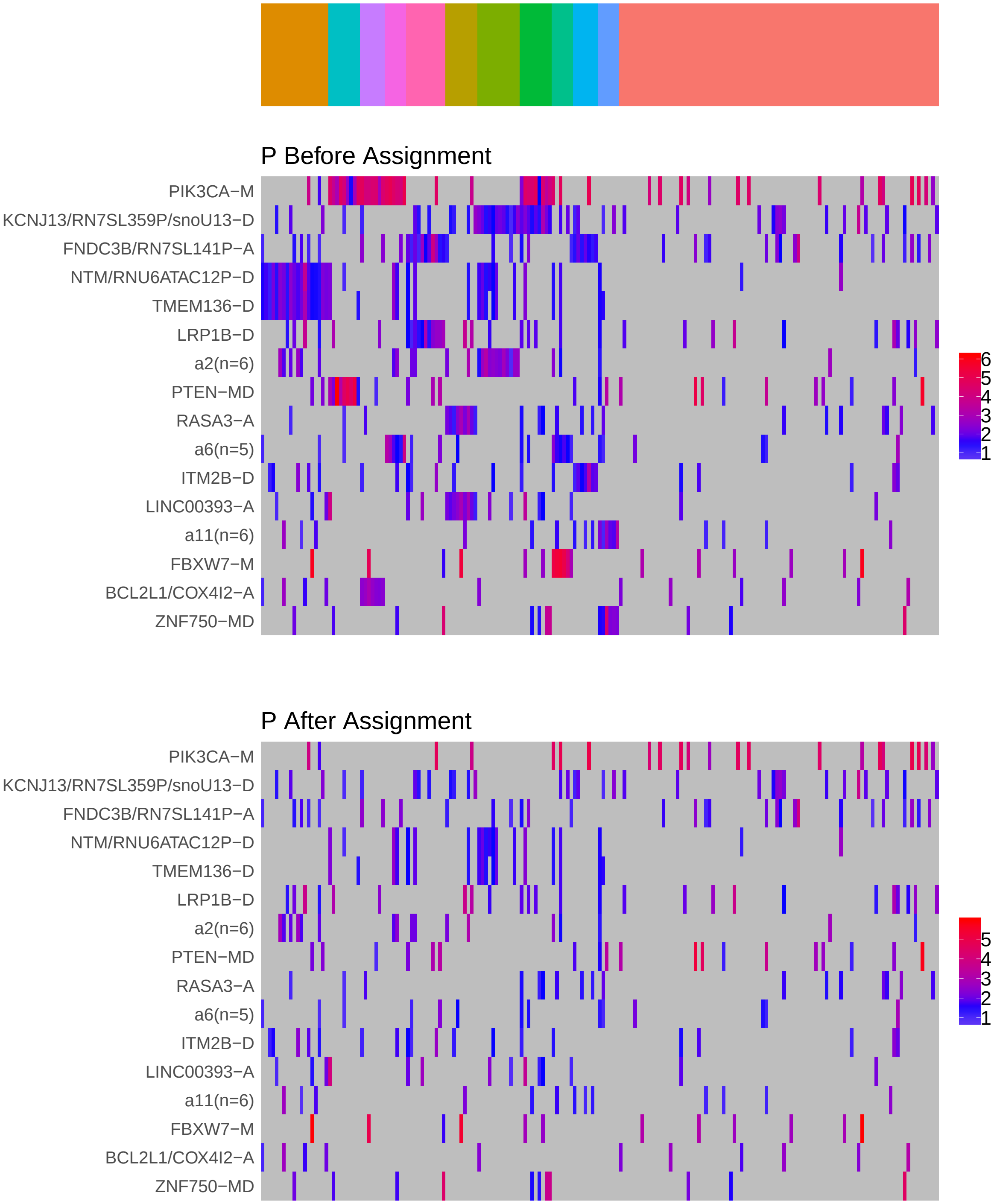

Figure 4: Heatmap of P before and after assignment to core rule set. Events are force ordered by frequency. Samples are force ordered according rule set membership, as indicated by the color bar. The right-most group of samples are not assigned to any rule.

### 4 Generalized Core Rules

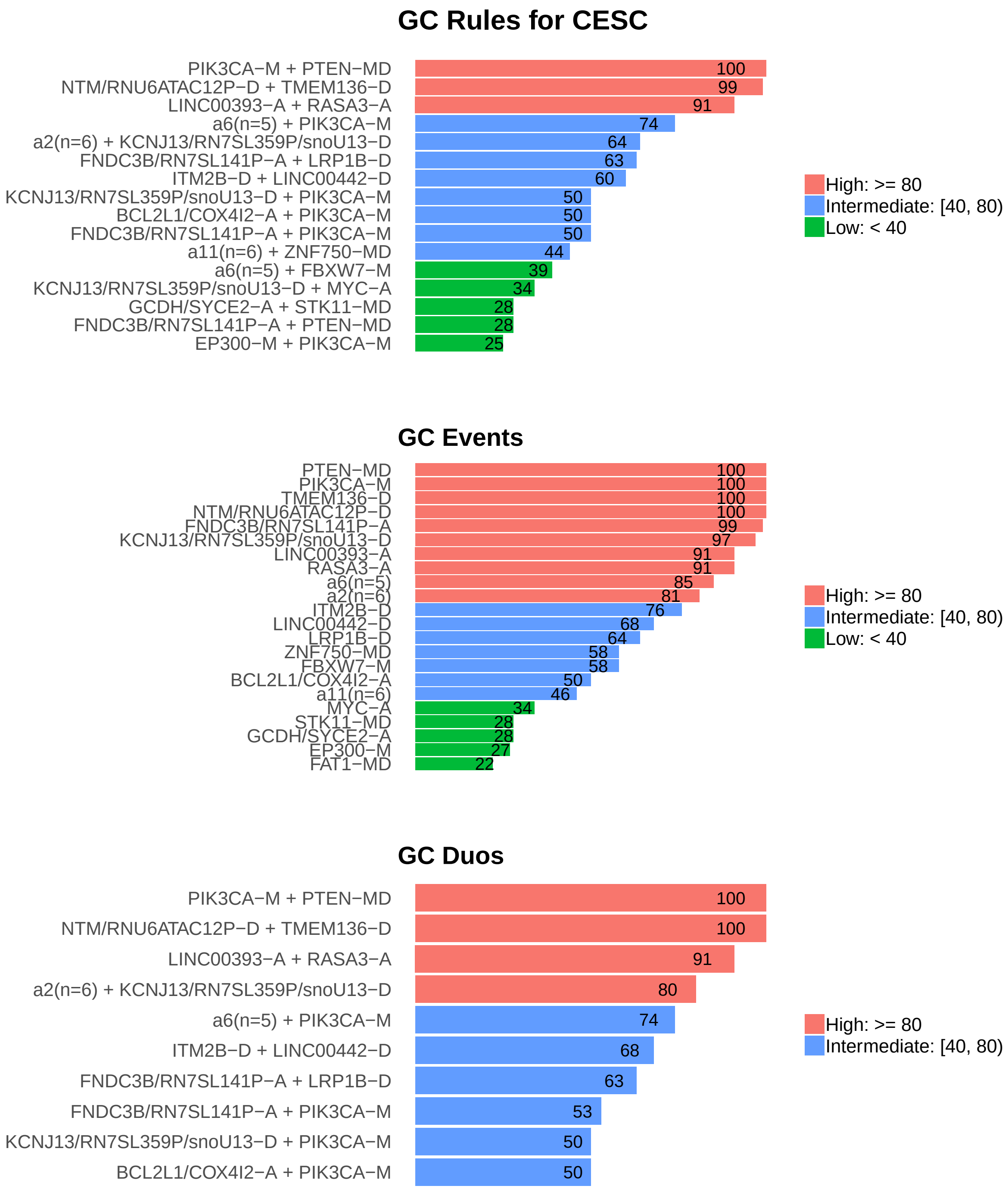

Figure 5: Summary of generalized core results. Bars show confidence levels, which are the percentage of sub-sample iterations containing the observation. Rules (top) and events (middle) that achieve a minimum confidence level of 20 are shown. Duos (bottom) with confidence of at least 50 are shown.

### 5 Dictionary of Copy Number Events

| CNV | Genes | Event__Name |
|---|---|---|
| a1 | RN7SL141P, FNDC3B | FNDC3B/RN7SL141P-A |
| a2 | snoU13\|ENSG00000239154.1, snoU13\|ENSG00000252679.1, BIRC2, BIRC3, YAP1, C11orf70 | a2(n=6) |
| a3 | RASA3 | RASA3-A |
| a4 | LINC00393 | LINC00393-A |
| a5 | MYC | MYC-A |
| a6 | NAA10, ARHGAP4, HCFC1, RENBP, TMEM187 | a6(n=5) |
| a7 | FGF3 | FGF3-A |
| a9 | PI4KB, PSMB4, RFX5, SELENBP1, POGZ | a9(n=5) |
| a11 | ACOX1, WBP2, MRPL38, FBF1, TRIM47, TRIM65 | a11(n=6) |
| a14 | BCL2L1, COX4I2 | BCL2L1/COX4I2-A |
| a17 | SPATS2L, KCTD18 | KCTD18/SPATS2L-A |
| a21 | SYCE2, GCDH | GCDH/SYCE2-A |
| d1 | snoU13\|ENSG00000239170.1, RN7SL359P, KCNJ13 | KCNJ13/RN7SL359P/snoU13-D |
| d2 | LRP1B | LRP1B-D |
| d3 | RNU6ATAC12P, NTM | NTM/RNU6ATAC12P-D |
| d4 | TMEM136 | TMEM136-D |
| d5 | ITM2B | ITM2B-D |
| d7 | LINC00442 | LINC00442-D |
| d11 | FKSG52, MIR582, PDE4D | FKSG52/MIR582/PDE4D-D |
| d12 | RN7SL729P | RN7SL729P-D |

Dictionary of important CNVs. Copy number events that are within any of the best K rule sets, or within the generalized core rules, or among the 20 most frequent events.
