## Supplementary material for "Identifying Combinations of Cancer Drivers in Individual Patients": TCGA Reports: CRSO_Report_COAD.pdf

### CRSO Output Report: COAD

Michael Klein

Apr 19 2019

**4 Generalized Core Rules** **6**

**5 Dictionary of Copy Number Events** **7**

#### 1 Heatmaps of D and P

**Number of samples = 362.**
**Number of events = 97.**
**Rule coverage requirement = 11 samples.**
**Rule library size = 986 rules.**

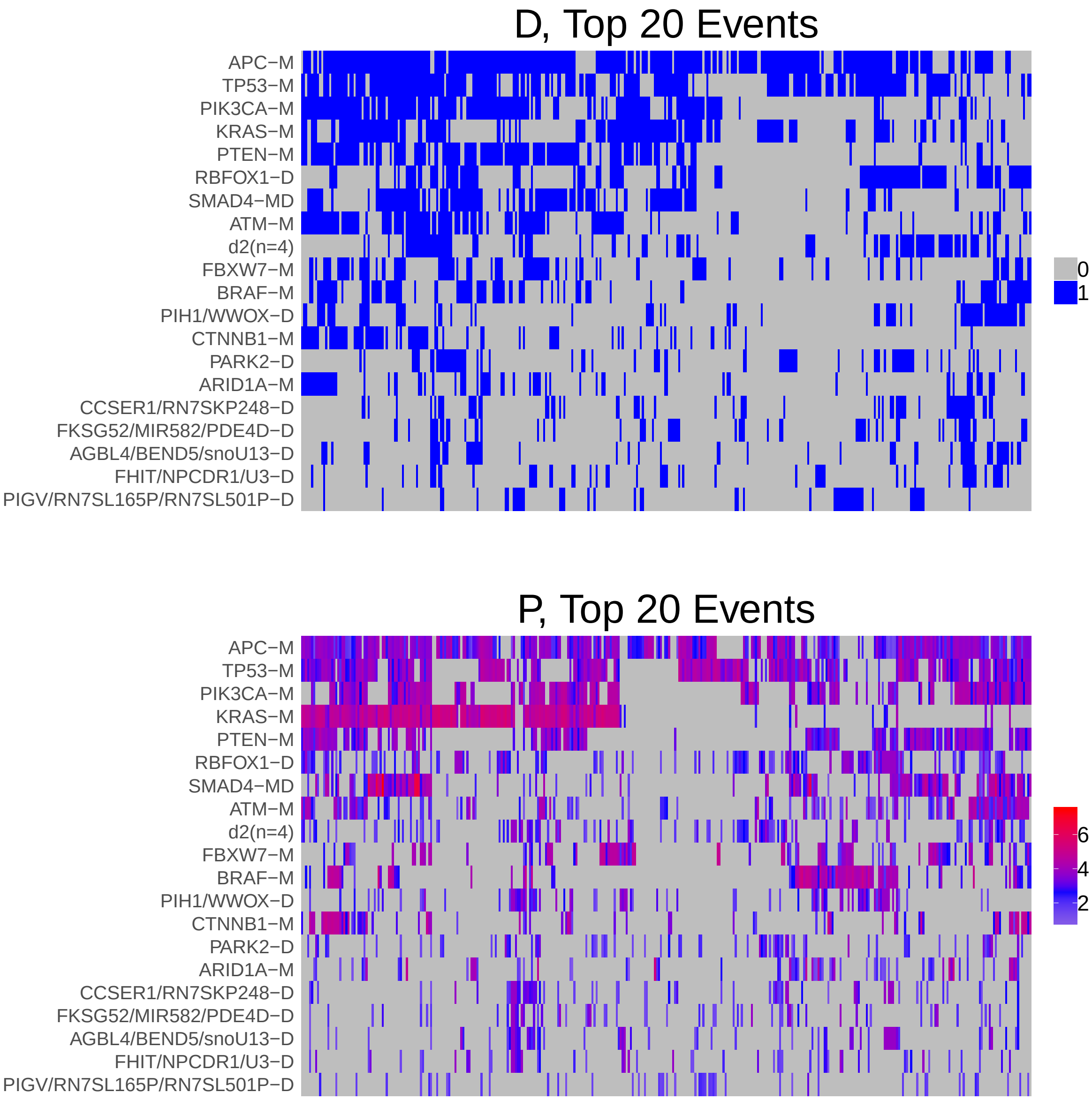

Figure 1: Heatmap of D and P. Events are ordered by frequency, from top to bottom. Event suffix -M: mutation, -A: amplification, -D: deletion, -MD: mutDel, -MA: mutAmp. Samples are ordered using hierarchical clustering. Wild-type events indicated in grey.

### 2 Summary of K Best Rule Sets

#### 2.1 Performance and coverage of best rule sets

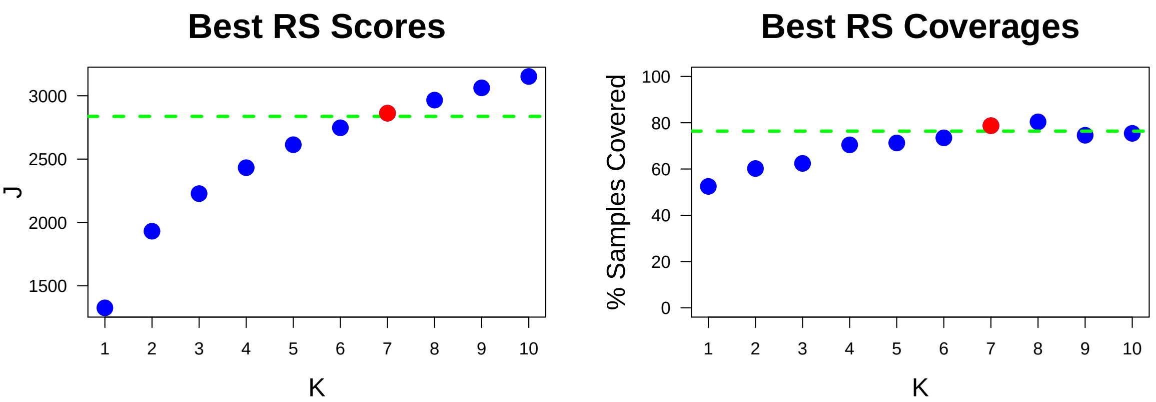

Figure 2: Performance and Coverage of Best Filtered RS for K. The core was determined to be the smallest RS that achieved at least 95% of maximum coverage and at least 90% of maximum performance.

#### 2.2 Table of rules that appear in any best rule set

| **ID** | **Rule** | **PC** | **Ks** |
|---|---|---|---|
| r1 | APC-M + TP53-M | 52 | 1,2,3,4,5,6,7,8 |
| r2 | APC-M + KRAS-M | 40 | 4,5,6,7,8 |
| r26 | APC-M + PIK3CA-M + PTEN-M | 28 | 4,5,6,7,8 |
| r101 | KRAS-M + PIK3CA-M + PTEN-M + TP53-M | 12 | 4,5,6,7,8 |
| r52 | ATM-M + PIK3CA-M + SMAD4-MD + TP53-M | 11 | 5,6,7,8 |
| r9 | APC-M + ATM-M + PIK3CA-M + PTEN-M | 17 | 3,9,10 |
| r70 | KRAS-M + RBFOX1-D + TP53-M | 11 | 6,7,8 |
| r3 | APC-M + KRAS-M + TP53-M | 26 | 9,10 |
| r4 | APC-M + ATM-M + PIK3CA-M + SMAD4-MD + TP53-M | 10 | 9,10 |
| r5 | RBFOX1-D + TP53-M | 25 | 9,10 |
| r6 | APC-M + PIK3CA-M + PTEN-M + TP53-M | 20 | 9,10 |
| r7 | BRAF-M + PIH1/WWOX-D + RBFOX1-D | 5.5 | 9,10 |
| r11 | KRAS-M + PIK3CA-M | 27 | 9,10 |
| r12 | APC-M + PTEN-M + SMAD4-MD | 20 | 9,10 |
| r15 | APC-M + KRAS-M + PIK3CA-M | 24 | 2,3 |
| r23 | APC-M + d2(n=4) | 23 | 9,10 |
| r111 | PIH1/WWOX-D + RBFOX1-D | 10 | 7,8 |
| r128 | BRAF-M + PTEN-M + SMAD4-MD | 9.7 | 8 |
| r237 | BRAF-M + PIK3CA-M + PTEN-M + SMAD4-MD + TP53-M | 5.2 | 10 |

**ID** = Rule IDs, rules are numbered according to importance rank determined from phase 1
**PC** = Percent of samples covered **Ks** = Membership in best RS

### 3 Core Rule Set

**Core K = 7.**
**Core rule set coverage = 78.7%.**

#### 3.1 Table of core rule set rules

| ID | Rule | CR | SJR | SJ | NSC | NSA | PC | PA | FracA |
|---|---|---|---|---|---|---|---|---|---|
| r1 | APC-M + TP53-M | 1 | 1 | 1330 | 190 | 58 | 52 | 16.0 | 0.31 |
| r2 | APC-M + KRAS-M | 3 | 2 | 1130 | 143 | 60 | 40 | 17.0 | 0.42 |
| r26 | APC-M + PIK3CA-M + PTEN-M | 13 | 6 | 1030 | 101 | 46 | 28 | 13.0 | 0.46 |
| r52 | ATM-M + PIK3CA-M + SMAD4-MD + TP53-M | 153 | 68 | 553 | 41 | 43 | 11 | 12.0 | 1.00 |
| r70 | KRAS-M + RBFOX1-D + TP53-M | 172 | 188 | 410 | 39 | 29 | 11 | 8.0 | 0.74 |
| r101 | KRAS-M + PIK3CA-M + PTEN-M + TP53-M | 133.5 | 45 | 642 | 43 | 27 | 12 | 7.5 | 0.63 |
| r111 | PIH1/WWOX-D + RBFOX1-D | 198 | 666 | 199 | 37 | 22 | 10 | 6.1 | 0.59 |

**ID** = Rule IDs, rules are numbered according to importance rank determined from phase 1
**CR** = Coverage rank
**SJ/SJR** = Single rule performance / performance rank
**NSC/PC** = Number/percent of samples covered
**NSA/PA** = Number/percent of samples assigned
**FracA** = Fraction of covered samples assigned to rule

#### 3.2 Event breakdown of core rule set

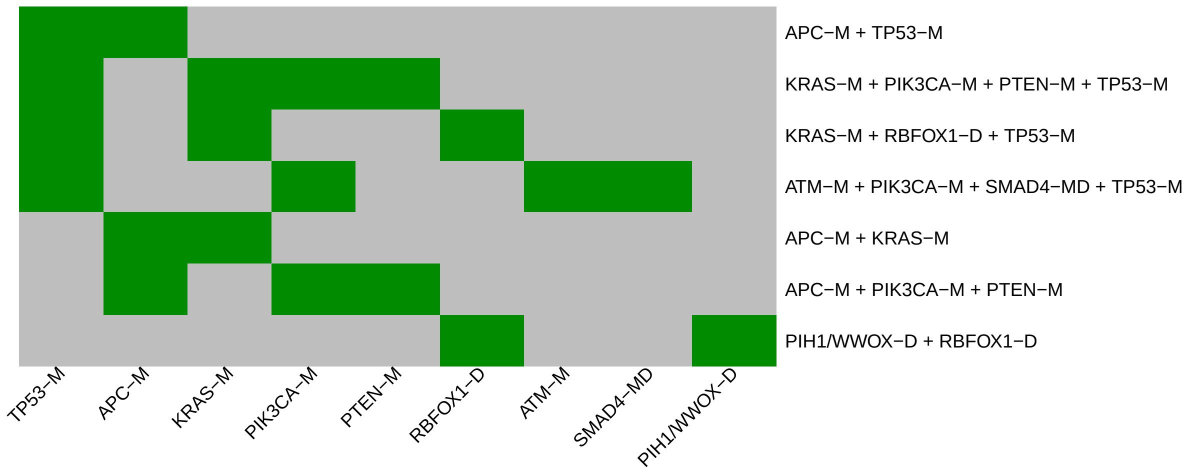

Figure 3: Visualization of core rule set. Rows are rules, columns are events. Events are ordered from left in decreasing rule membership frequency.

#### 3.3 Core Rule Set Penalties: Before and After Assignment

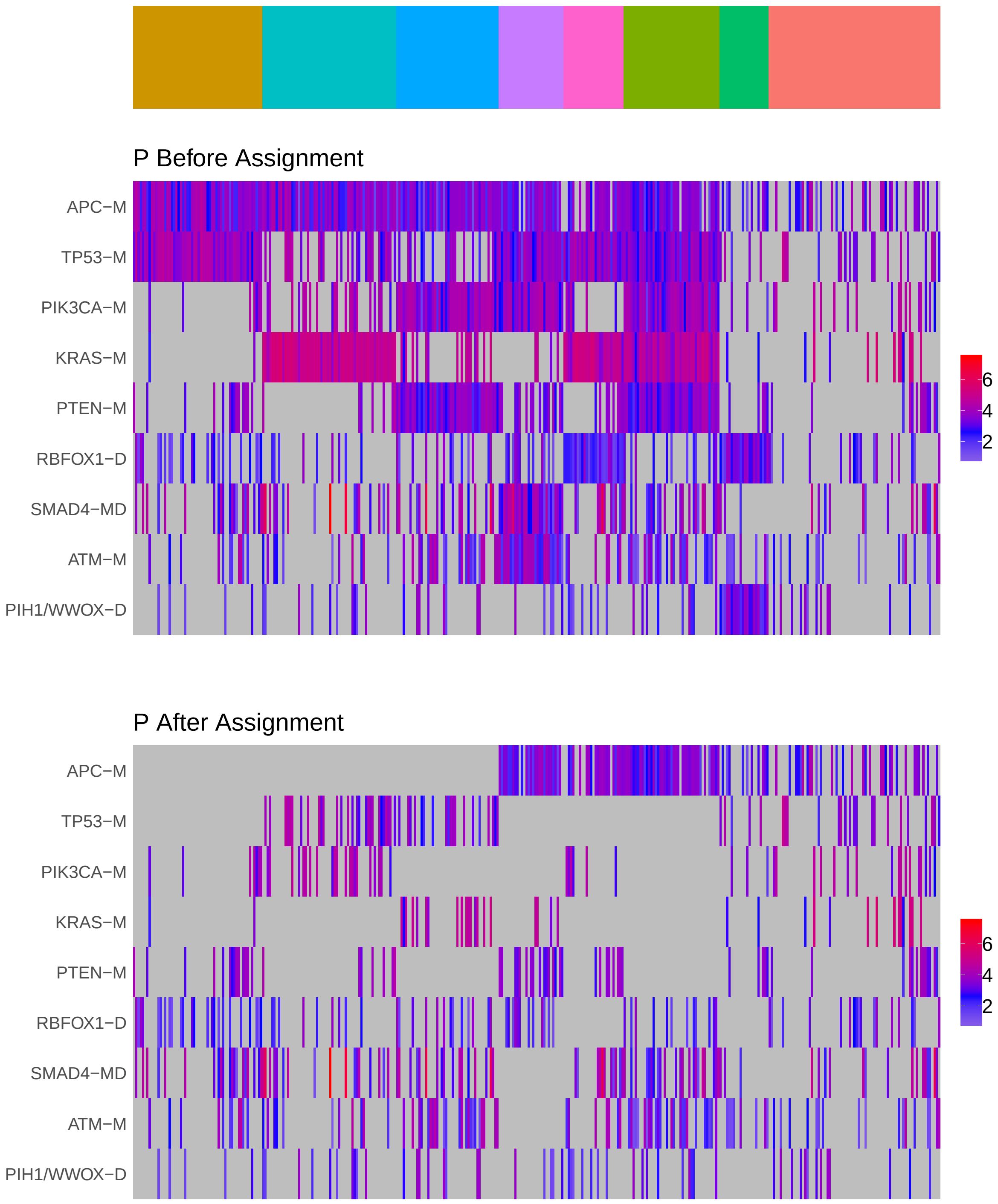

Figure 4: Heatmap of P before and after assignment to core rule set. Events are force ordered by frequency. Samples are force ordered according rule set membership, as indicated by the color bar. The right-most group of samples are not assigned to any rule.

### 4 Generalized Core Rules

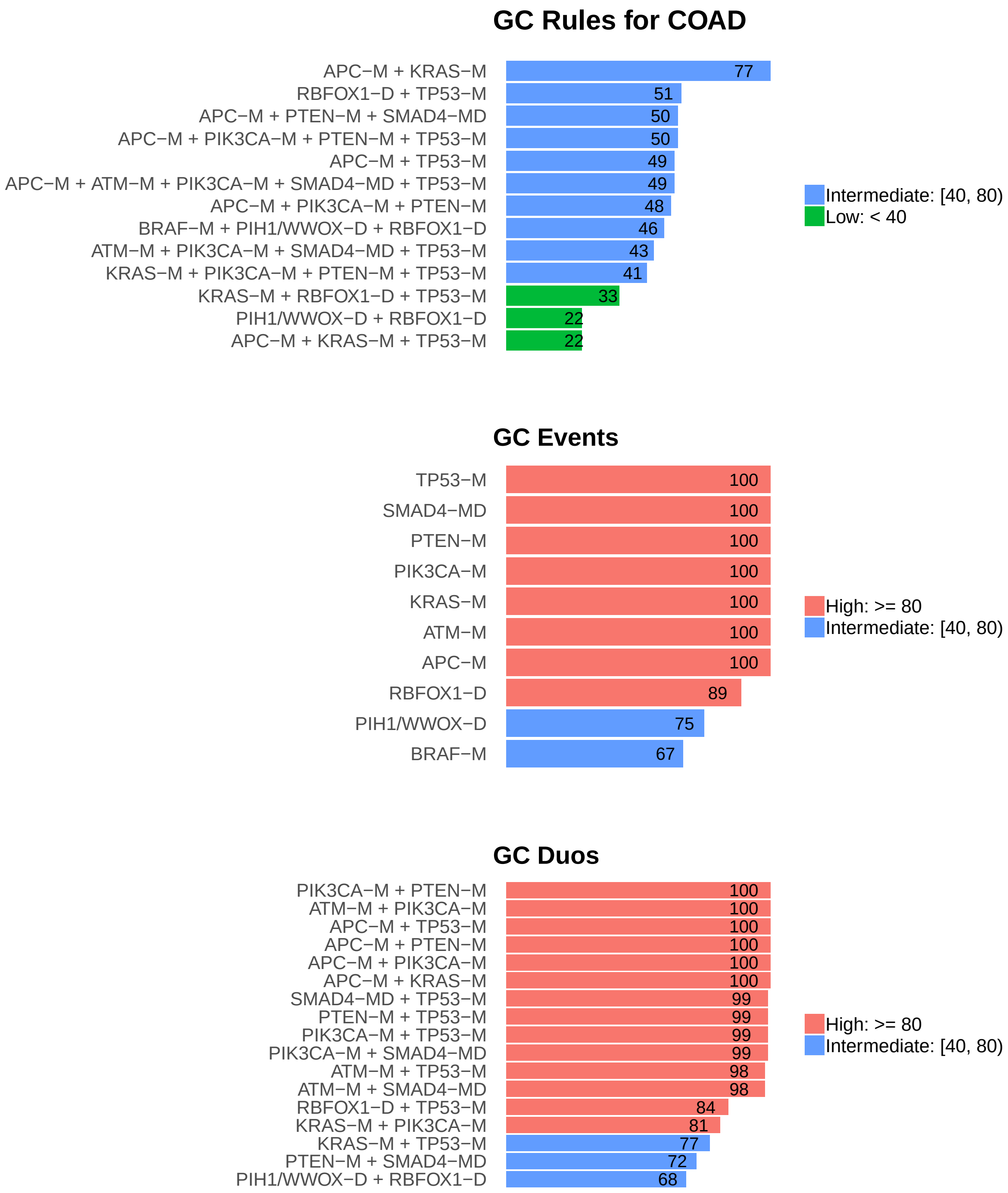

Figure 5: Summary of generalized core results. Bars show confidence levels, which are the percentage of sub-sample iterations containing the observation. Rules (top) and events (middle) that achieve a minimum confidence level of 20 are shown. Duos (bottom) with confidence of at least 60 are shown.

### 5 Dictionary of Copy Number Events

| CNV | Genes | Event__Name |
|---|---|---|
| a2 | SNORA26\|ENSG00000212224.1, CHD6 | CHD6/SNORA26-A |
| d1 | RBFOX1 | RBFOX1-D |
| d2 | RNA5SP475, RN7SL864P, FLRT3, MACROD2 | d2(n=4) |
| d3 | PIH1, WWOX | PIH1/WWOX-D |
| d4 | PARK2 | PARK2-D |
| d5 | RN7SKP248, CCSER1 | CCSER1/RN7SKP248-D |
| d6 | FKSG52, MIR582, PDE4D | FKSG52/MIR582/PDE4D-D |
| d7 | snoU13\|ENSG00000239144.1, BEND5, AGBL4 | AGBL4/BEND5/snoU13-D |
| d8 | U3\|ENSG00000212211.1, NPCDR1, FHIT | FHIT/NPCDR1/U3-D |
| d9 | RN7SL165P, RN7SL501P, PIGV | PIGV/RN7SL165P/RN7SL501P-D |

Dictionary of important CNVs. Copy number events that are within any of the best K rule sets, or within the generalized core rules, or among the 20 most frequent events.
