## Supplementary material for "Identifying Combinations of Cancer Drivers in Individual Patients": TCGA Reports: CRSO_Report_ESCA.pdf

### CRSO Output Report: ESCA

Michael Klein

Apr 19 2019

**4 Generalized Core Rules** **7**

**5 Dictionary of Copy Number Events** **8**

### 1 Heatmaps of D and P

**Number of samples = 184.**
**Number of events = 92.**
**Rule coverage requirement = 6 samples.**
**Rule library size = 827 rules.**

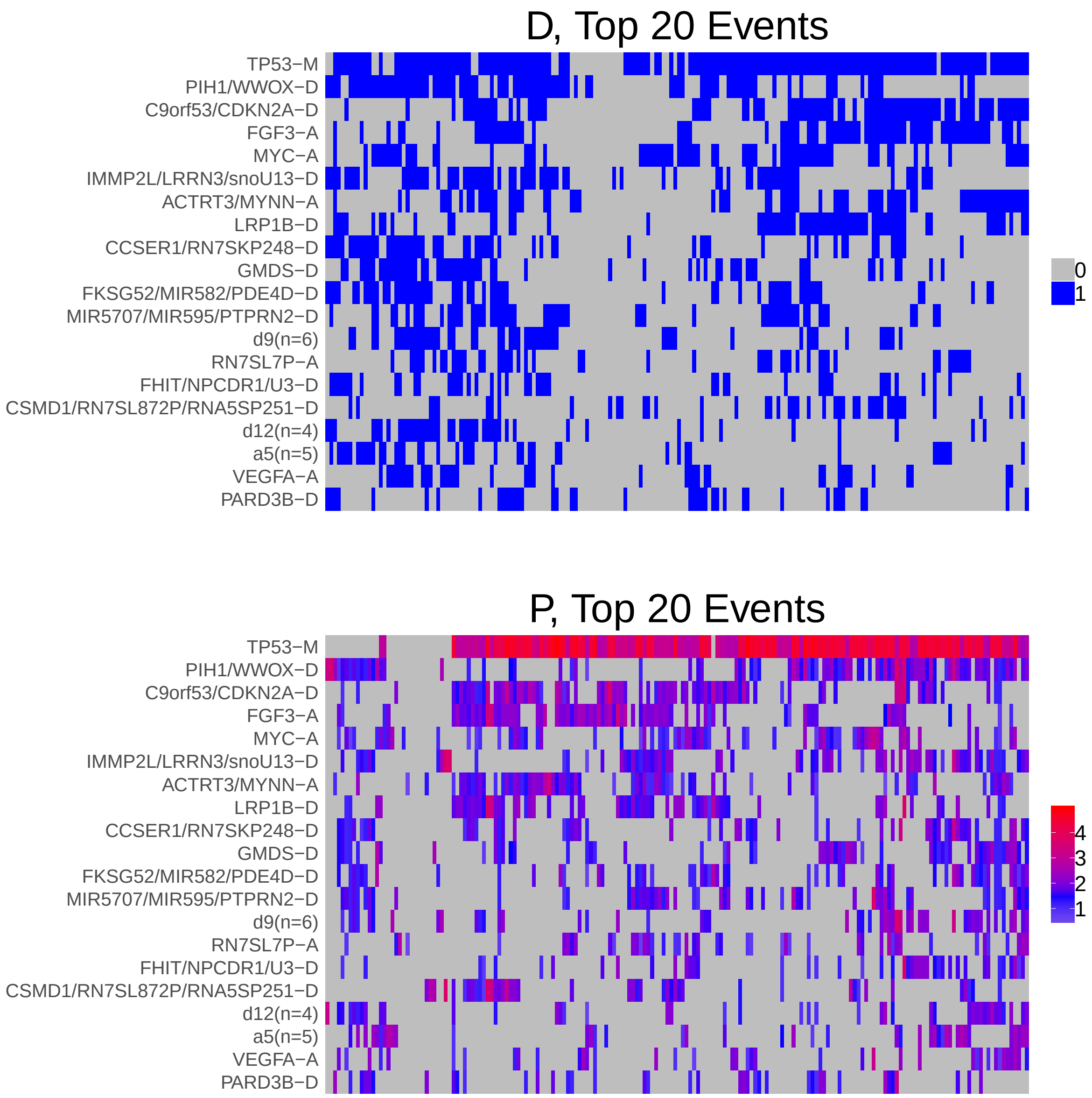

Figure 1: Heatmap of D and P. Events are ordered by frequency, from top to bottom. Event suffix -M: mutation, -A: amplification, -D: deletion, -MD: mutDel, -MA: mutAmp. Samples are ordered using hierarchical clustering. Wild-type events indicated in grey.

### 2 Summary of K Best Rule Sets

#### 2.1 Performance and coverage of best rule sets

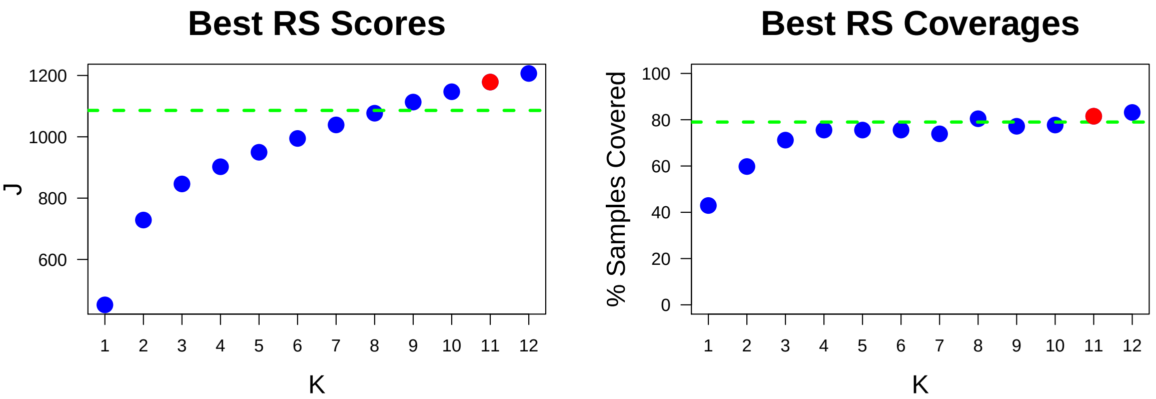

Figure 2: Performance and Coverage of Best Filtered RS for K. The core was determined to be the smallest RS that achieved at least 95% of maximum coverage and at least 90% of maximum performance.

#### 2.2 Table of rules that appear in any best rule set

| ID | Rule | PC | Ks |
|---|---|---|---|
| r7 | C9orf53/CDKN2A-D + FGF3-A + TP53-M | 27 | 2,3,4,5,6,7,8,10,11,12 |
| r2 | MYC-A + TP53-M | 33 | 4,5,6,7,8,9,10,11,12 |
| r6 | GMDS-D + PIH1/WWOX-D + TP53-M | 18 | 7,8,9,10,11,12 |
| r13 | LRP1B-D + TP53-M | 33 | 3,4,5,6,7,8 |
| r14 | SMAD4-MD + TP53-M | 19 | 7,8,9,10,11,12 |
| r1 | PIH1/WWOX-D + TP53-M | 40 | 2,3,4,5,6 |
| r17 | CCSER1/RN7SKP248-D + FKSG52/MIR582/PDE4D-D + IMMP2L/LRRN3/snoU13-D + PIH1/WWOX-D + TP53-M | 7.6 | 7,8,10,11,12 |
| r10 | d9(n=6) + IMMP2L/LRRN3/snoU13-D + PIH1/WWOX-D + TP53-M | 11 | 9,10,11,12 |
| r15 | MIR5707/MIR595/PTPRN2-D + TP53-M | 25 | 9,10,11,12 |
| r16 | ACTRT3/MYNN-A + FGF3-A + IMMP2L/LRRN3/snoU13-D + MIR5707/MIR595/PTPRN2-D + TP53-M | 7.1 | 5,6,7,8 |
| r20 | TP53-M + ZNF750-MD | 14 | 9,10,11,12 |
| r38 | NFE2L2-M + TP53-M | 8.2 | 10,11,12 |
| r28 | FGF3-A + LRP1B-D + TP53-M | 20 | 9,12 |
| r84 | ACTRT3/MYNN-A + CSMD1/RN7SL872P/RNA5SP251-D + FGF3-A + LRP1B-D + TP53-M | 8.7 | 10,11 |
| r268 | CCSER1/RN7SKP248-D + MIR5707/MIR595/PTPRN2-D + PIH1/WWOX-D | 8.2 | 11,12 |
| r5 | ACTRT3/MYNN-A + C9orf53/CDKN2A-D + FGF3-A + TP53-M | 18 | 9 |
| r8 | C9orf53/CDKN2A-D + TP53-M | 43 | 1 |
| r12 | C9orf53/CDKN2A-D + LRP1B-D + TP53-M | 21 | 9 |
| r118 | ACTRT3/MYNN-A + C9orf53/CDKN2A-D + CSMD1/RN7SL872P/RNA5SP251-D + LRP1B-D + TP53-M | 8.2 | 12 |
| r185 | MIR5707/MIR595/PTPRN2-D + PIH1/WWOX-D | 16 | 8 |
| r241 | CCSER1/RN7SKP248-D + d9(n=6) + IMMP2L/LRRN3/snoU13-D + TP53-M | 7.1 | 6 |

**ID** = Rule IDs, rules are numbered according to importance rank determined from phase 1
**PC** = Percent of samples covered **Ks** = Membership in best RS

### 3 Core Rule Set

**Core K = 11.**
**Core rule set coverage = 81.5%.**

#### 3.1 Table of core rule set rules

| ID | Rule | CR | SJR | SJ | NSC | NSA | PC | PA | FracA |
|---|---|---|---|---|---|---|---|---|---|
| r2 | MYC-A + TP53-M | 5 | 5 | 341 | 61 | 13 | 33 | 7.1 | 0.21 |
| r6 | GMDS-D + PIH1/WWOX-D + TP53-M | 40 | 19 | 248 | 34 | 17 | 18 | 9.2 | 0.50 |
| r7 | C9orf53/CDKN2A-D + FGF3-A + TP53-M | 9.5 | 4 | 377 | 49 | 31 | 27 | 17.0 | 0.63 |
| r10 | d9(n=6) + IMMP2L/LRRN3/snoU13-D + PIH1/WWOX-D + TP53-M | 213.5 | 45 | 189 | 20 | 12 | 11 | 6.5 | 0.60 |
| r14 | SMAD4-MD + TP53-M | 36.5 | 35 | 202 | 35 | 9 | 19 | 4.9 | 0.26 |
| r15 | MIR5707/MIR595/PTPRN2-D + TP53-M | 12.5 | 20 | 247 | 46 | 12 | 25 | 6.5 | 0.26 |
| r17 | CCSER1/RN7SKP248-D + FKSG52/MIR582/PDE4D-D + IMMP2L/LRRN3/snoU13-D + PIH1/WWOX-D + TP53-M | 587 | 118.5 | 147 | 14 | 13 | 7.6 | 7.1 | 0.93 |
| r20 | TP53-M + ZNF750-MD | 96 | 105 | 151 | 25 | 9 | 14 | 4.9 | 0.36 |
| r38 | NFE2L2-M + TP53-M | 482 | 244 | 118 | 15 | 11 | 8.2 | 6.0 | 0.73 |
| r84 | ACTRT3/MYNN-A + CSMD1/RN7SL872P/RNA5SP251-D + FGF3-A + LRP1B-D + TP53-M | 415 | 74 | 168 | 16 | 16 | 8.7 | 8.7 | 1.00 |
| r268 | CCSER1/RN7SKP248-D + MIR5707/MIR595/PTPRN2-D + PIH1/WWOX-D | 482 | 551.5 | 71.8 | 15 | 7 | 8.2 | 3.8 | 0.47 |

**ID** = Rule IDs, rules are numbered according to importance rank determined from phase 1
**CR** = Coverage rank
**SJ/SJR** = Single rule performance / performance rank
**NSC/PC** = Number/percent of samples covered
**NSA/PA** = Number/percent of samples assigned
**FracA** = Fraction of covered samples assigned to rule

#### 3.2 Event breakdown of core rule set

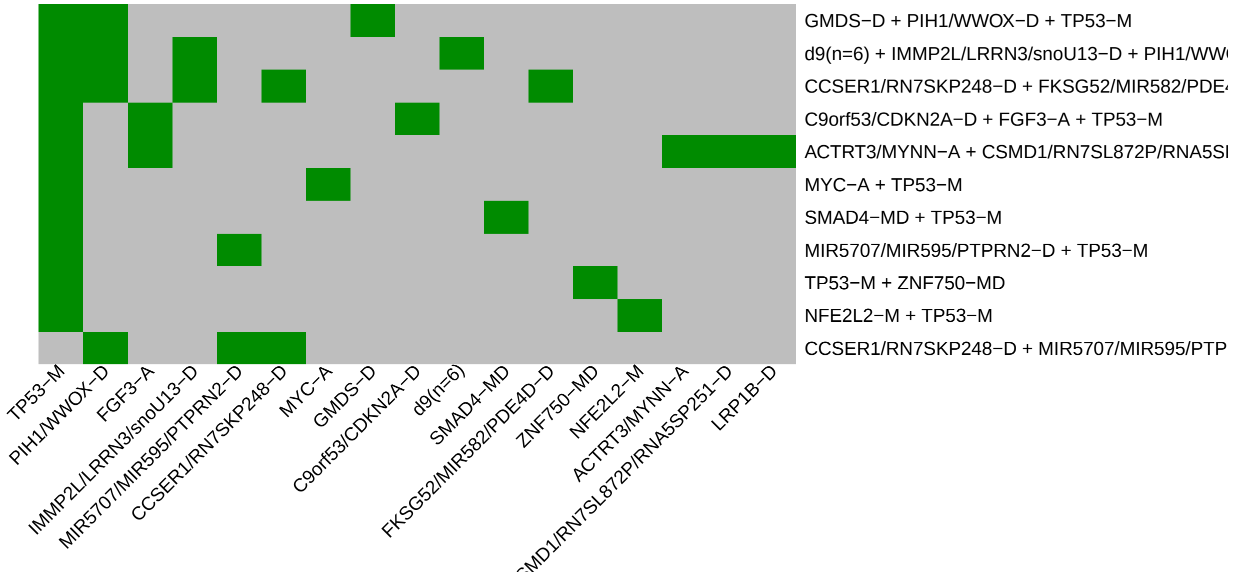

Figure 3: Visualization of core rule set. Rows are rules, columns are events. Events are ordered from left in decreasing rule membership frequency.

##### 3.3 Core Rule Set Penalties: Before and After Assignment

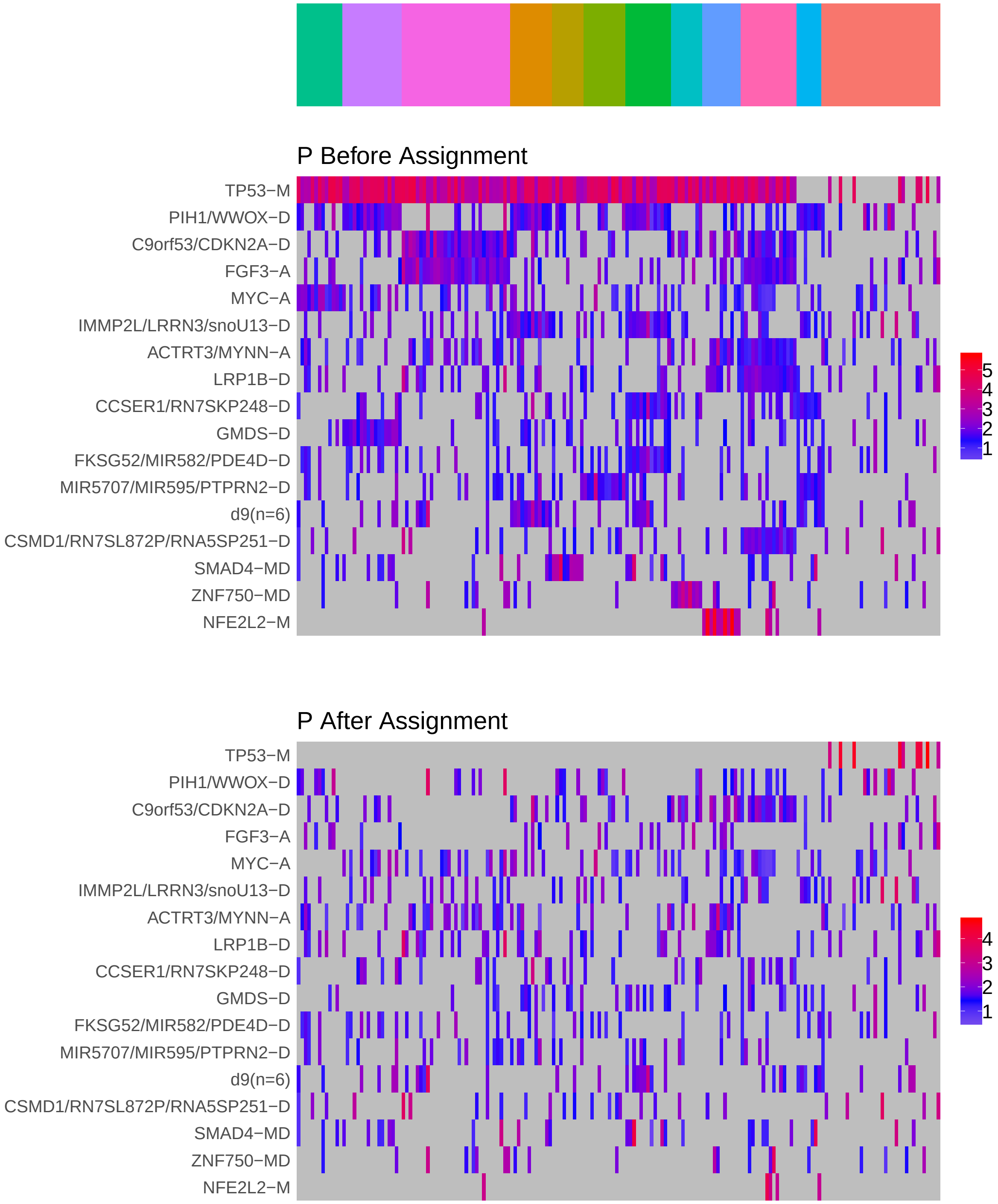

Figure 4: Heatmap of P before and after assignment to core rule set. Events are force ordered by frequency. Samples are force ordered according rule set membership, as indicated by the color bar. The right-most group of samples are not assigned to any rule.

### 4 Generalized Core Rules

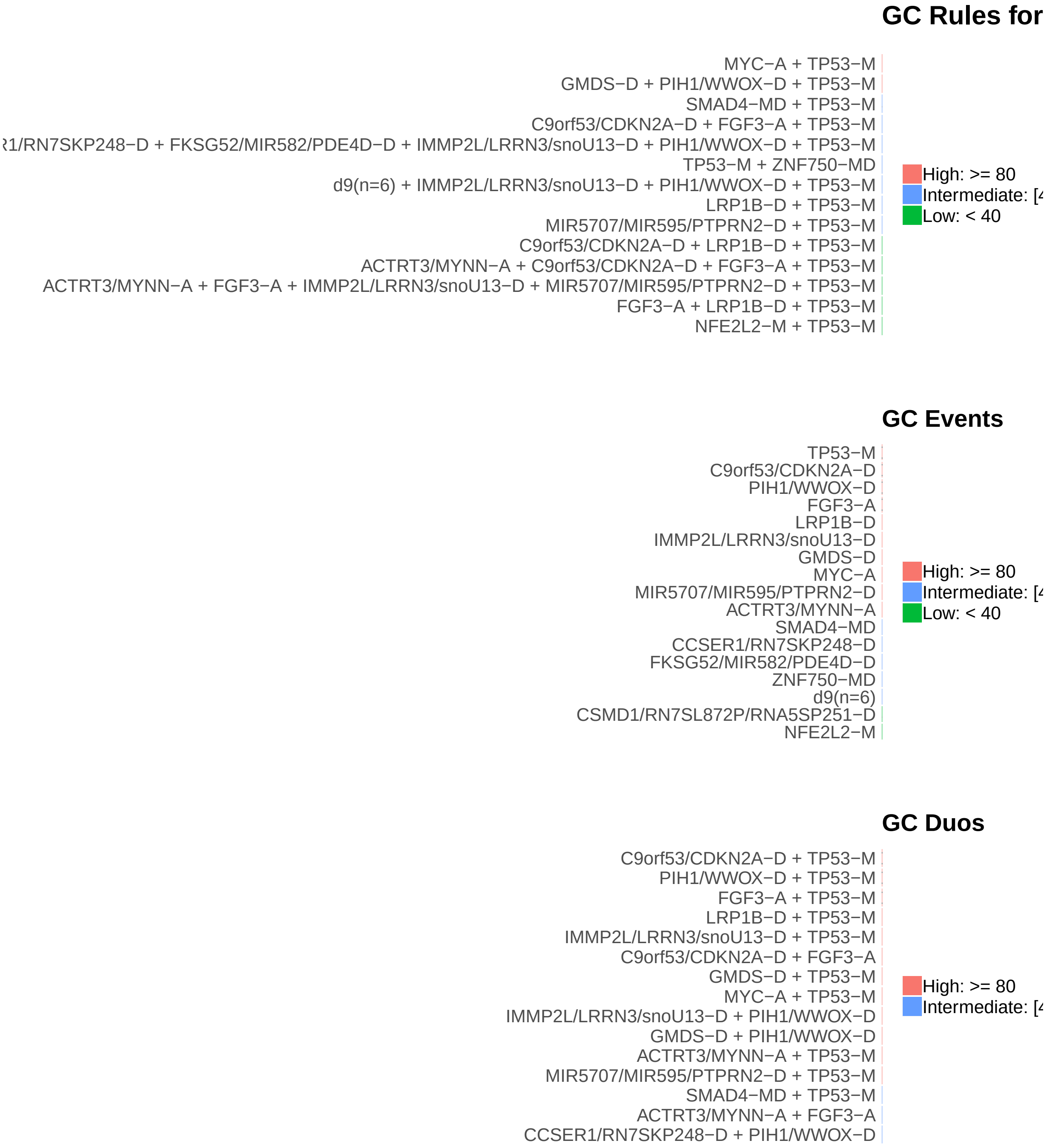

Figure 5: Summary of generalized core results. Bars show confidence levels, which are the percentage of sub-sample iterations containing the observation. Rules (top) and events (middle) that achieve a minimum confidence level of 20 are shown. Duos (bottom) with confidence of at least 60 are shown.

### 5 Dictionary of Copy Number Events

| CNV | Genes | Event_Name |
|---|---|---|
| a1 | FGF3 | FGF3-A |
| a2 | MYC | MYC-A |
| a3 | MYNN, ACTRT3 | ACTRT3/MYNN-A |
| a4 | RN7SL7P | RN7SL7P-A |
| a5 | IKZF3, MIR4728, ERBB2, GRB7, MIEN1 | a5(n=5) |
| a6 | VEGFA | VEGFA-A |
| a7 | RNY4P16 | RNY4P16-A |
| a8 | RNU6ATAC20P | RNU6ATAC20P-A |
| d1 | PIH1, WWOX | PIH1/WWOX-D |
| d2 | CDKN2A, C9orf53 | C9orf53/CDKN2A-D |
| d3 | snoU13\|ENSG00000238922.1, LRRN3, IMMP2L | IMMP2L/LRRN3/snoU13-D |
| d4 | LRP1B | LRP1B-D |
| d5 | RN7SKP248, CCSER1 | CCSER1/RN7SKP248-D |
| d6 | GMDS | GMDS-D |
| d7 | FKSG52, MIR582, PDE4D | FKSG52/MIR582/PDE4D-D |
| d8 | MIR595, PTPRN2, MIR5707 | MIR5707/MIR595/PTPRN2-D |
| d9 | snoU13\|ENSG00000238969.1, MIR548F5, MIR3915, RNA5SP501, DMD, FTHL17 | d9(n=6) |
| d10 | U3\|ENSG00000212211.1, NPCDR1, FHIT | FHIT/NPCDR1/U3-D |
| d11 | RN7SL872P, RNA5SP251, CSMD1 | CSMD1/RN7SL872P/RNA5SP251-D |
| d12 | RNA5SP475, RN7SL864P, FLRT3, MACROD2 | d12(n=4) |
| d13 | PARD3B | PARD3B-D |

Dictionary of important CNVs. Copy number events that are within any of the best K rule sets, or within the generalized core rules, or among the 20 most frequent events.
