## Supplementary material for "Identifying Combinations of Cancer Drivers in Individual Patients": TCGA Reports: CRSO_Report_GBM.pdf

### CRSO Output Report: GBM

Michael Klein

Apr 19 2019

**4 Generalized Core Rules** **6**

**5 Dictionary of Copy Number Events** **7**

### 1 Heatmaps of D and P

**Number of samples = 273.**
**Number of events = 78.**
**Rule coverage requirement = 9 samples.**
**Rule library size = 186 rules.**

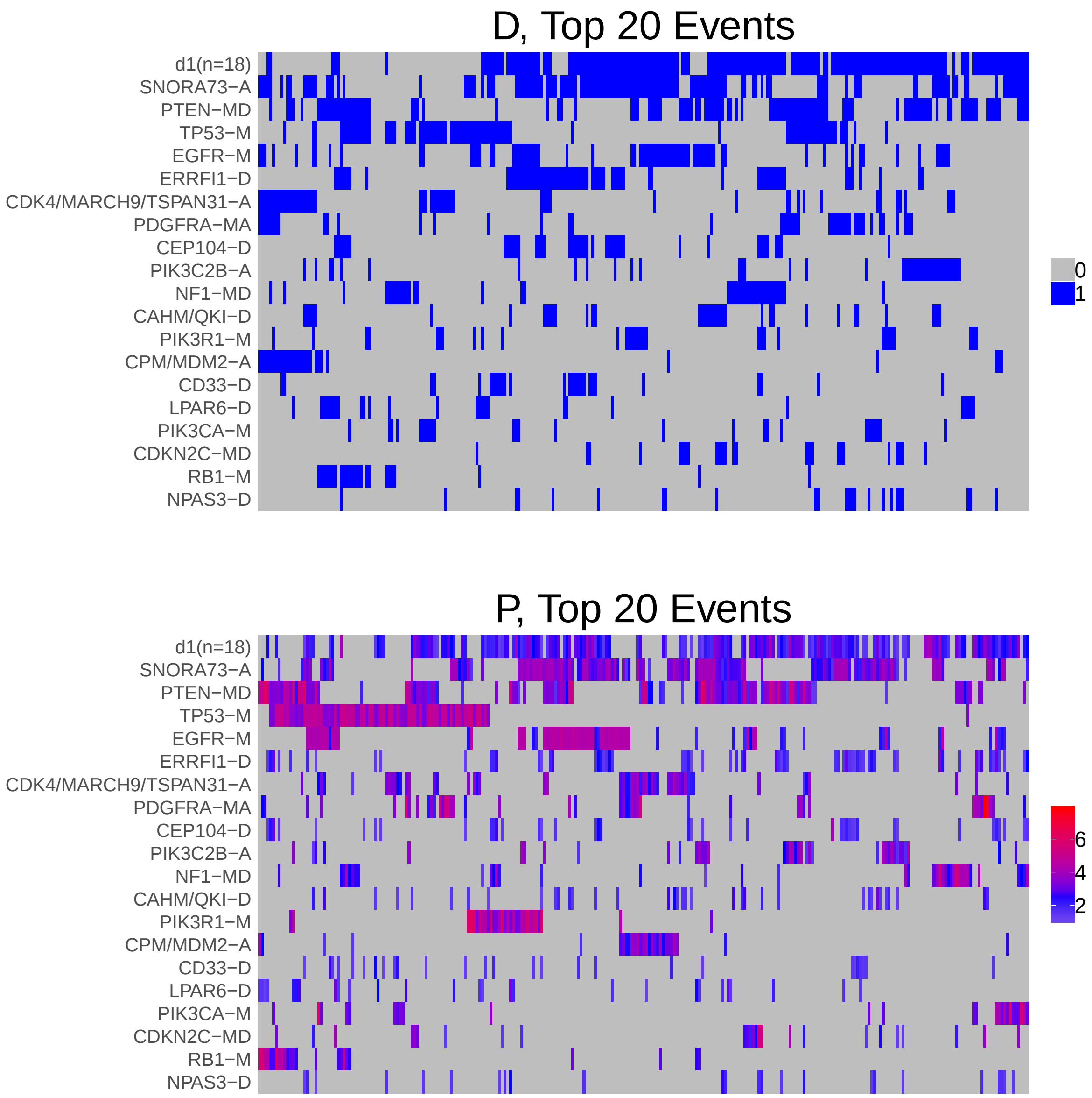

Figure 1: Heatmap of D and P. Events are ordered by frequency, from top to bottom. Event suffix -M: mutation, -A: amplification, -D: deletion, -MD: mutDel, -MA: mutAmp. Samples are ordered using hierarchical clustering. Wild-type events indicated in grey.

### 2 Summary of K Best Rule Sets

#### 2.1 Performance and coverage of best rule sets

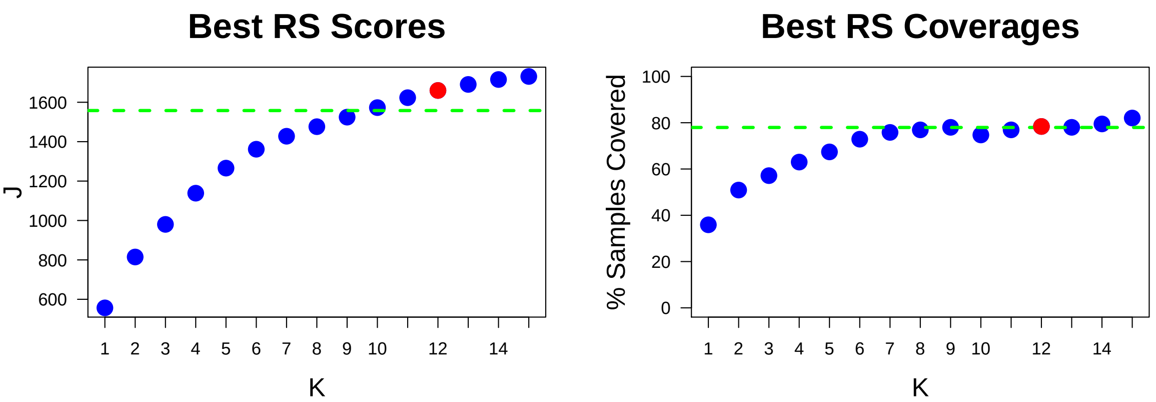

Figure 2: Performance and Coverage of Best Filtered RS for K. The core was determined to be the smallest RS that achieved at least 95% of maximum coverage and at least 90% of maximum performance.

#### 2.2 Table of rules that appear in any best rule set

| ID | Rule | PC | Ks |
|---|---|---|---|
| r1 | d1(n=18) + PTEN-MD | 28 | 2,3,4,5,6,7,8,9,10,11,12,13,14,15 |
| r6 | PTEN-MD + TP53-M | 12 | 3,4,5,6,7,8,9,10,11,12,13,14,15 |
| r5 | CDK4/MARCH9/TSPAN31-A + CPM/MDM2-A | 7.7 | 6,7,8,9,10,11,12,13,14,15 |
| r2 | d1(n=18) + SNORA73-A | 36 | 1,2,3,4,5,6,7,8,9 |
| r8 | CDK4/MARCH9/TSPAN31-A + TP53-M | 7 | 8,9,10,11,12,13,14,15 |
| r11 | ATRX-M + IDH1-M + TP53-M | 3.7 | 8,9,10,11,12,13,14,15 |
| r7 | d1(n=18) + PDGFRA-MA | 11 | 7,8,10,11,12,13,15 |
| r3 | d1(n=18) + EGFR-M + SNORA73-A | 15 | 10,11,12,13,14,15 |
| r4 | EGFR-M + SNORA73-A | 21 | 4,5,6,7,8,9 |
| r10 | d1(n=18) + NF1-MD | 9.9 | 10,11,12,13,14,15 |
| r13 | PTEN-MD + SNORA73-A | 17 | 10,11,12,14,15 |
| r14 | d1(n=18) + ERRFI1-D + SNORA73-A | 13 | 10,11,12,13,14 |
| r16 | d1(n=18) + PIK3R1-M | 8.1 | 11,12,13,14,15 |
| r24 | RB1-M + TP53-M | 5.9 | 12,13,14,15 |
| r12 | IDH1-M + TP53-M | 4.8 | 5,6,7 |
| r46 | d1(n=18) + PIK3C2B-A + SNORA73-A | 6.2 | 13,14,15 |
| r41 | d1(n=18) + PDGFRA-MA + TP53-M | 4 | 9,14 |
| r45 | d1(n=18) + PIK3CA-M | 5.5 | 9,14 |
| r25 | ERRFI1-D + SNORA73-A | 16 | 15 |
| r30 | EGFR-M + PTEN-MD + SNORA73-A | 7 | 13 |
| r40 | PDGFRA-MA + TP53-M | 5.9 | 15 |
| r49 | CEP104-D + d1(n=18) + SNORA73-A | 8.1 | 15 |

**ID** = Rule IDs, rules are numbered according to importance rank determined from phase 1
**PC** = Percent of samples covered **Ks** = Membership in best RS

### 3 Core Rule Set

**Core K = 12.**
**Core rule set coverage = 78.4%.**

#### 3.1 Table of core rule set rules

| ID | Rule | CR | SJR | SJ | NSC | NSA | PC | PA | FracA |
|---|---|---|---|---|---|---|---|---|---|
| r1 | d1(n=18) + PTEN-MD | 2 | 2 | 439 | 76 | 23 | 28 | 8.4 | 0.30 |
| r3 | d1(n=18) + EGFR-M + SNORA73-A | 8 | 4 | 382 | 40 | 37 | 15 | 14.0 | 0.92 |
| r5 | CDK4/MARCH9/TSPAN31-A + CPM/MDM2-A | 30.5 | 47 | 127 | 21 | 17 | 7.7 | 6.2 | 0.81 |
| r6 | PTEN-MD + TP53-M | 13 | 9 | 256 | 33 | 26 | 12 | 9.5 | 0.79 |
| r7 | d1(n=18) + PDGFRA-MA | 15.5 | 16 | 191 | 30 | 14 | 11 | 5.1 | 0.47 |
| r8 | CDK4/MARCH9/TSPAN31-A + TP53-M | 39.5 | 35.5 | 138 | 19 | 11 | 7 | 4.0 | 0.58 |
| r10 | d1(n=18) + NF1-MD | 20 | 24.5 | 149 | 27 | 13 | 9.9 | 4.8 | 0.48 |
| r11 | ATRX-M + IDH1-M + TP53-M | 137 | 43 | 132 | 10 | 10 | 3.7 | 3.7 | 1.00 |
| r13 | PTEN-MD + SNORA73-A | 6 | 6 | 304 | 47 | 20 | 17 | 7.3 | 0.43 |
| r14 | d1(n=18) + ERRFI1-D + SNORA73-A | 9 | 8 | 269 | 36 | 22 | 13 | 8.1 | 0.61 |
| r16 | d1(n=18) + PIK3R1-M | 26.5 | 30 | 143 | 22 | 12 | 8.1 | 4.4 | 0.55 |
| r24 | RB1-M + TP53-M | 55 | 51.5 | 122 | 16 | 9 | 5.9 | 3.3 | 0.56 |

**ID** = Rule IDs, rules are numbered according to importance rank determined from phase 1
**CR** = Coverage rank
**SJ/SJR** = Single rule performance / performance rank
**NSC/PC** = Number/percent of samples covered
**NSA/PA** = Number/percent of samples assigned
**FracA** = Fraction of covered samples assigned to rule

#### 3.2 Event breakdown of core rule set

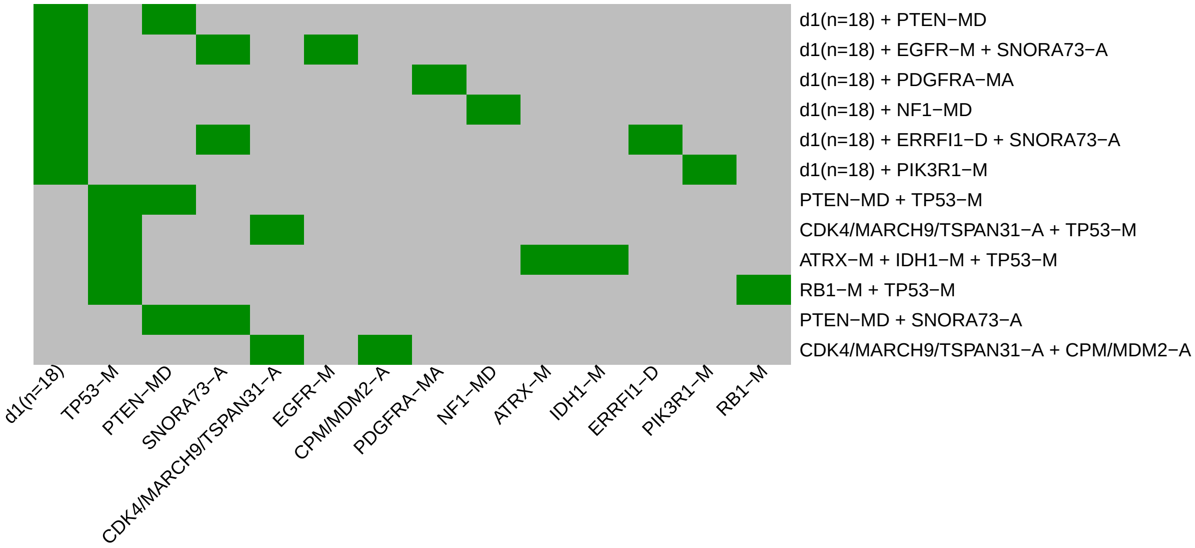

Figure 3: Visualization of core rule set. Rows are rules, columns are events. Events are ordered from left in decreasing rule membership frequency.

#### 3.3 Core Rule Set Penalties: Before and After Assignment

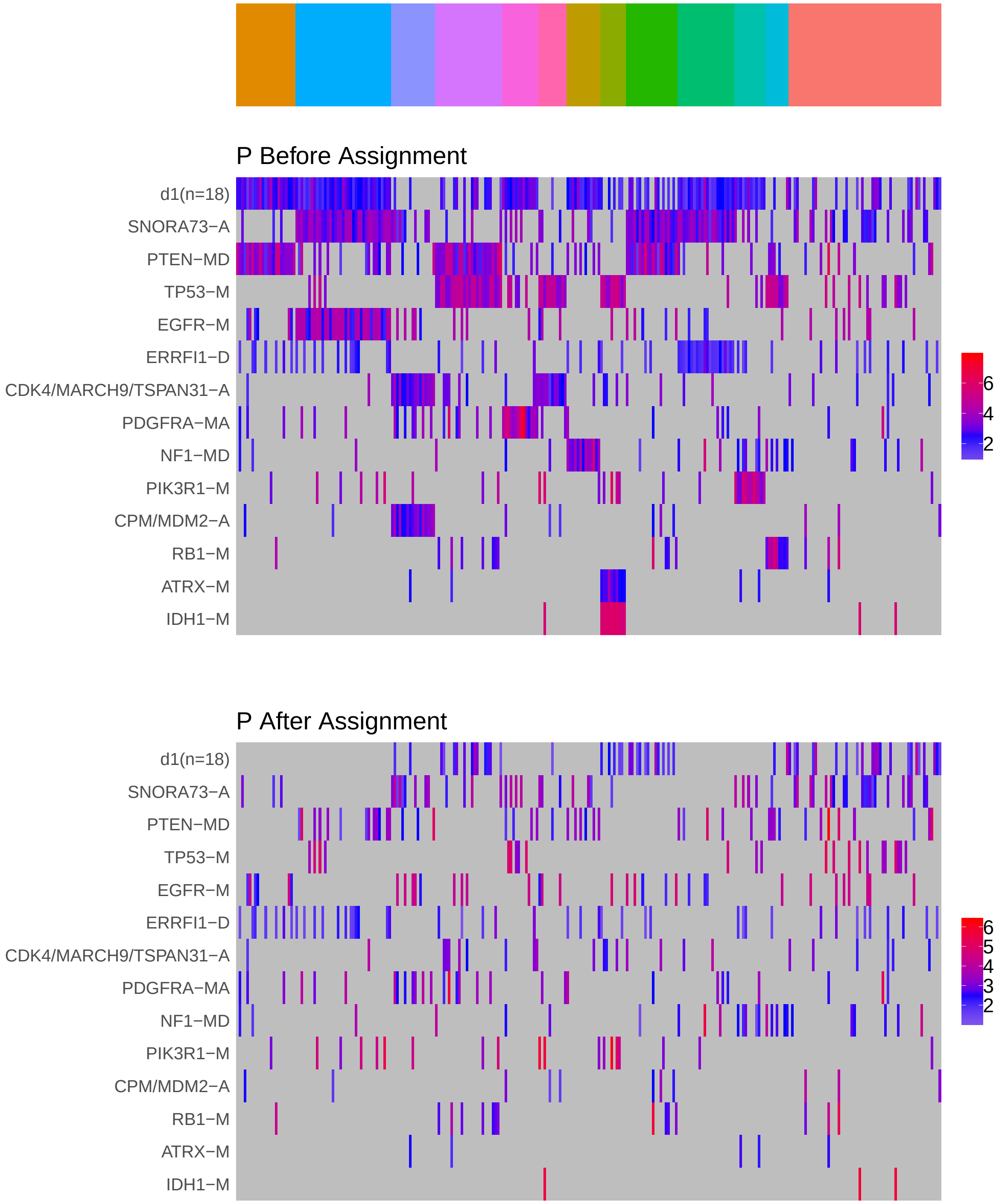

Figure 4: Heatmap of P before and after assignment to core rule set. Events are force ordered by frequency. Samples are force ordered according rule set membership, as indicated by the color bar. The right-most group of samples are not assigned to any rule.

### 4 Generalized Core Rules

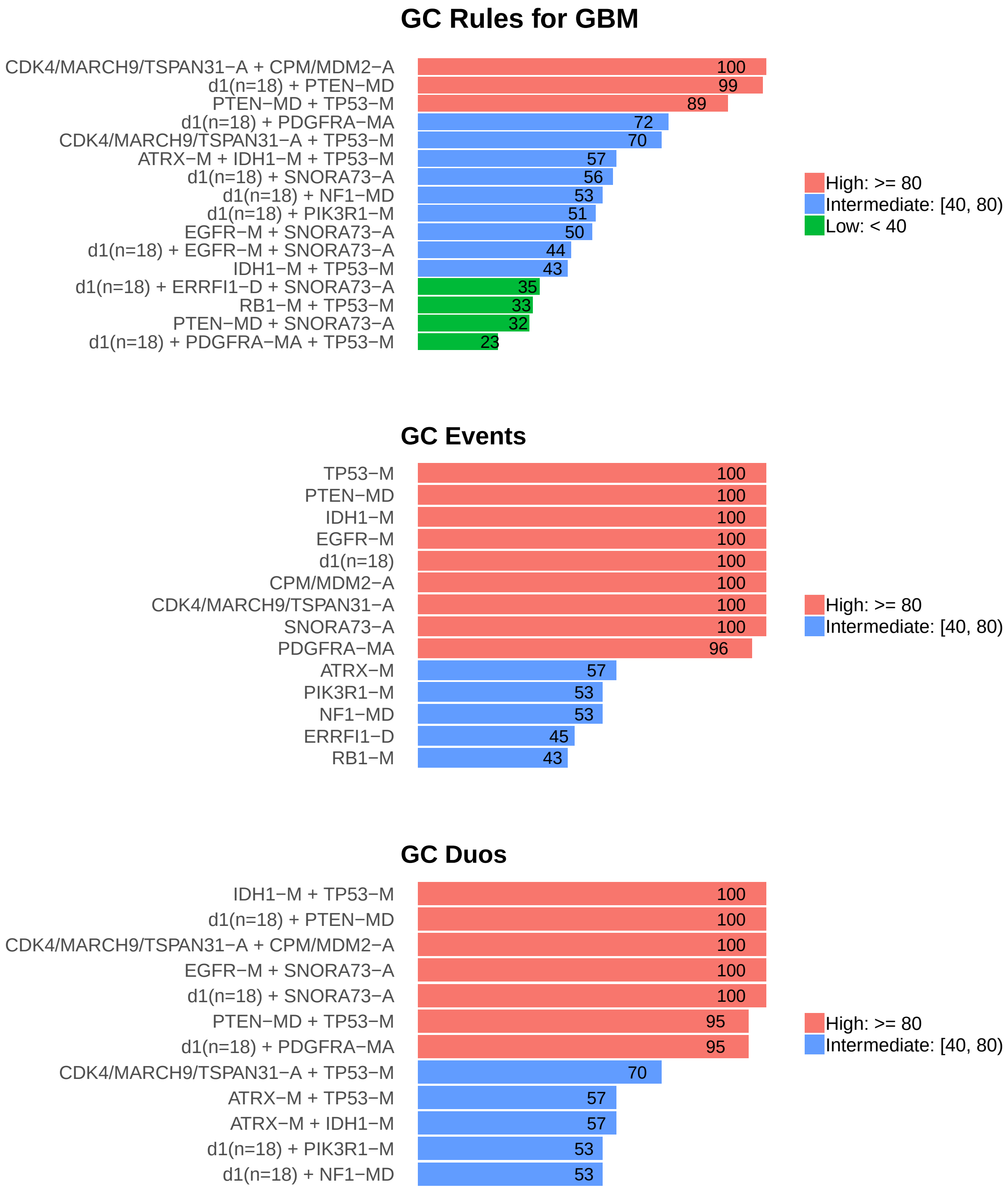

Figure 5: Summary of generalized core results. Bars show confidence levels, which are the percentage of sub-sample iterations containing the observation. Rules (top) and events (middle) that achieve a minimum confidence level of 20 are shown. Duos (bottom) with confidence of at least 50 are shown.

### 5 Dictionary of Copy Number Events

| CNV | Genes | Event_Name |
|---|---|---|
| a1 | SNORA73\|ENSG00000252054.1 | SNORA73-A |
| a2 | CDK4, TSPAN31, MARCH9 | CDK4/MARCH9/TSPAN31-A |
| a4 | PIK3C2B | PIK3C2B-A |
| a5 | CPM, MDM2 | CPM/MDM2-A |
| d1 | DMRTA1, RN7SL151P, SNORD39\|ENSG00000264379.1, MIR31HG, IFNA6, KLHL9, CDKN2A, CDKN2B, IFNA1, IFNA2, IFNA5, IFNA8, IFNA13, IFNA14, MTAP, C9orf53, IFNE, MIR31 | d1(n=18) |
| d2 | ERRFI1 | ERRFI1-D |
| d3 | CEP104 | CEP104-D |
| d4 | CAHM, QKI | CAHM/QKI-D |
| d6 | CD33 | CD33-D |
| d7 | LPAR6 | LPAR6-D |
| d8 | NPAS3 | NPAS3-D |

Dictionary of important CNVs. Copy number events that are within any of the best K rule sets, or within the generalized core rules, or among the 20 most frequent events.
