## Supplementary material for "Identifying Combinations of Cancer Drivers in Individual Patients": TCGA Reports: CRSO_Report_HNSC.pdf

### CRSO Output Report: HNSC

Michael Klein

Apr 19 2019

**4 Generalized Core Rules** **6**

**5 Dictionary of Copy Number Events** **7**

#### 1 Heatmaps of D and P

**Number of samples = 505.**
**Number of events = 94.**
**Rule coverage requirement = 16 samples.**
**Rule library size = 926 rules.**

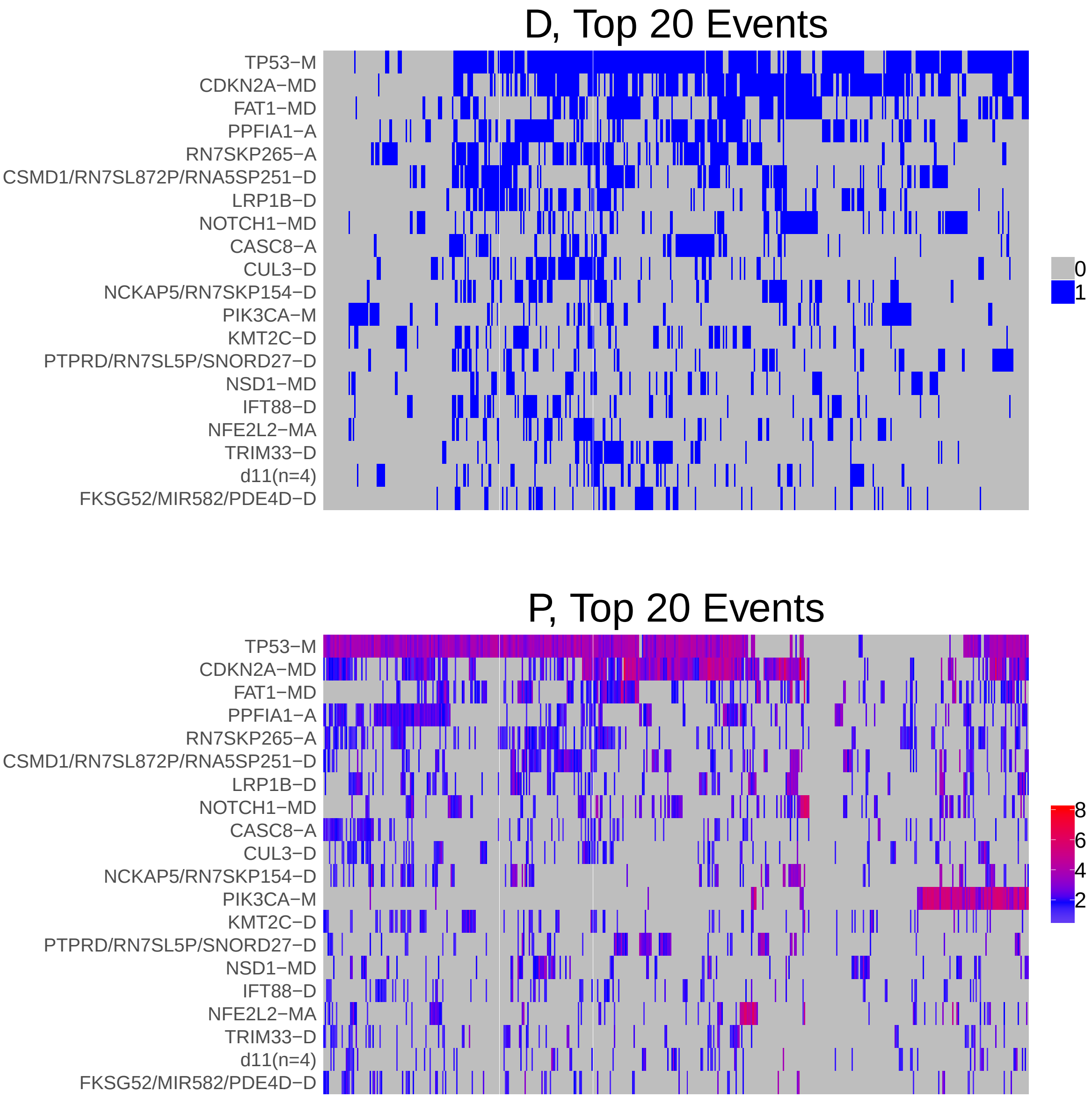

Figure 1: Heatmap of D and P. Events are ordered by frequency, from top to bottom. Event suffix -M: mutation, -A: amplification, -D: deletion, -MD: mutDel, -MA: mutAmp. Samples are ordered using hierarchical clustering. Wild-type events indicated in grey.

#### 2 Summary of K Best Rule Sets

##### 2.1 Performance and coverage of best rule sets

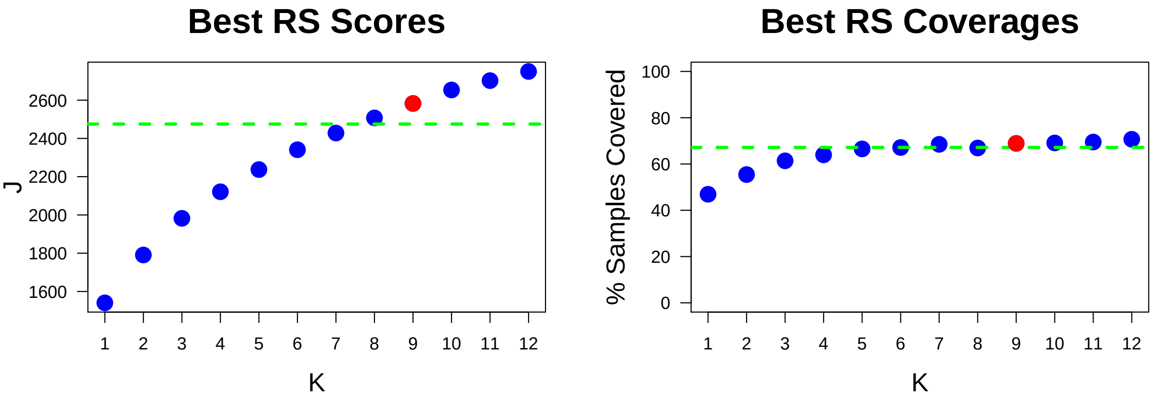

Figure 2: Performance and Coverage of Best Filtered RS for K. The core was determined to be the smallest RS that achieved at least 95% of maximum coverage and at least 90% of maximum performance.

##### 2.2 Table of rules that appear in any best rule set

| **ID** | **Rule** | **PC** | **Ks** |
|---|---|---|---|
| r4 | FAT1-MD + TP53-M | 27 | 2,3,4,5,6,7,8,9,10,11,12 |
| r6 | CASP8-M + HRAS-M | 3.8 | 5,6,7,8,9,10,11,12 |
| r1 | CDKN2A-MD + TP53-M | 47 | 1,2,3,4,5,6,7 |
| r2 | CDKN2A-MD + PPFIA1-A + TP53-M | 21 | 8,9,10,11,12 |
| r3 | PPFIA1-A + TP53-M | 29 | 3,4,5,6,7 |
| r7 | CDKN2A-MD + PIK3CA-M + TP53-M | 7.5 | 8,9,10,11,12 |
| r8 | RN7SKP265-A + TP53-M | 27 | 8,9,10,11,12 |
| r13 | CDKN2A-MD + LRP1B-D + TP53-M | 12 | 8,9,10,11,12 |
| r29 | CDKN2A-MD + FAT1-MD + NOTCH1-MD | 8.5 | 4,5,6,7,12 |
| r12 | CDKN2A-MD + NOTCH1-MD | 15 | 8,9,10,11 |
| r17 | EGFR-A + TP53-M | 11 | 9,10,11,12 |
| r9 | CSMD1/RN7SL872P/RNA5SP251-D + TP53-M | 22 | 8,9,12 |
| r14 | NOTCH1-MD + TP53-M | 15 | 10,11,12 |
| r11 | PIK3CA-M + TP53-M | 11 | 6,7 |
| r21 | CDKN2A-MD + CSMD1/RN7SL872P/RNA5SP251-D + TP53-M | 15 | 10,11 |
| r24 | CDKN2A-MD + NFE2L2-MA + TP53-M | 8.7 | 11,12 |
| r73 | CASC8-A + PPFIA1-A + TP53-M | 8.9 | 12 |
| r78 | LRP1B-D + NCKAP5/RN7SKP154-D + TP53-M | 6.9 | 7 |

**ID** = Rule IDs, rules are numbered according to importance rank determined from phase 1
**PC** = Percent of samples covered **Ks** = Membership in best RS

### 3 Core Rule Set

**Core K = 9.**
**Core rule set coverage = 68.9%.**

#### 3.1 Table of core rule set rules

| ID | Rule | CR | SJR | SJ | NSC | NSA | PC | PA | FracA |
|---|---|---|---|---|---|---|---|---|---|
| r2 | CDKN2A-MD + PPFIA1-A + TP53-M | 8 | 2 | 892 | 107 | 87 | 21 | 17.0 | 0.81 |
| r4 | FAT1-MD + TP53-M | 3 | 5 | 768 | 137 | 48 | 27 | 9.5 | 0.35 |
| r6 | CASP8-M + HRAS-M | 591.5 | 301.5 | 166 | 19 | 16 | 3.8 | 3.2 | 0.84 |
| r7 | CDKN2A-MD + PIK3CA-M + TP53-M | 109.5 | 18 | 426 | 38 | 37 | 7.5 | 7.3 | 0.97 |
| r8 | RN7SKP265-A + TP53-M | 4 | 6 | 695 | 136 | 39 | 27 | 7.7 | 0.29 |
| r9 | CSMD1/RN7SL872P/RNA5SP251-D + TP53-M | 7 | 8 | 621 | 112 | 39 | 22 | 7.7 | 0.35 |
| r12 | CDKN2A-MD + NOTCH1-MD | 17 | 20 | 418 | 76 | 26 | 15 | 5.1 | 0.34 |
| r13 | CDKN2A-MD + LRP1B-D + TP53-M | 25 | 13 | 516 | 63 | 34 | 12 | 6.7 | 0.54 |
| r17 | EGFR-A + TP53-M | 44 | 53 | 321 | 55 | 22 | 11 | 4.4 | 0.40 |

**ID** = Rule IDs, rules are numbered according to importance rank determined from phase 1
**CR** = Coverage rank
**SJ/SJR** = Single rule performance / performance rank
**NSC/PC** = Number/percent of samples covered
**NSA/PA** = Number/percent of samples assigned
**FracA** = Fraction of covered samples assigned to rule

#### 3.2 Event breakdown of core rule set

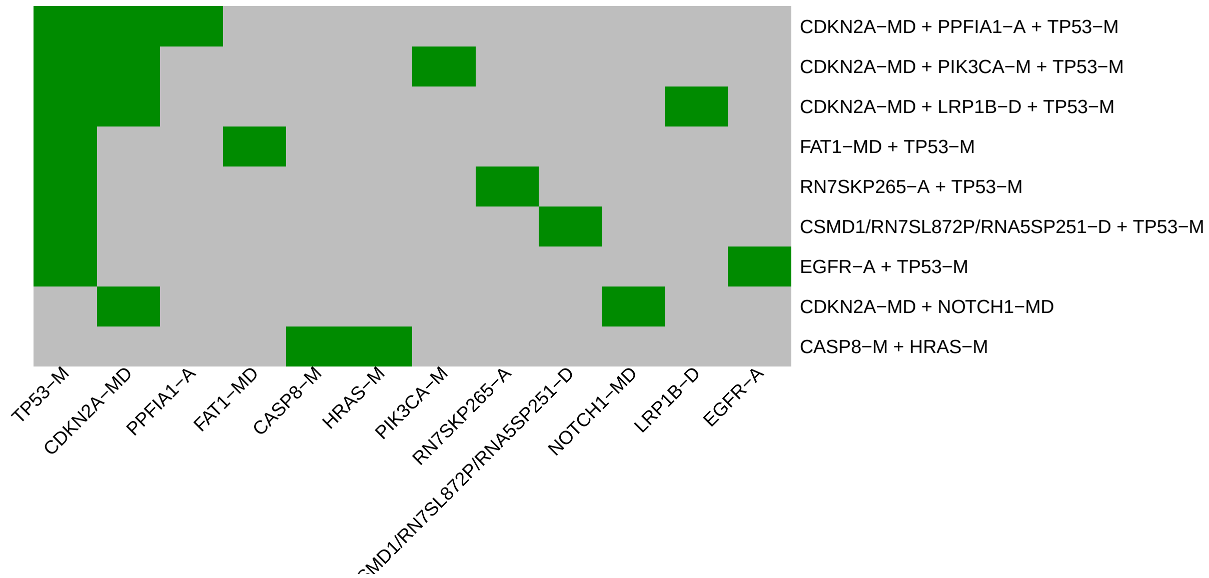

Figure 3: Visualization of core rule set. Rows are rules, columns are events. Events are ordered from left in decreasing rule membership frequency.

#### 3.3 Core Rule Set Penalties: Before and After Assignment

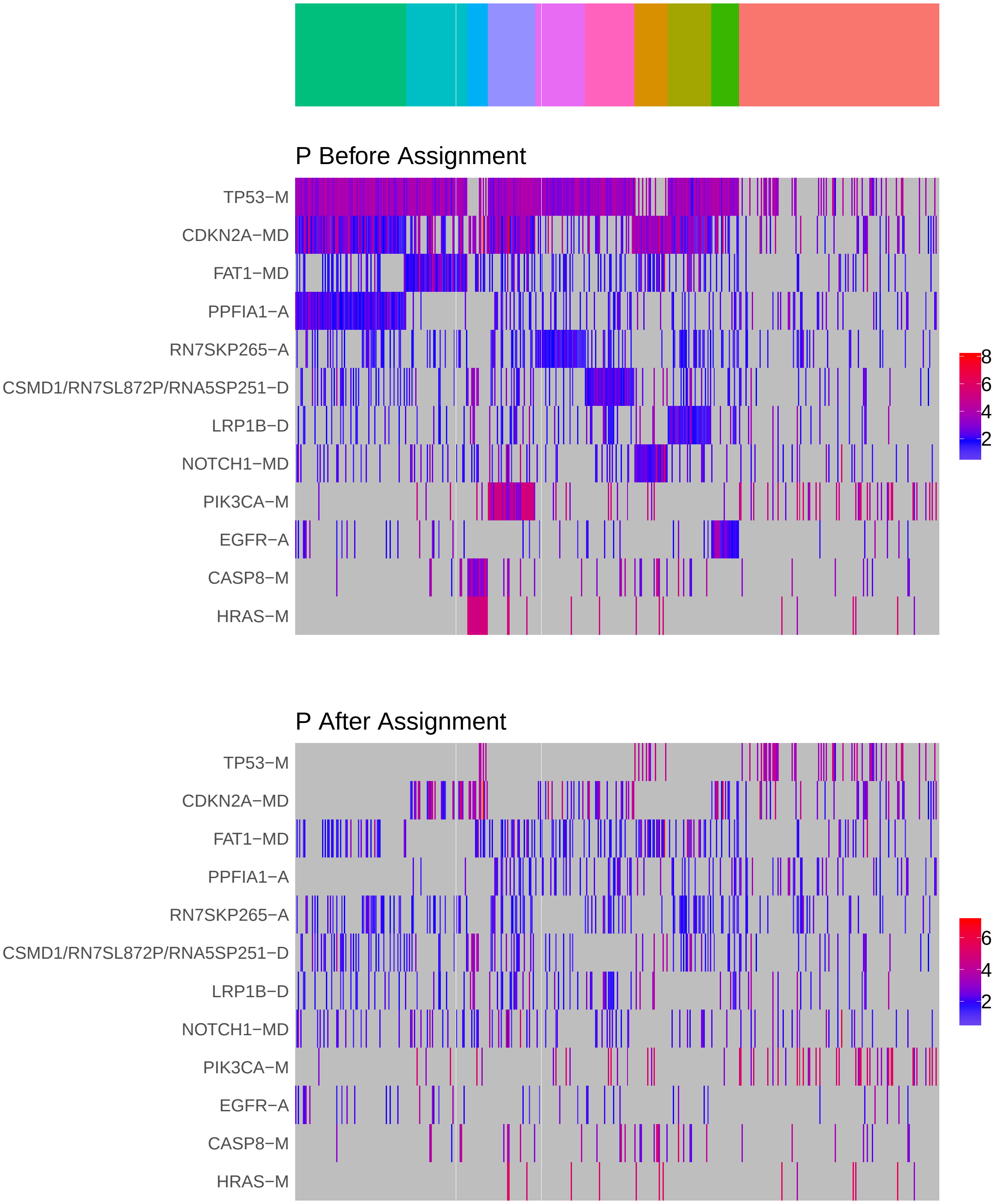

Figure 4: Heatmap of P before and after assignment to core rule set. Events are force ordered by frequency. Samples are force ordered according rule set membership, as indicated by the color bar. The right-most group of samples are not assigned to any rule.

### 4 Generalized Core Rules

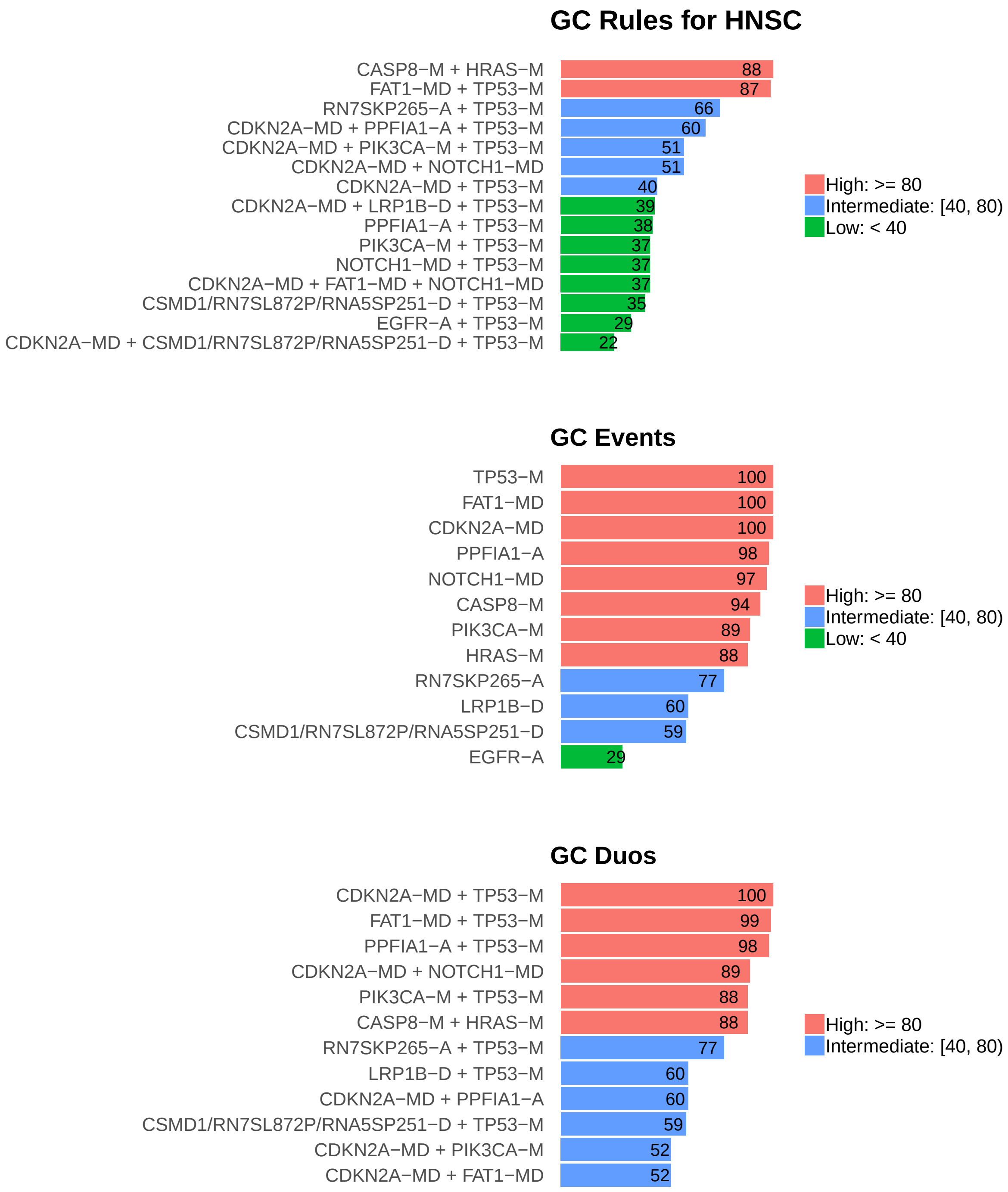

Figure 5: Summary of generalized core results. Bars show confidence levels, which are the percentage of sub-sample iterations containing the observation. Rules (top) and events (middle) that achieve a minimum confidence level of 20 are shown. Duos (bottom) with confidence of at least 50 are shown.

### 5 Dictionary of Copy Number Events

| CNV | Genes | Event__Name |
|---|---|---|
| a1 | PPFIA1 | PPFIA1-A |
| a2 | RN7SKP265 | RN7SKP265-A |
| a3 | CASC8 | CASC8-A |
| a4 | EGFR | EGFR-A |
| d2 | RN7SL872P, RNA5SP251, CSMD1 | CSMD1/RN7SL872P/RNA5SP251-D |
| d3 | LRP1B | LRP1B-D |
| d4 | CUL3 | CUL3-D |
| d5 | RN7SKP154, NCKAP5 | NCKAP5/RN7SKP154-D |
| d6 | KMT2C | KMT2C-D |
| d7 | RN7SL5P, SNORD27\|ENSG00000251699.1, PTPRD | PTPRD/RN7SL5P/SNORD27-D |
| d9 | IFT88 | IFT88-D |
| d10 | TRIM33 | TRIM33-D |
| d11 | ATP5D, STK11, MIDN, C19orf26 | d11(n=4) |
| d12 | FKSG52, MIR582, PDE4D | FKSG52/MIR582/PDE4D-D |

Dictionary of important CNVs. Copy number events that are within any of the best K rule sets, or within the generalized core rules, or among the 20 most frequent events.
