## Supplementary material for "Identifying Combinations of Cancer Drivers in Individual Patients": TCGA Reports: CRSO_Report_KIRC.pdf

### CRSO Output Report: KIRC

Michael Klein

Apr 19 2019

**4 Generalized Core Rules** **6**

**5 Dictionary of Copy Number Events** **7**

### 1 Heatmaps of D and P

**Number of samples = 433.**
**Number of events = 39.**
**Rule coverage requirement = 13 samples.**
**Rule library size = 18 rules.**

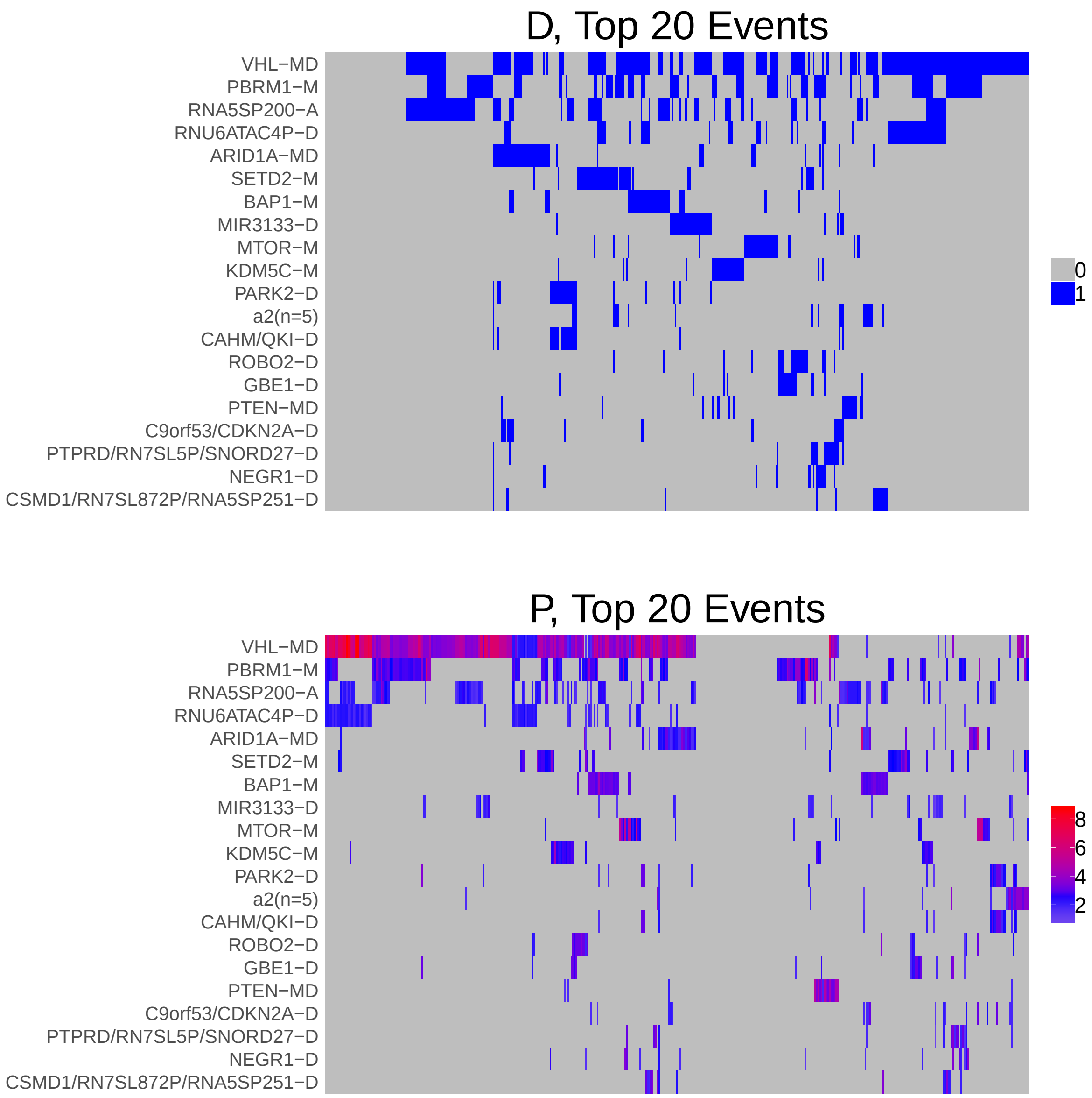

Figure 1: Heatmap of D and P. Events are ordered by frequency, from top to bottom. Event suffix -M: mutation, -A: amplification, -D: deletion, -MD: mutDel, -MA: mutAmp. Samples are ordered using hierarchical clustering. Wild-type events indicated in grey.

### 2 Summary of K Best Rule Sets

#### 2.1 Performance and coverage of best rule sets

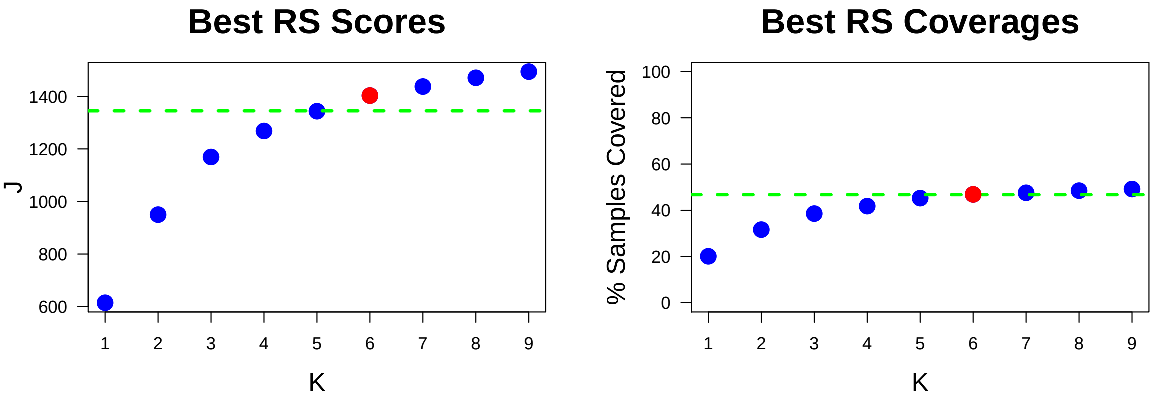

Figure 2: Performance and Coverage of Best Filtered RS for K. The core was determined to be the smallest RS that achieved at least 95% of maximum coverage and at least 90% of maximum performance.

#### 2.2 Table of rules that appear in any best rule set

| ID | Rule | PC | Ks |
|---|---|---|---|
| r1 | PBRM1-M + VHL-MD | 20 | 1,2,3,4,5,6,7,8,9 |
| r3 | RNA5SP200-A + VHL-MD | 17 | 2,3,4,5,6,7,8,9 |
| r2 | RNU6ATAC4P-D + VHL-MD | 15 | 3,4,5,6,7,8,9 |
| r4 | ARID1A-MD + VHL-MD | 7.2 | 4,5,6,7,8,9 |
| r6 | CAHM/QKI-D + PARK2-D | 4.4 | 5,6,7,8,9 |
| r7 | BAP1-M + VHL-MD | 5.5 | 6,7,8,9 |
| r5 | MTOR-M + VHL-MD | 3.9 | 7,8,9 |
| r8 | SETD2-M + VHL-MD | 5.5 | 8,9 |
| r13 | KDM5C-M + VHL-MD | 3.7 | 9 |

**ID** = Rule IDs, rules are numbered according to importance rank determined from phase 1
**PC** = Percent of samples covered **Ks** = Membership in best RS

### 3 Core Rule Set

**Core K = 6.**
**Core rule set coverage = 46.9%.**

#### 3.1 Table of core rule set rules

| ID | Rule | CR | SJR | SJ | NSC | NSA | PC | PA | FracA |
|---|---|---|---|---|---|---|---|---|---|
| r1 | PBRM1-M + VHL-MD | 1 | 1 | 615 | 87 | 77 | 20 | 18.0 | 0.89 |
| r2 | RNU6ATAC4P-D + VHL-MD | 3 | 3 | 459 | 65 | 38 | 15 | 8.8 | 0.58 |
| r3 | RNA5SP200-A + VHL-MD | 2 | 2 | 475 | 72 | 34 | 17 | 7.9 | 0.47 |
| r4 | ARID1A-MD + VHL-MD | 4.5 | 5 | 201 | 31 | 17 | 7.2 | 3.9 | 0.55 |
| r6 | CAHM/QKI-D + PARK2-D | 13 | 16 | 95.3 | 19 | 15 | 4.4 | 3.5 | 0.79 |
| r7 | BAP1-M + VHL-MD | 6.5 | 8 | 169 | 24 | 22 | 5.5 | 5.1 | 0.92 |

**ID** = Rule IDs, rules are numbered according to importance rank determined from phase 1
**CR** = Coverage rank
**SJ/SJR** = Single rule performance / performance rank
**NSC/PC** = Number/percent of samples covered
**NSA/PA** = Number/percent of samples assigned
**FracA** = Fraction of covered samples assigned to rule

#### 3.2 Event breakdown of core rule set

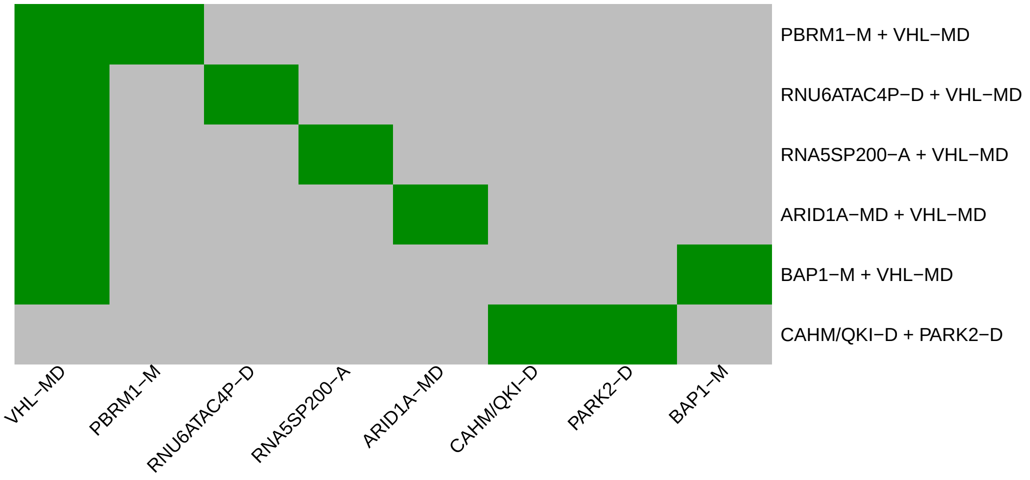

Figure 3: Visualization of core rule set. Rows are rules, columns are events. Events are ordered from left in decreasing rule membership frequency.

##### 3.3 Core Rule Set Penalties: Before and After Assignment

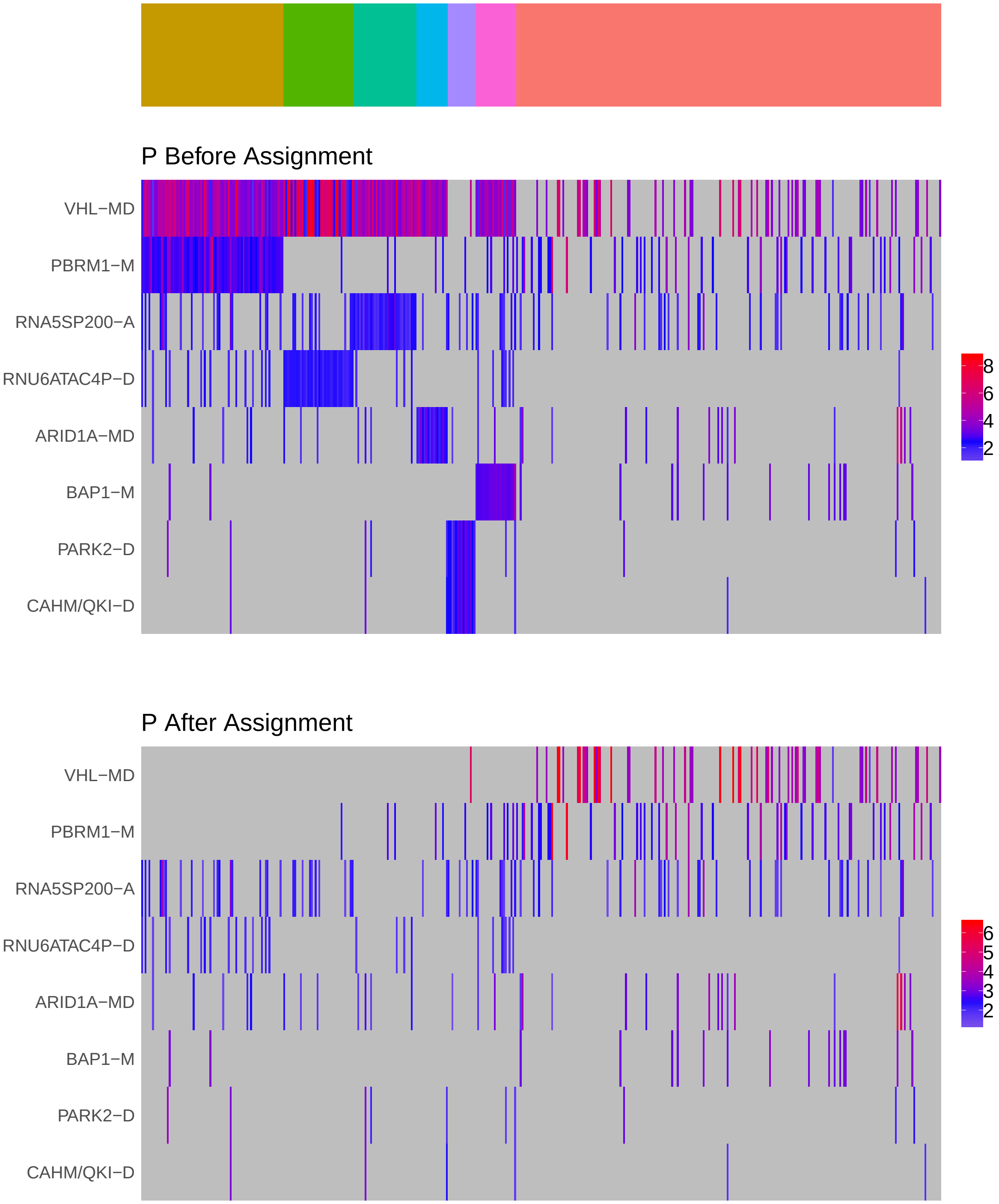

Figure 4: Heatmap of P before and after assignment to core rule set. Events are force ordered by frequency. Samples are force ordered according rule set membership, as indicated by the color bar. The right-most group of samples are not assigned to any rule.

### 4 Generalized Core Rules

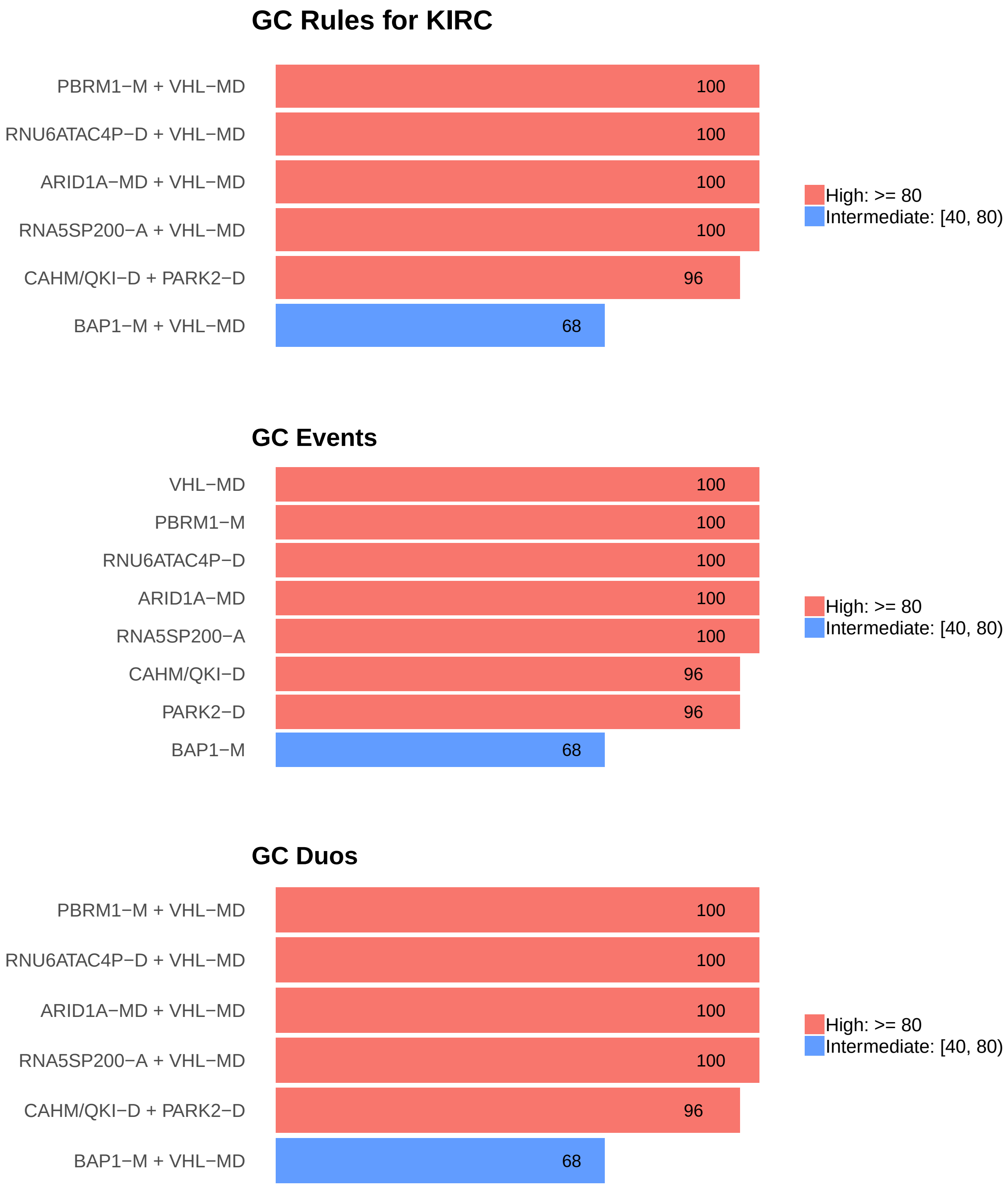

Figure 5: Summary of generalized core results. Bars show confidence levels, which are the percentage of sub-sample iterations containing the observation. Rules (top) and events (middle) that achieve a minimum confidence level of 20 are shown. Duos (bottom) with confidence of at least 50 are shown.

### 5 Dictionary of Copy Number Events

| CNV | Genes | Event_Name |
|---|---|---|
| a1 | RNA5SP200 | RNA5SP200-A |
| a2 | GNB4, PIK3CA, KCNMB3, ZNF639, MFN1 | a2(n=5) |
| d2 | RNU6ATAC4P | RNU6ATAC4P-D |
| d4 | MIR3133 | MIR3133-D |
| d5 | PARK2 | PARK2-D |
| d6 | CAHM, QKI | CAHM/QKI-D |
| d7 | ROBO2 | ROBO2-D |
| d8 | GBE1 | GBE1-D |
| d9 | CDKN2A, C9orf53 | C9orf53/CDKN2A-D |
| d10 | RN7SL5P, SNORD27\|ENSG00000251699.1, PTPRD | PTPRD/RN7SL5P/SNORD27-D |
| d11 | RN7SL872P, RNA5SP251, CSMD1 | CSMD1/RN7SL872P/RNA5SP251-D |
| d12 | NEGR1 | NEGR1-D |

Dictionary of important CNVs. Copy number events that are within any of the best K rule sets, or within the generalized core rules, or among the 20 most frequent events.
