## Supplementary material for "Identifying Combinations of Cancer Drivers in Individual Patients": TCGA Reports: CRSO_Report_LGG.pdf

### CRSO Output Report: LGG

Michael Klein

Apr 19 2019

**4 Generalized Core Rules** **6**

**5 Dictionary of Copy Number Events** **7**

### 1 Heatmaps of D and P

**Number of samples = 513.**
**Number of events = 67.**
**Rule coverage requirement = 16 samples.**
**Rule library size = 210 rules.**

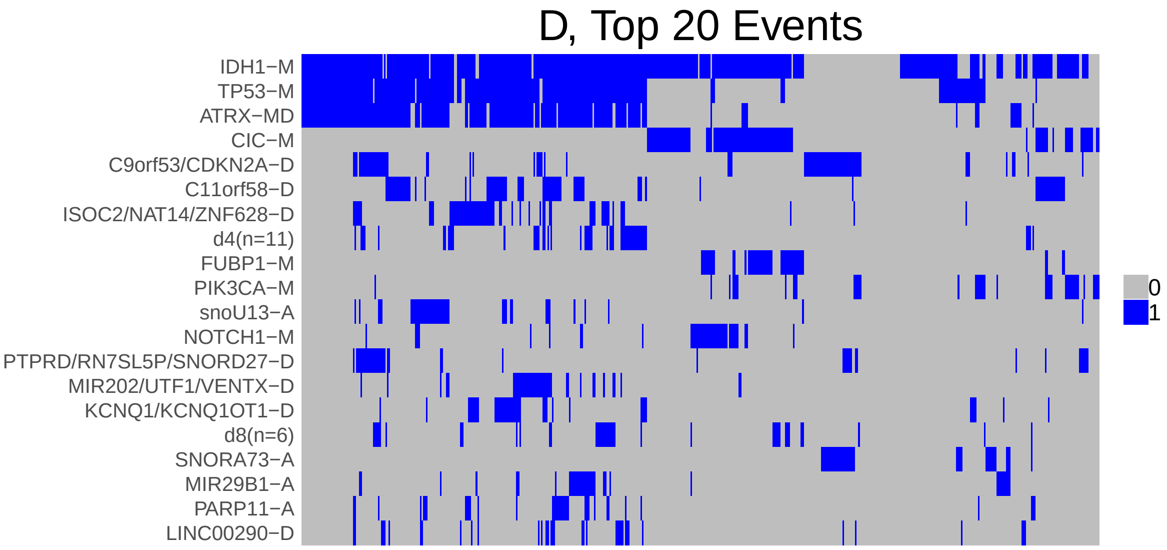

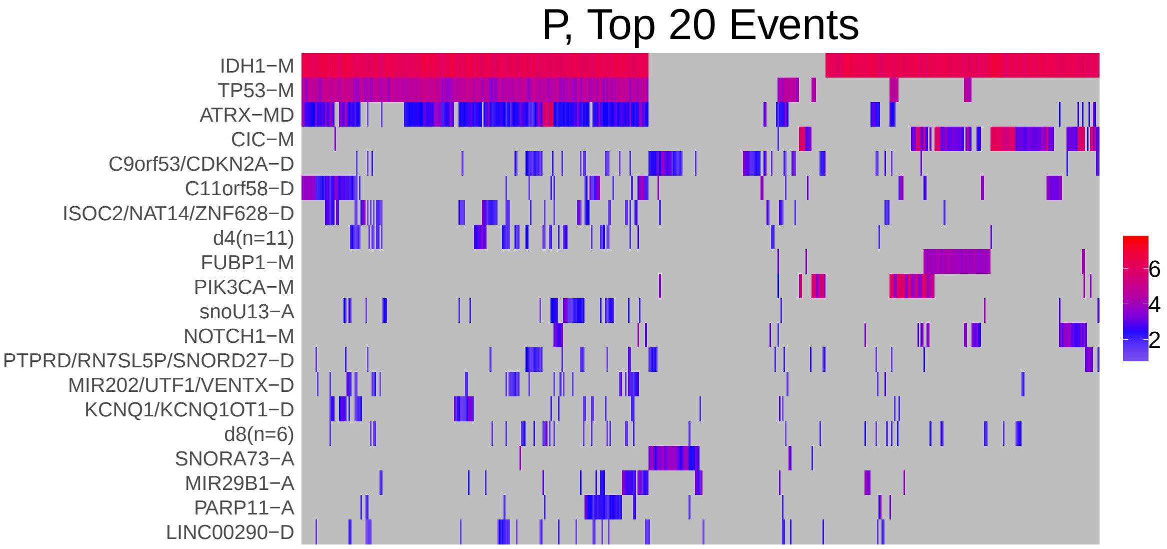

Figure 1: Heatmap of D and P. Events are ordered by frequency, from top to bottom. Event suffix -M: mutation, -A: amplification, -D: deletion, -MD: mutDel, -MA: mutAmp. Samples are ordered using hierarchical clustering. Wild-type events indicated in grey.

### 2 Summary of K Best Rule Sets

#### 2.1 Performance and coverage of best rule sets

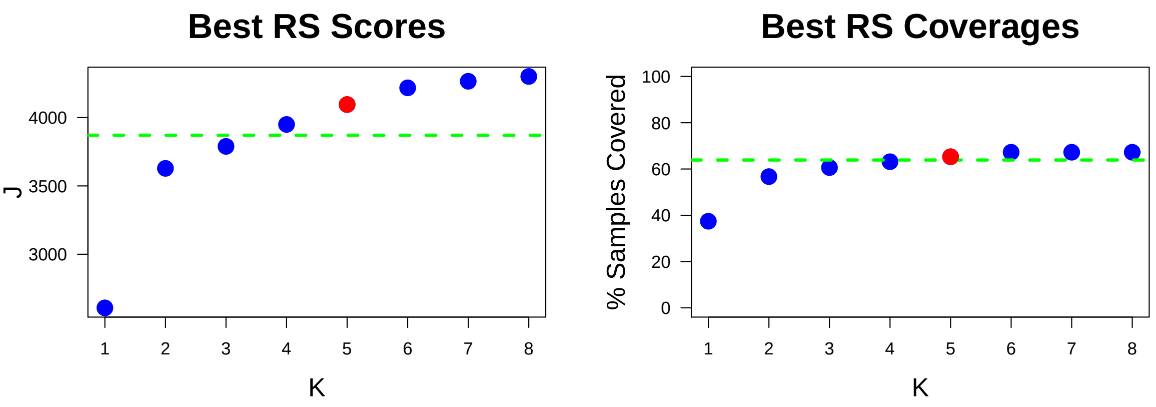

Figure 2: Performance and Coverage of Best Filtered RS for K. The core was determined to be the smallest RS that achieved at least 95% of maximum coverage and at least 90% of maximum performance.

#### 2.2 Table of rules that appear in any best rule set

| ID | Rule | PC | Ks |
|---|---|---|---|
| r3 | ATRX-MD + IDH1-M + TP53-M | 37 | 1,2,3,4,5,6,7,8 |
| r1 | CIC-M + IDH1-M | 19 | 2,3,4,5,6,7,8 |
| r9 | EGFR-M + SNORA73-A | 3.9 | 3,4,5,6,7,8 |
| r7 | IDH1-M + PIK3CA-M | 6 | 4,5,6,7,8 |
| r13 | IDH1-M + ISOC2/NAT14/ZNF628-D + TP53-M | 10 | 5,6,7,8 |
| r6 | FUBP1-M + IDH1-M | 8.8 | 6,7,8 |
| r42 | a8(n=7) + HUWE1/PHF8/RNA5SP505-A + IDH1-M + TP53-M | 4.1 | 7,8 |
| r96 | CAPZA2/MET/RNA5SP239-A + IDH1-M + MIR29B1-A + TP53-M | 3.5 | 8 |

**ID** = Rule IDs, rules are numbered according to importance rank determined from phase 1
**PC** = Percent of samples covered **Ks** = Membership in best RS

### 3 Core Rule Set

**Core K = 5.**
**Core rule set coverage = 65.3%.**

#### 3.1 Table of core rule set rules

| ID | Rule | CR | SJR | SJ | NSC | NSA | PC | PA | FracA |
|---|---|---|---|---|---|---|---|---|---|
| r1 | CIC-M + IDH1-M | 5 | 5 | 1020 | 99 | 89 | 19 | 17.0 | 0.90 |
| r3 | ATRX-MD + IDH1-M + TP53-M | 4 | 1 | 2610 | 192 | 184 | 37 | 36.0 | 0.96 |
| r7 | IDH1-M + PIK3CA-M | 60 | 42.5 | 346 | 31 | 23 | 6 | 4.5 | 0.74 |
| r9 | EGFR-M + SNORA73-A | 143 | 152.5 | 161 | 20 | 20 | 3.9 | 3.9 | 1.00 |
| r13 | IDH1-M + ISOC2/NAT14/ZNF628-D + TP53-M | 15 | 9 | 674 | 53 | 19 | 10 | 3.7 | 0.36 |

**ID** = Rule IDs, rules are numbered according to importance rank determined from phase 1
**CR** = Coverage rank
**SJ/SJR** = Single rule performance / performance rank
**NSC/PC** = Number/percent of samples covered
**NSA/PA** = Number/percent of samples assigned
**FracA** = Fraction of covered samples assigned to rule

#### 3.2 Event breakdown of core rule set

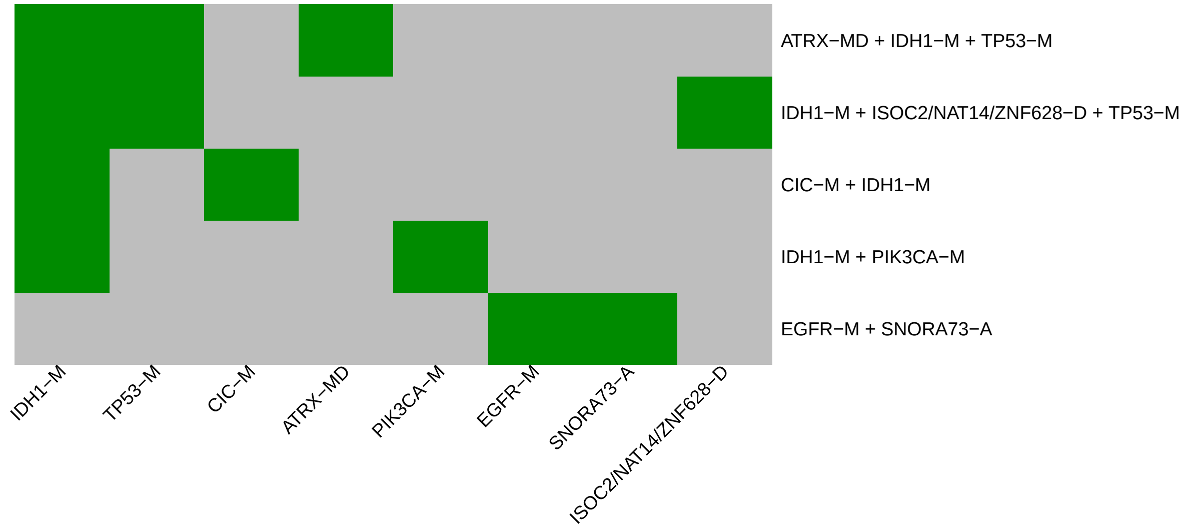

Figure 3: Visualization of core rule set. Rows are rules, columns are events. Events are ordered from left in decreasing rule membership frequency.

##### 3.3 Core Rule Set Penalties: Before and After Assignment

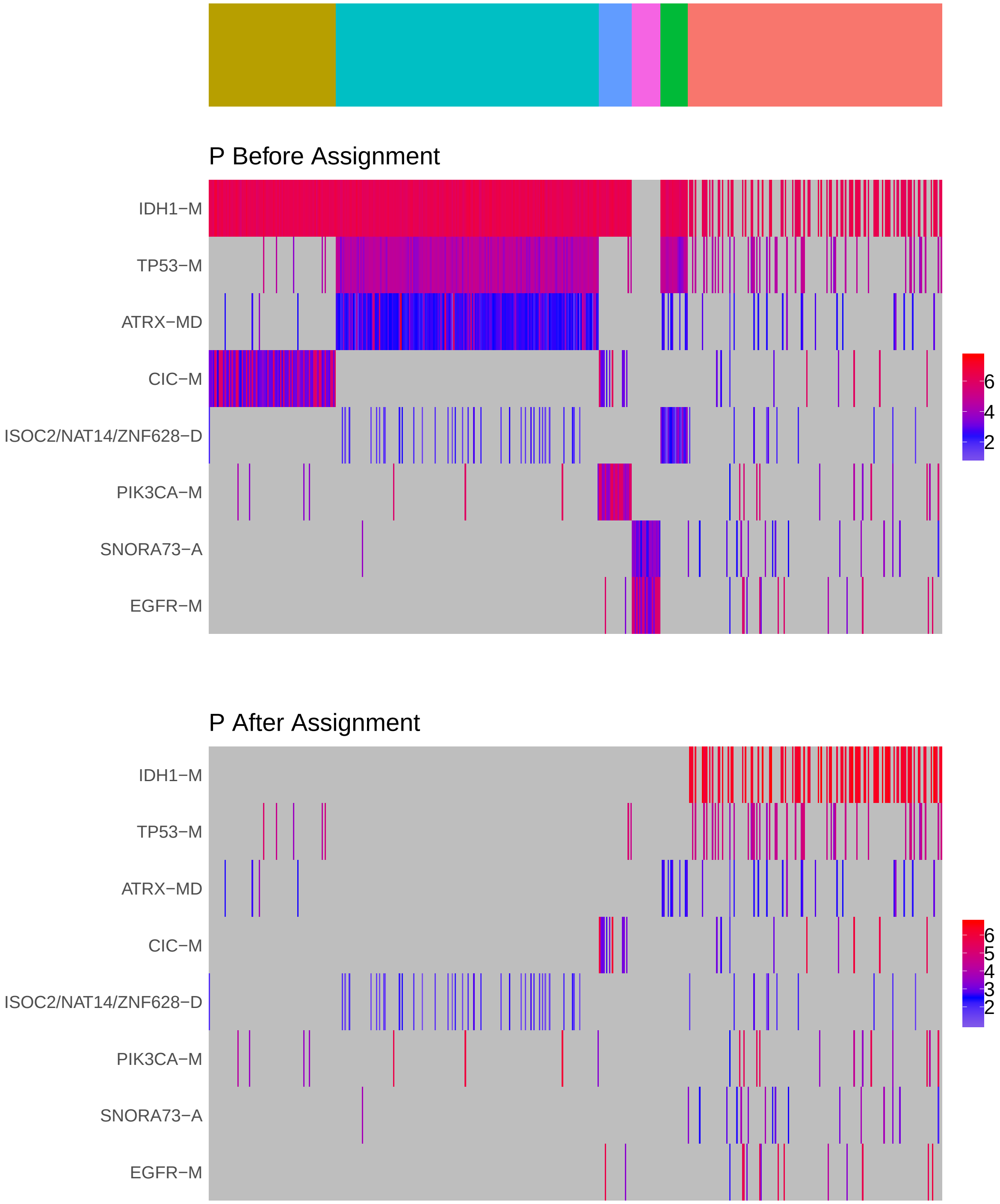

Figure 4: Heatmap of P before and after assignment to core rule set. Events are force ordered by frequency. Samples are force ordered according rule set membership, as indicated by the color bar. The right-most group of samples are not assigned to any rule.

### 4 Generalized Core Rules

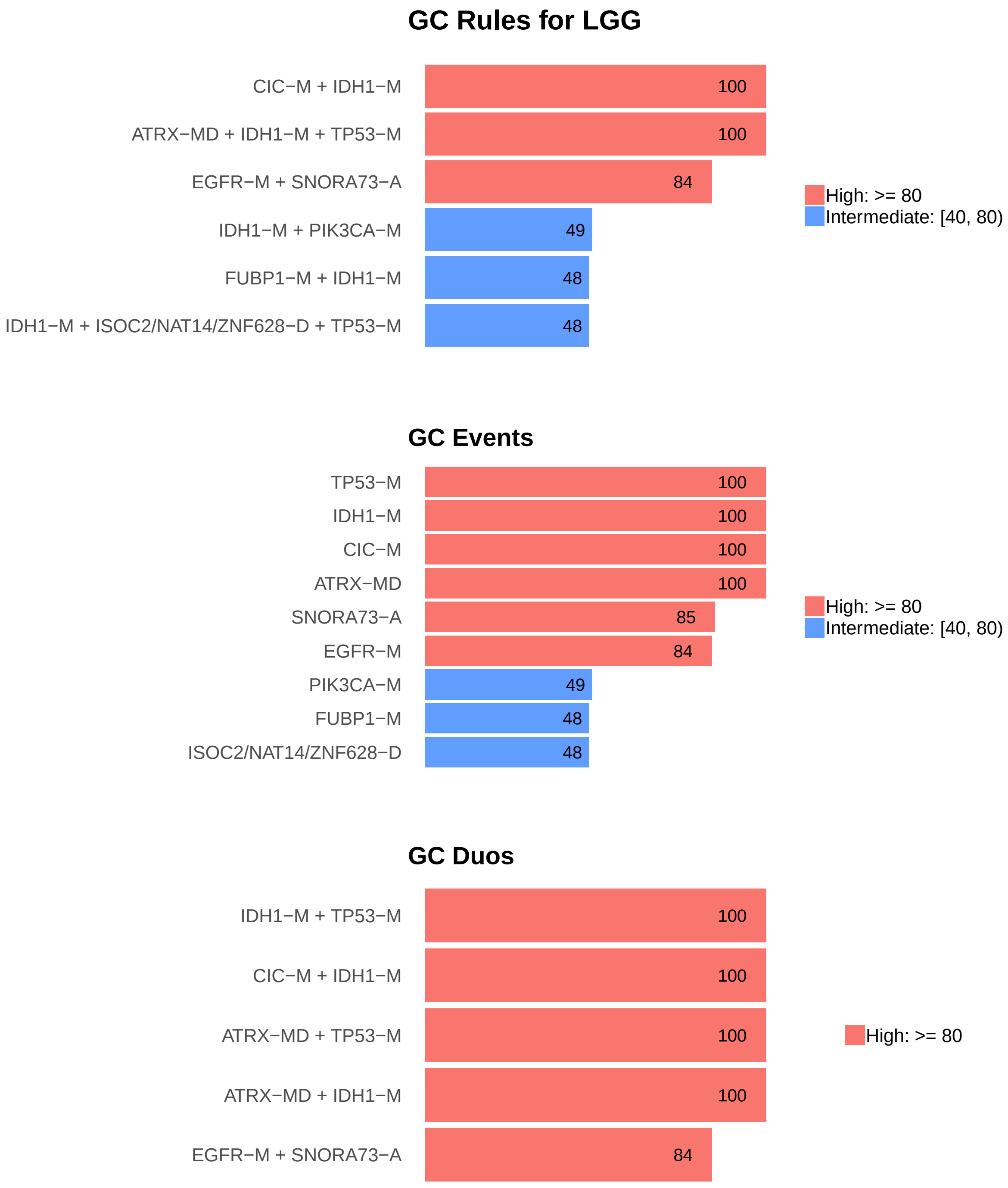

Figure 5: Summary of generalized core results. Bars show confidence levels, which are the percentage of sub-sample iterations containing the observation. Rules (top) and events (middle) that achieve a minimum confidence level of 20 are shown. Duos (bottom) with confidence of at least 50 are shown.

### 5 Dictionary of Copy Number Events

| CNV | Genes | Event_Name |
|---|---|---|
| a1 | snoU13\|ENSG00000238901.1 | snoU13-A |
| a2 | SNORA73\|ENSG00000252054.1 | SNORA73-A |
| a3 | MIR29B1 | MIR29B1-A |
| a4 | PARP11 | PARP11-A |
| a6 | RNA5SP239, CAPZA2, MET | CAPZA2/MET/RNA5SP239-A |
| a7 | RNA5SP505, HUWE1, PHF8 | HUWE1/PHF8/RNA5SP505-A |
| a8 | GNL3L, U3\|ENSG00000252175.1, FGD1, PHF8, FAM120C, WNK3, TSR2 | a8(n=7) |
| d1 | CDKN2A, C9orf53 | C9orf53/CDKN2A-D |
| d2 | C11orf58 | C11orf58-D |
| d3 | NAT14, ISOC2, ZNF628 | ISOC2/NAT14/ZNF628-D |
| d4 | CXXC11, RNA5SP122, BOK, DTYMK, PDCD1, ATG4B, THAP4, GAL3ST2, ING5, NEU4, D2HGDH | d4(n=11) |
| d5 | RN7SL5P, SNORD27\|ENSG00000251699.1, PTPRD | PTPRD/RN7SL5P/SNORD27-D |
| d6 | UTF1, VENTX, MIR202 | MIR202/UTF1/VENTX-D |
| d7 | KCNQ1OT1, KCNQ1 | KCNQ1/KCNQ1OT1-D |
| d8 | SNORA7\|ENSG00000222604.1, ISCA2, MIR4709, LTBP2, AREL1, NPC2 | d8(n=6) |
| d9 | LINC00290 | LINC00290-D |

Dictionary of important CNVs. Copy number events that are within any of the best K rule sets, or within the generalized core rules, or among the 20 most frequent events.
