## Supplementary material for "Identifying Combinations of Cancer Drivers in Individual Patients": TCGA Reports: CRSO_Report_LIHC.pdf

### CRSO Output Report: LIHC

Michael Klein

Apr 19 2019

**4 Generalized Core Rules** **6**

**5 Dictionary of Copy Number Events** **7**

### 1 Heatmaps of D and P

**Number of samples = 366.**
**Number of events = 75.**
**Rule coverage requirement = 11 samples.**
**Rule library size = 146 rules.**

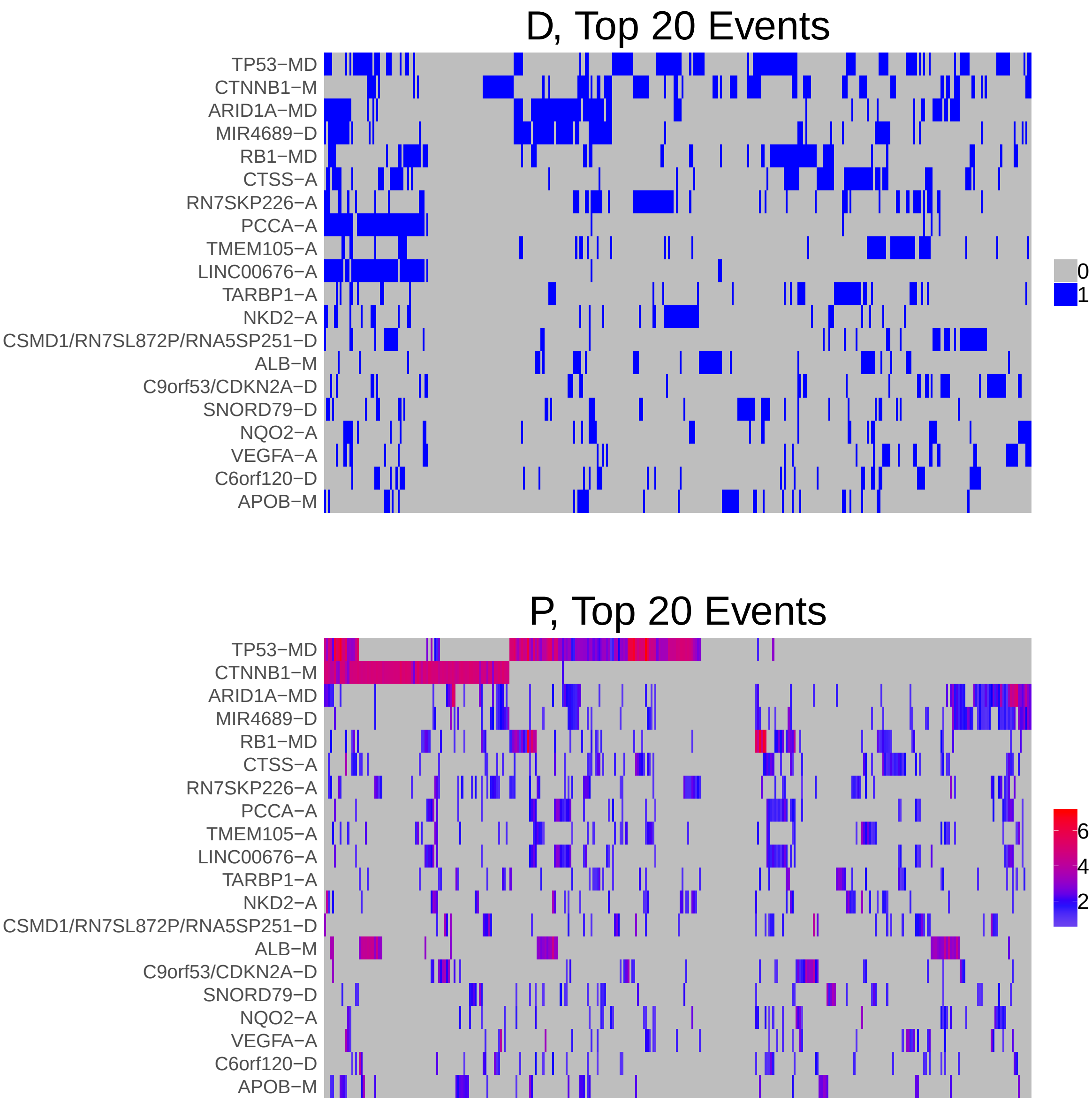

Figure 1: Heatmap of D and P. Events are ordered by frequency, from top to bottom. Event suffix -M: mutation, -A: amplification, -D: deletion, -MD: mutDel, -MA: mutAmp. Samples are ordered using hierarchical clustering. Wild-type events indicated in grey.

#### 2 Summary of K Best Rule Sets

##### 2.1 Performance and coverage of best rule sets

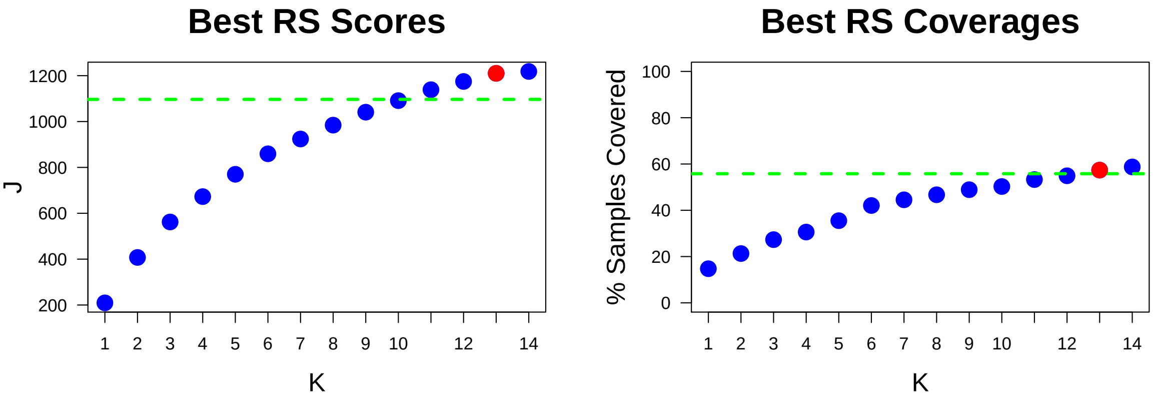

Figure 2: Performance and Coverage of Best Filtered RS for K. The core was determined to be the smallest RS that achieved at least 95% of maximum coverage and at least 90% of maximum performance.

##### 2.2 Table of rules that appear in any best rule set

| **ID** | **Rule** | **PC** | **Ks** |
|---|---|---|---|
| r1 | ARID1A-MD + MIR4689-D | 15 | 1,2,3,4,5,6,7,8,9,10,11,12,13,14 |
| r2 | RB1-MD + TP53-MD | 8.5 | 3,4,5,6,7,8,9,10,11,12,13,14 |
| r5 | CTNNB1-M + TP53-MD | 6.6 | 2,3,4,5,6,7,8,9,10,11,12,13 |
| r3 | ALB-M + CTNNB1-M | 4.4 | 4,5,6,7,8,9,10,11,12,13,14 |
| r4 | LINC00676-A + PCCA-A | 13 | 6,7,8,9,10,11,12,13,14 |
| r7 | CTSS-A + TP53-MD | 7.9 | 7,8,9,10,11,12,13,14 |
| r12 | RN7SKP226-A + TP53-MD | 8.7 | 5,6,9,10,11,12,13,14 |
| r8 | CTNNB1-M + RN7SKP226-A | 6.3 | 8,9,10,11,12,13,14 |
| r9 | ARID1A-MD + CTNNB1-M | 6.3 | 10,11,12,13,14 |
| r10 | C19orf77/NFIC-D + TP53-MD | 6.8 | 7,8,12,13,14 |
| r13 | CTSS-A + RB1-MD | 6.8 | 11,12,13,14 |
| r15 | ALB-M + TP53-MD | 3.6 | 9,10,11,14 |
| r11 | CCND1/ORAOV1-A + TP53-MD | 4.6 | 12,13,14 |
| r26 | RN7SKP96-D + TACR3-D | 4.6 | 13,14 |
| r30 | ARID1A-MD + TP53-MD | 6.6 | 14 |

**ID** = Rule IDs, rules are numbered according to importance rank determined from phase 1
**PC** = Percent of samples covered **Ks** = Membership in best RS

### 3 Core Rule Set

**Core K = 13.**
**Core rule set coverage = 57.4%.**

#### 3.1 Table of core rule set rules

| ID | Rule | CR | SJR | SJ | NSC | NSA | PC | PA | FracA |
|---|---|---|---|---|---|---|---|---|---|
| r1 | ARID1A-MD + MIR4689-D | 1 | 1 | 209 | 54 | 37 | 15 | 10.0 | 0.69 |
| r2 | RB1-MD + TP53-MD | 4 | 2 | 201 | 31 | 19 | 8.5 | 5.2 | 0.61 |
| r3 | ALB-M + CTNNB1-M | 45.5 | 12 | 134 | 16 | 14 | 4.4 | 3.8 | 0.88 |
| r4 | LINC00676-A + PCCA-A | 2 | 5 | 176 | 49 | 22 | 13 | 6.0 | 0.45 |
| r5 | CTNNB1-M + TP53-MD | 11.5 | 3 | 198 | 24 | 21 | 6.6 | 5.7 | 0.88 |
| r7 | CTSS-A + TP53-MD | 5 | 7 | 162 | 29 | 15 | 7.9 | 4.1 | 0.52 |
| r8 | CTNNB1-M + RN7SKP226-A | 14.5 | 9 | 143 | 23 | 11 | 6.3 | 3.0 | 0.48 |
| r9 | ARID1A-MD + CTNNB1-M | 14.5 | 8 | 152 | 23 | 13 | 6.3 | 3.6 | 0.57 |
| r10 | C19orf77/NFIC-D + TP53-MD | 8 | 10 | 139 | 25 | 11 | 6.8 | 3.0 | 0.44 |
| r11 | CCND1/ORAOV1-A + TP53-MD | 35 | 17 | 113 | 17 | 11 | 4.6 | 3.0 | 0.65 |
| r12 | RN7SKP226-A + TP53-MD | 3 | 4 | 177 | 32 | 13 | 8.7 | 3.6 | 0.41 |
| r13 | CTSS-A + RB1-MD | 8 | 23.5 | 100 | 25 | 12 | 6.8 | 3.3 | 0.48 |
| r26 | RN7SKP96-D + TACR3-D | 35 | 77 | 58.1 | 17 | 11 | 4.6 | 3.0 | 0.65 |

**ID** = Rule IDs, rules are numbered according to importance rank determined from phase 1
**CR** = Coverage rank
**SJ/SJR** = Single rule performance / performance rank
**NSC/PC** = Number/percent of samples covered
**NSA/PA** = Number/percent of samples assigned
**FracA** = Fraction of covered samples assigned to rule

#### 3.2 Event breakdown of core rule set

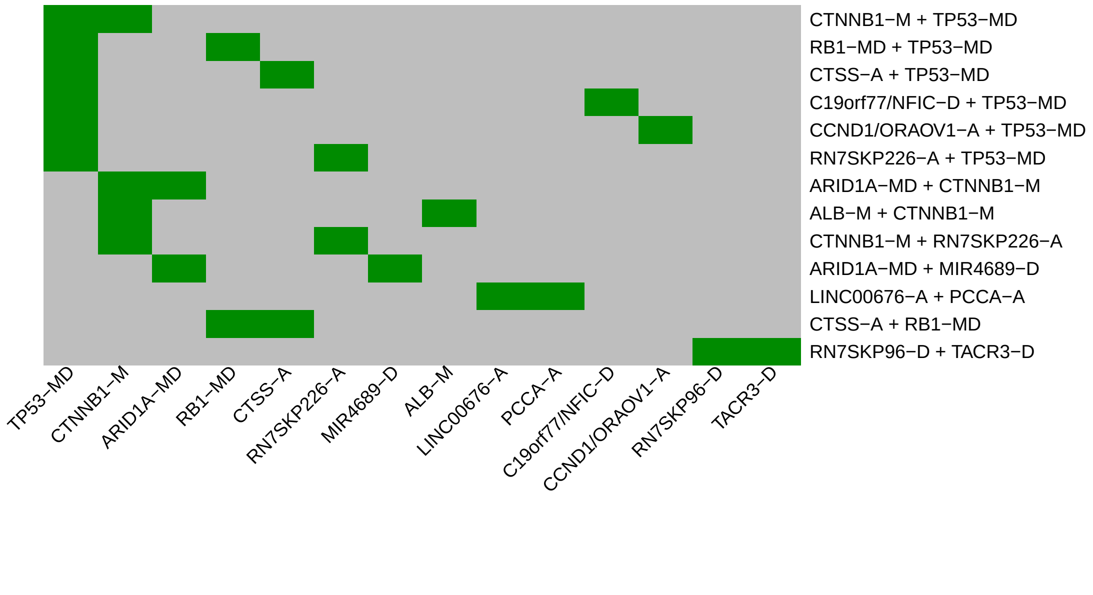

Figure 3: Visualization of core rule set. Rows are rules, columns are events. Events are ordered from left in decreasing rule membership frequency.

#### 3.3 Core Rule Set Penalties: Before and After Assignment

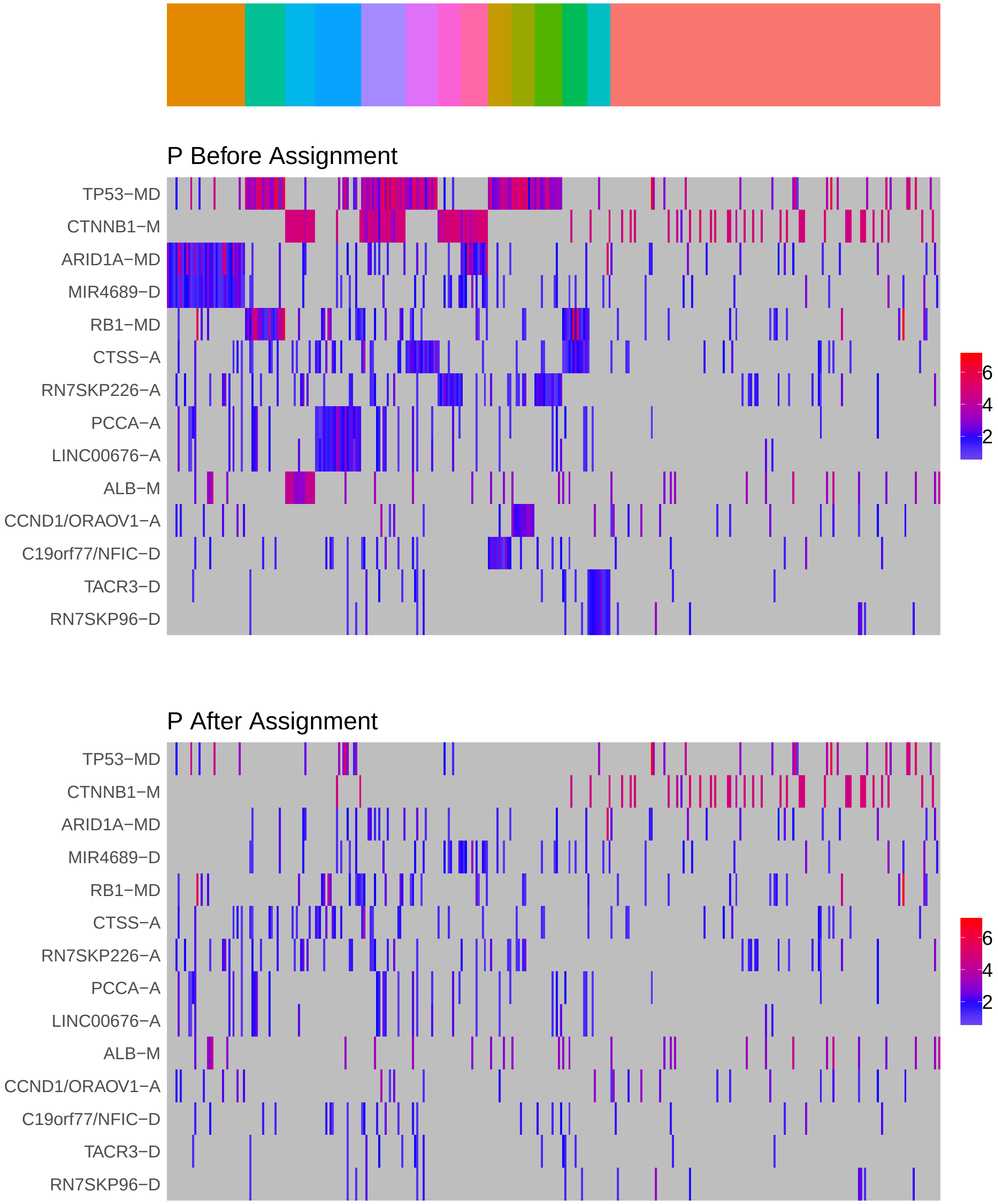

Figure 4: Heatmap of P before and after assignment to core rule set. Events are force ordered by frequency. Samples are force ordered according rule set membership, as indicated by the color bar. The right-most group of samples are not assigned to any rule.

### 4 Generalized Core Rules

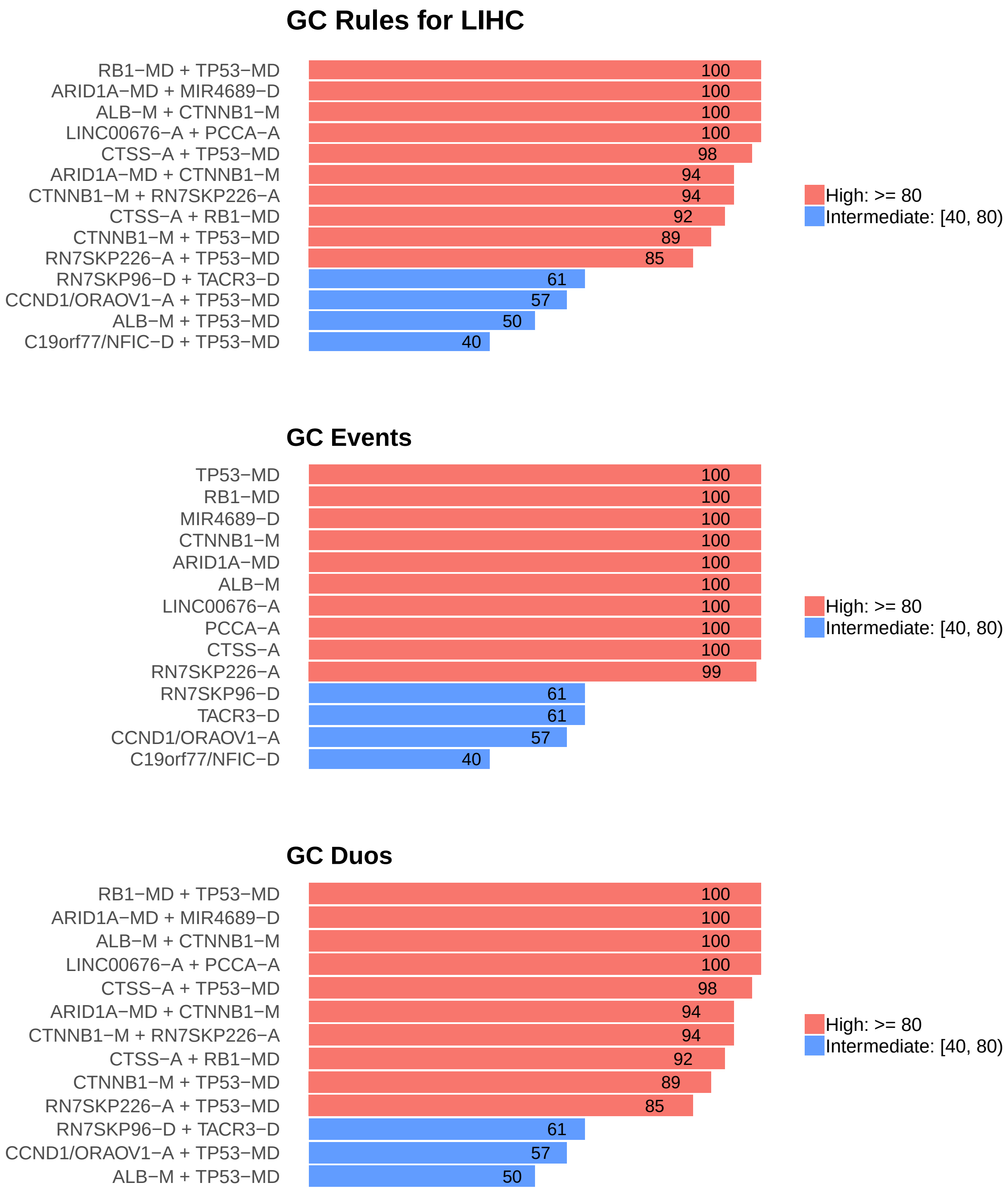

Figure 5: Summary of generalized core results. Bars show confidence levels, which are the percentage of sub-sample iterations containing the observation. Rules (top) and events (middle) that achieve a minimum confidence level of 20 are shown. Duos (bottom) with confidence of at least 50 are shown.

### 5 Dictionary of Copy Number Events

| CNV | Genes | Event__Name |
|---|---|---|
| a1 | CTSS | CTSS-A |
| a2 | RN7SKP226 | RN7SKP226-A |
| a3 | PCCA | PCCA-A |
| a4 | TMEM105 | TMEM105-A |
| a5 | LINC00676 | LINC00676-A |
| a6 | TARBP1 | TARBP1-A |
| a7 | NKD2 | NKD2-A |
| a8 | NQO2 | NQO2-A |
| a9 | VEGFA | VEGFA-A |
| a12 | CCND1, ORAOV1 | CCND1/ORAOV1-A |
| d1 | MIR4689 | MIR4689-D |
| d4 | RN7SL872P, RNA5SP251, CSMD1 | CSMD1/RN7SL872P/RNA5SP251-D |
| d5 | CDKN2A, C9orf53 | C9orf53/CDKN2A-D |
| d6 | SNORD79 | SNORD79-D |
| d7 | C6orf120 | C6orf120-D |
| d9 | NFIC, C19orf77 | C19orf77/NFIC-D |
| d16 | TACR3 | TACR3-D |
| d19 | RN7SKP96 | RN7SKP96-D |

Dictionary of important CNVs. Copy number events that are within any of the best K rule sets, or within the generalized core rules, or among the 20 most frequent events.
