## Supplementary material for "Identifying Combinations of Cancer Drivers in Individual Patients": TCGA Reports: CRSO_Report_LUAD.pdf

### CRSO Output Report: LUAD

Michael Klein

Apr 19 2019

**4 Generalized Core Rules** **6**

**5 Dictionary of Copy Number Events** **7**

#### 1 Heatmaps of D and P

**Number of samples = 478.**
**Number of events = 93.**
**Rule coverage requirement = 15 samples.**
**Rule library size = 291 rules.**

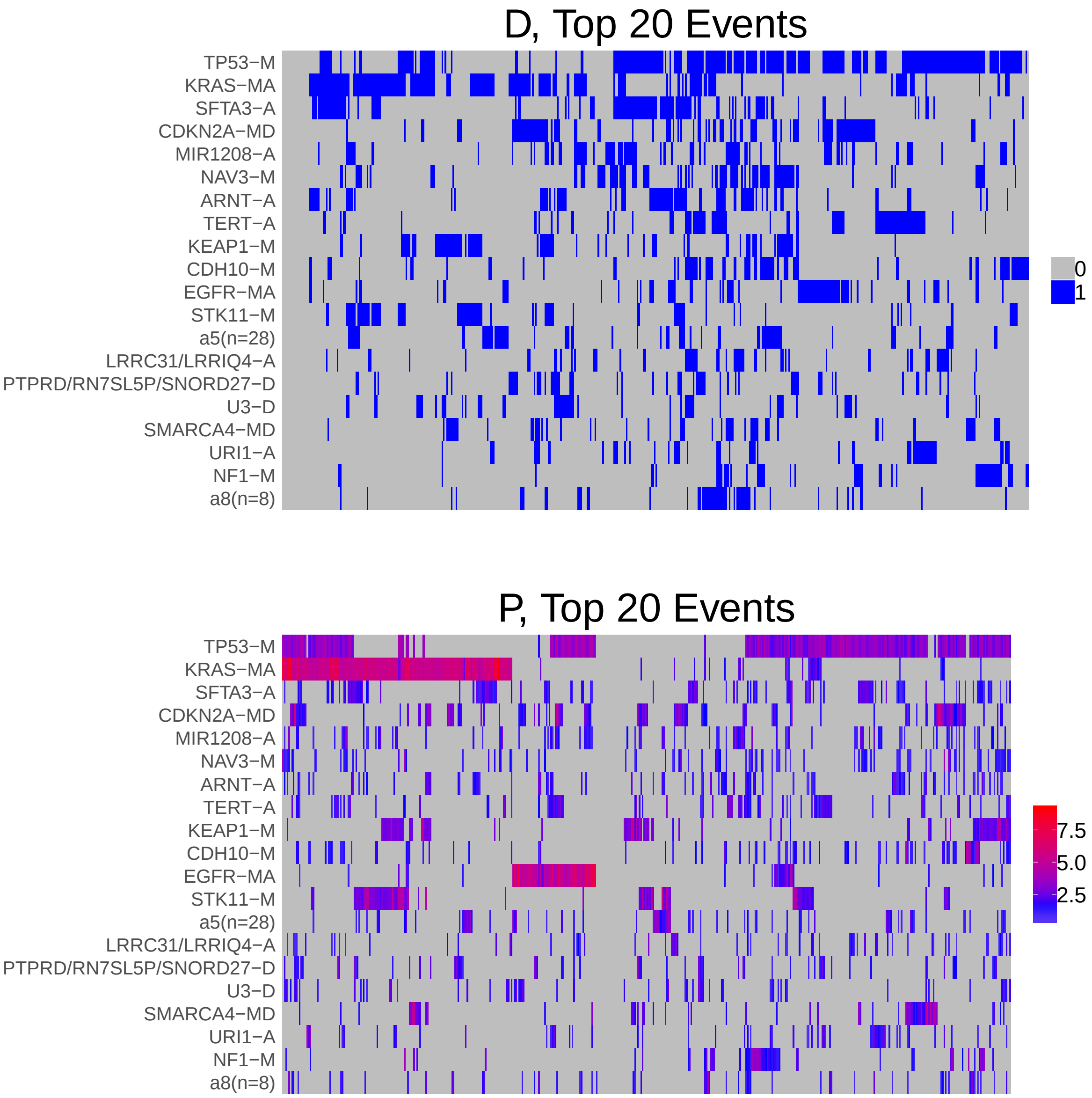

Figure 1: Heatmap of D and P. Events are ordered by frequency, from top to bottom. Event suffix -M: mutation, -A: amplification, -D: deletion, -MD: mutDel, -MA: mutAmp. Samples are ordered using hierarchical clustering. Wild-type events indicated in grey.

#### 2 Summary of K Best Rule Sets

##### 2.1 Performance and coverage of best rule sets

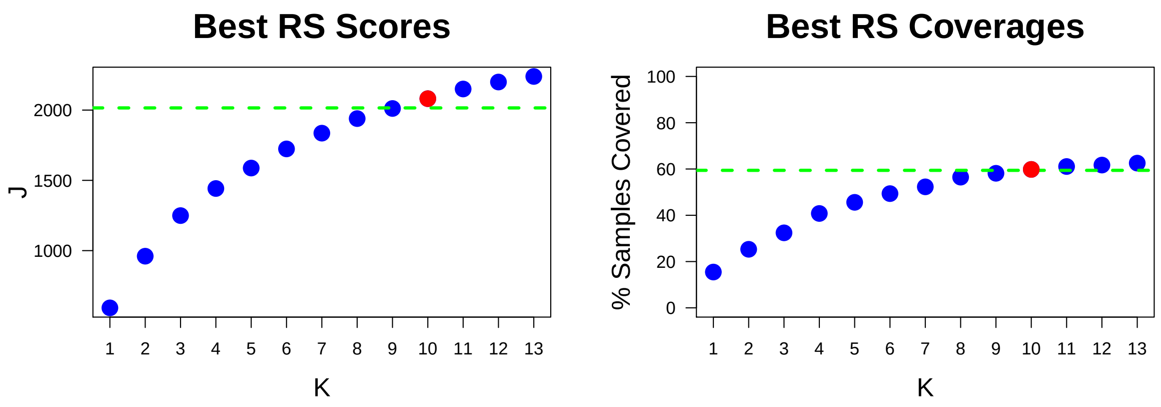

Figure 2: Performance and Coverage of Best Filtered RS for K. The core was determined to be the smallest RS that achieved at least 95% of maximum coverage and at least 90% of maximum performance.

##### 2.2 Table of rules that appear in any best rule set

| ID | Rule | PC | Ks |
|---|---|---|---|
| r1 | KRAS-MA + TP53-M | 15 | 1,2,3,4,5,6,7,8,9,10,11,12,13 |
| r2 | EGFR-MA + TP53-M | 12 | 2,3,4,5,6,7,8,9,10,11,12,13 |
| r3 | KRAS-MA + STK11-M | 10 | 3,4,5,6,7,8,9,10,11,12,13 |
| r6 | SFTA3-A + TP53-M | 15 | 4,5,6,7,8,9,10,11,12,13 |
| r8 | CDKN2A-MD + TP53-M | 13 | 5,6,7,8,9,10,11,12,13 |
| r4 | KRAS-MA + SFTA3-A | 11 | 6,7,8,9,10,11,12,13 |
| r5 | CDKN2A-MD + KRAS-MA | 8.2 | 7,8,9,10,11,12,13 |
| r7 | ATM-M + KRAS-MA | 6.1 | 10,11,12,13 |
| r10 | CDH10-M + TP53-M | 14 | 8,9,10 |
| r11 | NF1-M + TP53-M | 8.8 | 11,12,13 |
| r16 | KEAP1-M + TP53-M | 8.2 | 11,12,13 |
| r31 | a14(n=4) + TERT-A + TP53-M | 5.4 | 9,10,11 |
| r13 | BRAF-M + TP53-M | 5.2 | 12,13 |
| r18 | SMARCA4-MD + TP53-M | 6.7 | 12,13 |
| r30 | RB1-M + TP53-M | 4.8 | 13 |

**ID** = Rule IDs, rules are numbered according to importance rank determined from phase 1
**PC** = Percent of samples covered **Ks** = Membership in best RS

### 3 Core Rule Set

**Core K = 10.**
**Core rule set coverage = 59.8%.**

#### 3.1 Table of core rule set rules

| ID | Rule | CR | SJR | SJ | NSC | NSA | PC | PA | FracA |
|---|---|---|---|---|---|---|---|---|---|
| r1 | KRAS-MA + TP53-M | 1.5 | 1 | 593 | 74 | 60 | 15 | 13.0 | 0.81 |
| r2 | EGFR-MA + TP53-M | 9 | 2 | 408 | 55 | 42 | 12 | 8.8 | 0.76 |
| r3 | KRAS-MA + STK11-M | 11 | 3 | 380 | 48 | 29 | 10 | 6.1 | 0.60 |
| r4 | KRAS-MA + SFTA3-A | 10 | 6 | 338 | 51 | 15 | 11 | 3.1 | 0.29 |
| r5 | CDKN2A-MD + KRAS-MA | 16.5 | 10 | 289 | 39 | 16 | 8.2 | 3.3 | 0.41 |
| r6 | SFTA3-A + TP53-M | 1.5 | 4.5 | 362 | 74 | 28 | 15 | 5.9 | 0.38 |
| r7 | ATM-M + KRAS-MA | 48 | 15.5 | 223 | 29 | 20 | 6.1 | 4.2 | 0.69 |
| r8 | CDKN2A-MD + TP53-M | 5 | 4.5 | 362 | 64 | 31 | 13 | 6.5 | 0.48 |
| r10 | CDH10-M + TP53-M | 4 | 8 | 305 | 66 | 28 | 14 | 5.9 | 0.42 |
| r31 | a14(n=4) + TERT-A + TP53-M | 62 | 37.5 | 174 | 26 | 17 | 5.4 | 3.6 | 0.65 |

**ID** = Rule IDs, rules are numbered according to importance rank determined from phase 1
**CR** = Coverage rank
**SJ/SJR** = Single rule performance / performance rank
**NSC/PC** = Number/percent of samples covered
**NSA/PA** = Number/percent of samples assigned
**FracA** = Fraction of covered samples assigned to rule

#### 3.2 Event breakdown of core rule set

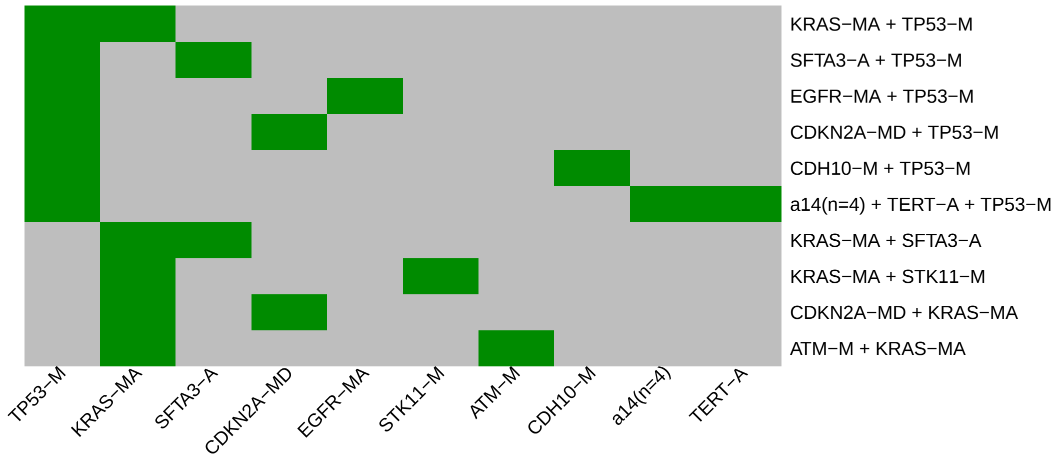

Figure 3: Visualization of core rule set. Rows are rules, columns are events. Events are ordered from left in decreasing rule membership frequency.

#### 3.3 Core Rule Set Penalties: Before and After Assignment

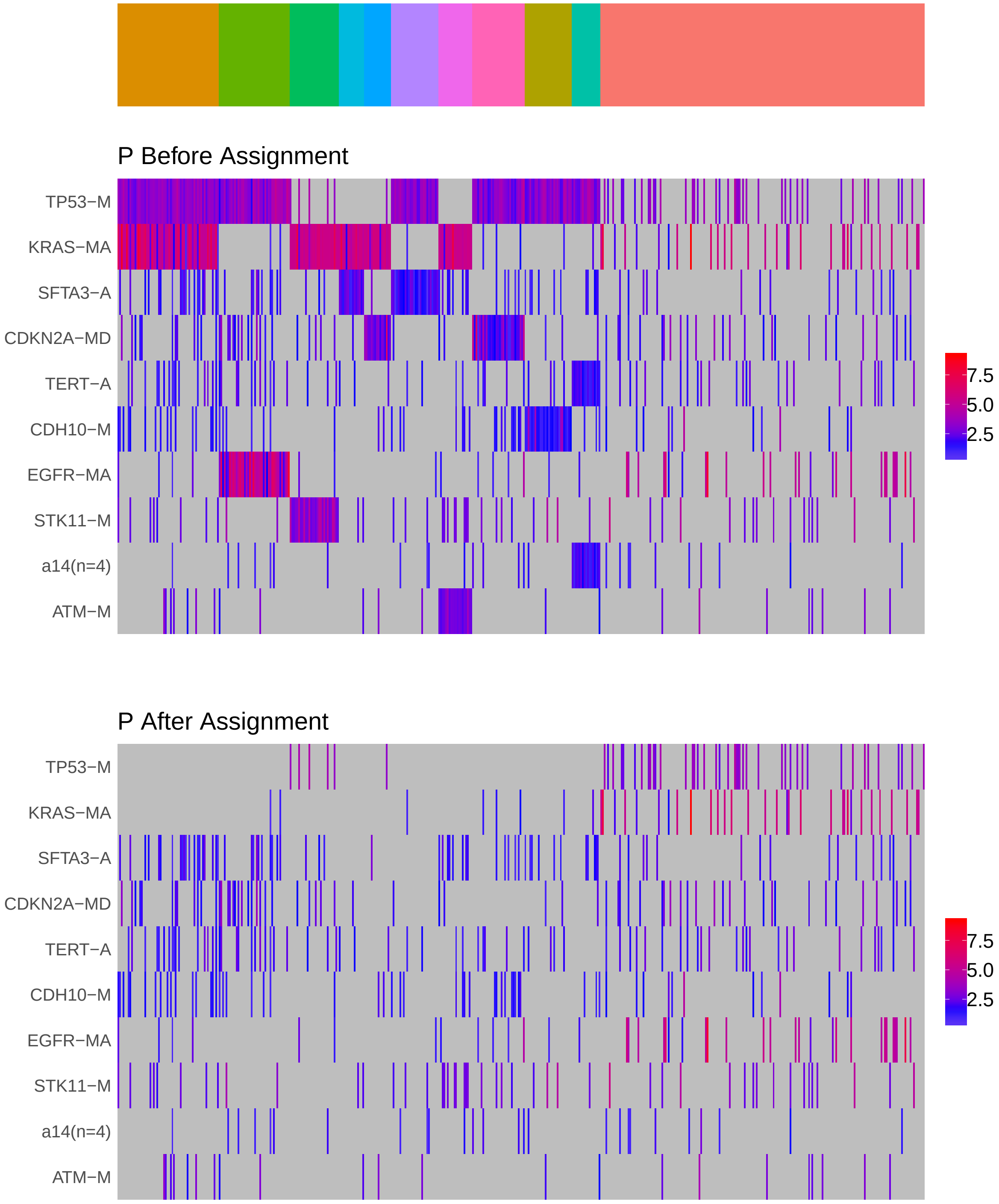

Figure 4: Heatmap of P before and after assignment to core rule set. Events are force ordered by frequency. Samples are force ordered according rule set membership, as indicated by the color bar. The right-most group of samples are not assigned to any rule.

### 4 Generalized Core Rules

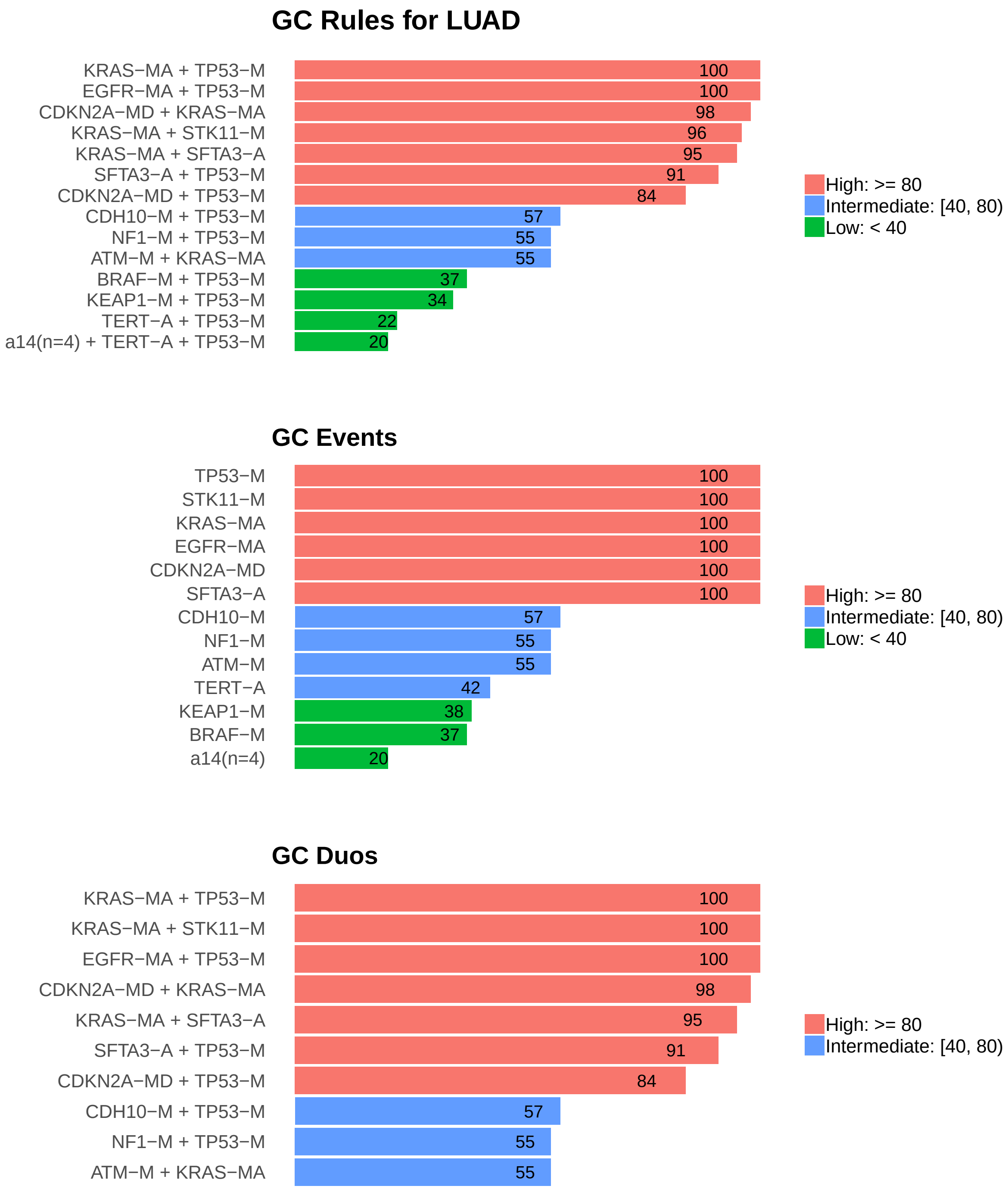

Figure 5: Summary of generalized core results. Bars show confidence levels, which are the percentage of sub-sample iterations containing the observation. Rules (top) and events (middle) that achieve a minimum confidence level of 20 are shown. Duos (bottom) with confidence of at least 50 are shown.

### 5 Dictionary of Copy Number Events

| CNV | Genes | Event_Name |
|---|---|---|
| a1 | SFTA3 | SFTA3-A |
| a2 | MIR1208 | MIR1208-A |
| a3 | ARNT | ARNT-A |
| a4 | TERT | TERT-A |
| a5 | a5(n=28) | a5(n=28) |
| a6 | LRRC31, LRRIQ4 | LRRC31/LRRIQ4-A |
| a7 | URI1 | URI1-A |
| a8 | RN7SL742P, RN7SL697P, LAGE3, G6PD, UBL4A, SLC10A3, PLXNA3, FAM3A | a8(n=8) |
| a14 | SNORA57\|ENSG00000212567.1, PRKAA1, PTGER4, TTC33 | a14(n=4) |
| d2 | RN7SL5P, SNORD27\|ENSG00000251699.1, PTPRD | PTPRD/RN7SL5P/SNORD27-D |
| d3 | U3\|ENSG00000221040.1 | U3-D |

Dictionary of important CNVs. Copy number events that are within any of the best K rule sets, or within the generalized core rules, or among the 20 most frequent events.
