## Supplementary material for "Identifying Combinations of Cancer Drivers in Individual Patients": TCGA Reports: CRSO_Report_LUSC.pdf

### CRSO Output Report: LUSC

Michael Klein

Apr 19 2019

**4 Generalized Core Rules** **8**

**5 Dictionary of Copy Number Events** **9**

### 1 Heatmaps of D and P

**Number of samples = 178.**
**Number of events = 112.**
**Rule coverage requirement = 6 samples.**
**Rule library size = 828 rules.**

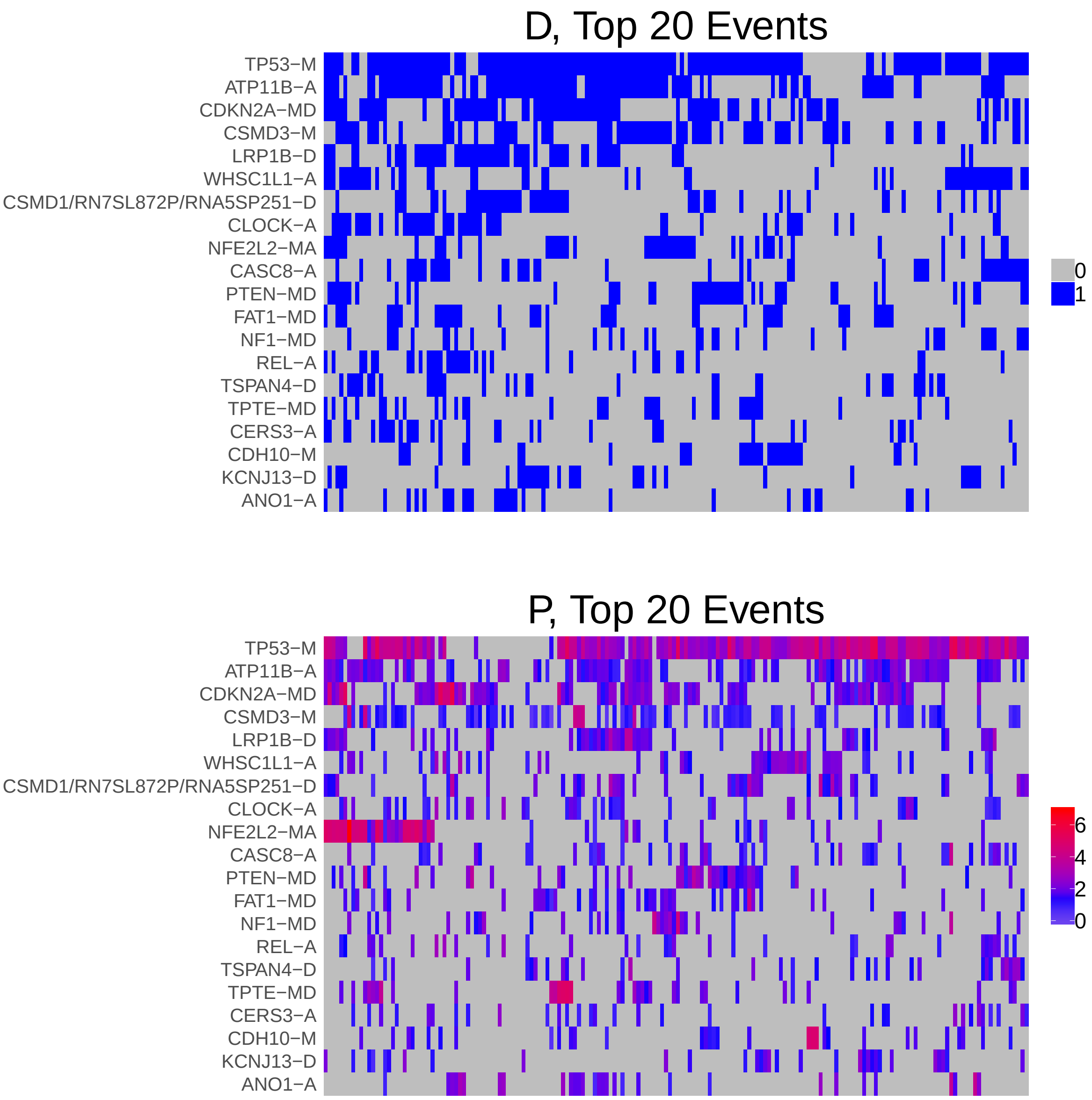

Figure 1: Heatmap of D and P. Events are ordered by frequency, from top to bottom. Event suffix -M: mutation, -A: amplification, -D: deletion, -MD: mutDel, -MA: mutAmp. Samples are ordered using hierarchical clustering. Wild-type events indicated in grey.

### 2 Summary of K Best Rule Sets

#### 2.1 Performance and coverage of best rule sets

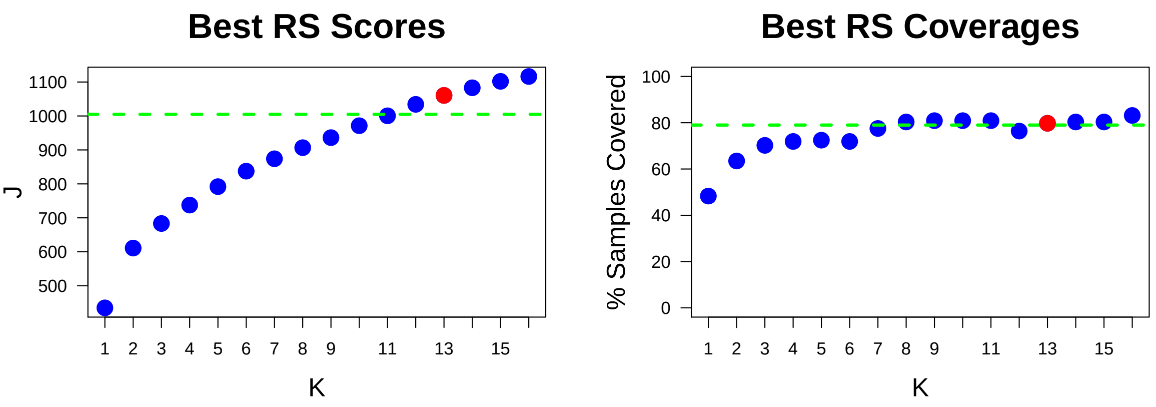

Figure 2: Performance and Coverage of Best Filtered RS for K. The core was determined to be the smallest RS that achieved at least 95% of maximum coverage and at least 90% of maximum performance.

#### 2.2 Table of rules that appear in any best rule set

| ID | Rule | PC | Ks |
|---|---|---|---|
| r3 | TP53-M + WHSC1L1-A | 21 | 3,4,5,6,7,8,9,10,11,12,13,14,15,16 |
| r11 | CDH10-M + TP53-M | 16 | 5,7,8,9,10,11,12,13,14,15,16 |
| r4 | FOXP1/MIR1284-D + MIR3923/RN7SL751P/ROBO1-D + PROS1/STX19-D + ROBO2-D + TP53-M | 6.7 | 6,7,8,10,11,12,13,14,15,16 |
| r25 | TP53-M + TSPAN4-D | 16 | 8,9,10,11,12,13,14,15,16 |
| r12 | NF1-MD + TP53-M | 16 | 9,10,11,12,13,14,15,16 |
| r19 | CDKN2A-MD + PTEN-MD + TP53-M | 12 | 9,10,11,12,13,14,15,16 |
| r24 | CDKN2A-MD + LRP1B-D + NFE2L2-MA + TP53-M | 6.2 | 9,10,11,12,13,14,15,16 |
| r1 | CDKN2A-MD + TP53-M | 39 | 2,3,4,5,6,7,8 |
| r17 | ATP11B-A + NFE2L2-MA + TP53-M | 15 | 6,7,8,9,10,11 |
| r8 | ATP11B-A + CSMD3-M | 28 | 7,8,9,10,11 |
| r9 | ATP11B-A + PIK3CA-M + TP53-M | 8.4 | 12,13,14,15,16 |
| r13 | ATP11B-A + RB1-MD + TP53-M | 8.4 | 12,13,14,15,16 |
| r20 | ATP11B-A + CDKN2A-MD + EGFR-A + LRP1B-D + TP53-M | 6.2 | 12,13,14,15,16 |
| r33 | ATP11B-A + CSMD3-M + NFE2L2-MA | 8.4 | 12,13,14,15,16 |
| r2 | ATP11B-A + TP53-M | 48 | 1,2,3,4 |
| r16 | ATP11B-A + LRP1B-D + TP53-M | 22 | 5,6,7,8 |
| r6 | ATP11B-A + CDKN2A-MD + TP53-M | 24 | 9,10,11 |
| r34 | CDKN2A-MD + TP53-M + TPTE-MD | 9 | 12,14,15 |
| r62 | CDKN2A-MD + CLOCK-A | 15 | 14,15 |
| r80 | ATP11B-A + CSMD1/RN7SL872P/RNA5SP251-D + CSMD3-M + TP53-M | 7.9 | 15,16 |
| r82 | ANO1-A + CDKN2A-MD | 10 | 13,16 |
| r85 | ATP11B-A + CASC8-A + LRP1B-D + TP53-M | 7.9 | 14,15 |
| r5 | CSMD3-M + TP53-M | 36 | 6 |
| r18 | NFE2L2-MA + TP53-M | 22 | 4 |
| r21 | ATP11B-A + CSMD3-M + TP53-M | 22 | 5 |
| r42 | ATP11B-A + CSMD1/RN7SL872P/RNA5SP251-D + LRP1B-D + TP53-M | 11 | 11 |
| r48 | CDKN2A-MD + CSMD3-M | 22 | 16 |
| r75 | ATP11B-A + KEAP1-M + TP53-M | 7.9 | 16 |
| r111 | ATP11B-A + LRP1B-D + PLEKHO1/VPS45-A + TP53-M | 7.3 | 13 |
| r112 | ATP11B-A + KDM6A-MD + TP53-M | 7.3 | 16 |

**ID** = Rule IDs, rules are numbered according to importance rank determined from phase 1
**PC** = Percent of samples covered **Ks** = Membership in best RS

### 3 Core Rule Set

**Core K = 13.**
**Core rule set coverage = 79.8%.**

#### 3.1 Table of core rule set rules

| ID | Rule | CR | SJR | SJ | NSC | NSA | PC | PA | FracA |
|---|---|---|---|---|---|---|---|---|---|
| r3 | TP53-M + WHSC1L1-A | 16 | 17 | 193 | 38 | 17 | 21 | 9.6 | 0.45 |
| r4 | FOXP1/MIR1284-D + MIR3923/RN7SL751P/ROBO1-D + PROS1/STX19-D + ROBO2-D + TP53-M | 439.5 | 110.5 | 106 | 12 | 9 | 6.7 | 5.1 | 0.75 |
| r9 | ATP11B-A + PIK3CA-M + TP53-M | 230 | 63.5 | 125 | 15 | 13 | 8.4 | 7.3 | 0.87 |
| r11 | CDH10-M + TP53-M | 28 | 34.5 | 149 | 29 | 13 | 16 | 7.3 | 0.45 |
| r12 | NF1-MD + TP53-M | 32.5 | 38.5 | 144 | 28 | 11 | 16 | 6.2 | 0.39 |
| r13 | ATP11B-A + RB1-MD + TP53-M | 230 | 59.5 | 127 | 15 | 10 | 8.4 | 5.6 | 0.67 |
| r19 | CDKN2A-MD + PTEN-MD + TP53-M | 80 | 29 | 152 | 21 | 15 | 12 | 8.4 | 0.71 |
| r20 | ATP11B-A + CDKN2A-MD + EGFR-A + LRP1B-D + TP53-M | 547 | 84.5 | 112 | 11 | 11 | 6.2 | 6.2 | 1.00 |
| r24 | CDKN2A-MD + LRP1B-D + NFE2L2-MA + TP53-M | 547 | 68.5 | 120 | 11 | 10 | 6.2 | 5.6 | 0.91 |
| r25 | TP53-M + TSPAN4-D | 32.5 | 34.5 | 149 | 28 | 12 | 16 | 6.7 | 0.43 |
| r33 | ATP11B-A + CSMD3-M + NFE2L2-MA | 230 | 125 | 102 | 15 | 8 | 8.4 | 4.5 | 0.53 |
| r82 | ANO1-A + CDKN2A-MD | 128.5 | 283.5 | 75.7 | 18 | 6 | 10 | 3.4 | 0.33 |
| r111 | ATP11B-A + LRP1B-D + PLEKHO1/VPS45-A + TP53-M | 348 | 117 | 104 | 13 | 7 | 7.3 | 3.9 | 0.54 |

**ID** = Rule IDs, rules are numbered according to importance rank determined from phase 1
**CR** = Coverage rank
**SJ/SJR** = Single rule performance / performance rank
**NSC/PC** = Number/percent of samples covered
**NSA/PA** = Number/percent of samples assigned
**FracA** = Fraction of covered samples assigned to rule

#### 3.2 Event breakdown of core rule set

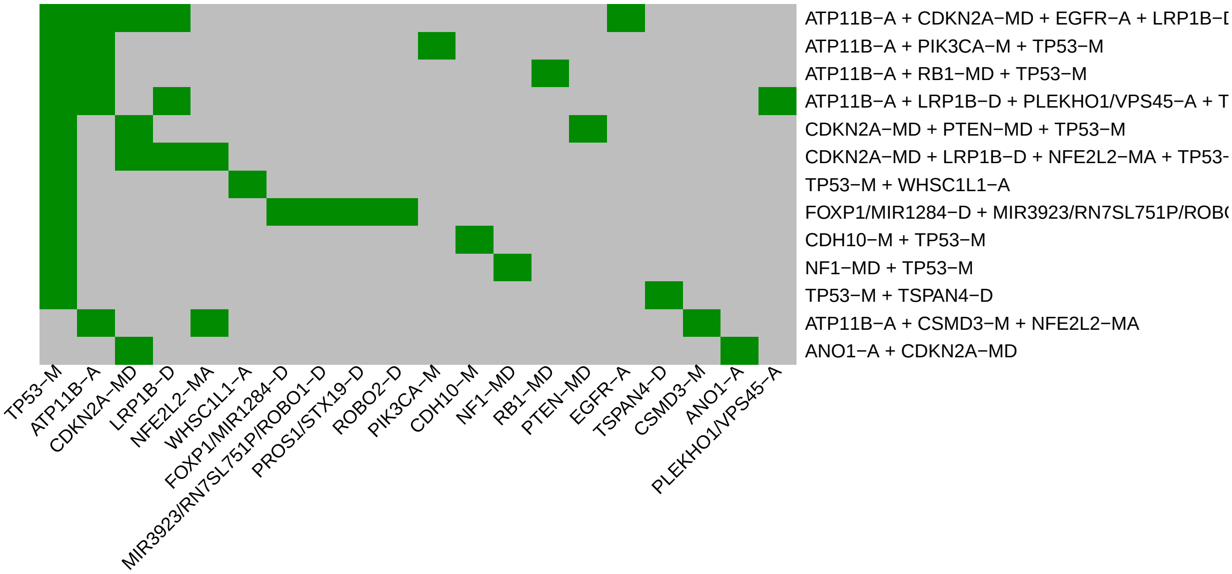

Figure 3: Visualization of core rule set. Rows are rules, columns are events. Events are ordered from left in decreasing rule membership frequency.

#### 3.3 Core Rule Set Penalties: Before and After Assignment

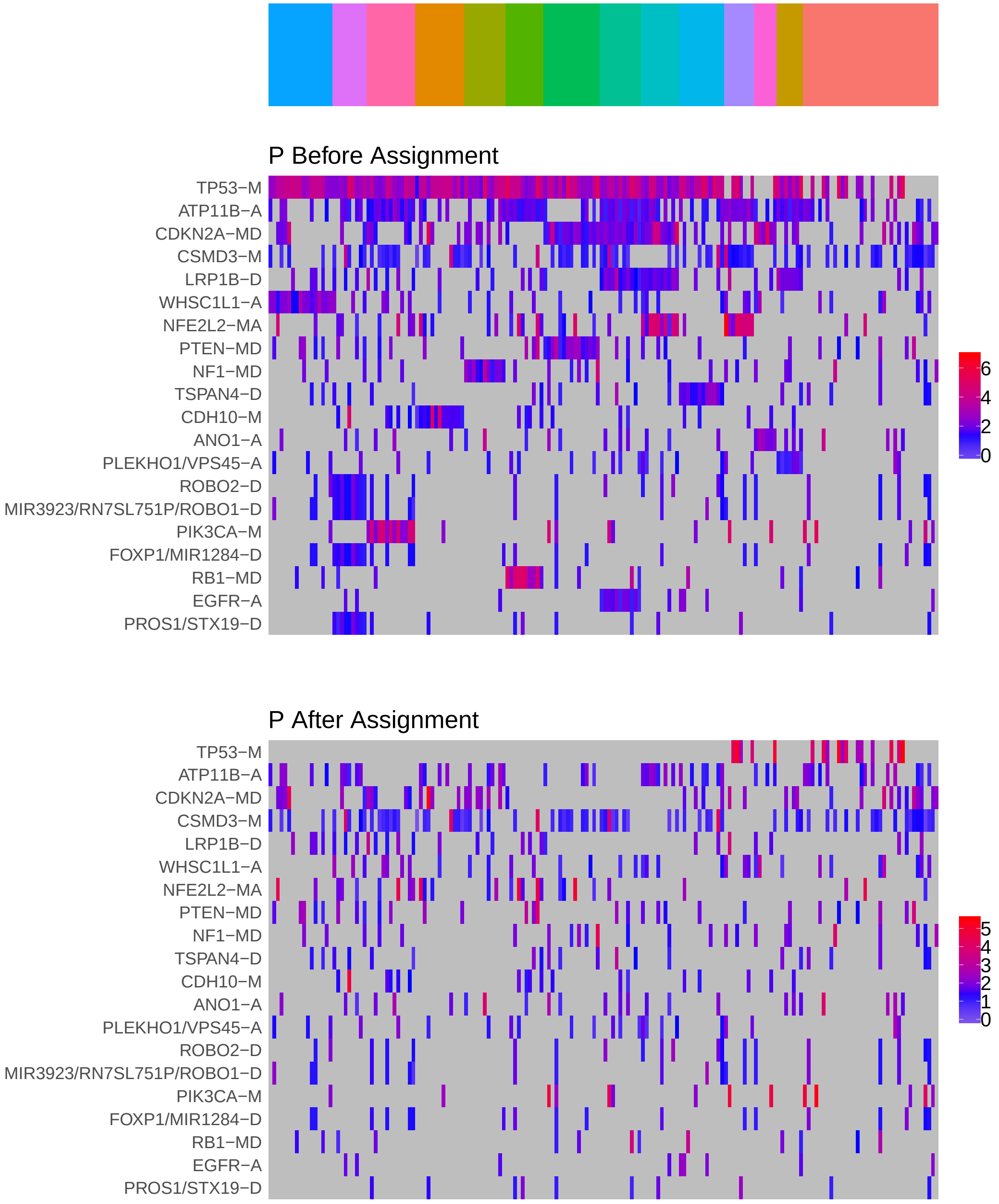

Figure 4: Heatmap of P before and after assignment to core rule set. Events are force ordered by frequency. Samples are force ordered according rule set membership, as indicated by the color bar. The right-most group of samples are not assigned to any rule.

### 4 Generalized Core Rules

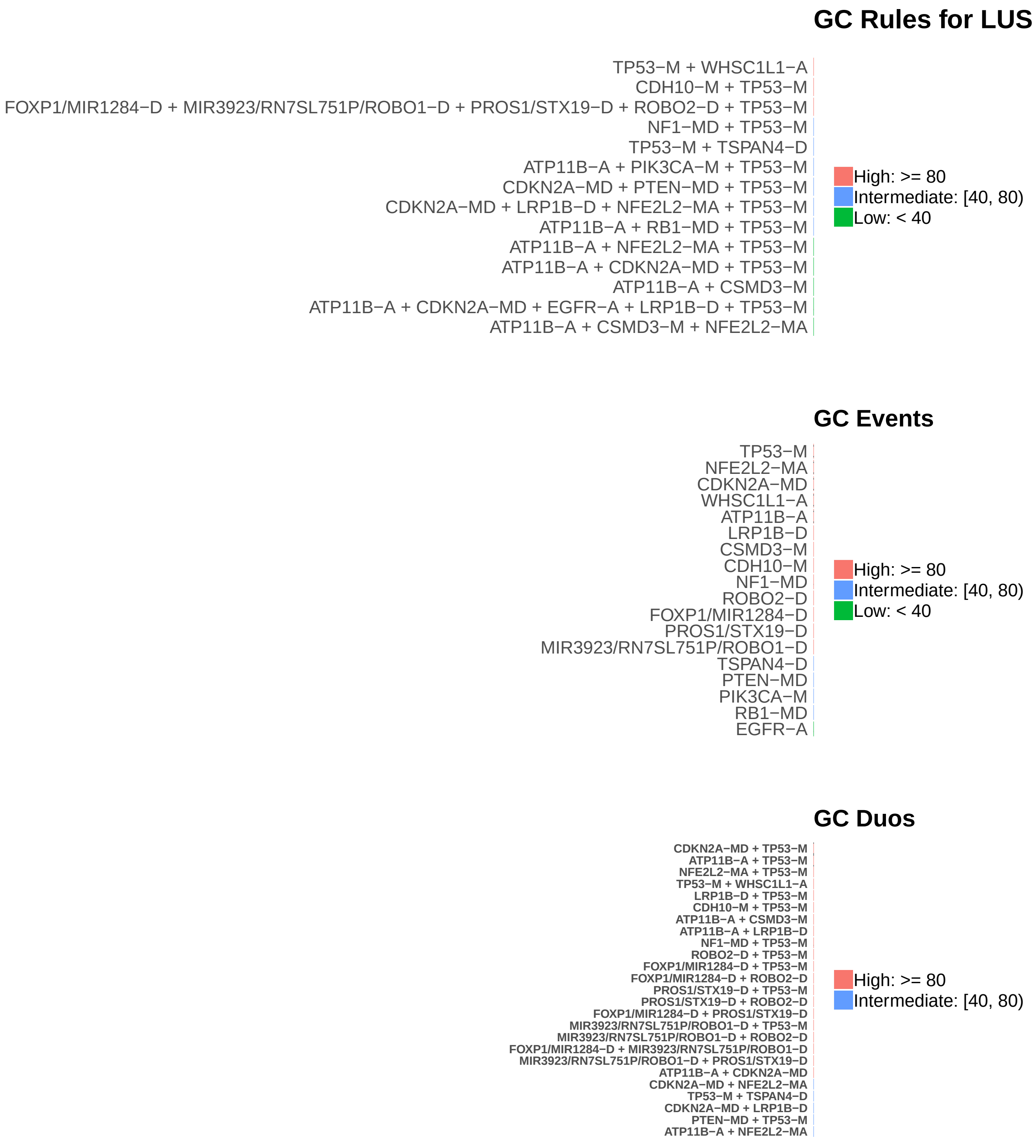

Figure 5: Summary of generalized core results. Bars show confidence levels, which are the percentage of sub-sample iterations containing the observation. Rules (top) and events (middle) that achieve a minimum confidence level of 30 are shown. Duos (bottom) with confidence of at least 70 are shown.

### 5 Dictionary of Copy Number Events

| CNV | Genes | Event_Name |
|---|---|---|
| a1 | ATP11B | ATP11B-A |
| a2 | CASC8 | CASC8-A |
| a3 | WHSC1L1 | WHSC1L1-A |
| a4 | CLOCK | CLOCK-A |
| a6 | ANO1 | ANO1-A |
| a7 | KCNK6, SPINT2, CATSPERG, C19orf33, YIF1B, PPP1R14A | a7(n=6) |
| a8 | UNC13B, ATP8B5P | ATP8B5P/UNC13B-A |
| a9 | CCNE1 | CCNE1-A |
| a11 | CERS3 | CERS3-A |
| a12 | REL | REL-A |
| a13 | VPS45, PLEKHO1 | PLEKHO1/VPS45-A |
| a14 | CDC42EP4 | CDC42EP4-A |
| a21 | EGFR | EGFR-A |
| d2 | LRP1B | LRP1B-D |
| d3 | RN7SL872P, RNA5SP251, CSMD1 | CSMD1/RN7SL872P/RNA5SP251-D |
| d4 | KCNJ13 | KCNJ13-D |
| d5 | TSPAN4 | TSPAN4-D |
| d6 | FOXP1, MIR1284 | FOXP1/MIR1284-D |
| d7 | ROBO2 | ROBO2-D |
| d10 | RN7SL751P, ROBO1, MIR3923 | MIR3923/RN7SL751P/ROBO1-D |
| d21 | PROS1, STX19 | PROS1/STX19-D |

Dictionary of important CNVs. Copy number events that are within any of the best K rule sets, or within the generalized core rules, or among the 20 most frequent events.
