## Supplementary material for "Identifying Combinations of Cancer Drivers in Individual Patients": TCGA Reports: CRSO_Report_OV.pdf

### CRSO Output Report: OV

Michael Klein

Apr 19 2019

**4 Generalized Core Rules** **7**

**5 Dictionary of Copy Number Events** **8**

#### 1 Heatmaps of D and P

**Number of samples = 455.**
**Number of events = 77.**
**Rule coverage requirement = 14 samples.**
**Rule library size = 932 rules.**

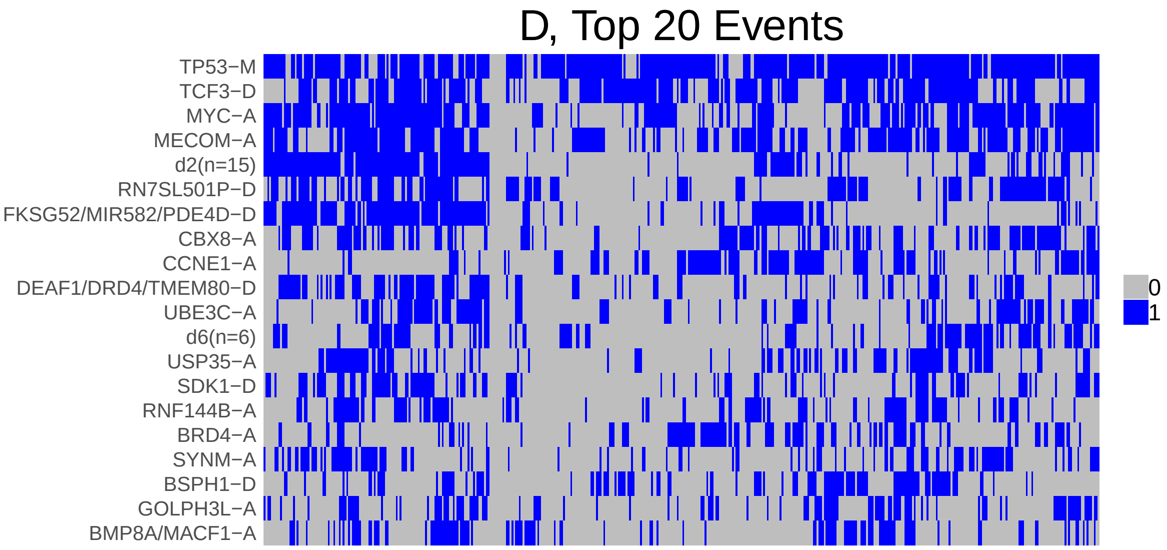

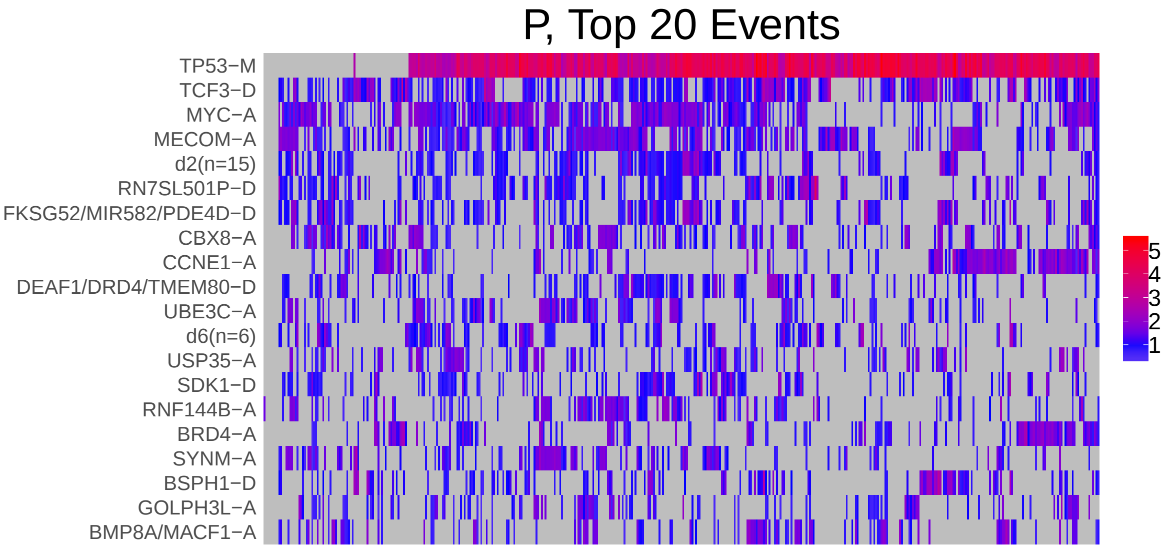

Figure 1: Heatmap of D and P. Events are ordered by frequency, from top to bottom. Event suffix -M: mutation, -A: amplification, -D: deletion, -MD: mutDel, -MA: mutAmp. Samples are ordered using hierarchical clustering. Wild-type events indicated in grey.

### 2 Summary of K Best Rule Sets

#### 2.1 Performance and coverage of best rule sets

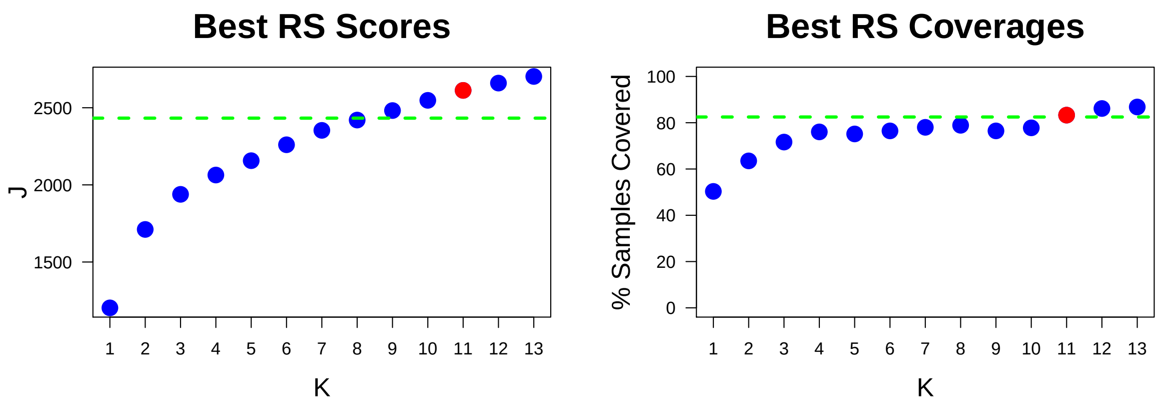

Figure 2: Performance and Coverage of Best Filtered RS for K. The core was determined to be the smallest RS that achieved at least 95% of maximum coverage and at least 90% of maximum performance.

#### 2.2 Table of rules that appear in any best rule set

| ID | Rule | PC | Ks |
|---|---|---|---|
| r3 | BSPH1-D + RN7SL526P-D + TCF3-D + TP53-M | 12 | 6,7,8,9,10,11,12,13 |
| r4 | RN7SL501P-D + TP53-M | 31 | 6,7,8,9,10,11,12,13 |
| r5 | BRD4-A + TP53-M | 24 | 6,7,8,9,10,11,12,13 |
| r19 | MYC-A + TCF3-D + TP53-M | 31 | 6,7,8,9,10,11,12,13 |
| r11 | d15(n=5) + PPP2R2A-D + TP53-M | 13 | 5,8,9,10,11,12,13 |
| r15 | CCNE1-A + RN7SL566P/SAMD4B-A + TP53-M | 13 | 7,8,9,10,11,12,13 |
| r2 | MECOM-A + TP53-M | 49 | 3,4,5,6,7,8 |
| r18 | d2(n=15) + FKSG52/MIR582/PDE4D-D + MYC-A + TP53-M | 18 | 3,4,5,6,7,8 |
| r1 | TCF3-D + TP53-M | 50 | 1,2,3,4,5 |
| r7 | d2(n=15) + FKSG52/MIR582/PDE4D-D + MECOM-A + MYC-A + TP53-M | 13 | 9,10,11,12,13 |
| r12 | MECOM-A + TCF3-D + TP53-M | 29 | 9,10,11,12,13 |
| r23 | d6(n=6) + MECOM-A + MYC-A + TP53-M | 13 | 9,10,11,12,13 |
| r16 | ANKS1B/FAM71C/RNA5SP366-D + d13(n=19) + TP53-M | 13 | 10,11,12,13 |
| r179 | CBX8-A + MYC-A | 25 | 11,12,13 |
| r185 | d2(n=15) + FKSG52/MIR582/PDE4D-D + TCF3-D | 18 | 12,13 |
| r9 | CCNE1-A + TP53-M | 27 | 4 |
| r10 | MECOM-A + MYC-A + TP53-M | 33 | 2 |
| r30 | BRD4-A + CCNE1-A + TP53-M | 12 | 5 |
| r176 | d2(n=15) + DEAF1/DRD4/TMEM80-D + FKSG52/MIR582/PDE4D-D + TP53-M | 12 | 13 |

**ID** = Rule IDs, rules are numbered according to importance rank determined from phase 1
**PC** = Percent of samples covered **Ks** = Membership in best RS

### 3 Core Rule Set

**Core K = 11.**
**Core rule set coverage = 83.3%.**

#### 3.1 Table of core rule set rules

| ID | Rule | CR | SJR | SJ | NSC | NSA | PC | PA | FracA |
|---|---|---|---|---|---|---|---|---|---|
| r3 | BSPH1-D + RN7SL526P-D + TCF3-D + TP53-M | 460.5 | 85 | 441 | 54 | 42 | 12 | 9.2 | 0.78 |
| r4 | RN7SL501P-D + TP53-M | 9 | 8 | 717 | 143 | 26 | 31 | 5.7 | 0.18 |
| r5 | BRD4-A + TP53-M | 27.5 | 24 | 579 | 110 | 26 | 24 | 5.7 | 0.24 |
| r7 | d2(n=15) + FKSG52/MIR582/PDE4D-D + MECOM-A + MYC-A + TP53-M | 329.5 | 39 | 520 | 60 | 56 | 13 | 12.0 | 0.93 |
| r11 | d15(n=5) + PPP2R2A-D + TP53-M | 386.5 | 160.5 | 374 | 57 | 32 | 13 | 7.0 | 0.56 |
| r12 | MECOM-A + TCF3-D + TP53-M | 13.5 | 6 | 862 | 133 | 39 | 29 | 8.6 | 0.29 |
| r15 | CCNE1-A + RN7SL566P/SAMD4B-A + TP53-M | 348 | 119 | 404 | 59 | 31 | 13 | 6.8 | 0.53 |
| r16 | ANKS1B/FAM71C/RNA5SP366-D + d13(n=19) + TP53-M | 366 | 202.5 | 350 | 58 | 26 | 13 | 5.7 | 0.45 |
| r19 | MYC-A + TCF3-D + TP53-M | 11 | 5 | 895 | 139 | 37 | 31 | 8.1 | 0.27 |
| r23 | d6(n=6) + MECOM-A + MYC-A + TP53-M | 366 | 108 | 415 | 58 | 39 | 13 | 8.6 | 0.67 |
| r179 | CBX8-A + MYC-A | 24 | 337.5 | 289 | 114 | 25 | 25 | 5.5 | 0.22 |

**ID** = Rule IDs, rules are numbered according to importance rank determined from phase 1
**CR** = Coverage rank
**SJ/SJR** = Single rule performance / performance rank
**NSC/PC** = Number/percent of samples covered
**NSA/PA** = Number/percent of samples assigned
**FracA** = Fraction of covered samples assigned to rule

#### 3.2 Event breakdown of core rule set

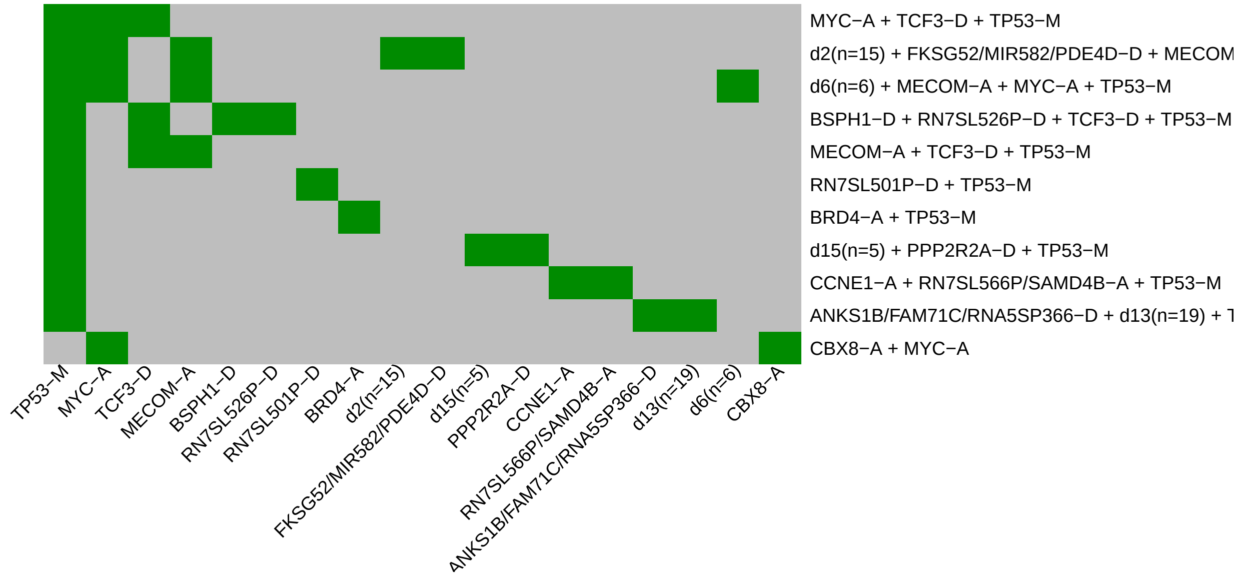

Figure 3: Visualization of core rule set. Rows are rules, columns are events. Events are ordered from left in decreasing rule membership frequency.

#### 3.3 Core Rule Set Penalties: Before and After Assignment

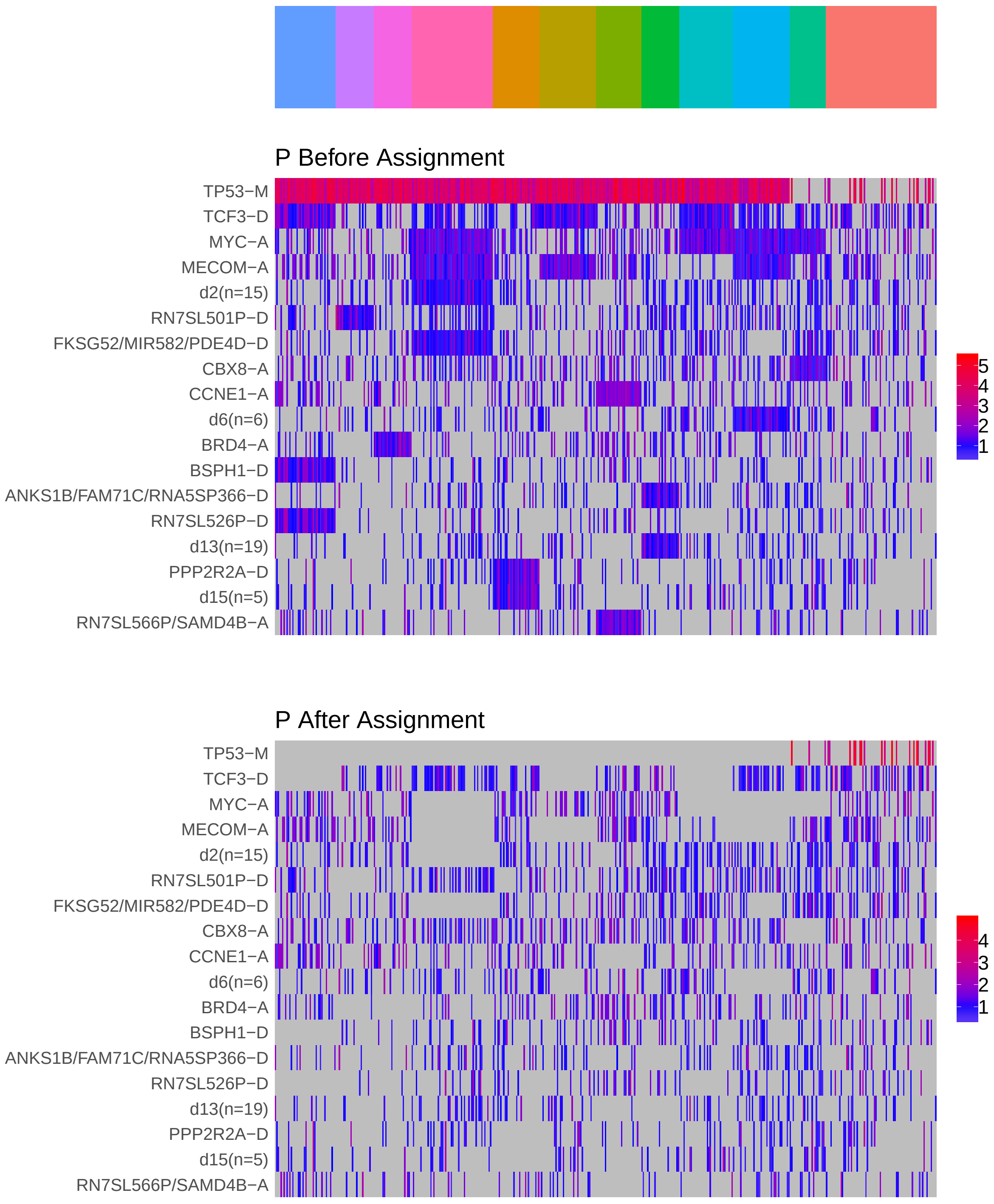

Figure 4: Heatmap of P before and after assignment to core rule set. Events are force ordered by frequency. Samples are force ordered according rule set membership, as indicated by the color bar. The right-most group of samples are not assigned to any rule.

#### 4 Generalized Core Rules

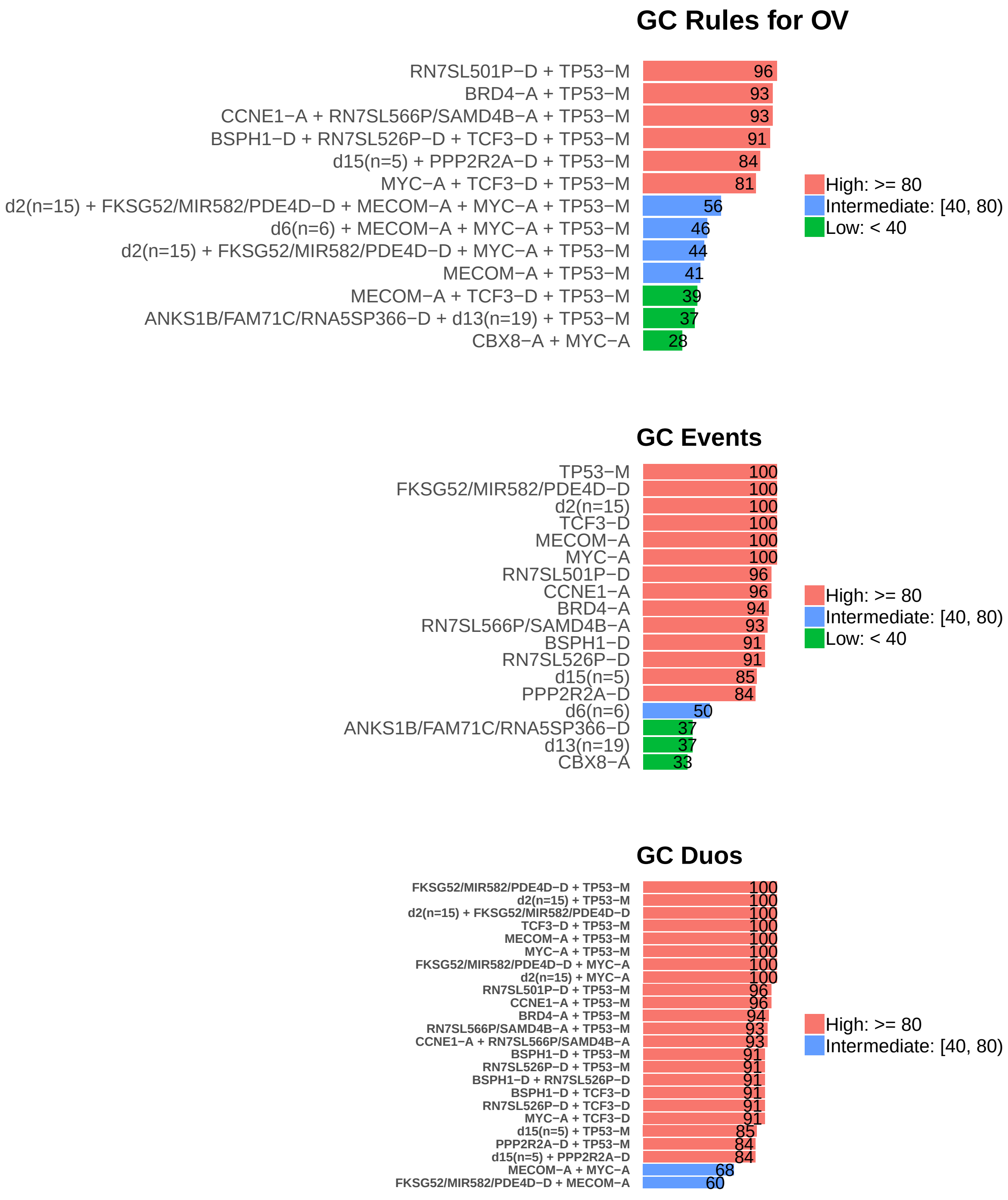

Figure 5: Summary of generalized core results. Bars show confidence levels, which are the percentage of sub-sample iterations containing the observation. Rules (top) and events (middle) that achieve a minimum confidence level of 20 are shown. Duos (bottom) with confidence of at least 60 are shown.

### 5 Dictionary of Copy Number Events

| CNV | Genes | Event__Name |
|---|---|---|
| a1 | MYC | MYC-A |
| a2 | MECOM | MECOM-A |
| a3 | CBX8 | CBX8-A |
| a4 | CCNE1 | CCNE1-A |
| a5 | GOLPH3L | GOLPH3L-A |
| a6 | USP35 | USP35-A |
| a7 | RNF144B | RNF144B-A |
| a8 | BRD4 | BRD4-A |
| a9 | UBE3C | UBE3C-A |
| a10 | BMP8A, MACF1 | BMP8A/MACF1-A |
| a11 | SYNM | SYNM-A |
| a13 | HELZ2 | HELZ2-A |
| a14 | RN7SL566P, SAMD4B | RN7SL566P/SAMD4B-A |
| d1 | TCF3 | TCF3-D |
| d2 | snoU13\|ENSG00000238451.1, GTF2H2B, RN7SL9P, snoU13\|ENSG00000238740.1, GUSBP3, RN7SL616P, RN7SL476P, GTF2H2, NAIP, SMN1, SMN2, SERF1A, GTF2H2C, SERF1B, OCLN | d2(n=15) |
| d3 | FKSG52, MIR582, PDE4D | FKSG52/MIR582/PDE4D-D |
| d4 | RN7SL501P | RN7SL501P-D |
| d5 | TMEM80, DRD4, DEAF1 | DEAF1/DRD4/TMEM80-D |
| d6 | TCP10, TTLL2, GPR31, C6orf123, UNC93A, TCP10L2 | d6(n=6) |
| d7 | SDK1 | SDK1-D |
| d8 | BSPH1 | BSPH1-D |
| d9 | LINC00901, TUSC7, RN7SL582P, LINC00903, RN7SL815P, LSAMP, MIR4447 | d9(n=7) |
| d10 | RN7SL526P | RN7SL526P-D |
| d11 | LRP1B | LRP1B-D |
| d12 | PPP2R2A | PPP2R2A-D |
| d13 | ANHX, ZNF891, ZNF140, RNU4ATAC12P, RNA5SP379, LRCOL1, GOLGA3, POLE, PXMP2, ZNF10, ZNF26, ZNF84, ZNF268, P2RX2, ANKLE2, CHFR, FBRSL1, PGAM5, ZNF605 | d13(n=19) |
| d14 | RNA5SP366, ANKS1B, FAM71C | ANKS1B/FAM71C/RNA5SP366-D |
| d15 | RPL23AP53, OR4F21, FBXO25, TDRP, ZNF596 | d15(n=5) |

Dictionary of important CNVs. Copy number events that are within any of the best K rule sets, or within the generalized core rules, or among the 20 most frequent events.
