## Supplementary material for "Identifying Combinations of Cancer Drivers in Individual Patients": TCGA Reports: CRSO_Report_PAAD.pdf

### CRSO Output Report: PAAD

Michael Klein

Apr 19 2019

**4 Generalized Core Rules** **6**

**5 Dictionary of Copy Number Events** **7**

### 1 Heatmaps of D and P

**Number of samples = 126.**
**Number of events = 65.**
**Rule coverage requirement = 6 samples.**
**Rule library size = 210 rules.**

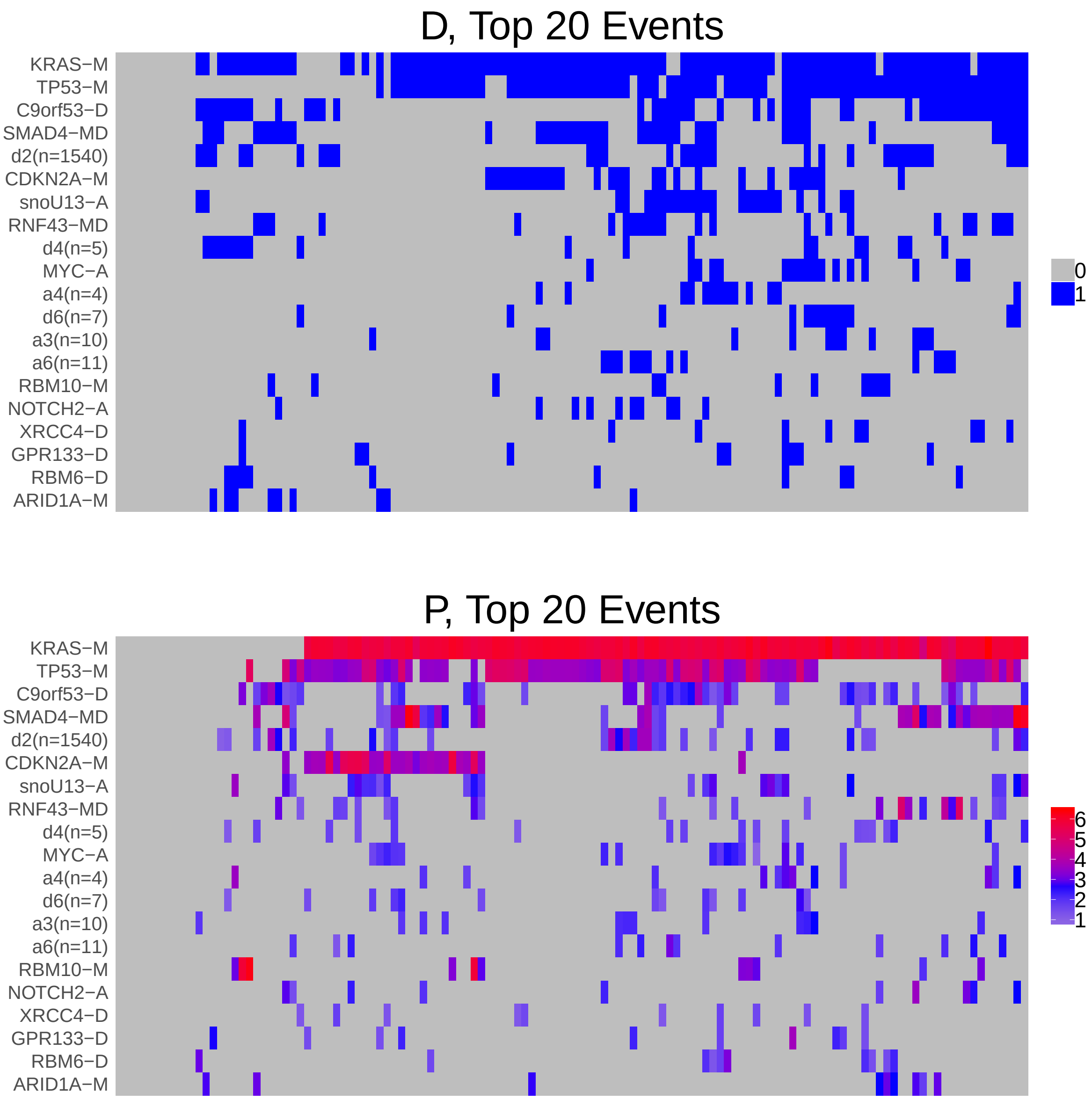

Figure 1: Heatmap of D and P. Events are ordered by frequency, from top to bottom. Event suffix -M: mutation, -A: amplification, -D: deletion, -MD: mutDel, -MA: mutAmp. Samples are ordered using hierarchical clustering. Wild-type events indicated in grey.

### 2 Summary of K Best Rule Sets

#### 2.1 Performance and coverage of best rule sets

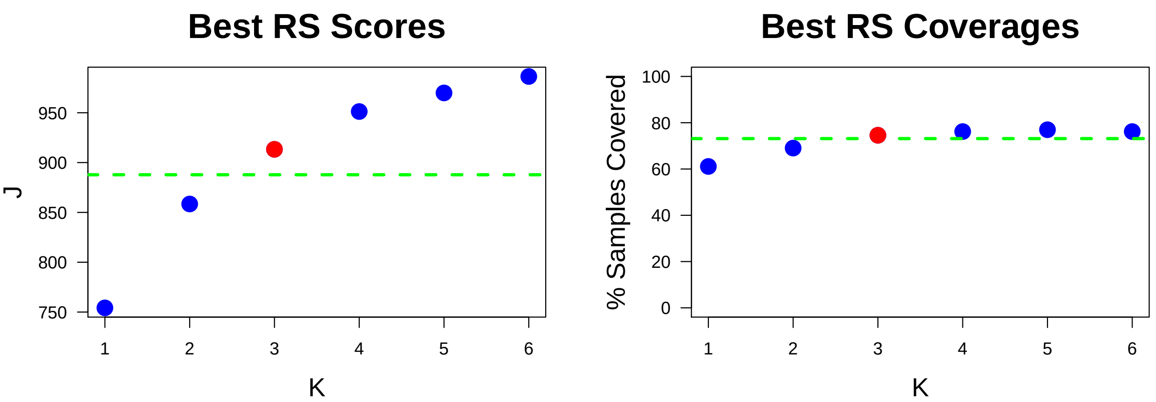

Figure 2: Performance and Coverage of Best Filtered RS for K. The core was determined to be the smallest RS that achieved at least 95% of maximum coverage and at least 90% of maximum performance.

#### 2.2 Table of rules that appear in any best rule set

| ID | Rule | PC | Ks |
|---|---|---|---|
| r1 | KRAS-M + TP53-M | 61 | 1,2,3,4,5,6 |
| r3 | KRAS-M + SMAD4-MD | 29 | 2,3,4,5,6 |
| r4 | CDKN2A-M + KRAS-M | 21 | 4,5,6 |
| r5 | C9orf53-D + KRAS-M | 29 | 3,4,5 |
| r60 | CDKN2A-M + SMAD4-MD + TP53-M | 8.7 | 5,6 |
| r39 | C9orf53-D + KRAS-M + RBM6-D | 6.3 | 6 |
| r40 | C9orf53-D + d2(n=1540) + KRAS-M | 11 | 6 |

**ID** = Rule IDs, rules are numbered according to importance rank determined from phase 1
**PC** = Percent of samples covered **Ks** = Membership in best RS

### 3 Core Rule Set

**Core K = 3.**
**Core rule set coverage = 74.6%.**

#### 3.1 Table of core rule set rules

| ID | Rule | CR | SJR | SJ | NSC | NSA | PC | PA | FracA |
|---|---|---|---|---|---|---|---|---|---|
| r1 | KRAS-M + TP53-M | 1 | 1 | 754 | 77 | 67 | 61 | 53.0 | 0.87 |
| r3 | KRAS-M + SMAD4-MD | 3 | 3 | 331 | 36 | 20 | 29 | 16.0 | 0.56 |
| r5 | C9orf53-D + KRAS-M | 2 | 5 | 294 | 37 | 7 | 29 | 5.6 | 0.19 |

**ID** = Rule IDs, rules are numbered according to importance rank determined from phase 1
**CR** = Coverage rank
**SJ/SJR** = Single rule performance / performance rank
**NSC/PC** = Number/percent of samples covered
**NSA/PA** = Number/percent of samples assigned
**FracA** = Fraction of covered samples assigned to rule

#### 3.2 Event breakdown of core rule set

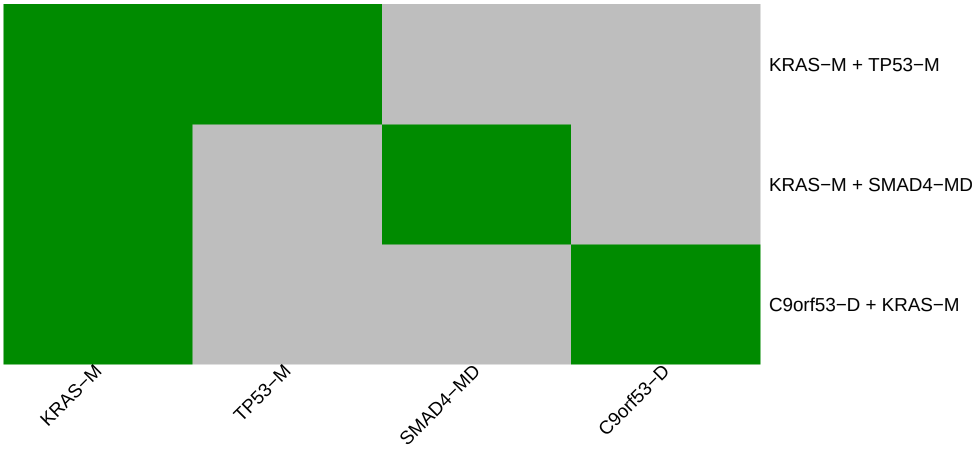

Figure 3: Visualization of core rule set. Rows are rules, columns are events. Events are ordered from left in decreasing rule membership frequency.

#### 3.3 Core Rule Set Penalties: Before and After Assignment

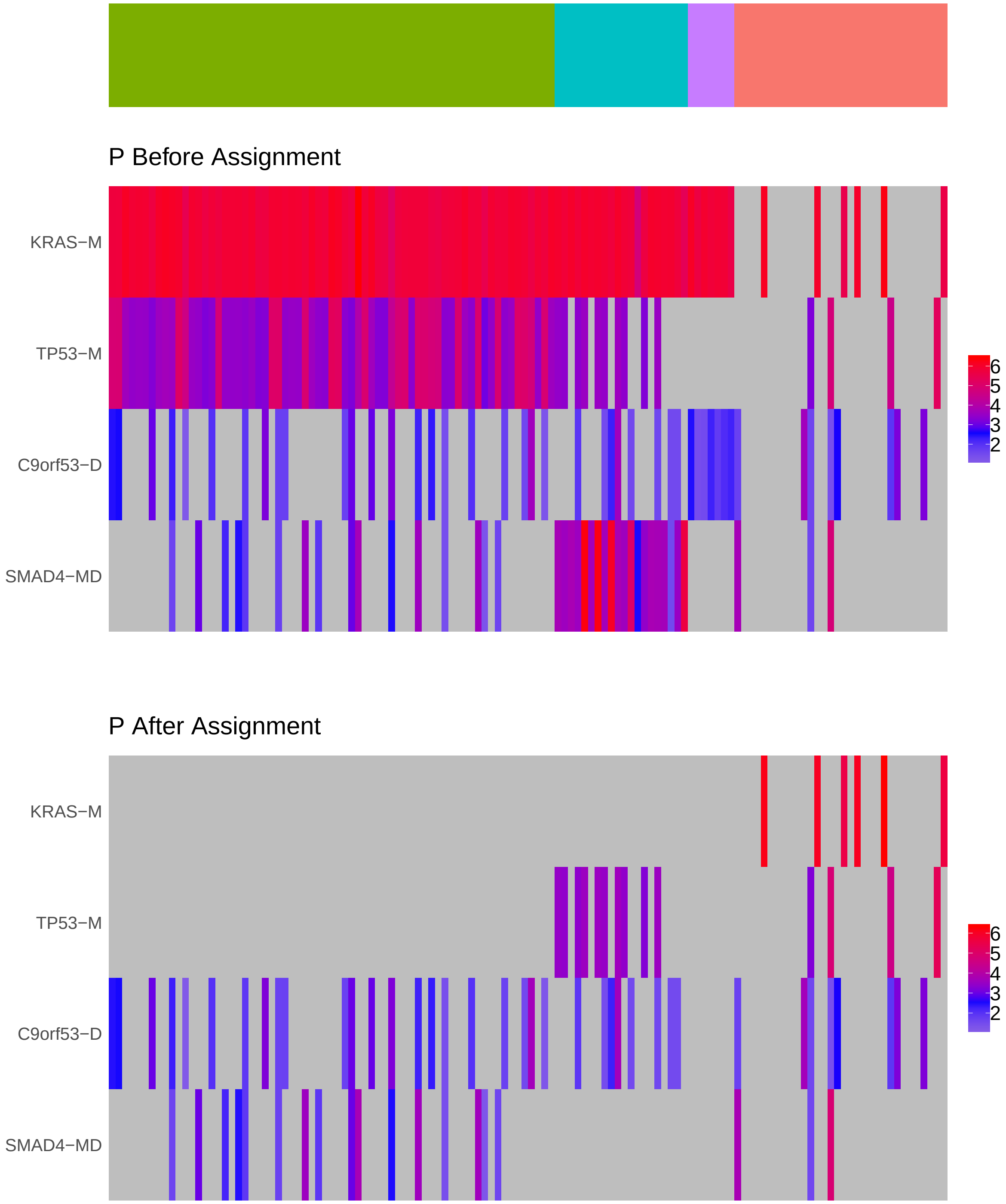

Figure 4: Heatmap of P before and after assignment to core rule set. Events are force ordered by frequency. Samples are force ordered according rule set membership, as indicated by the color bar. The right-most group of samples are not assigned to any rule.

### 4 Generalized Core Rules

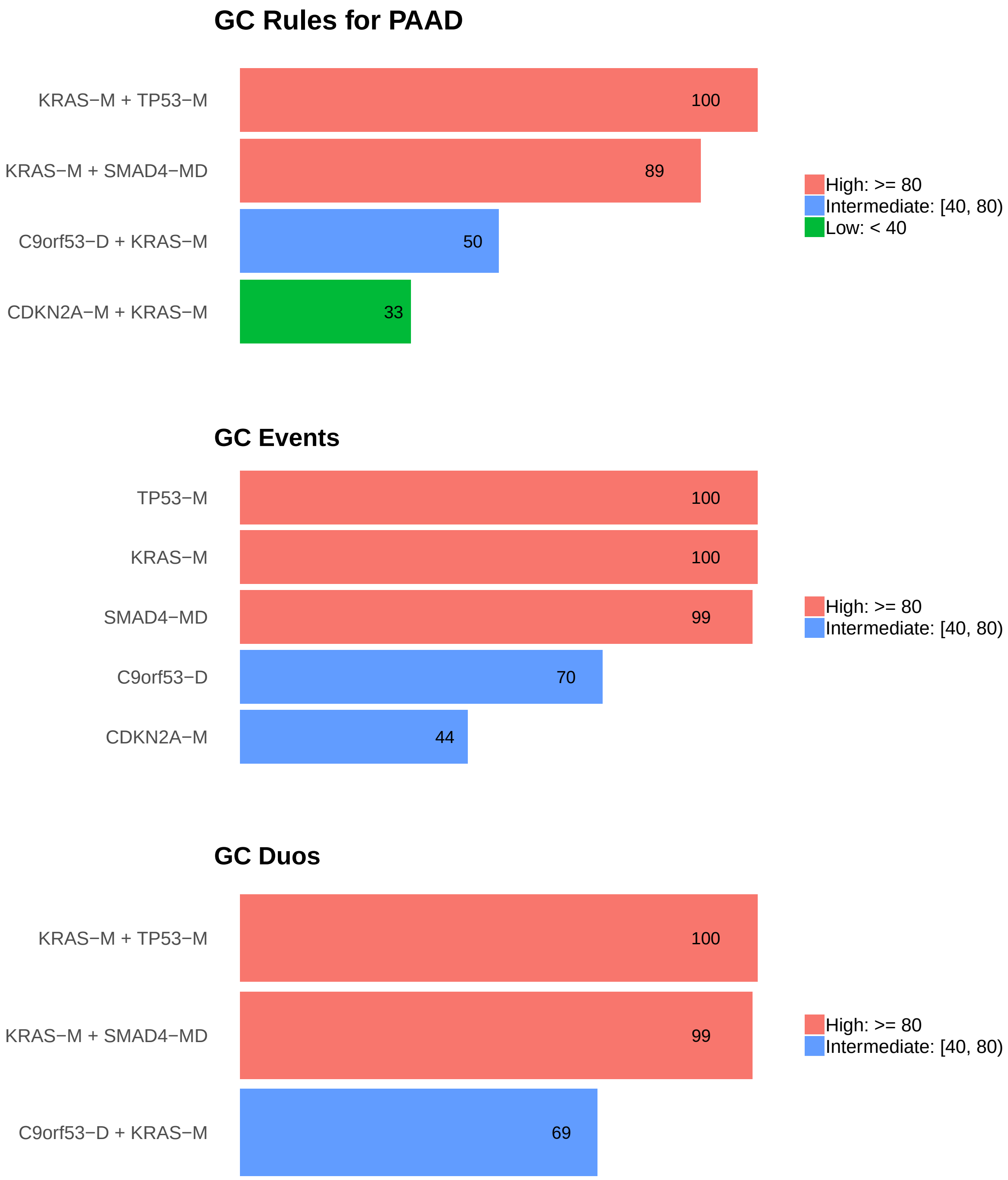

Figure 5: Summary of generalized core results. Bars show confidence levels, which are the percentage of sub-sample iterations containing the observation. Rules (top) and events (middle) that achieve a minimum confidence level of 20 are shown. Duos (bottom) with confidence of at least 50 are shown.

### 5 Dictionary of Copy Number Events

| CNV | Genes | Event_Name |
|---|---|---|
| a1 | snoU13\|ENSG00000238907.1 | snoU13-A |
| a2 | MYC | MYC-A |
| a3 | RN7SL566P, RPS16, SUPT5H, ZFP36, GMFG, PAF1, SAMD4B, MED29, PLEKHG2, MIR4530 | a3(n=10) |
| a4 | RN7SL22P, CA9, TPM2, ARHGEF39 | a4(n=4) |
| a5 | NOTCH2 | NOTCH2-A |
| a6 | IKZF3, MIR4728, PNMT, TCAP, NEUROD2, ERBB2, GRB7, STARD3, PPP1R1B, MIEN1, PGAP3 | a6(n=11) |
| d1 | C9orf53 | C9orf53-D |
| d2 | d2(n=1540) | d2(n=1540) |
| d4 | GPN2, ZDHHC18, RN7SL165P, SFN, PIGV | d4(n=5) |
| d6 | GAMT, RPS15, APC2, DAZAP1, PCSK4, REEP6, C19orf25 | d6(n=7) |
| d8 | XRCC4 | XRCC4-D |
| d9 | GPR133 | GPR133-D |
| d13 | RBM6 | RBM6-D |

Dictionary of important CNVs. Copy number events that are within any of the best K rule sets, or within the generalized core rules, or among the 20 most frequent events.
