## Supplementary material for "Identifying Combinations of Cancer Drivers in Individual Patients": TCGA Reports: CRSO_Report_PRAD.pdf

### CRSO Output Report: PRAD

Michael Klein

Apr 19 2019

**4 Generalized Core Rules** **6**

**5 Dictionary of Copy Number Events** **7**

#### 1 Heatmaps of D and P

**Number of samples = 492.**
**Number of events = 73.**
**Rule coverage requirement = 15 samples.**
**Rule library size = 301 rules.**

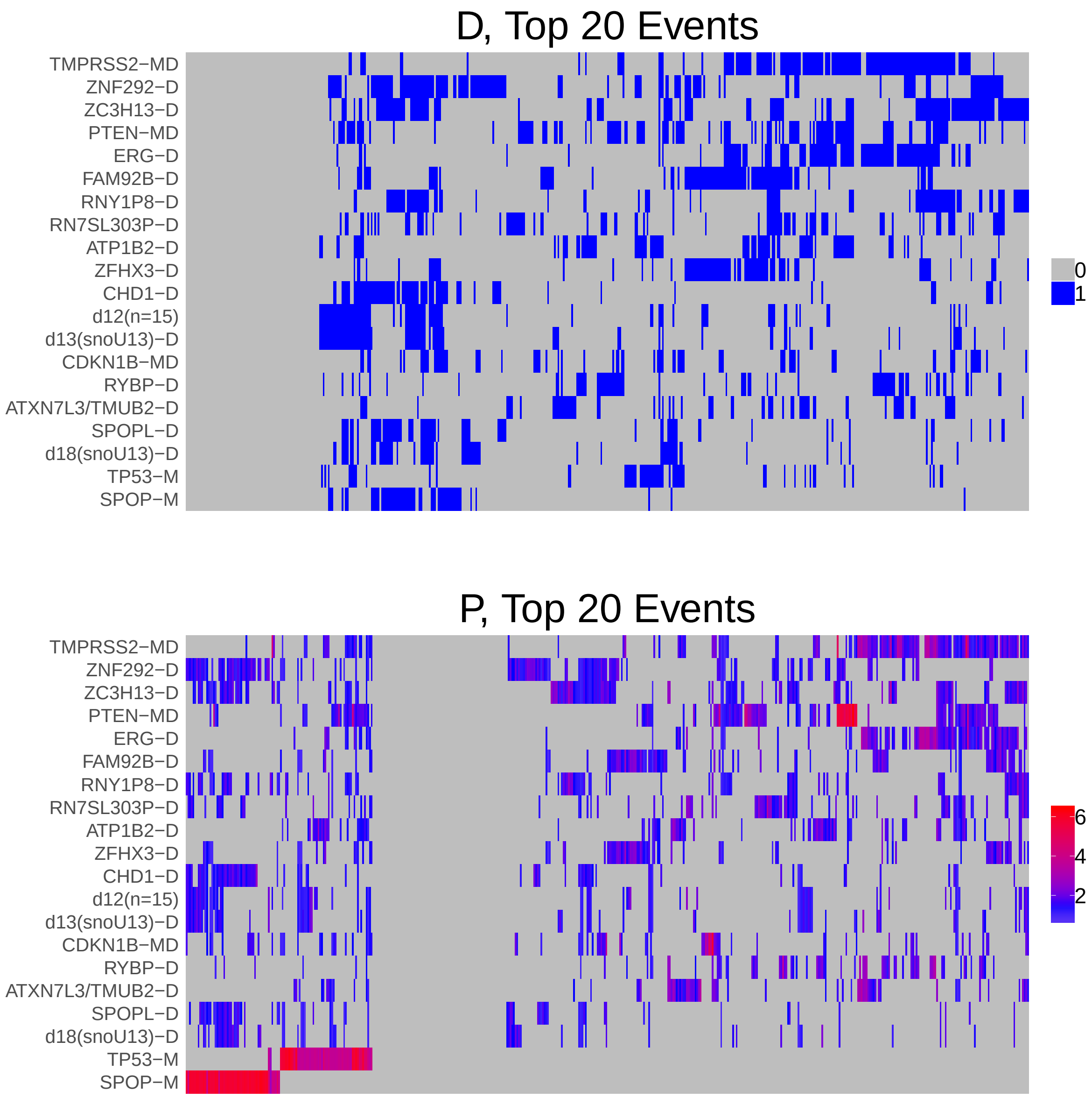

Figure 1: Heatmap of D and P. Events are ordered by frequency, from top to bottom. Event suffix -M: mutation, -A: amplification, -D: deletion, -MD: mutDel, -MA: mutAmp. Samples are ordered using hierarchical clustering. Wild-type events indicated in grey.

### 2 Summary of K Best Rule Sets

#### 2.1 Performance and coverage of best rule sets

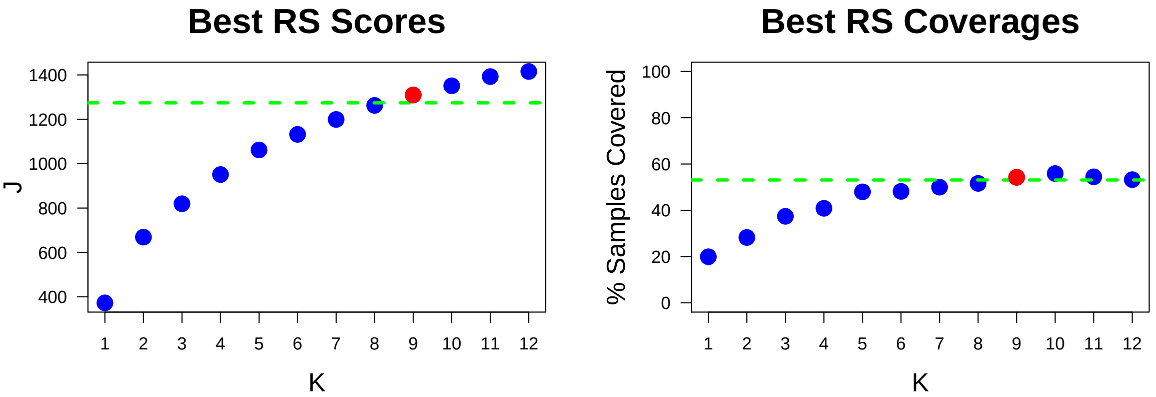

Figure 2: Performance and Coverage of Best Filtered RS for K. The core was determined to be the smallest RS that achieved at least 95% of maximum coverage and at least 90% of maximum performance.

#### 2.2 Table of rules that appear in any best rule set

| ID | Rule | PC | Ks |
|---|---|---|---|
| r5 | SPOP-M + ZNF292-D | 8.3 | 2,3,4,5,6,7,8,9,10,11,12 |
| r1 | ERG-D + TMPRSS2-MD | 20 | 1,2,3,4,5,6,7,8,9,10 |
| r2 | FAM92B-D + ZFHX3-D | 14 | 3,4,5,6,7,8,9,10,11,12 |
| r3 | RNY1P8-D + ZC3H13-D | 16 | 5,6,7,8,9,10,11 |
| r4 | PTEN-MD + TP53-M | 5.3 | 4,5,6,7,8,9,10 |
| r10 | FOXA1-M + ZNF292-D | 3.9 | 8,9,10,11,12 |
| r58 | CHD1-D + SPOP-M + SPOPL-D + ZC3H13-D | 3.5 | 6,7,9,11 |
| r7 | PTEN-MD + TMPRSS2-MD | 13 | 7,8,9 |
| r16 | PTEN-MD + RN7SL303P-D | 7.1 | 10,11,12 |
| r17 | CHD1-D + d18(snoU13)-D + SPOP-M + SPOPL-D | 3.3 | 8,10,12 |
| r29 | CDKN1B-MD + ZNF292-D | 7.9 | 9,10,11 |
| r6 | ATP1B2-D + TP53-M | 4.7 | 11,12 |
| r9 | ERG-D + PTEN-MD + TMPRSS2-MD | 9.3 | 11,12 |
| r15 | ERG-D + RYBP-D + TMPRSS2-MD | 5.9 | 11,12 |
| r24 | ATXN7L3/TMUB2-D + TMPRSS2-MD | 7.9 | 11,12 |
| r11 | ZC3H13-D + ZNF292-D | 12 | 12 |
| r13 | d12(n=15) + d13(snoU13)-D | 12 | 12 |
| r14 | ERG-D + RNY1P8-D + TMPRSS2-MD + ZC3H13-D | 4.5 | 12 |
| r74 | ATP1B2-D + PTEN-MD + TMPRSS2-MD | 4.7 | 10 |

**ID** = Rule IDs, rules are numbered according to importance rank determined from phase 1
**PC** = Percent of samples covered **Ks** = Membership in best RS

### 3 Core Rule Set

**Core K = 9.**
**Core rule set coverage = 54.3%.**

#### 3.1 Table of core rule set rules

| ID | Rule | CR | SJR | SJ | NSC | NSA | PC | PA | FracA |
|---|---|---|---|---|---|---|---|---|---|
| r1 | ERG-D + TMPRSS2-MD | 1 | 1 | 373 | 98 | 72 | 20 | 15.0 | 0.73 |
| r2 | FAM92B-D + ZFHX3-D | 3 | 8 | 224 | 68 | 40 | 14 | 8.1 | 0.59 |
| r3 | RNY1P8-D + ZC3H13-D | 2 | 6 | 250 | 79 | 24 | 16 | 4.9 | 0.30 |
| r4 | PTEN-MD + TP53-M | 86 | 35 | 158 | 26 | 26 | 5.3 | 5.3 | 1.00 |
| r5 | SPOP-M + ZNF292-D | 19 | 2 | 296 | 41 | 28 | 8.3 | 5.7 | 0.68 |
| r7 | PTEN-MD + TMPRSS2-MD | 4 | 5 | 258 | 66 | 29 | 13 | 5.9 | 0.44 |
| r10 | FOXA1-M + ZNF292-D | 182.5 | 102.5 | 111 | 19 | 16 | 3.9 | 3.3 | 0.84 |
| r29 | CDKN1B-MD + ZNF292-D | 21 | 84.5 | 120 | 39 | 15 | 7.9 | 3.0 | 0.38 |
| r58 | CHD1-D + SPOP-M + SPOPL-D + ZC3H13-D | 227.5 | 22.5 | 172 | 17 | 17 | 3.5 | 3.5 | 1.00 |

**ID** = Rule IDs, rules are numbered according to importance rank determined from phase 1
**CR** = Coverage rank
**SJ/SJR** = Single rule performance / performance rank
**NSC/PC** = Number/percent of samples covered
**NSA/PA** = Number/percent of samples assigned
**FracA** = Fraction of covered samples assigned to rule

#### 3.2 Event breakdown of core rule set

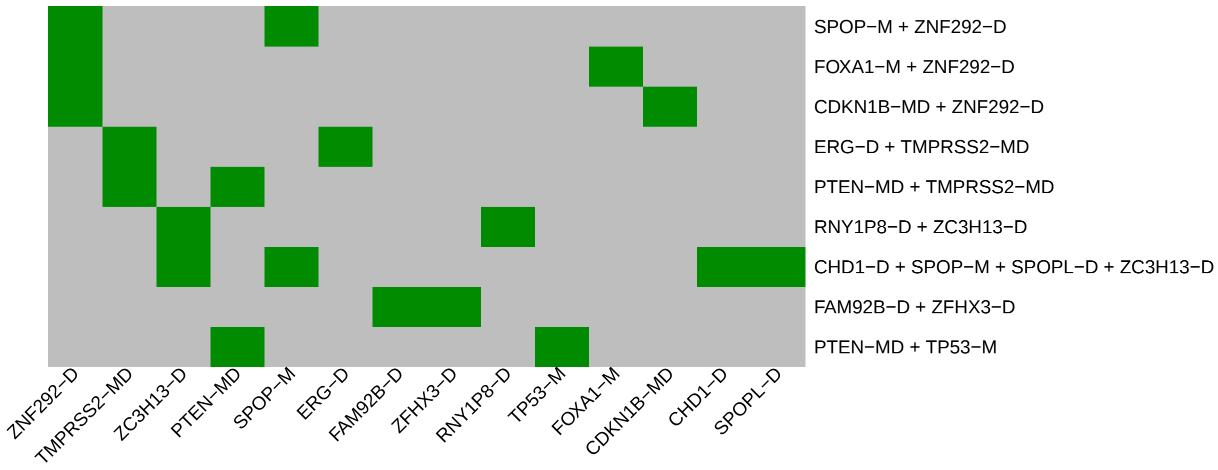

Figure 3: Visualization of core rule set. Rows are rules, columns are events. Events are ordered from left in decreasing rule membership frequency.

##### 3.3 Core Rule Set Penalties: Before and After Assignment

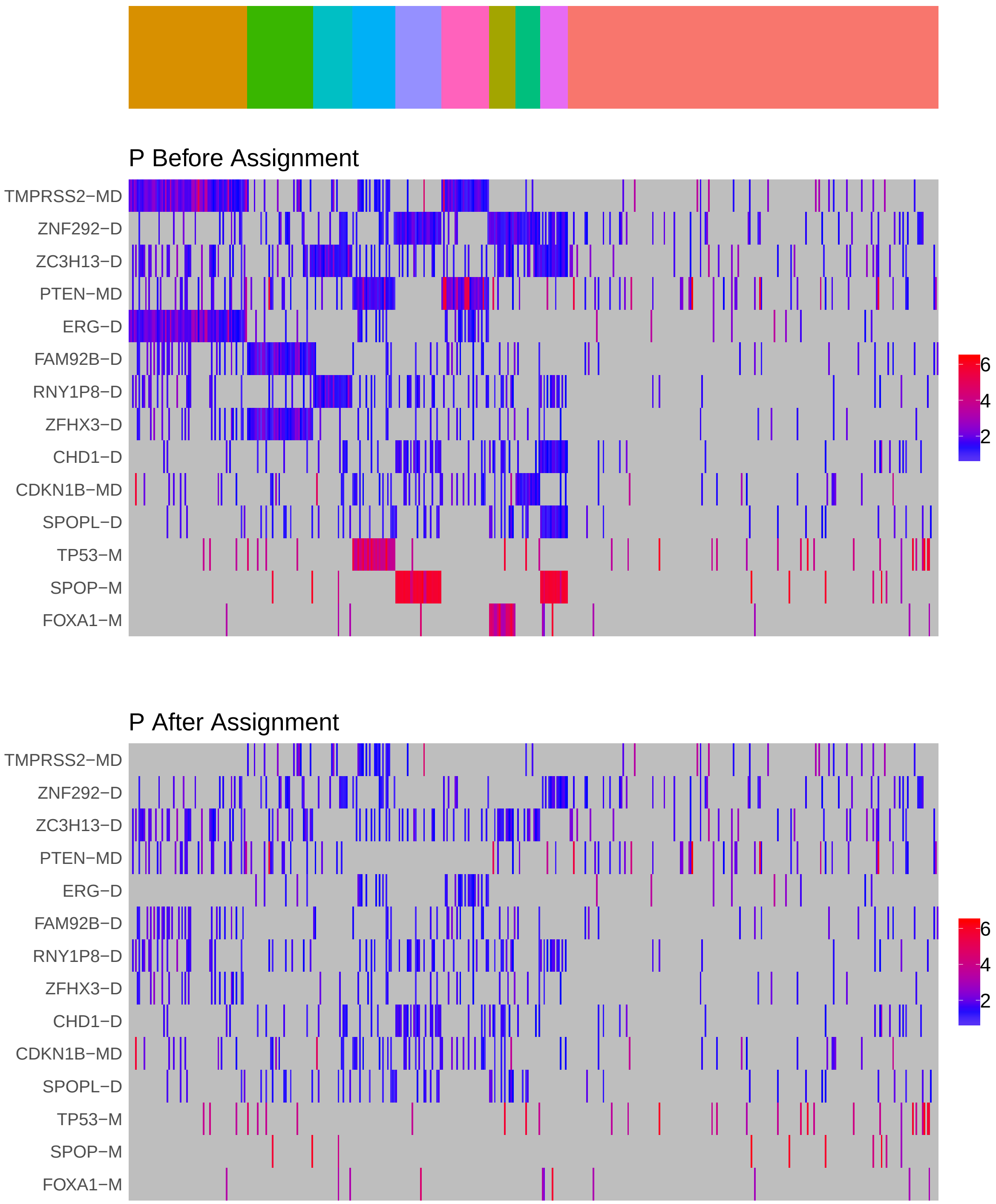

Figure 4: Heatmap of P before and after assignment to core rule set. Events are force ordered by frequency. Samples are force ordered according rule set membership, as indicated by the color bar. The right-most group of samples are not assigned to any rule.

### 4 Generalized Core Rules

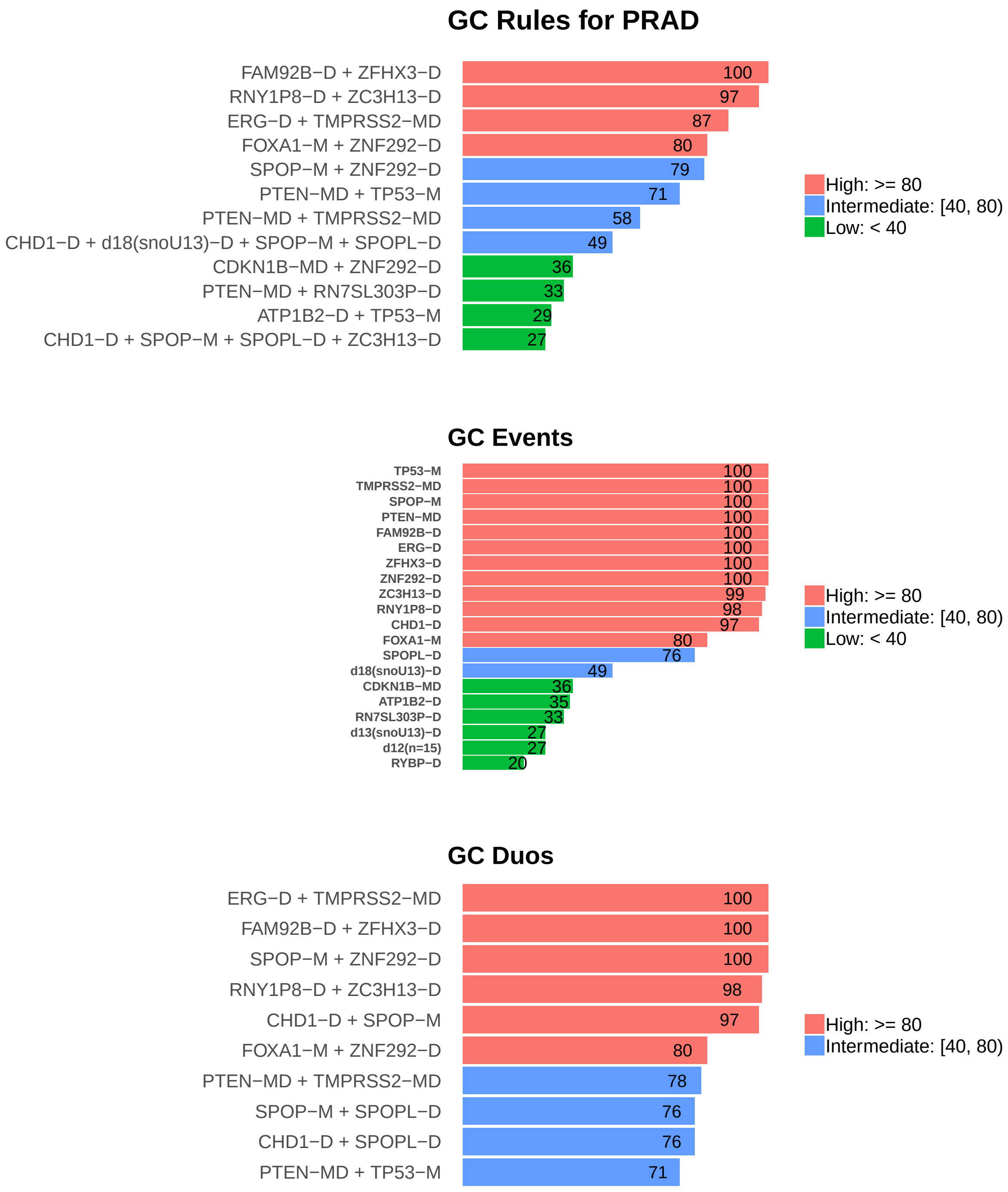

Figure 5: Summary of generalized core results. Bars show confidence levels, which are the percentage of sub-sample iterations containing the observation. Rules (top) and events (middle) that achieve a minimum confidence level of 20 are shown. Duos (bottom) with confidence of at least 50 are shown.

### 5 Dictionary of Copy Number Events

| CNV | Genes | Event_Name |
|---|---|---|
| d1 | ZNF292 | ZNF292-D |
| d2 | ZC3H13 | ZC3H13-D |
| d5 | ERG | ERG-D |
| d6 | FAM92B | FAM92B-D |
| d7 | RNY1P8 | RNY1P8-D |
| d8 | RN7SL303P | RN7SL303P-D |
| d9 | ATP1B2 | ATP1B2-D |
| d10 | ZFHX3 | ZFHX3-D |
| d11 | CHD1 | CHD1-D |
| d12 | snoU13\|ENSG00000238451.1, GTF2H2B, RN7SL9P, snoU13\|ENSG00000238740.1, GUSBP3, RN7SL616P, RN7SL476P, GTF2H2, NAIP, SMN1, SMN2, SERF1A, GTF2H2C, SERF1B, OCLN | d12(n=15) |
| d13 | snoU13\|ENSG00000238717.1 | d13(snoU13)-D |
| d14 | RYBP | RYBP-D |
| d16 | ATXN7L3, TMUB2 | ATXN7L3/TMUB2-D |
| d17 | SPOPL | SPOPL-D |
| d18 | snoU13\|ENSG00000238860.1 | d18(snoU13)-D |

Dictionary of important CNVs. Copy number events that are within any of the best K rule sets, or within the generalized core rules, or among the 20 most frequent events.
