## Supplementary material for "Identifying Combinations of Cancer Drivers in Individual Patients": TCGA Reports: CRSO_Report_READ.pdf

### CRSO Output Report: READ

Michael Klein

Apr 19 2019

**4 Generalized Core Rules** **6**

**5 Dictionary of Copy Number Events** **7**

### 1 Heatmaps of D and P

**Number of samples = 120.**
**Number of events = 67.**
**Rule coverage requirement = 6 samples.**
**Rule library size = 735 rules.**

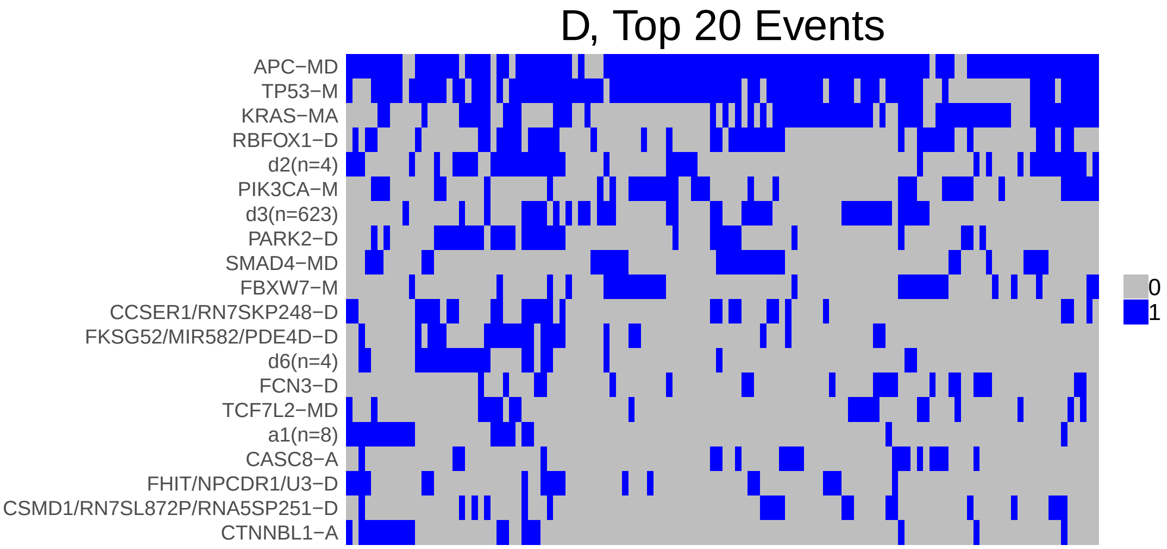

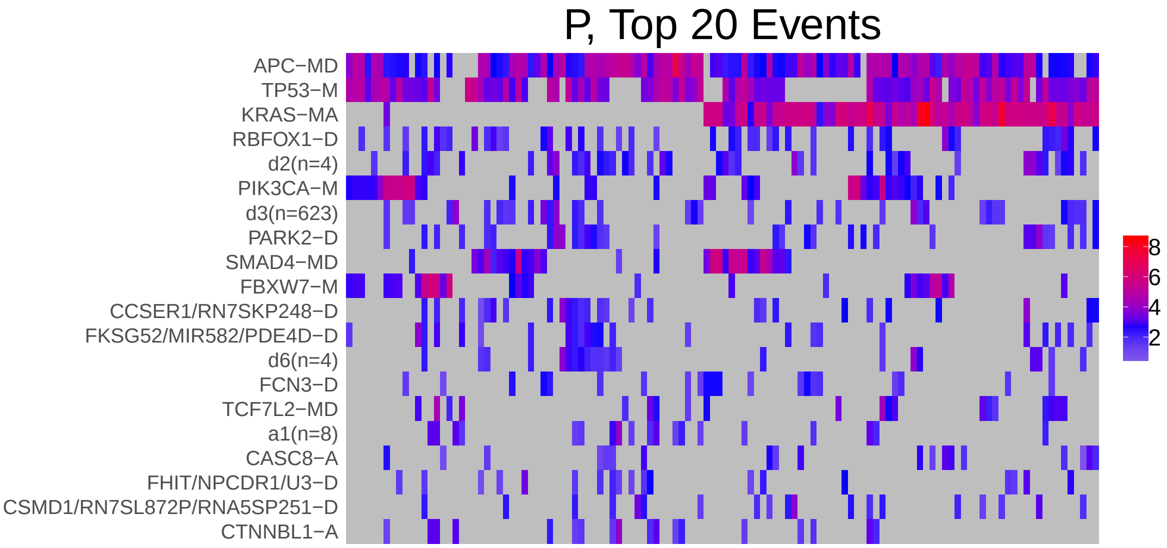

Figure 1: Heatmap of D and P. Events are ordered by frequency, from top to bottom. Event suffix -M: mutation, -A: amplification, -D: deletion, -MD: mutDel, -MA: mutAmp. Samples are ordered using hierarchical clustering. Wild-type events indicated in grey.

### 2 Summary of K Best Rule Sets

#### 2.1 Performance and coverage of best rule sets

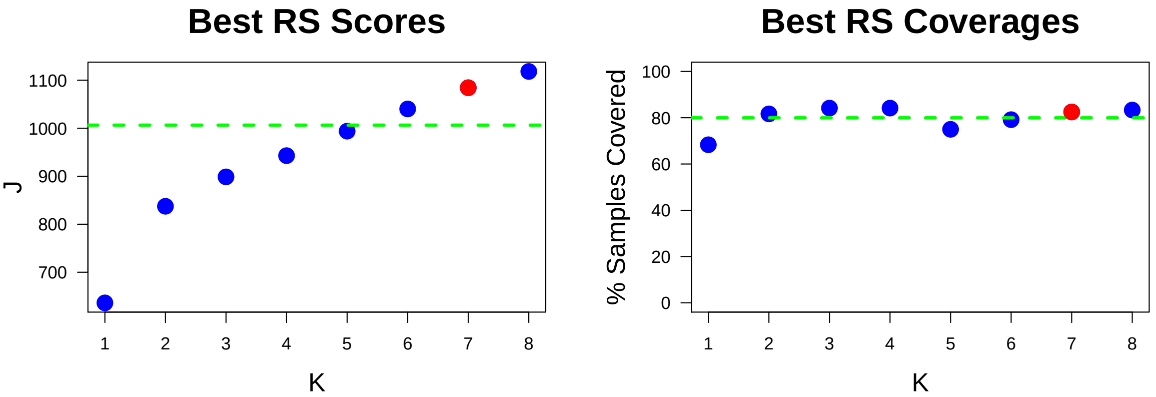

Figure 2: Performance and Coverage of Best Filtered RS for K. The core was determined to be the smallest RS that achieved at least 95% of maximum coverage and at least 90% of maximum performance.

#### 2.2 Table of rules that appear in any best rule set

| **ID** | **Rule** | **PC** | **Ks** |
|---|---|---|---|
| r1 | APC-MD + TP53-M | 68 | 1,2,3,4 |
| r3 | APC-MD + KRAS-MA + TP53-M | 36 | 5,6,7,8 |
| r4 | APC-MD + PIK3CA-M + TP53-M | 26 | 5,6,7,8 |
| r7 | APC-MD + d2(n=4) | 31 | 5,6,7,8 |
| r17 | APC-MD + SMAD4-MD | 22 | 5,6,7,8 |
| r2 | APC-MD + KRAS-MA | 49 | 2,3,4 |
| r105 | INS/MIR4686/TH-A + KRAS-MA | 6.7 | 6,7,8 |
| r5 | KRAS-MA + TP53-M | 38 | 3,4 |
| r16 | APC-MD + d3(n=623) + TP53-M | 22 | 7,8 |
| r23 | a1(n=8) + APC-MD + CTNNBL1-A + TP53-M | 8.3 | 7,8 |
| r65 | APC-MD + CCSER1/RN7SKP248-D + RBFOX1-D + TP53-M | 12 | 5,6 |
| r104 | APC-MD + KRAS-MA + PIK3CA-M + RBFOX1-D | 7.5 | 8 |
| r338 | CCSER1/RN7SKP248-D + d2(n=4) + FKSG52/MIR582/PDE4D-D + PARK2-D + TP53-M | 5.8 | 4 |

**ID** = Rule IDs, rules are numbered according to importance rank determined from phase 1
**PC** = Percent of samples covered **Ks** = Membership in best RS

### 3 Core Rule Set

**Core K = 7.**
**Core rule set coverage = 82.5%.**

#### 3.1 Table of core rule set rules

| ID | Rule | CR | SJR | SJ | NSC | NSA | PC | PA | FracA |
|---|---|---|---|---|---|---|---|---|---|
| r3 | APC-MD + KRAS-MA + TP53-M | 4 | 2 | 553 | 43 | 41 | 36 | 34.0 | 0.95 |
| r4 | APC-MD + PIK3CA-M + TP53-M | 11.5 | 5 | 339 | 31 | 16 | 26 | 13.0 | 0.52 |
| r7 | APC-MD + d2(n=4) | 6 | 14.5 | 230 | 37 | 9 | 31 | 7.5 | 0.24 |
| r16 | APC-MD + d3(n=623) + TP53-M | 18.5 | 8 | 251 | 26 | 9 | 22 | 7.5 | 0.35 |
| r17 | APC-MD + SMAD4-MD | 18.5 | 28 | 184 | 26 | 11 | 22 | 9.2 | 0.42 |
| r23 | a1(n=8) + APC-MD + CTNNBL1-A + TP53-M | 183 | 89 | 125 | 10 | 7 | 8.3 | 5.8 | 0.70 |
| r105 | INS/MIR4686/TH-A + KRAS-MA | 311.5 | 365 | 66.8 | 8 | 6 | 6.7 | 5.0 | 0.75 |

**ID** = Rule IDs, rules are numbered according to importance rank determined from phase 1
**CR** = Coverage rank
**SJ/SJR** = Single rule performance / performance rank
**NSC/PC** = Number/percent of samples covered
**NSA/PA** = Number/percent of samples assigned
**FracA** = Fraction of covered samples assigned to rule

#### 3.2 Event breakdown of core rule set

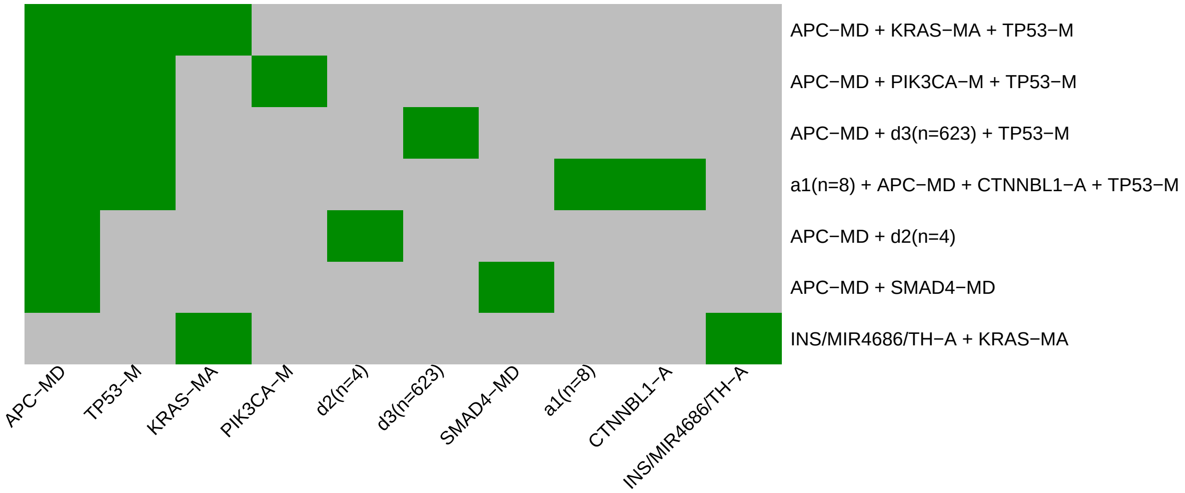

Figure 3: Visualization of core rule set. Rows are rules, columns are events. Events are ordered from left in decreasing rule membership frequency.

##### 3.3 Core Rule Set Penalties: Before and After Assignment

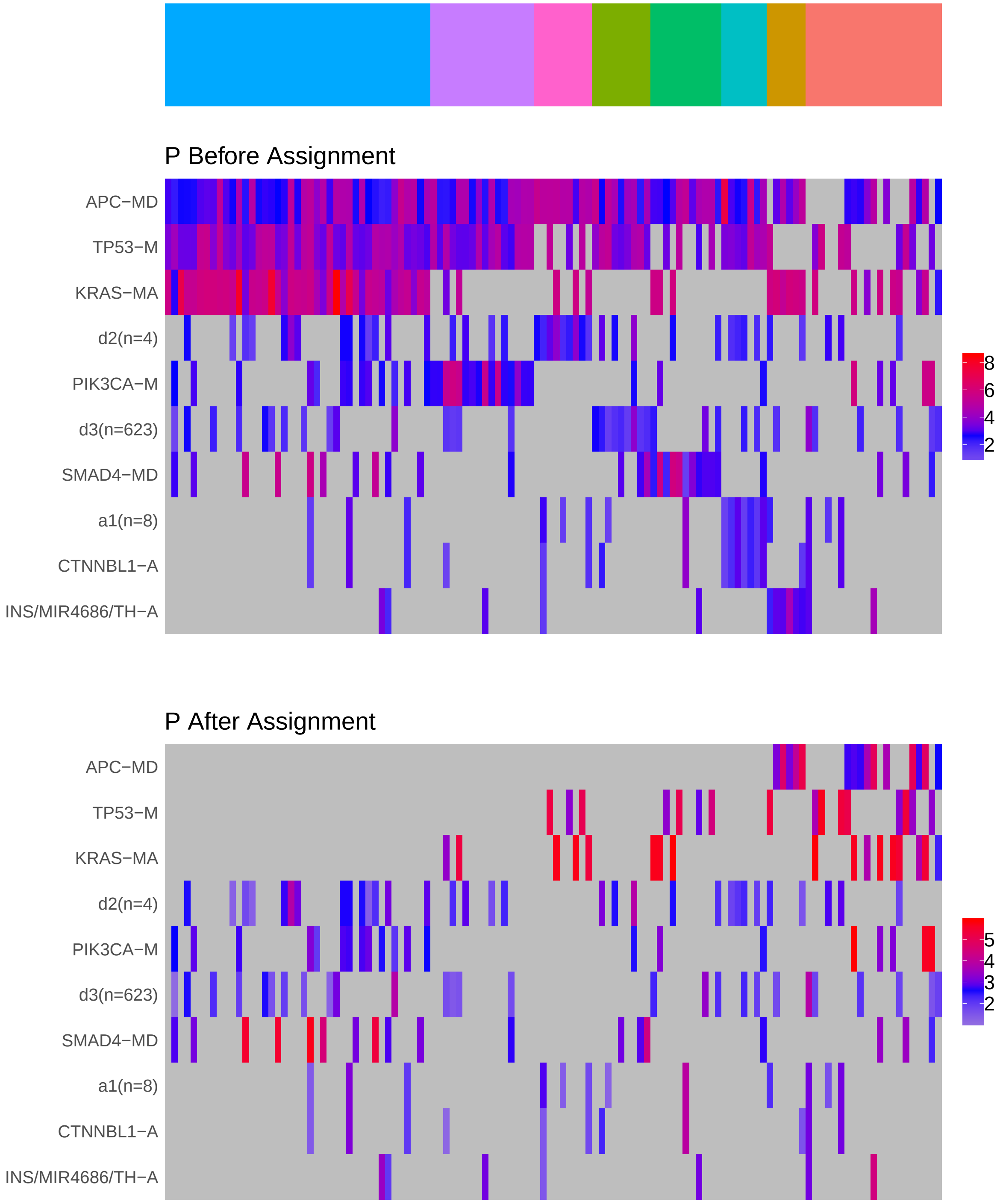

Figure 4: Heatmap of P before and after assignment to core rule set. Events are force ordered by frequency. Samples are force ordered according rule set membership, as indicated by the color bar. The right-most group of samples are not assigned to any rule.

### 4 Generalized Core Rules

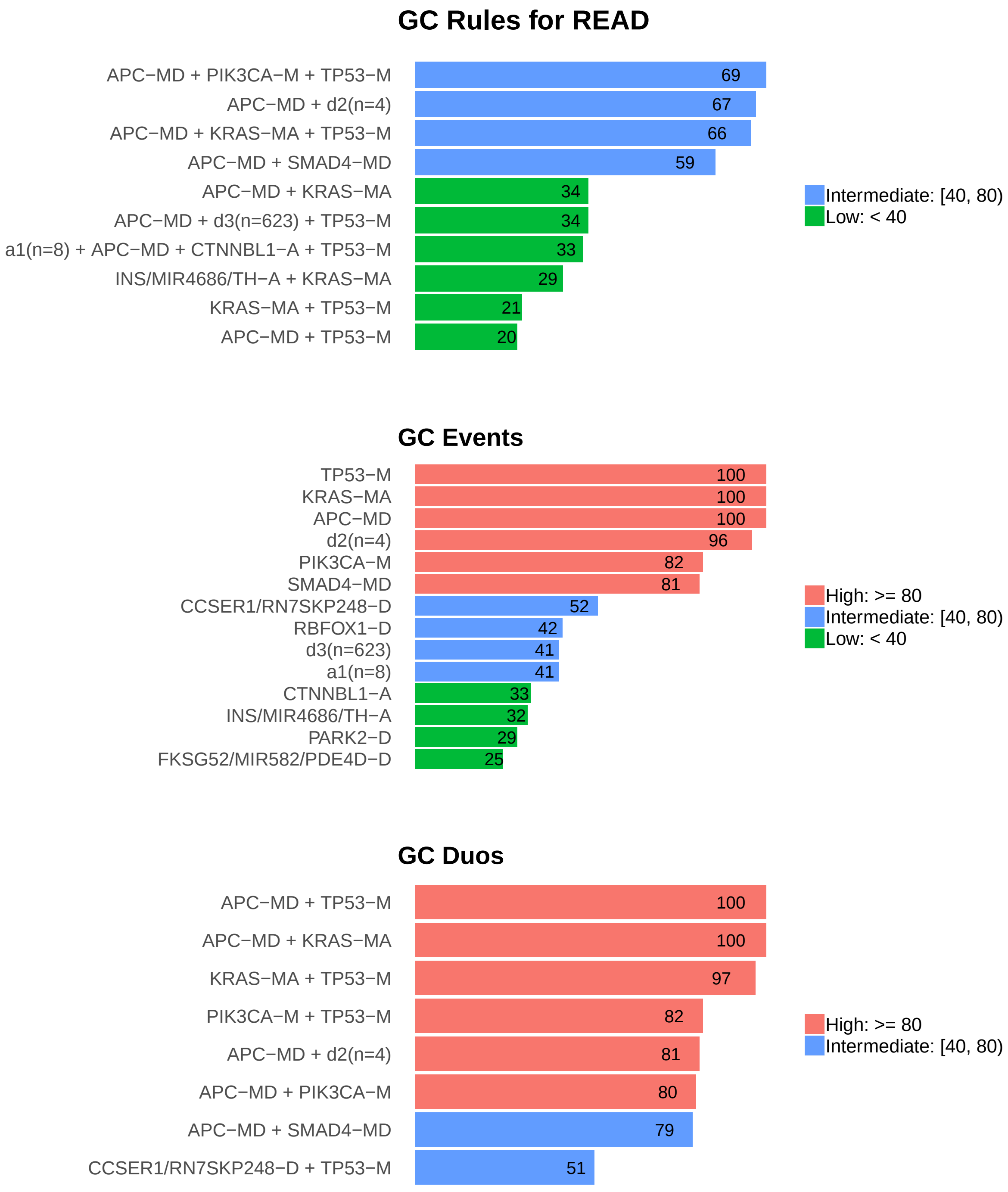

Figure 5: Summary of generalized core results. Bars show confidence levels, which are the percentage of sub-sample iterations containing the observation. Rules (top) and events (middle) that achieve a minimum confidence level of 20 are shown. Duos (bottom) with confidence of at least 50 are shown.

### 5 Dictionary of Copy Number Events

| CNV | Genes | Event_Name |
|---|---|---|
| a1 | HCK, PLAGL2, KIF3B, TM9SF4, POFUT1, TSPY26P, ASXL1, MIR1825 | a1(n=8) |
| a2 | CTNNBL1 | CTNNBL1-A |
| a3 | CASC8 | CASC8-A |
| a8 | INS, TH, MIR4686 | INS/MIR4686/TH-A |
| d1 | RBFOX1 | RBFOX1-D |
| d2 | RNA5SP475, RN7SL864P, FLRT3, MACROD2 | d2(n=4) |
| d3 | d3(n=623) | d3(n=623) |
| d4 | PARK2 | PARK2-D |
| d5 | RN7SKP248, CCSER1 | CCSER1/RN7SKP248-D |
| d6 | snoU13\|ENSG00000271842.1, MIR4789, RN7SKP40, NAALADL2 | d6(n=4) |
| d7 | FCN3 | FCN3-D |
| d8 | FKSG52, MIR582, PDE4D | FKSG52/MIR582/PDE4D-D |
| d9 | U3\|ENSG00000212211.1, NPCDR1, FHIT | FHIT/NPCDR1/U3-D |
| d10 | RN7SL872P, RNA5SP251, CSMD1 | CSMD1/RN7SL872P/RNA5SP251-D |
| d16 | snoU13\|ENSG00000238922.1, LRRN3, IMMP2L | IMMP2L/LRRN3/snoU13-D |

Dictionary of important CNVs. Copy number events that are within any of the best K rule sets, or within the generalized core rules, or among the 20 most frequent events.
