## Supplementary material for "Identifying Combinations of Cancer Drivers in Individual Patients": TCGA Reports: CRSO_Report_SKCM.pdf

### CRSO Output Report: SKCM

Michael Klein

Apr 19 2019

**4 Generalized Core Rules** **6**

**5 Dictionary of Copy Number Events** **7**

#### 1 Heatmaps of D and P

**Number of samples = 290.**
**Number of events = 71.**
**Rule coverage requirement = 9 samples.**
**Rule library size = 197 rules.**

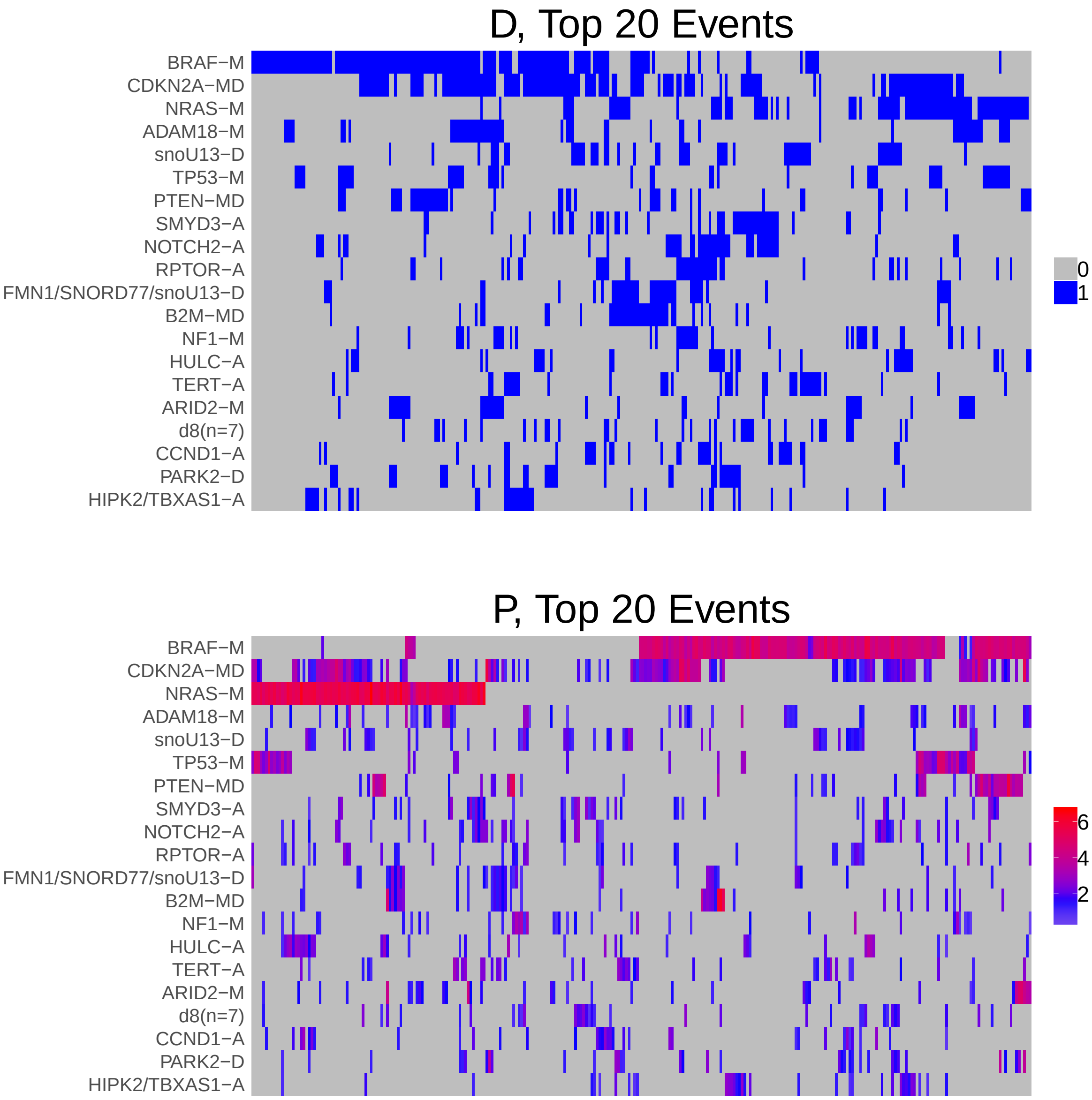

Figure 1: Heatmap of D and P. Events are ordered by frequency, from top to bottom. Event suffix -M: mutation, -A: amplification, -D: deletion, -MD: mutDel, -MA: mutAmp. Samples are ordered using hierarchical clustering. Wild-type events indicated in grey.

### 2 Summary of K Best Rule Sets

#### 2.1 Performance and coverage of best rule sets

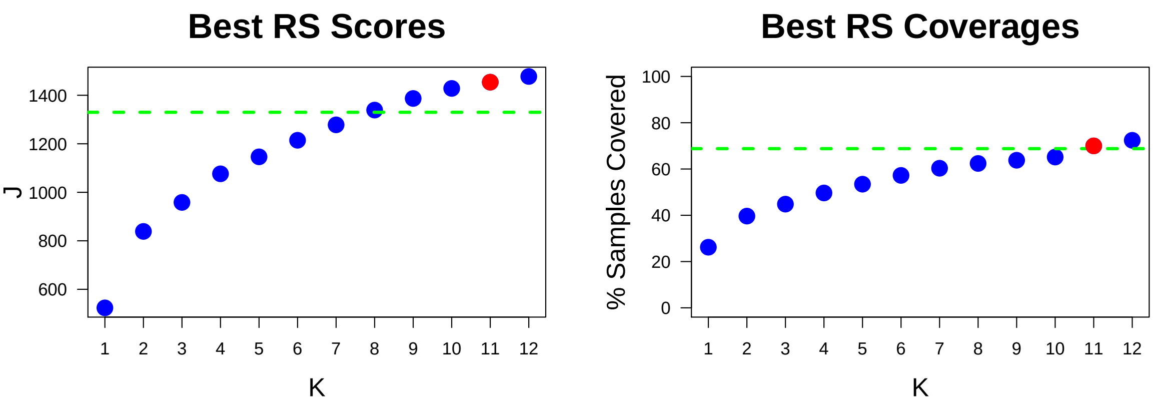

Figure 2: Performance and Coverage of Best Filtered RS for K. The core was determined to be the smallest RS that achieved at least 95% of maximum coverage and at least 90% of maximum performance.

#### 2.2 Table of rules that appear in any best rule set

| **ID** | **Rule** | **PC** | **Ks** |
|---|---|---|---|
| r1 | BRAF-M + CDKN2A-MD | 26 | 1,2,3,4,5,6,7,8,9,10,11,12 |
| r2 | CDKN2A-MD + NRAS-M | 14 | 2,3,4,5,6,7,8,9,10,11,12 |
| r4 | BRAF-M + PTEN-MD | 11 | 3,4,5,6,7,8,9,10,11,12 |
| r3 | NRAS-M + TP53-M | 6.6 | 4,5,6,7,8,9,10,11,12 |
| r5 | BRAF-M + HIPK2/TBXAS1-A | 7.9 | 5,6,7,8,9,10,11,12 |
| r12 | BRAF-M + RN7SKP254-A | 6.6 | 6,7,8,9,10,11,12 |
| r20 | ARID2-M + NRAS-M | 5.2 | 7,8,9,10 |
| r30 | KCNN3-A + NOTCH2-A + NRAS-M | 3.4 | 8,9,10,12 |
| r7 | BRAF-M + TP53-M | 8.3 | 10,11,12 |
| r6 | HULC-A + NRAS-M | 6.6 | 11,12 |
| r8 | ADAM18-M + NRAS-M | 6.9 | 11,12 |
| r17 | B2M-MD + FMN1/SNORD77/snoU13-D | 9 | 11,12 |
| r18 | B2M-MD + FMN1/SNORD77/snoU13-D + NRAS-M | 3.4 | 9,10 |
| r16 | NRAS-M + SMYD3-A | 5.5 | 11 |
| r112 | d8(n=7) + SMYD3-A | 4.8 | 12 |

**ID** = Rule IDs, rules are numbered according to importance rank determined from phase 1
**PC** = Percent of samples covered **Ks** = Membership in best RS

### 3 Core Rule Set

**Core K = 11.**
**Core rule set coverage = 70%.**

#### 3.1 Table of core rule set rules

| ID | Rule | CR | SJR | SJ | NSC | NSA | PC | PA | FracA |
|---|---|---|---|---|---|---|---|---|---|
| r1 | BRAF-M + CDKN2A-MD | 1 | 1 | 524 | 76 | 62 | 26 | 21.0 | 0.82 |
| r2 | CDKN2A-MD + NRAS-M | 2 | 2 | 327 | 41 | 28 | 14 | 9.7 | 0.68 |
| r3 | NRAS-M + TP53-M | 29 | 8 | 156 | 19 | 17 | 6.6 | 5.9 | 0.89 |
| r4 | BRAF-M + PTEN-MD | 3.5 | 3 | 227 | 31 | 18 | 11 | 6.2 | 0.58 |
| r5 | BRAF-M + HIPK2/TBXAS1-A | 11.5 | 10 | 144 | 23 | 10 | 7.9 | 3.4 | 0.43 |
| r6 | HULC-A + NRAS-M | 29 | 9 | 146 | 19 | 11 | 6.6 | 3.8 | 0.58 |
| r7 | BRAF-M + TP53-M | 9 | 6.5 | 161 | 24 | 13 | 8.3 | 4.5 | 0.54 |
| r8 | ADAM18-M + NRAS-M | 22.5 | 11 | 137 | 20 | 9 | 6.9 | 3.1 | 0.45 |
| r12 | BRAF-M + RN7SKP254-A | 29 | 21 | 115 | 19 | 11 | 6.6 | 3.8 | 0.58 |
| r16 | NRAS-M + SMYD3-A | 42.5 | 20 | 119 | 16 | 10 | 5.5 | 3.4 | 0.62 |
| r17 | B2M-MD + FMN1/SNORD77/snoU13-D | 6.5 | 36 | 96.4 | 26 | 14 | 9 | 4.8 | 0.54 |

**ID** = Rule IDs, rules are numbered according to importance rank determined from phase 1
**CR** = Coverage rank
**SJ/SJR** = Single rule performance / performance rank
**NSC/PC** = Number/percent of samples covered
**NSA/PA** = Number/percent of samples assigned
**FracA** = Fraction of covered samples assigned to rule

#### 3.2 Event breakdown of core rule set

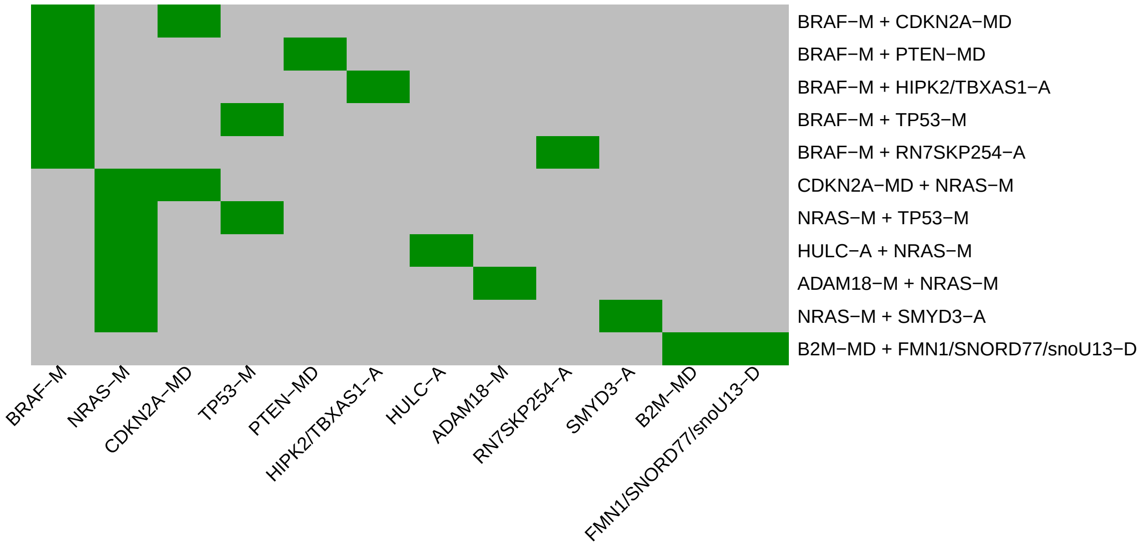

Figure 3: Visualization of core rule set. Rows are rules, columns are events. Events are ordered from left in decreasing rule membership frequency.

##### 3.3 Core Rule Set Penalties: Before and After Assignment

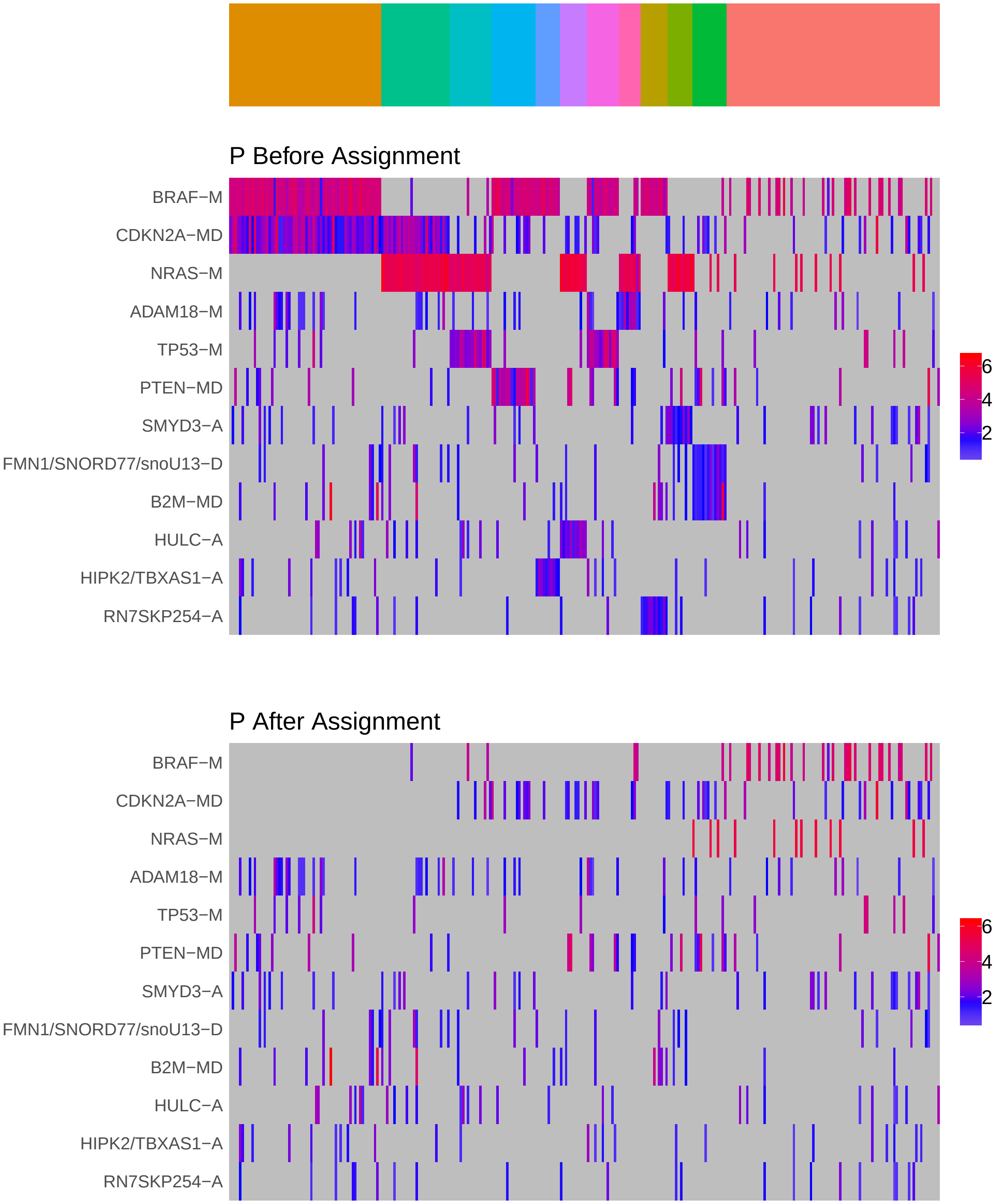

Figure 4: Heatmap of P before and after assignment to core rule set. Events are force ordered by frequency. Samples are force ordered according rule set membership, as indicated by the color bar. The right-most group of samples are not assigned to any rule.

### 4 Generalized Core Rules

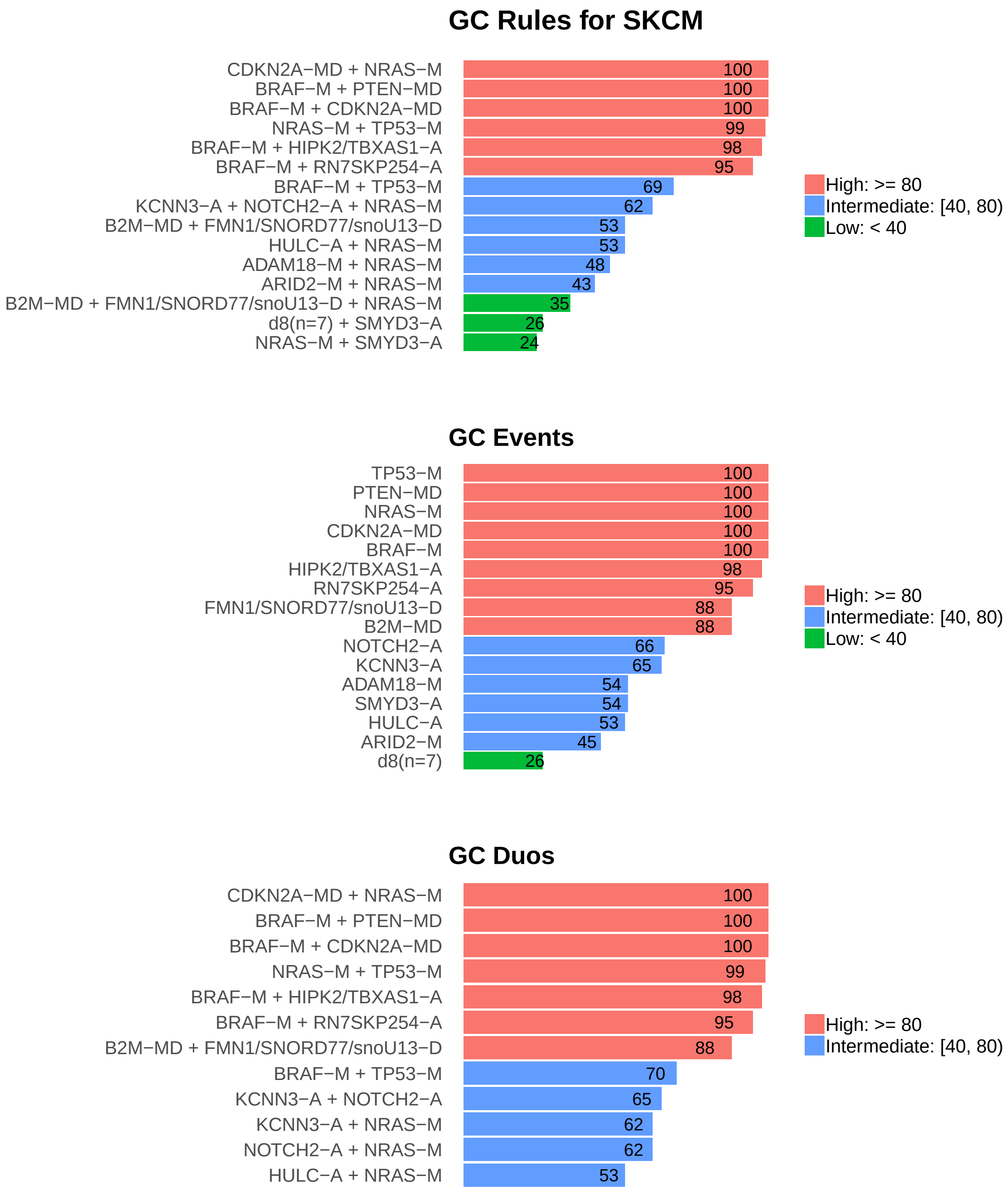

Figure 5: Summary of generalized core results. Bars show confidence levels, which are the percentage of sub-sample iterations containing the observation. Rules (top) and events (middle) that achieve a minimum confidence level of 20 are shown. Duos (bottom) with confidence of at least 50 are shown.

### 5 Dictionary of Copy Number Events

| CNV | Genes | Event_Name |
|---|---|---|
| a1 | SMYD3 | SMYD3-A |
| a2 | RPTOR | RPTOR-A |
| a3 | NOTCH2 | NOTCH2-A |
| a4 | HULC | HULC-A |
| a5 | TBXAS1, HIPK2 | HIPK2/TBXAS1-A |
| a6 | CCND1 | CCND1-A |
| a7 | MITF | MITF-A |
| a8 | TERT | TERT-A |
| a9 | KCNN3 | KCNN3-A |
| a10 | RN7SKP254 | RN7SKP254-A |
| d2 | snoU13\|ENSG00000239153.1 | snoU13-D |
| d3 | PARK2 | PARK2-D |
| d7 | SNORD77\|ENSG00000212415.1, snoU13\|ENSG00000238342.1, FMN1 | FMN1/SNORD77/snoU13-D |
| d8 | RPL22, KCNAB2, CHD5, NPHP4, RNF207, MIR4689, MIR4417 | d8(n=7) |

Dictionary of important CNVs. Copy number events that are within any of the best K rule sets, or within the generalized core rules, or among the 20 most frequent events.
