## Supplementary material for "Identifying Combinations of Cancer Drivers in Individual Patients": TCGA Reports: CRSO_Report_STAD.pdf

### CRSO Output Report: STAD

Michael Klein

Apr 19 2019

**4 Generalized Core Rules** **8**

**5 Dictionary of Copy Number Events** **9**

#### 1 Heatmaps of D and P

**Number of samples = 391.**
**Number of events = 121.**
**Rule coverage requirement = 12 samples.**
**Rule library size = 850 rules.**

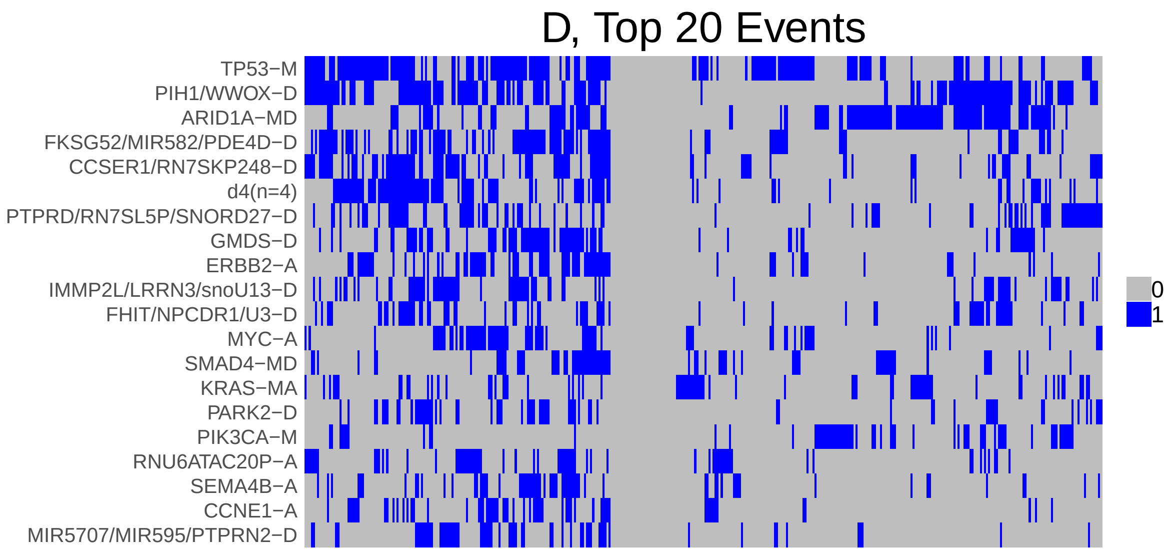

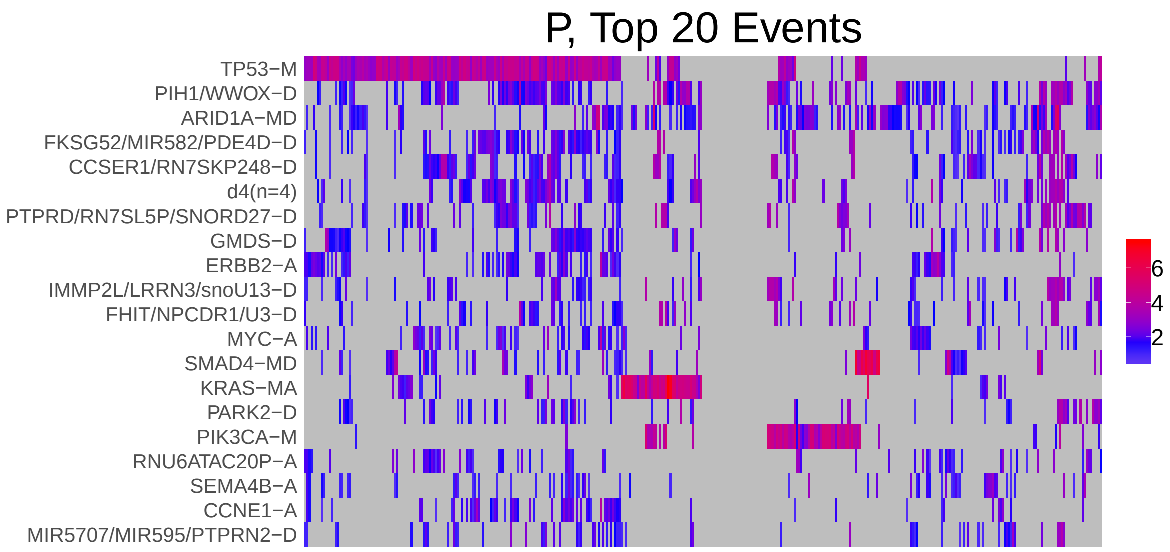

Figure 1: Heatmap of D and P. Events are ordered by frequency, from top to bottom. Event suffix -M: mutation, -A: amplification, -D: deletion, -MD: mutDel, -MA: mutAmp. Samples are ordered using hierarchical clustering. Wild-type events indicated in grey.

#### 2 Summary of K Best Rule Sets

##### 2.1 Performance and coverage of best rule sets

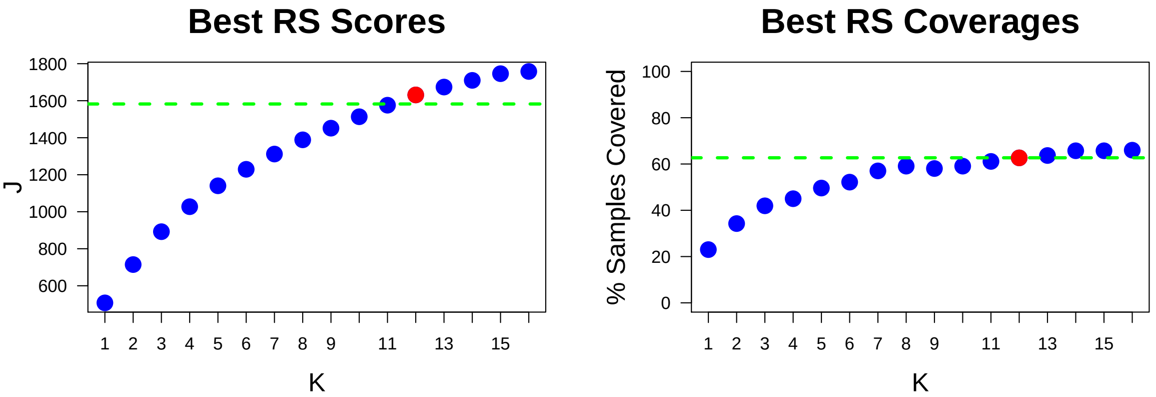

Figure 2: Performance and Coverage of Best Filtered RS for K. The core was determined to be the smallest RS that achieved at least 95% of maximum coverage and at least 90% of maximum performance.

##### 2.2 Table of rules that appear in any best rule set

| ID | Rule | PC | Ks |
|---|---|---|---|
| r1 | ARID1A-MD + PIK3CA-M | 10 | 5,6,7,8,9,10,11,12,13,14,15,16 |
| r4 | CCSER1/RN7SKP248-D + PIH1/WWOX-D + TP53-M | 10 | 5,6,7,8,9,10,11,12,13,14,15,16 |
| r2 | ARID1A-MD + PIH1/WWOX-D | 17 | 2,3,4,5,6,7,8,9,10,11,12 |
| r9 | d4(n=4) + TP53-M | 18 | 3,5,6,7,8,11,12,13,14,15,16 |
| r8 | SMAD4-MD + TP53-M | 12 | 6,8,9,10,11,12,13,14,15,16 |
| r5 | ARID1A-MD + TP53-M | 14 | 4,7,8,11,12,13,14,15,16 |
| r7 | KRAS-MA + TP53-M | 8.2 | 9,10,11,12,13,14,15,16 |
| r10 | PIH1/WWOX-D + PTPRD/RN7SL5P/SNORD27-D | 14 | 7,8,13,14,15,16 |
| r3 | ARID1A-MD + KRAS-MA | 8.2 | 12,13,14,15,16 |
| r15 | ERBB2-A + TP53-M | 16 | 9,10,11,12,13 |
| r6 | PIH1/WWOX-D + TP53-M | 23 | 1,2,3,4 |
| r13 | ARID1A-MD + FHIT/NPCDR1/U3-D + PIH1/WWOX-D | 7.4 | 13,14,15,16 |
| r16 | GMDS-D + TP53-M | 15 | 5,6,7,8 |
| r19 | ARID1A-MD + SMAD4-MD | 7.9 | 13,14,15,16 |
| r20 | PIH1/WWOX-D + PIK3CA-M | 7.4 | 9,10,11,12 |
| r18 | MYC-A + TP53-M | 12 | 14,15,16 |
| r84 | ERBB2-A + GMDS-D + TP53-M | 7.2 | 14,15,16 |
| r86 | d4(n=4) + FKSG52/MIR582/PDE4D-D + IMMP2L/LRRN3/snoU13-D + PIH1/WWOX-D | 4.3 | 10,12,13 |
| r89 | ERBB2-A + FKSG52/MIR582/PDE4D-D + PIH1/WWOX-D + TP53-M | 5.1 | 14,15,16 |
| r101 | CCNE1-A + FKSG52/MIR582/PDE4D-D + GMDS-D + TP53-M | 4.3 | 11,12,13 |
| r11 | FKSG52/MIR582/PDE4D-D + TP53-M | 18 | 9,10 |
| r12 | IMMP2L/LRRN3/snoU13-D + PIH1/WWOX-D | 14 | 14,15 |
| r70 | d4(n=4) + PTPRD/RN7SL5P/SNORD27-D + TP53-M | 6.6 | 9,10 |
| r166 | CCNE1-A + CCSER1/RN7SKP248-D + FKSG52/MIR582/PDE4D-D + TP53-M | 4.3 | 15,16 |
| r17 | CCSER1/RN7SKP248-D + d4(n=4) + TP53-M | 9.7 | 4 |
| r22 | d4(n=4) + IMMP2L/LRRN3/snoU13-D + PIH1/WWOX-D | 7.2 | 11 |
| r38 | PIK3CA-M + PTPRD/RN7SL5P/SNORD27-D | 4.1 | 16 |
| r123 | FKSG52/MIR582/PDE4D-D + IMMP2L/LRRN3/snoU13-D | 9.7 | 16 |

**ID** = Rule IDs, rules are numbered according to importance rank determined from phase 1
**PC** = Percent of samples covered **Ks** = Membership in best RS

### 3 Core Rule Set

**Core K = 12.**
**Core rule set coverage = 62.7%.**

#### 3.1 Table of core rule set rules

| ID | Rule | CR | SJR | SJ | NSC | NSA | PC | PA | FracA |
|---|---|---|---|---|---|---|---|---|---|
| r1 | ARID1A-MD + PIK3CA-M | 38 | 33 | 220 | 39 | 15 | 10 | 3.8 | 0.38 |
| r2 | ARID1A-MD + PIH1/WWOX-D | 5 | 10 | 296 | 67 | 25 | 17 | 6.4 | 0.37 |
| r3 | ARID1A-MD + KRAS-MA | 77.5 | 71.5 | 176 | 32 | 19 | 8.2 | 4.9 | 0.59 |
| r4 | CCSER1/RN7SKP248-D + PIH1/WWOX-D + TP53-M | 30.5 | 7 | 315 | 41 | 30 | 10 | 7.7 | 0.73 |
| r5 | ARID1A-MD + TP53-M | 13.5 | 13 | 287 | 54 | 20 | 14 | 5.1 | 0.37 |
| r7 | KRAS-MA + TP53-M | 77.5 | 39 | 211 | 32 | 19 | 8.2 | 4.9 | 0.59 |
| r8 | SMAD4-MD + TP53-M | 23 | 17 | 275 | 45 | 20 | 12 | 5.1 | 0.44 |
| r9 | d4(n=4) + TP53-M | 3 | 3 | 399 | 70 | 27 | 18 | 6.9 | 0.39 |
| r15 | ERBB2-A + TP53-M | 8 | 5 | 340 | 61 | 22 | 16 | 5.6 | 0.36 |
| r20 | PIH1/WWOX-D + PIK3CA-M | 105 | 55 | 196 | 29 | 19 | 7.4 | 4.9 | 0.66 |
| r86 | d4(n=4) + FKSG52/MIR582/PDE4D-D + IMMP2L/LRRN3/snoU13-D + PIH1/WWOX-D | 549.5 | 123.5 | 153 | 17 | 12 | 4.3 | 3.1 | 0.71 |
| r101 | CCNE1-A + FKSG52/MIR582/PDE4D-D + GMDS-D + TP53-M | 549.5 | 108 | 157 | 17 | 17 | 4.3 | 4.3 | 1.00 |

**ID** = Rule IDs, rules are numbered according to importance rank determined from phase 1
**CR** = Coverage rank
**SJ/SJR** = Single rule performance / performance rank
**NSC/PC** = Number/percent of samples covered
**NSA/PA** = Number/percent of samples assigned
**FracA** = Fraction of covered samples assigned to rule

#### 3.2 Event breakdown of core rule set

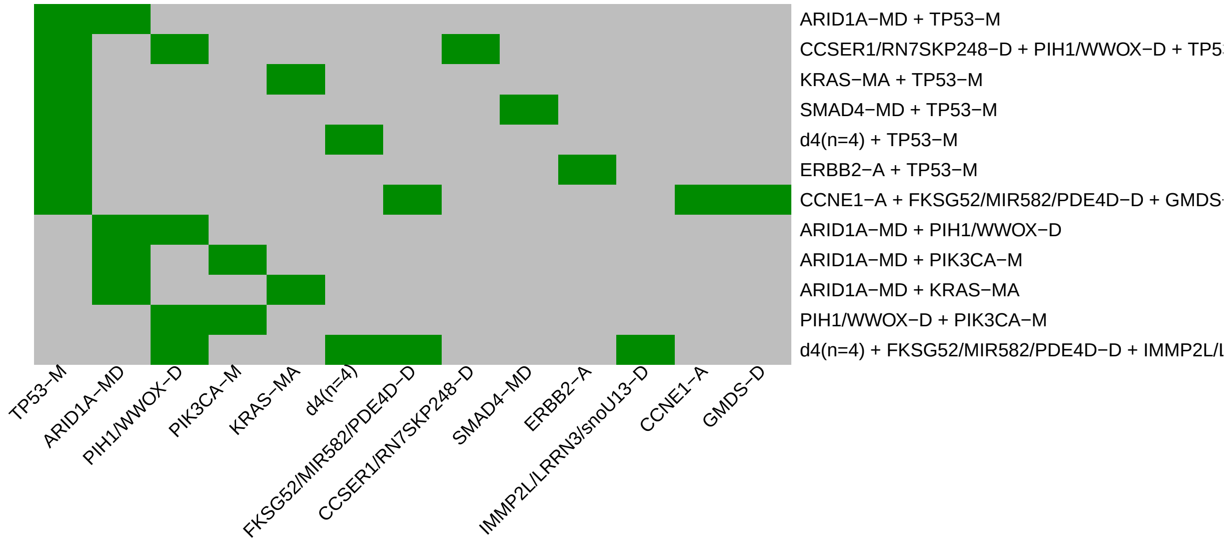

Figure 3: Visualization of core rule set. Rows are rules, columns are events. Events are ordered from left in decreasing rule membership frequency.

##### 3.3 Core Rule Set Penalties: Before and After Assignment

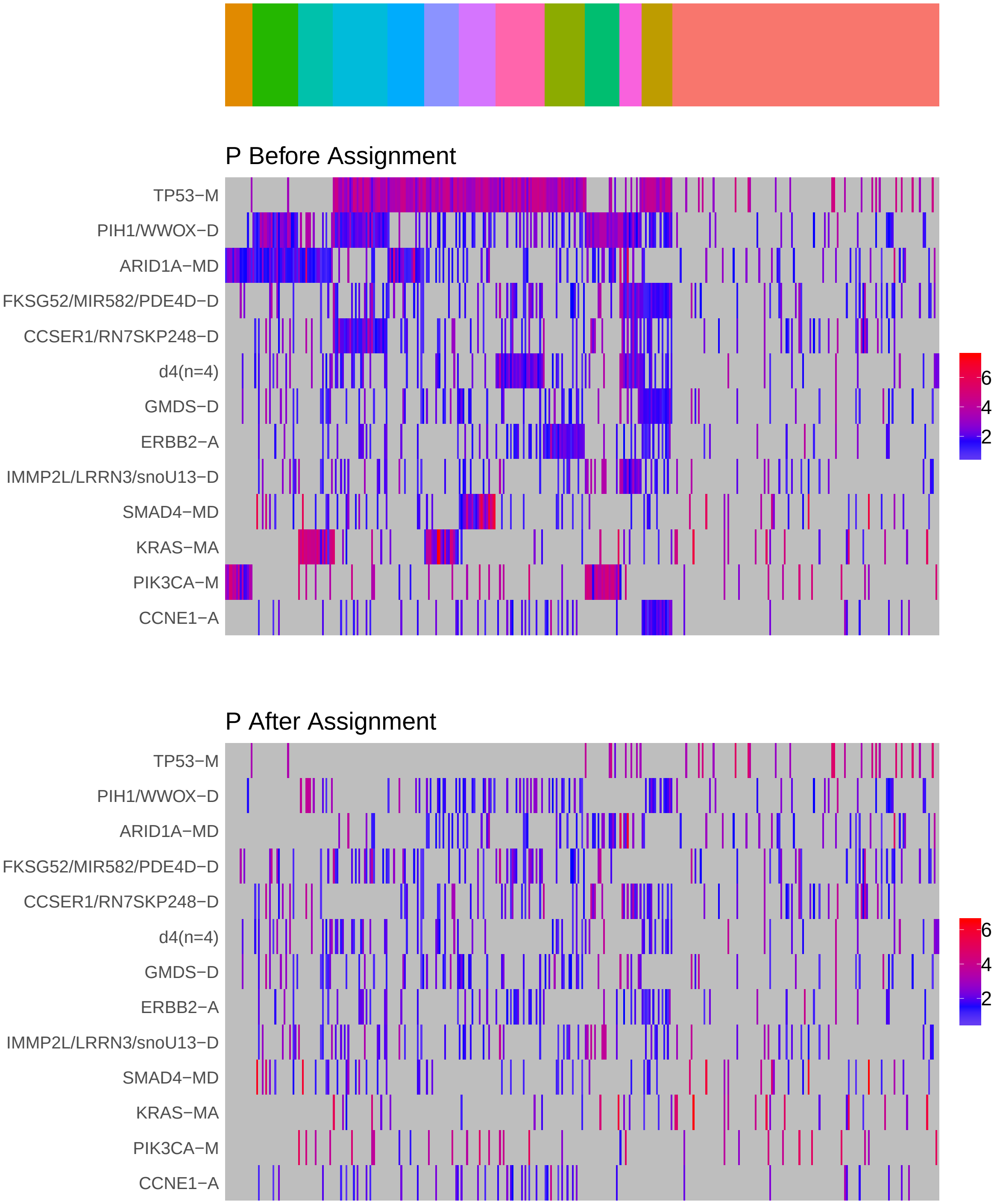

Figure 4: Heatmap of P before and after assignment to core rule set. Events are force ordered by frequency. Samples are force ordered according rule set membership, as indicated by the color bar. The right-most group of samples are not assigned to any rule.

### 4 Generalized Core Rules

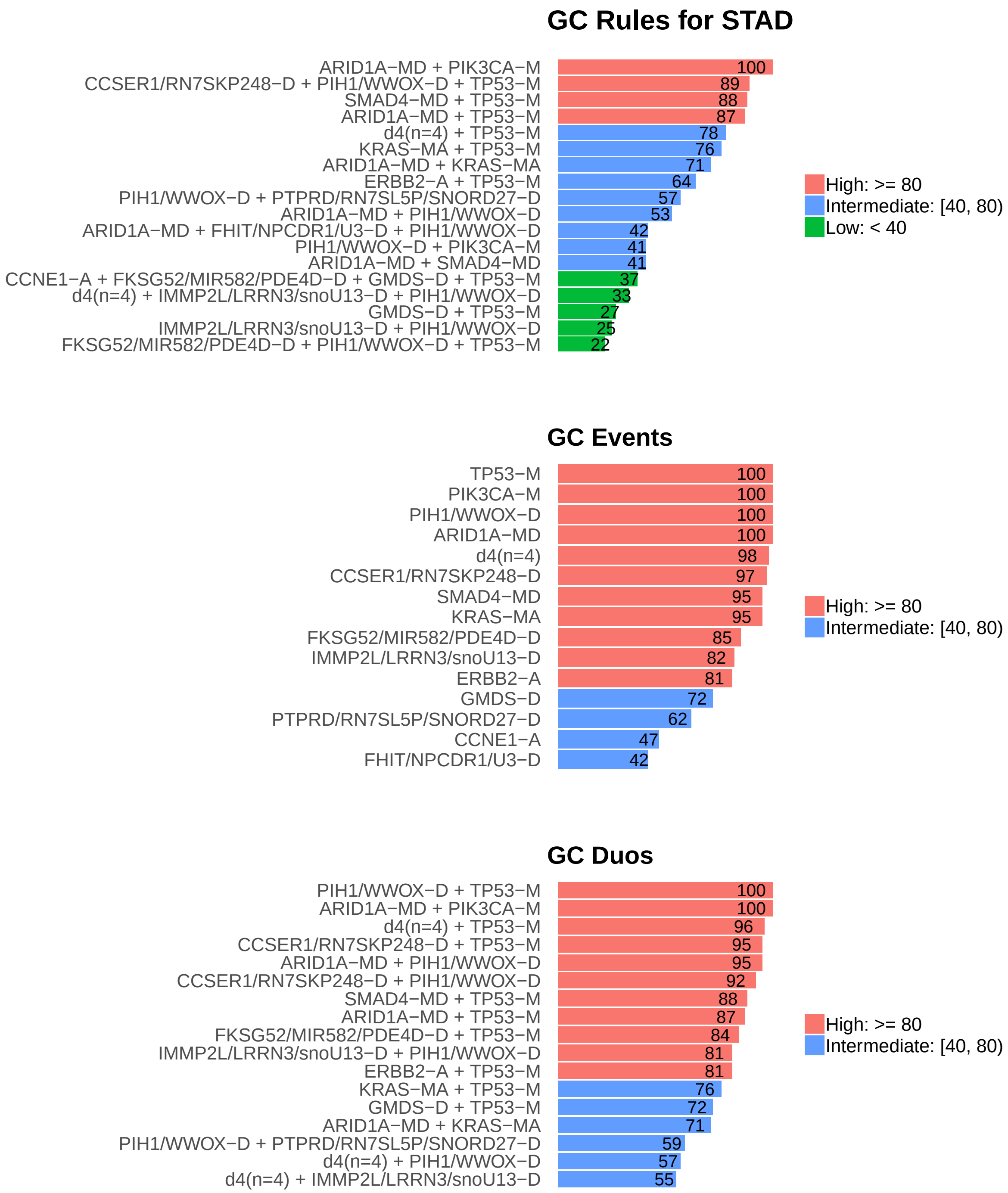

Figure 5: Summary of generalized core results. Bars show confidence levels, which are the percentage of sub-sample iterations containing the observation. Rules (top) and events (middle) that achieve a minimum confidence level of 20 are shown. Duos (bottom) with confidence of at least 50 are shown.

### 5 Dictionary of Copy Number Events

| CNV | Genes | Event_Name |
|---|---|---|
| a1 | ERBB2 | ERBB2-A |
| a2 | MYC | MYC-A |
| a3 | RNU6ATAC20P | RNU6ATAC20P-A |
| a4 | SEMA4B | SEMA4B-A |
| a6 | CCNE1 | CCNE1-A |
| d1 | PIH1, WWOX | PIH1/WWOX-D |
| d2 | FKSG52, MIR582, PDE4D | FKSG52/MIR582/PDE4D-D |
| d3 | RN7SKP248, CCSER1 | CCSER1/RN7SKP248-D |
| d4 | RNA5SP475, RN7SL864P, FLRT3, MACROD2 | d4(n=4) |
| d5 | RN7SL5P, SNORD27\|ENSG00000251699.1, PTPRD | PTPRD/RN7SL5P/SNORD27-D |
| d6 | GMDS | GMDS-D |
| d7 | snoU13\|ENSG00000238922.1, LRRN3, IMMP2L | IMMP2L/LRRN3/snoU13-D |
| d8 | U3\|ENSG00000212211.1, NPCDR1, FHIT | FHIT/NPCDR1/U3-D |
| d9 | PARK2 | PARK2-D |
| d10 | MIR595, PTPRN2, MIR5707 | MIR5707/MIR595/PTPRN2-D |

Dictionary of important CNVs. Copy number events that are within any of the best K rule sets, or within the generalized core rules, or among the 20 most frequent events.
