## Supplementary material for "Identifying Combinations of Cancer Drivers in Individual Patients": TCGA Reports: CRSO_Report_UCEC.pdf

### CRSO Output Report: UCEC

Michael Klein

Apr 19 2019

**4 Generalized Core Rules** **6**

**5 Dictionary of Copy Number Events** **7**

#### 1 Heatmaps of D and P

**Number of samples = 242.**
**Number of events = 136.**
**Rule coverage requirement = 8 samples.**
**Rule library size = 930 rules.**

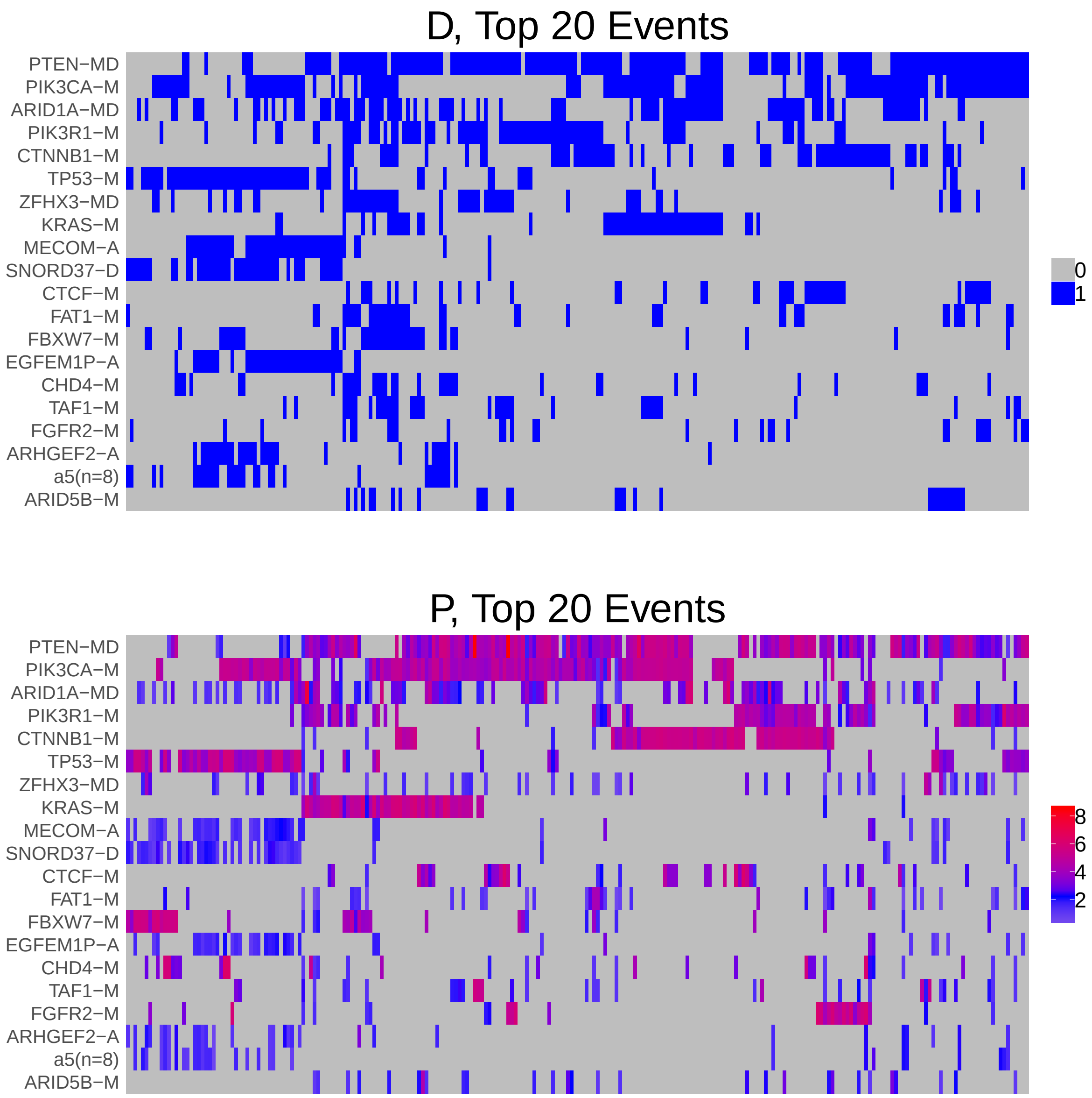

Figure 1: Heatmap of D and P. Events are ordered by frequency, from top to bottom. Event suffix -M: mutation, -A: amplification, -D: deletion, -MD: mutDel, -MA: mutAmp. Samples are ordered using hierarchical clustering. Wild-type events indicated in grey.

### 2 Summary of K Best Rule Sets

#### 2.1 Performance and coverage of best rule sets

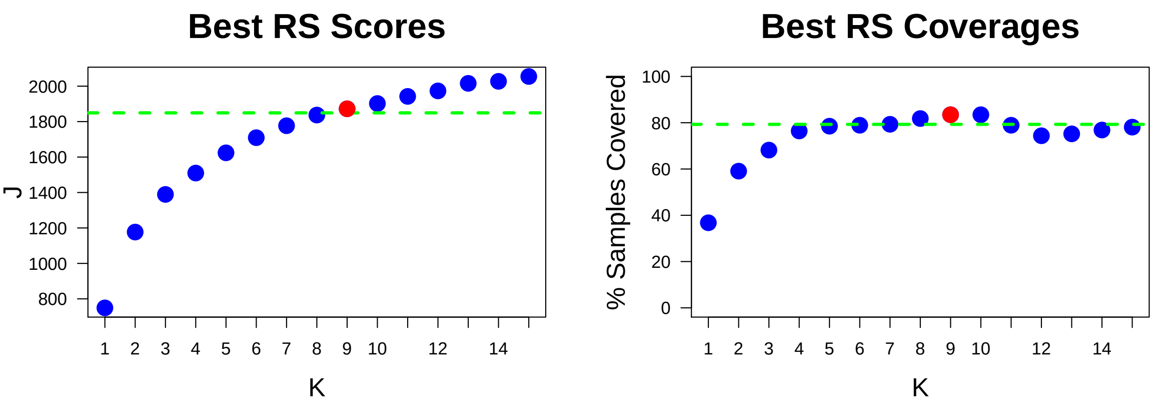

Figure 2: Performance and Coverage of Best Filtered RS for K. The core was determined to be the smallest RS that achieved at least 95% of maximum coverage and at least 90% of maximum performance.

#### 2.2 Table of rules that appear in any best rule set

| **ID** | **Rule** | **PC** | **Ks** |
|---|---|---|---|
| r4 | SNORD37-D + TP53-M | 16 | 4,5,6,7,8,9,10,11,12,13,14,15 |
| r1 | PIK3R1-M + PTEN-MD | 29 | 2,3,4,5,6,7,8,9,10,11 |
| r2 | PIK3CA-M + PTEN-MD | 37 | 1,2,3,4,5,6,7,8,9,10 |
| r7 | EGFEM1P-A + MECOM-A + PIK3CA-M + TP53-M | 7 | 6,7,8,9,10,11,12,13 |
| r33 | ARID1A-MD + KRAS-M + PIK3CA-M | 9.9 | 5,7,8,9,10,11 |
| r3 | PIK3CA-M + TP53-M | 13 | 3,4,5,14,15 |
| r6 | CTNNB1-M + PIK3CA-M + PTEN-MD | 13 | 11,12,13,14,15 |
| r9 | CTNNB1-M + PIK3CA-M | 17 | 6,7,8,9,10 |
| r10 | CTCF-M + PIK3CA-M + PTEN-MD | 9.5 | 11,12,13,14,15 |
| r15 | FGFR2-M + PTEN-MD | 9.1 | 11,12,13,14,15 |
| r5 | CTNNB1-M + PIK3R1-M + PTEN-MD | 11 | 12,13,14,15 |
| r11 | FBXW7-M + TP53-M | 5.8 | 11,13,14,15 |
| r24 | PIK3R1-M + PTEN-MD + ZFHX3-MD | 9.9 | 12,13,14,15 |
| r25 | PTEN-MD + TP53-M | 11 | 12,13,14,15 |
| r32 | FBXW7-M + PTEN-MD | 9.5 | 12,13,14,15 |
| r43 | ARID1A-MD + KRAS-M + PIK3R1-M + PTEN-MD | 4.5 | 12,13,14,15 |
| r45 | ARID1A-MD + CTNNB1-M + PTEN-MD | 9.9 | 7,8,9,10 |
| r16 | ARID1A-MD + PIK3CA-M + PTEN-MD | 16 | 13,14,15 |
| r20 | KRAS-M + PIK3CA-M | 15 | 13,14,15 |
| r62 | CHD4-M + FBXW7-M | 6.6 | 8,9,10 |
| r17 | KRAS-M + PIK3CA-M + PTEN-MD | 11 | 11,12 |
| r38 | PPP2R1A-M + TP53-M | 5.4 | 10,14 |
| r14 | ARID1A-MD + PTEN-MD | 28 | 11 |
| r18 | ARID1A-MD + PIK3CA-M | 24 | 12 |
| r34 | ARID1A-MD + KRAS-M + PTEN-MD | 11 | 6 |
| r36 | ARID1A-MD + CTNNB1-M | 13 | 11 |
| r57 | FAT1-M + PIK3CA-M + PTEN-MD | 8.7 | 15 |
| r61 | MYC-A + RN7SKP226-A + TP53-M | 5.8 | 10 |
| r107 | a5(n=8) + CCNE1-A + TP53-M | 5.8 | 9 |
| r131 | MECOM-A + MYC-A + RN7SKP226-A + TP53-M | 4.5 | 15 |

**ID** = Rule IDs, rules are numbered according to importance rank determined from phase 1
**PC** = Percent of samples covered **Ks** = Membership in best RS

### 3 Core Rule Set

**Core K = 9.**
**Core rule set coverage = 83.5%.**

#### 3.1 Table of core rule set rules

| ID | Rule | CR | SJR | SJ | NSC | NSA | PC | PA | FracA |
|---|---|---|---|---|---|---|---|---|---|
| r1 | PIK3R1-M + PTEN-MD | 2 | 2 | 534 | 70 | 49 | 29 | 20.0 | 0.70 |
| r2 | PIK3CA-M + PTEN-MD | 1 | 1 | 749 | 89 | 39 | 37 | 16.0 | 0.44 |
| r4 | SNORD37-D + TP53-M | 10.5 | 25 | 229 | 38 | 17 | 16 | 7.0 | 0.45 |
| r7 | EGFEM1P-A + MECOM-A + PIK3CA-M + TP53-M | 115 | 30 | 212 | 17 | 17 | 7 | 7.0 | 1.00 |
| r9 | CTNNB1-M + PIK3CA-M | 7.5 | 8 | 366 | 40 | 22 | 17 | 9.1 | 0.55 |
| r33 | ARID1A-MD + KRAS-M + PIK3CA-M | 42.5 | 17 | 277 | 24 | 23 | 9.9 | 9.5 | 0.96 |
| r45 | ARID1A-MD + CTNNB1-M + PTEN-MD | 42.5 | 17 | 277 | 24 | 19 | 9.9 | 7.9 | 0.79 |
| r62 | CHD4-M + FBXW7-M | 143 | 244.5 | 100 | 16 | 8 | 6.6 | 3.3 | 0.50 |
| r107 | a5(n=8) + CCNE1-A + TP53-M | 229 | 255 | 98.9 | 14 | 8 | 5.8 | 3.3 | 0.57 |

**ID** = Rule IDs, rules are numbered according to importance rank determined from phase 1
**CR** = Coverage rank
**SJ/SJR** = Single rule performance / performance rank
**NSC/PC** = Number/percent of samples covered
**NSA/PA** = Number/percent of samples assigned
**FracA** = Fraction of covered samples assigned to rule

#### 3.2 Event breakdown of core rule set

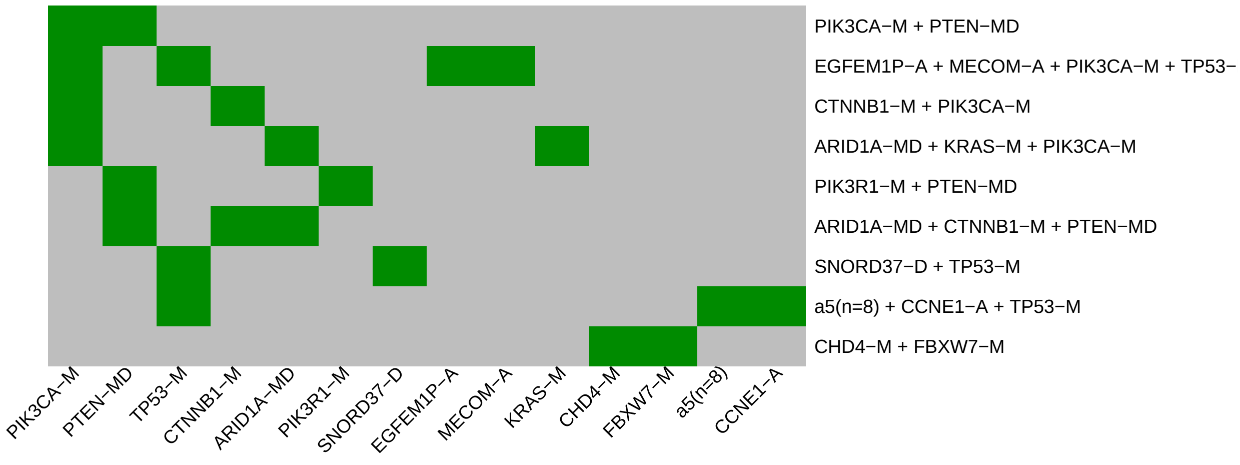

Figure 3: Visualization of core rule set. Rows are rules, columns are events. Events are ordered from left in decreasing rule membership frequency.

#### 3.3 Core Rule Set Penalties: Before and After Assignment

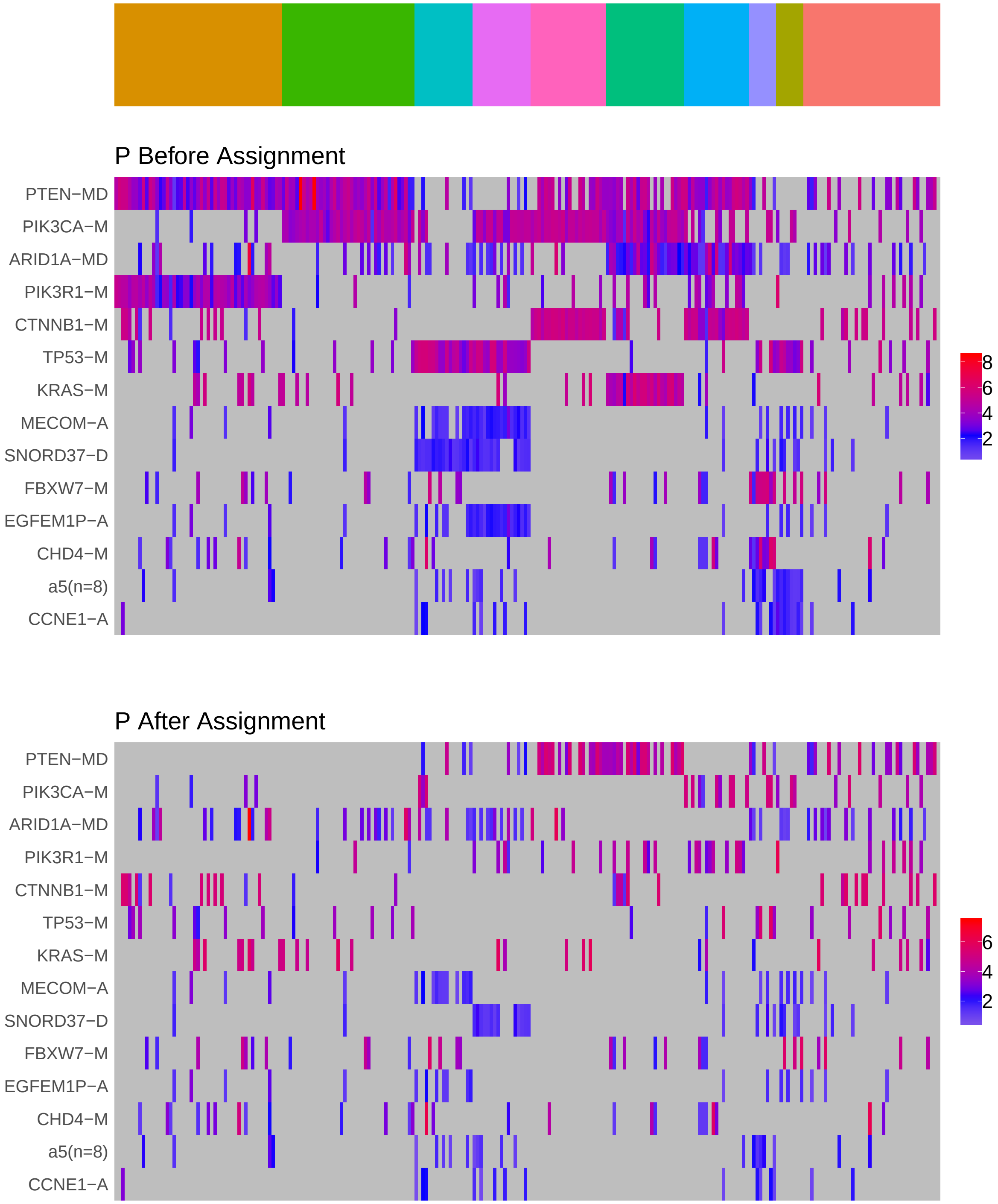

Figure 4: Heatmap of P before and after assignment to core rule set. Events are force ordered by frequency. Samples are force ordered according rule set membership, as indicated by the color bar. The right-most group of samples are not assigned to any rule.

### 4 Generalized Core Rules

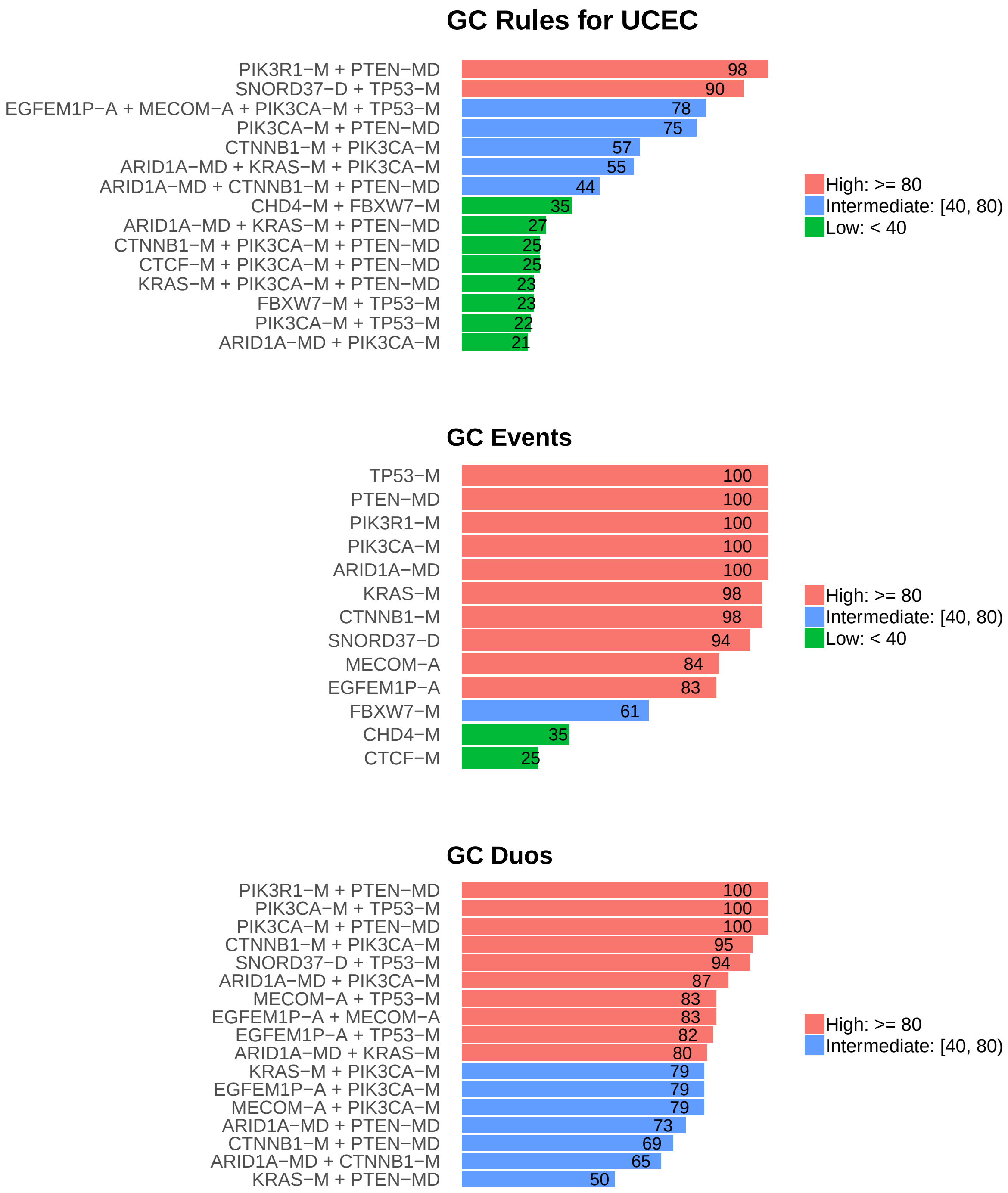

Figure 5: Summary of generalized core results. Bars show confidence levels, which are the percentage of sub-sample iterations containing the observation. Rules (top) and events (middle) that achieve a minimum confidence level of 20 are shown. Duos (bottom) with confidence of at least 50 are shown.

### 5 Dictionary of Copy Number Events

| CNV | Genes | Event_Name |
|---|---|---|
| a1 | MECOM | MECOM-A |
| a2 | EGFEM1P | EGFEM1P-A |
| a3 | ARHGEF2 | ARHGEF2-A |
| a5 | SNORA40\|ENSG00000253047.1, RN7SL600P, RN7SL473P, C1orf138, ENSA, MCL1, ADAMTSL4, MIR4257 | a5(n=8) |
| a6 | MYC | MYC-A |
| a7 | CCNE1 | CCNE1-A |
| a8 | RN7SKP226 | RN7SKP226-A |
| a11 | DNM2 | DNM2-A |
| d1 | SNORD37\|ENSG00000206775.1 | SNORD37-D |

Dictionary of important CNVs. Copy number events that are within any of the best K rule sets, or within the generalized core rules, or among the 20 most frequent events.
